# Prioritizing 2nd order interactions via support vector ranking using sensitivity indices on time series Wnt measurements

**DOI:** 10.1101/060228

**Authors:** shriprakash sinha

## Abstract

It is widely known that the sensitivity analysis plays a major role in computing the strength of the influence of involved factors in any phenomena under investigation. When applied to expression profiles of various intra/extracellular factors that form an integral part of a signaling pathway, the variance and density based analysis yields a range of sensitivity indices for individual as well as various combinations of factors. These combinations denote the higher order interactions among the involved factors, that might be of interest in the working mechanism of the pathway. For example, there are 19 types of WNTs and 10 FZDs with their 2^*nd*^ order combinations high enough and it is not possible to know which one to test first (except for those for which wet lab validations have been confirmed). But the effect of these combinations vary over time as measurements of fold changes and deviations in fold changes vary. In this work, after estimating the individual effects of factors for a higher order combination, the individual indices are considered as discriminative features. A combination, then is a multivariate feature set in higher order (>=2). With an excessively large number of factors involved in the pathway, it is difficult to search for important combinations in a wide search space over different orders. Exploiting the analogy of prioritizing webpages using ranking algorithms, for a particular order, a full set of combinations of interactions can then be prioritized based on these features using a powerful ranking algorithm via support vectors. Recording the changing rankings of the combinations over time points and durations, reveals how higher order interactions behave within the pathway and when and where an intervention might be necessary to influence the pathway. This could lead to development of time based therapeutic interventions. Based on a small dataset in time, we were able to generate the rankings of the 2^*nd*^ order combinations between WNTs and FZDs at different time snap shots and for different duration or time periods. Code has been made available on Google drive at https://drive.google.com/folderview?id=0B7Kkv8wlhPU-V1Fkd1dMSTd5ak0&usp=sharing

**Significance:** The search and wet lab testing of unknown biological hypotheses in the form of combinations of various intra/extracellular factors that are involved in a signaling pathway, costs a lot in terms of time, investment and energy. To reduce this cost of search in a vast combinatorial space, a pipeline has been developed that prioritises these list of combinations so that a biologist can narrow down their investigation. The pipeline uses kernel based sensitivity indices to capture the influence of the factors in a pathway and employs powerful support vector ranking algorithm. The generic workflow and future improvements are bound to cut down the cost for many wet lab experiments and reveal unknown/untested biological hypothesis.

## 1 Introduction

### 1.1 A short review

Sharma^1^’s accidental discovery of the Wingless played a pioneering role in the emergence of a widely expanding research field of the Wnt signaling pathway. A majority of the work has focused on issues related to • the discovery of genetic and epigenetic factors affecting the pathway (Thorstensen *et al*^2^. & Baron and Kneissel^3^), • implications of mutations in the pathway and its dominant role on cancer and other diseases (Clevers^4^), • investigation into the pathway’s contribution towards embryo development (Sokol^5^), homeostasis (Pinto *et al*.^6^, Zhong *et al*.^7^) and apoptosis (Pećina-Šlaus^8^) and • safety and feasibility of drug design for the Wnt pathway (Kahn^9^, Garber^10^, Voronkov and Krauss^11^, Blagodatski *et al*.^12^ & Curtin and Lorenzi^13^). Approximately forty years after the discovery, important strides have been made in the research work involving several wet lab experiments and cancer clinical trials (Kahn^9^, Curtin and Lorenzi^13^) which have been augmented by the recent developments in the various advanced computational modeling techniques of the pathway. More recent informative reviews have touched on various issues related to the different types of the Wnt signaling pathway and have stressed not only the activation of the Wnt signaling pathway via the Wnt proteins (Rao and Kühl^14^) but also the on the secretion mechanism that plays a major role in the initiation of the Wnt activity as a prelude (Yu and Virshup^15^).

The work in this paper investigates some of the current aspects of research regarding the pathway via sensitivity analysis while using static (Jiang *et al*.^16^) and time series (Gujral and MacBeath^17^) gene expression data retrieved from colorectal cancer samples.

### 1.2 Canonical Wnt signaling pathway

Before delving into the problem statement, a brief introduction to the Wnt pathway is given here. From the recent work of Sinha^18^, the canonical Wnt signaling pathway is a transduction mechanism that contributes to embryo development and controls homeostatic self renewal in several tissues (Clevers^4^). Somatic mutations in the pathway are known to be associated with cancer in different parts of the human body. Prominent among them is the colorectal cancer case (Gregorieff and Clevers^19^). In a succinct overview, the Wnt signaling pathway works when the Wnt ligand gets attached to the Frizzled(*FZD*)/*LRP* coreceptor complex. *FZD* may interact with the Dishevelled (*DVL*) causing phosphorylation. It is also thought that Wnts cause phosphorylation of the *LRP* via casein kinase 1 (*CK*1) and kinase *GSK*3. These developments further lead to attraction of Axin which causes inhibition of the formation of the degradation complex. The degradation complex constitutes of *AXIN*, the *β*-*catenin* transportation complex *APC*, *CK*1 and *GSK*3. When the pathway is active the dissolution of the degradation complex leads to stabilization in the concentration of *β*-*catenin* in the cytoplasm. As *β*-*catenin* enters into the nucleus it displaces the *GROUCHO* and binds with transcription cell factor *TCF* thus instigating transcription of Wnt target genes. *GROUCHO* acts as lock on *TCF* and prevents the transcription of target genes which may induce cancer. In cases when the Wnt ligands are not captured by the coreceptor at the cell membrane, *AXIN* helps in formation of the degradation complex. The degradation complex phosphorylates *β*-*catenin* which is then recognized by *FBOX*/*WD* repeat protein *β*-*TRCP*. *β*-*TRCP* is a component of ubiquitin ligase complex that helps in ubiquitination of *β*-*catenin* thus marking it for degradation via the proteasome. Cartoons depicting the phenomena of Wnt being inactive and active are shown in figures 1(A) and 1(B), respectively.

**Fig. 1.**
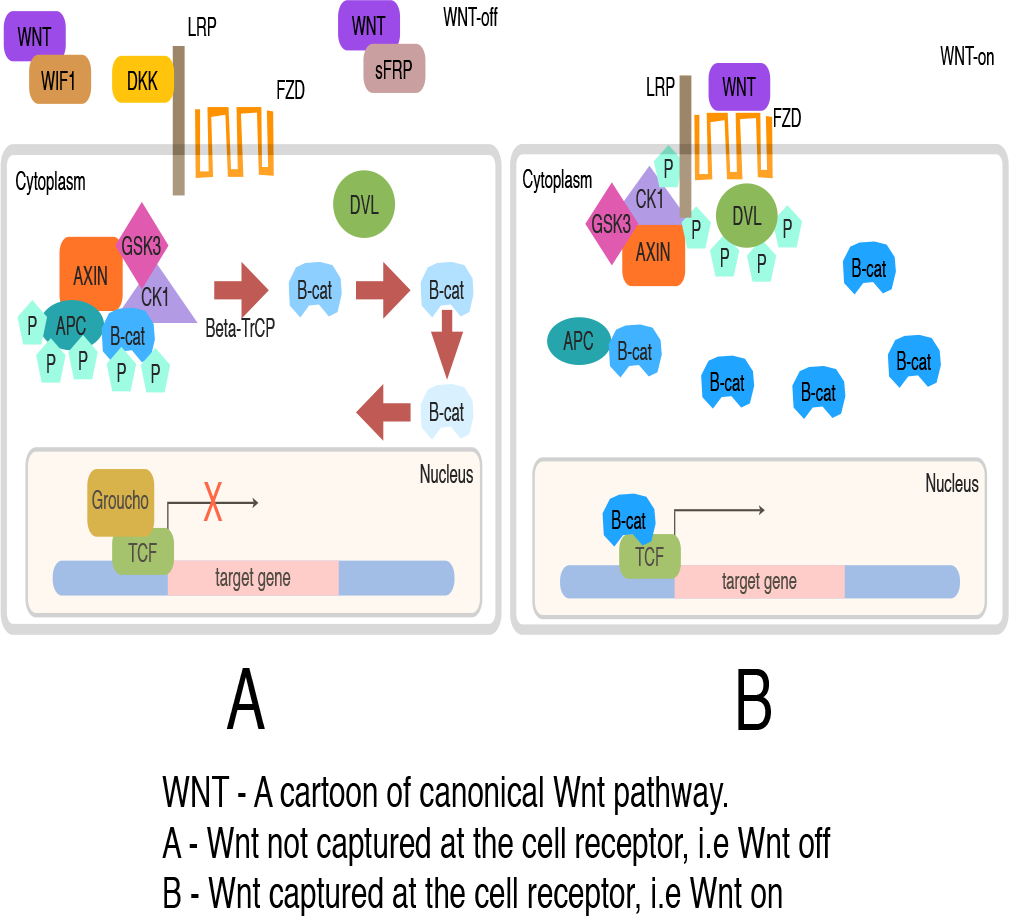
A cartoon of Wnt signaling pathway. Part (A) represents the destruction of *γ-catenin* leading to the inactivation of the Wnt target gene. Part (B) represents activation of Wnt target gene.

## 2 Problem statement & sensitivity analysis

It is widely known that the sensitivity analysis plays a major role in computing the strength of the influence of involved factors in any phenomena under investigation. When applied to expression profiles of various intra/extracellular factors that form an integral part of a signaling pathway, the variance and density based analysis yields a range of sensitivity indices for individual as well as various combinations of factors. These combinations denote the higher order interactions among the involved factors. Computation of higher order interactions is often time consuming but they give a chance to explore the various combinations that might be of interest in the working mechanism of the pathway. For example, in a range of fourth order combinations among the various factors of the Wnt pathway, it would be easy to assess the influence of the destruction complex formed by APC, AXIN, CSKI and GSK3 interaction. But the effect of these combinations vary over time as measurements of fold changes and deviations in fold changes vary. So it is imperative to know how an interaction or a combination of the involved factors behave in time and a procedure needs to be developed that can track the behaviour by exploiting the influences of the involved factors.

In this work, after estimating the individual effects of factors for a higher order combination, the individual indices are considered as discriminative features. A combination, then is a multivariate feature set in higher order (>2). With an excessively large number of factors involved in the pathway, it is difficult to search for important combinations in a wide search space over different orders. Exploiting the analogy with the issues of prioritizing webpages using ranking algorithms, for a particular order, a full set of combinations of interactions can then be prioritized based on these features using a powerful ranking algorithm via support vectors (Joachims^20^). Recording the changing rankings of the combinations over time reveals of how higher order interactions behave within the pathway and when an intervention might be necessary to influence the interaction within the pathway. This could lead to development of time based therapeutic interventions. A graphical abstract of how the search engine works to find the crucial interactions by ranking in time has been represented in figure 2.

**Fig. 2.**
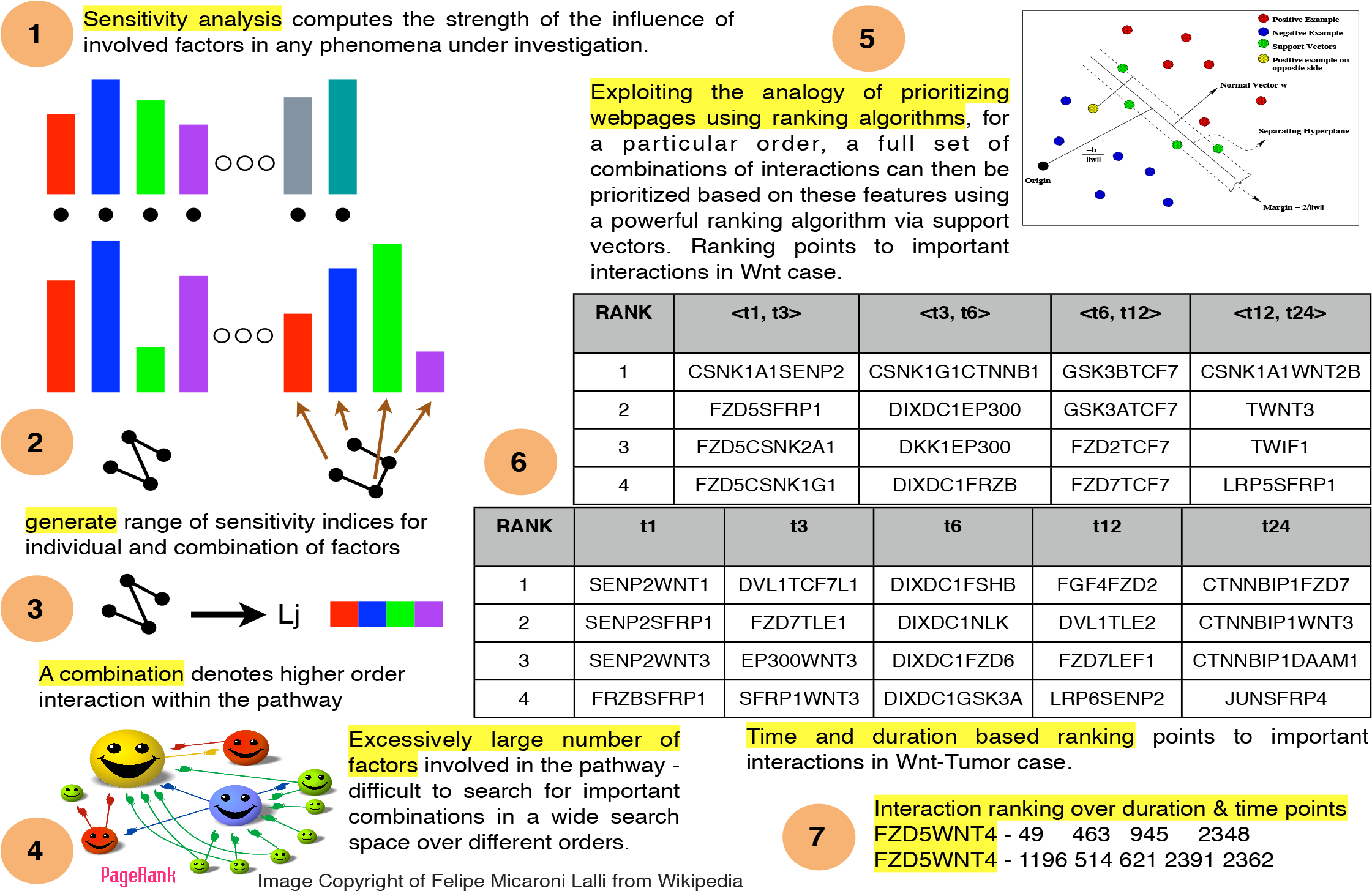
Graphical asbtract of how the whole search engine works to find out crucial interactions within the components of the pathway under consideration as different time points and durations.

### 2.1 Sensitivity analysis

Seminal work by Russian mathematician Sobol’^21^ lead to development as well as employment of SA methods to study various complex systems where it was tough to measure the contribution of various input parameters in the behaviour of the output. A recent unpublished review on the global SA methods by Iooss and Lemaître^22^ categorically delineates these methods with the following functionality • screening for sorting influential measures (Morris^23^ method, Group screening in Moon *et al*.^24^ & Dean and Lewis^25^, Iterated factorial design in Andres and Hajas^26^, Sequential bifurcation in Bettonvil and Kleijnen^27^ and Cotter^28^ design), • quantitative indicies for measuring the importance of contributing input factors in linear models (Christensen^29^, Saltelli *et al*.^30^, Helton and Davis^31^ and McKay *et al*.^32^) and nonlinear models (Homma and Saltelli^33^, Sobol^34^, Saltelli^35^, Saltelli *et al*.^36^, Saltelli *et al*.^37^, Cukier *et al*.^38^, Saltelli *et al*.^39^, & Tarantola *et al*.^40^ Saltelli *et al*.^41^, Janon *et al*.^42^, Owen^43^, Tissot and Prieur^44^, Da Veiga and Gamboa^45^, Archer *et al*.^46^, Tarantola *et al*.^47^, Saltelli and Annoni^48^ and Jansen^49^) and • exploring the model behaviour over a range on input values (Storlie and Helton^50^ and Da Veiga *et al*.^51^, Li *et al*.^52^ and Hajikolaei and Wang^53^). Iooss and Lemaître^22^ also provide various criteria in a flowchart for adapting a method or a combination of the methods for sensitivity analysis. Figure 3 shows the general flow of the mathematical formulation for computing the indices in the variance based Sobol method. The general idea is as follows - A model could be represented as a mathematical function with a multidimensional input vector where each element of a vector is an input factor. This function needs to be defined in a unit dimensional cube. Based on ANOVA decomposition, the function can then be broken down into *ƒ*_0_ and summands of different dimensions, if *ƒ*_0_ is a constant and integral of summands with respect to their own variables is 0. This implies that orthogonality follows in between two functions of different dimensions, if at least one of the variables is not repeated. By applying these properties, it is possible to show that the function can be written into a unique expansion. Next, assuming that the function is square integrable variances can be computed. The ratio of variance of a group of input factors to the variance of the total set of input factors constitute the sensitivity index of a particular group. Detailed derivation is present in the Appendix.

**Fig. 3.**
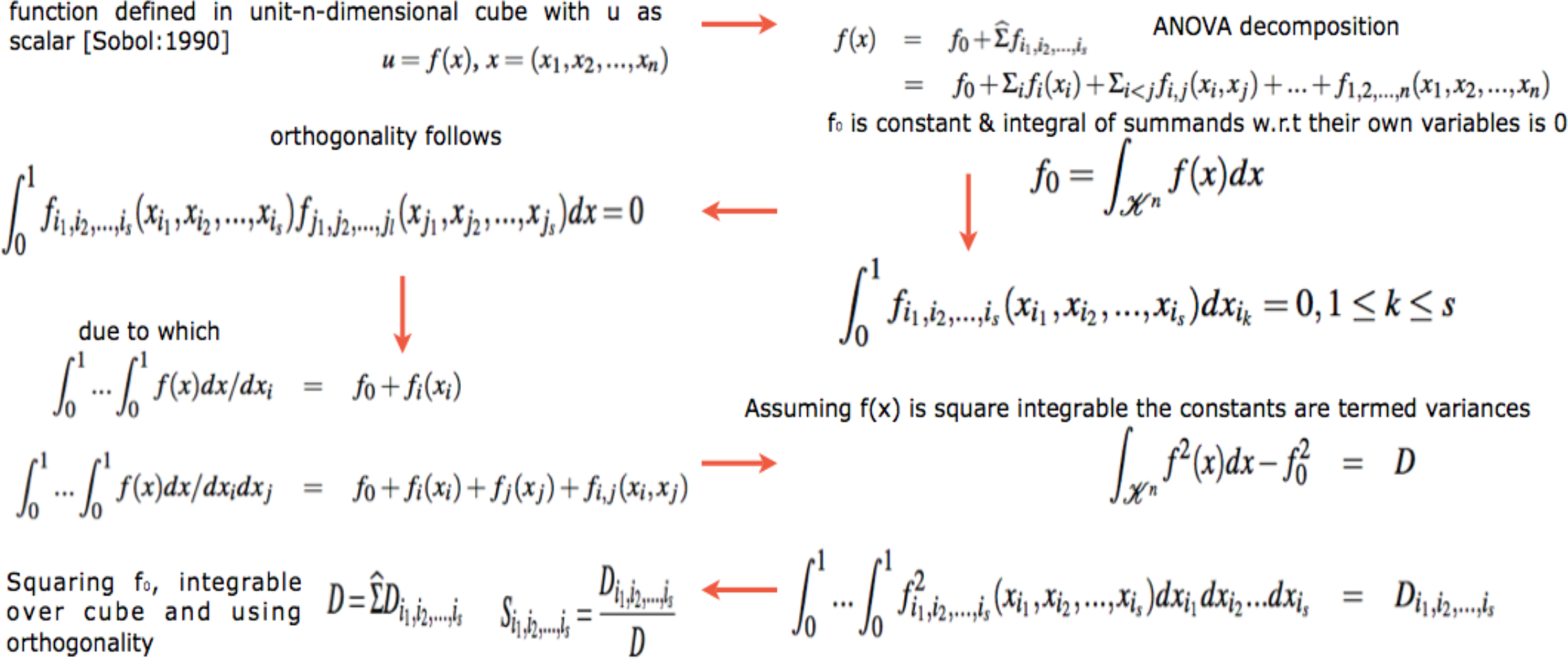
Computation of variance based sobol sensitivity indices. For detailed notations, see appendix.

Besides the above Sobol’^21^’s variance based indicies, more recent developments regarding new indicies based on density, derivative and goal-oriented can be found in Borgonovo^54^, Sobol and Kucherenko^55^ and Fort *et al*.^56^, respectively. In a latest development, Da Veiga^57^ propose new class of indicies based on density ratio estimation (Borgonovo^54^) that are special cases of dependence measures. This in turn helps in exploiting measures like distance correlation (Székely *et al*.^58^) and Hilbert-Schmidt independence criterion (Gretton *et al*.^59^) as new sensitivity indicies. The framework of these indicies is based on use of Csiszár *et al*.^60^ f-divergence, concept of dissimilarity measure and kernel trick Aizerman *et al*.^61^. Finally, Da Veiga^57^ propose feature selection as an alternative to screening methods in sensitivity analysis. The main issue with variance based indicies (Sobol’^21^) is that even though they capture importance information regarding the contribution of the input factors, they • do not handle multivariate random variables easily and • are only invariant under linear transformations. In comparison to these variance methods, the newly proposed indicies based on density estimations (Borgonovo^54^) and dependence measures are more robust. Figure 4 shows the general flow of the mathematical formulation for computing the indices in the density based HSIC method. The general idea is as follows - The sensitivity index is actually a distance correlation which incorporates the kernel based Hilbert-Schmidt Information Criterion between two input vectors in higher dimension. The criterion is nothing but the Hilbert-Schmidt norm of cross-covariance operator which generalizes the covariance matrix by representing higher order correlations between the input vectors through nonlinear kernels. For every operator and provided the sum converges, the Hilbert-Schmidt norm is the dot product of the orthonormal bases. For a finite dimensional input vectors, the Hilbert-Schmidt Information Criterion estimator is a trace of product of two kernel matrices (or the Gram matrices) with a centering matrix such that HSIC evalutes to a summation of different kernel values. Detailed derivation is present in the Appendix.

**Fig. 4.**
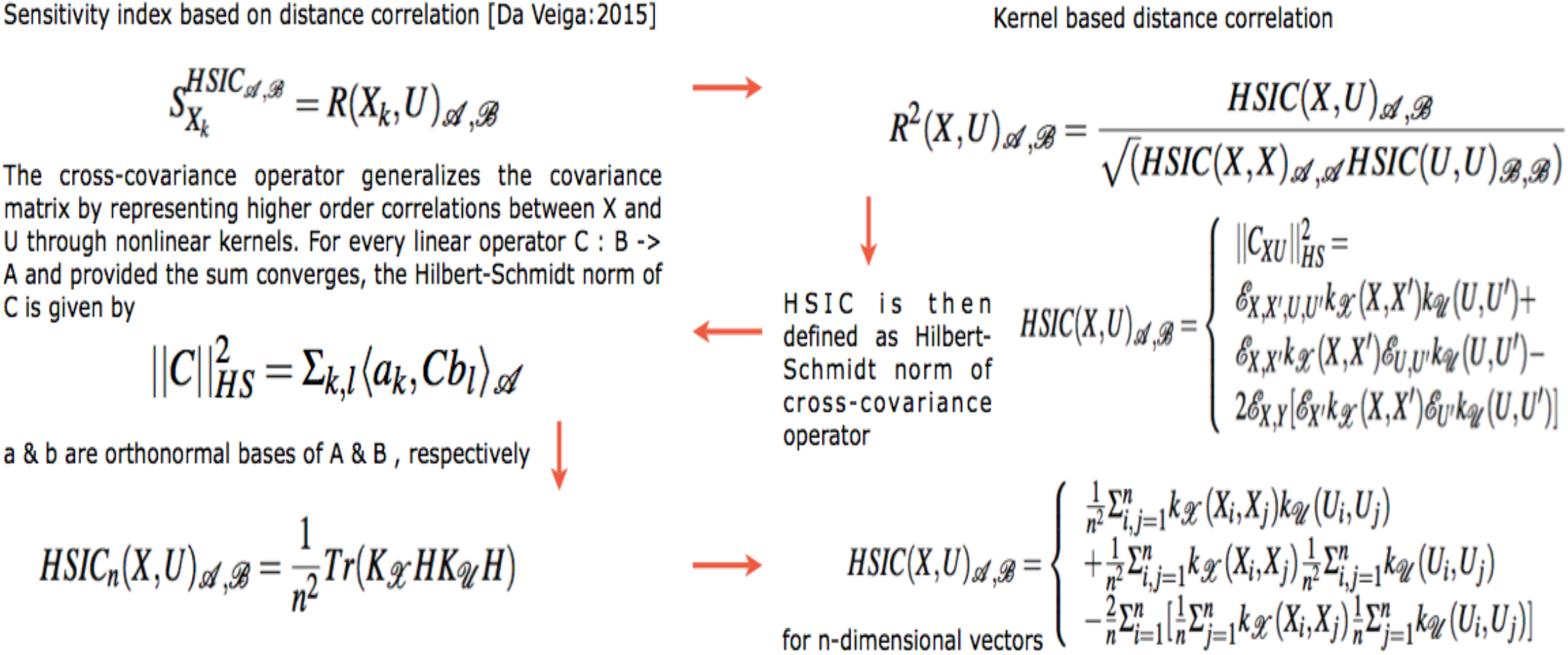
Computation of density based hsic sensitivity indices. For detailed notations, see appendix.

It is this strength of the kernel methods that HSIC is able to capture the deep nonlinearities in the biological data and provide reasonable information regarding the degree of influence of the involved factors within the pathway. Improvements in variance based methods also provide ways to cope with these nonlinearities but do not exploit the available strength of kernel methods. Results in the later sections provide experimental evidence for the same.

### 2.2 Relevance in systems biology

Recent efforts in systems biology to understand the importance of various factors apropos output behaviour has gained prominence. Sumner *et al*.^62^ compares the use of Sobol’^21^ variance based indices versus Morris^23^ screening method which uses a One-at-a-time (OAT) approach to analyse the sensitivity of *GSK*3 dynamics to uncertainty in an insulin signaling model. Similar efforts, but on different pathways can be found in Zheng and Rundell^63^ and Marino *et al*.^64^.

SA provides a way of analyzing various factors taking part in a biological phenomena and deals with the effects of these factors on the output of the biological system under consideration. Usually, the model equations are differential in nature with a set of inputs and the associated set of parameters that guide the output. SA helps in observing how the variance in these parameters and inputs leads to changes in the output behaviour. The goal of this manuscript is not to analyse differential equations and the parameters associated with it. Rather, the aim is to observe which input genotypic factors have greater contribution to observed phenotypic behaviour like a sample being normal or cancerous in both static and time series data. In this process, the effect of fold changes and deviations in fold changes in time is also considered for analysis in the light of the recently observed psychophysical laws acting downstream of the Wnt pathway (Goentoro and Kirschner^65^).

There are two approaches to sensitivity analysis. The first is the local sensitivity analysis in which if there is a required solution, then the sensitivity of a function apropos a set of variables is estimated via a partial derivative for a fixed point in the input space. In global sensitivity, the input solution is not specified. This implies that the model function lies inside a cube and the sensitivity indices are regarded as tools for studying the model instead of the solution. The general form of g-function (as the model or output variable) is used to test the sensitivity of each of the input factor (i.e expression profile of each of the genes). This is mainly due to its non-linearity, non-monotonicity as well as the capacity to produce analytical sensitivity indices. The g-function takes the form

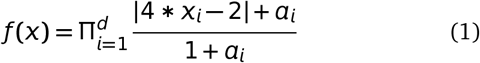

were, *d* is the total number of dimensions and *a_i_* ≥ 0 are the indicators of importance of the input variable *x_i_*. Note that lower values of *a_i_* indicate higher importance of *x_i_*. In our formulation, we randomly assign values of *x_i_* ∈ [0, 1]. For the static (time series) data *d* **=** 71 (factors affecting the pathway). The value of *d* varies from 2 to 70, depending on the order of the combination one might be interested in. Thus the expression profiles of the various genetic factors in the pathway are considered as input factors and the global analysis conducted. Note that in the predefined dataset, the working of the signaling pathway is governed by a preselected set of genes that affect the pathway. For comparison purpose, the local sensitivity analysis method is also used to study how the individual factor is behaving with respect to the remaining factors while working of the pathway is observed in terms of expression profiles of the various factors.

Finally, in context of Goentoro and Kirschner^65^’s work regarding the recent development of observation of Weber’s law working downstream of the pathway, it has been found that the law is governed by the ratio of the deviation in the input and the absolute input value. More importantly, it is these deviations in input that are of significance in studing such a phemomena. The current manuscript explores the sensitivity of deviation in the fold changes between measurements of fold changes at consecutive time points to explore in what duration of time, a particular factor is affecting the pathway in a major way. This has deeper implications in the fact that one is now able to observe when in time an intervention can be made or a gene be perturbed to study the behaviour of the pathway in tumorous cases. Thus sensitivity analysis of deviations in the mathematical formulation of the psychophysical law can lead to insights into the time period based influence of the involved factors in the pathway. Thus, both global and local anaylsis methods are employed to observe the entire behaviour of the pathway as well as the local behaviour of the input factors with respect to the other factors, respectively, via analysis of fold changes and deviations in fold changes, in time.

Given the range of estimators available for testing the sensitivity, it might be useful to list a few which are going to be employed in this research study. Also, a brief introduction into the fundamentals of the derivation of the three main indicies and the choice of sensitivity packages which are already available in literature, has been described in the Appendix.

### 2.3 Ranking Support Vector Machines

Learning to rank is a machine learning approach with the idea that the model is trained to learn how to rank. A good introduction to this work can be found in^66^. Existing methods involve pointwise, pairwise and listwise approaches. In all these approaches, Support Vector Machines (SVM) can be employed to rank the required query. SVMs for pointwise approach build various hyperplanes to segregate the data and rank them. Pairwise approach uses ordered pair of objects to classify the objects and then utilize the classifyer to rank the objects. In this approach, the group structure of the ranking is not taken into account. Finally, the listwise ranking approach uses ranking list as instances for learning and prediction. In this case the ranking is straightforward and the group structure of ranking is maintained. Various different designs of SVMs have been developed and the research in this field is still in preliminary stages. In context of the gene expression data set employed in this manuscript, the objects are the genes with their recorded FOLD CHANGE EXPRESSION VALUES AT DIFFERENT TIME SNAP SHOTS. Another case is the genes with their recorded DEVIATIONS IN FOLD CHANGE EXPRESSION VALUES IN BETWEEN TIME SNAP SHOTS.

Note that rankings algorithms have been developed to be employed in the genomic datasets but to the author’s awareness, these algorithms do not rank the range of combinations in a wide combinatorial search space in time. Also, they do not take into account the ranking of unexplored biological hypothesis which are assigned to a particular sensitivity value or vector that can be used for prioritization. For example, Kolde *et al*.^67^ presents a ranking algorithm that betters existing ranking model based on the assignment of *P*-value. As stated by Kolde *et al*.^67^ it detects genes that are ranked consistently better than expected under null hypothesis of uncorrelated inputs and assigns a significance score for each gene. The underlying probabilistic model makes the algorithm parameter free and robust to outliers, noise and errors. Significance scores also provide a rigorous way to keep only the statistically relevant genes in the final list. The proposed work here develops on sensitivity analysis and computes the influences of the factors for a system under investigation. These sensitivity indices give a much realistic view of the biological influence than the proposed *P*-value assignment and the probabilistic model. The manuscript at the current stage does not compare the algorithms as it is a pipeline to investigate and conduct a systems wide study. Instead of using SVM-Ranking it is possible to use other algorithms also, but the author has restricted to the development of the pipeline per se. Finally, the current work tests the effectiveness of the variance based (SOBOL) sensitivity indices apropos the density and kernel based (HSIC) sensitivity indices. Finally, Blondel *et al*.^68^ provides a range of comparison for 10 different regression methods and a score to measure the models. Compared to the frame provided in Blondel *et al*.^68^, the current pipeline takes into account biological information an converts into sensitivity scores and uses them as discriminative features to provide rankings. Thus the proposed method is algorithm independent.

## 3 Description of the dataset & design of experiments

Gujral and MacBeath^17^ present a bigger set of 71 Wnt-related gene expression values for 6 different times points over a range of 24-hour period using qPCR. The changes represent the fold-change in the expression levels of genes in 200 ng/mL *WNT*3*A*-stimulated HEK 293 cells in time relative to their levels in un-stimulated, serum-starved cells at 0-hour. The data are the means of three biological replicates. Only genes whose mean transcript levels changed by more than two-fold at one or more time points were considered significant. Positive (negative) numbers represent up (down) -regulation. Note that green (red) represents activation (repression) in the heat maps of data in Gujral and MacBeath^17^. For the case of time series data, interactions among the contributing factors are studied by comparing (1) pairs of fold-changes at single time points and (2) pairs of deviations in fold changes between pairs of time points.

The prodecure begins with the generation of distribution around measurements at single time points with added noise is done to estimate the indices. Two types of distributions are generated; One for the fold changes at single time points and the other for deviations in fold changes that recorded between fold changes at two time points. Then for every gene, there is a vector of values representing fold changes as well as deviations in fold changes for different time points and durations between time points, respectively. Next a listing of all 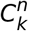 combinations for *k* number of genes from a total of *n* genes is generated. *k* is ≥ 2 and ≤ (*n*– 1). Each of the combination of order *k* represents a unique set of interaction between the involved genetic factors. After this, the datasets are combined in a specifed format which go as input as per the requirement of a particular sensitivity analysis method. Thus for each *p^th^* combination in 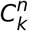 combinations, the dataset is prepared in the required format from the distributions for two separate cases which have been discussed above. (See .R code in mainScript-1-1.R in google drive and step 4 in figure 5). After the data has been transformed, vectorized programming is employed for density based sensitivity analysis and looping is employed for variance based sensitivity analysis to compute the required sensitivity indices for each of the *p* combinations. This procedure is done for different kinds of sensitivity analysis methods.

**Fig. 5.**
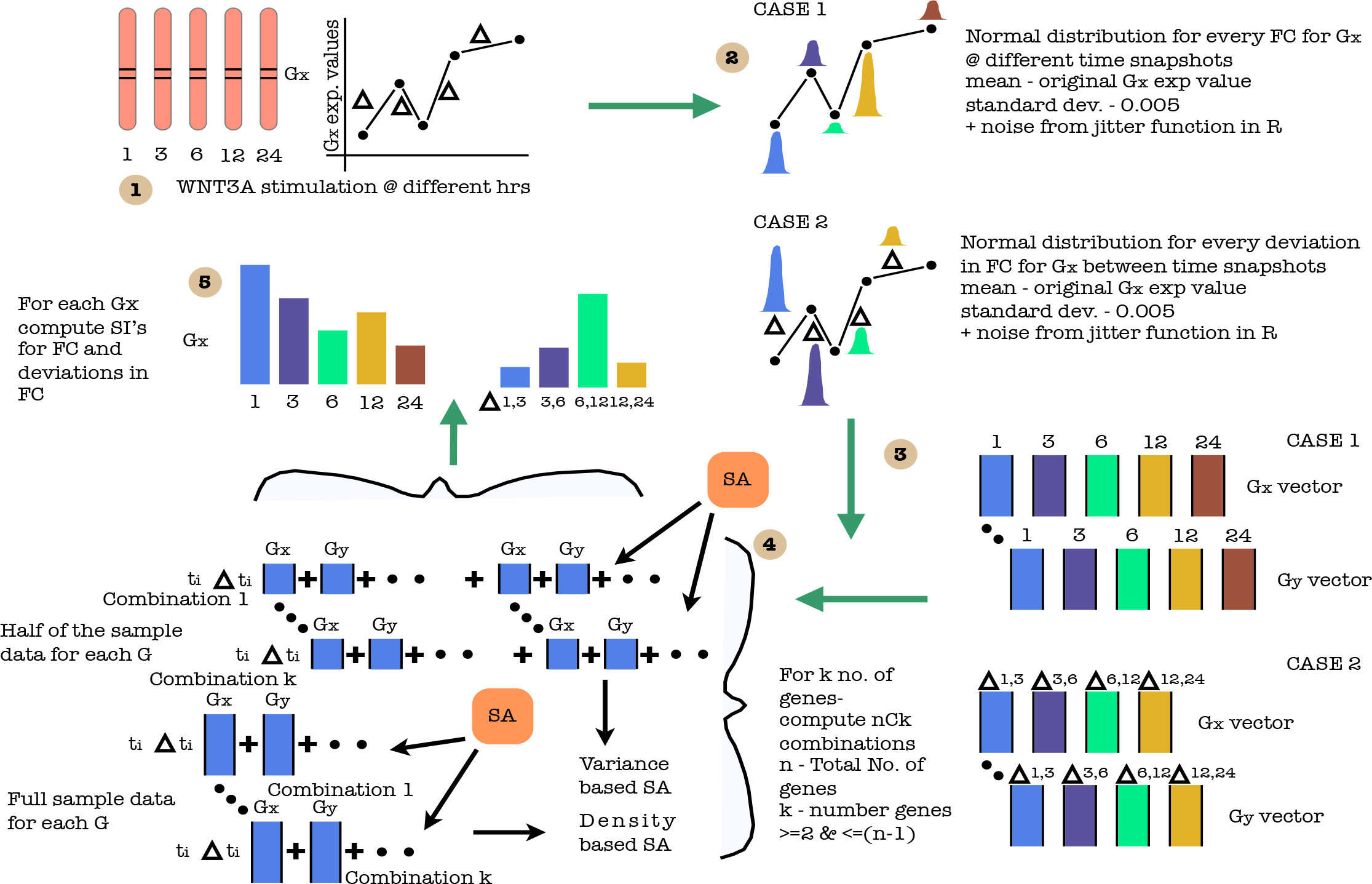
A cartoon of experimental setup. Step - (1) Time recordings of different gene expression values after WNT3A stimulation at different hrs. Generate deviations in fold changes per gene using the fold change recordings pers time point. (2) Generate of normal distribution for every FC & ΔFC for Gx at & between different time snapshots, respectively. mean - original Gx exp value; standard dev. - 0.005 + noise from jitter function in R (3) Generation of data set for FC & ΔFC. (4) For *k* number of genes, compute 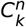 combinations and for every SA method, combine data in a specified format for both FC & ΔFC. (5) Compute SI for FC & ΔFC for different.

After the above sensitivity indices have been stored for each of the *p^th^* combination, the next step in the design of experiment is conducted. Since there is only one recording of sensitivity index per combination, each combination forms a training example which is alloted a training index and the sensitivity indices of the individual genetic factors form the training example. Thus there are 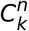 training examples for *k^th^* order interaction. Using this training set 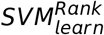 (Joachims^20^) is used to generate a model on default value *C* value of 20. In the current experiment on toy model *C* value has not been tunned.The training set helps in the generation of the model as the different gene combinations are numbered in order which are used as rank indices. The model is then used to generate score on the observations in the testing set using the 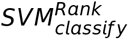 (Joachims^20^). Note that due to availability of only one example per combination, after the model has been built, the same training data is used as test data to generates the scores. This procedure is executed for each and every sensitivity analysis method. This is followed by sorting of these scores along with the rank indices (i.e the training indices) already assigned to the gene combinations. The end result is a sorted order of the gene combinations based on the ranking score learned by the *SVM^Rank^* algorithm. Finally, this entire procedure is computed for sensitivity indices generated for each and every fold change at time point and deviations in fold change at different durations. Observing the changing rank of a particular combination at different times and different time periods will reveal how a combination is behaving. These steps are depicted in figure 6.

**Fig. 6.**
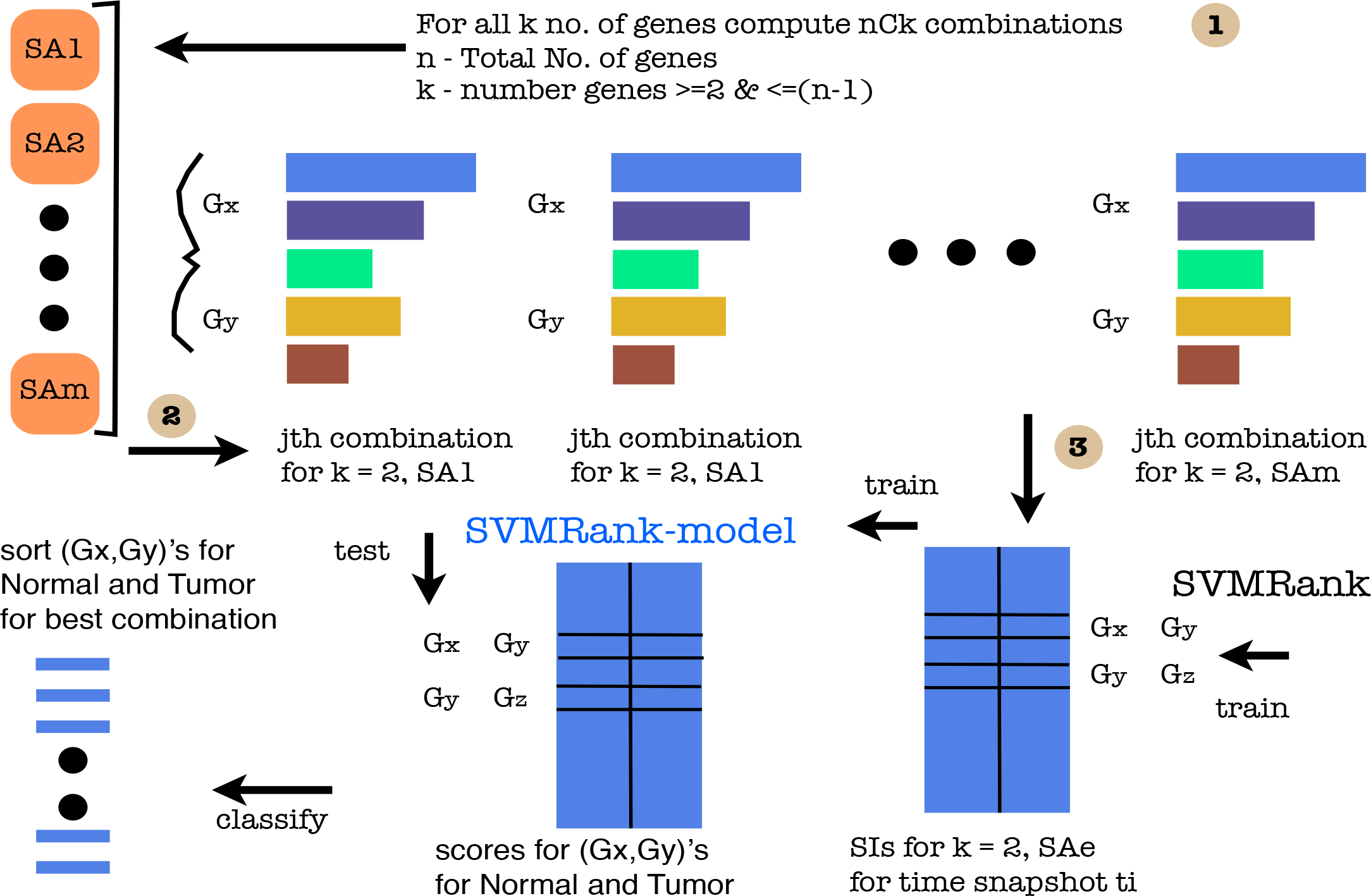
A cartoon of experimental setup. Step - (1) For *k^th^* order genes in 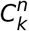 combinations and for every SA, combine (say for *k* = 2, i.e interaction level 2) SI’s of genetic factors form training data. (3) Using indices in (1) 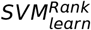 is employed on training data to generate a model. This model is used to generate a ranking score on the training data again via 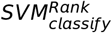. Finally the combinations are ranked based on sorting of training set. This is generated for each and every FC as well as mFC. Observing changing ranking in time & time periods reveals how a particular combination is behaving.

**Fig. 7.**
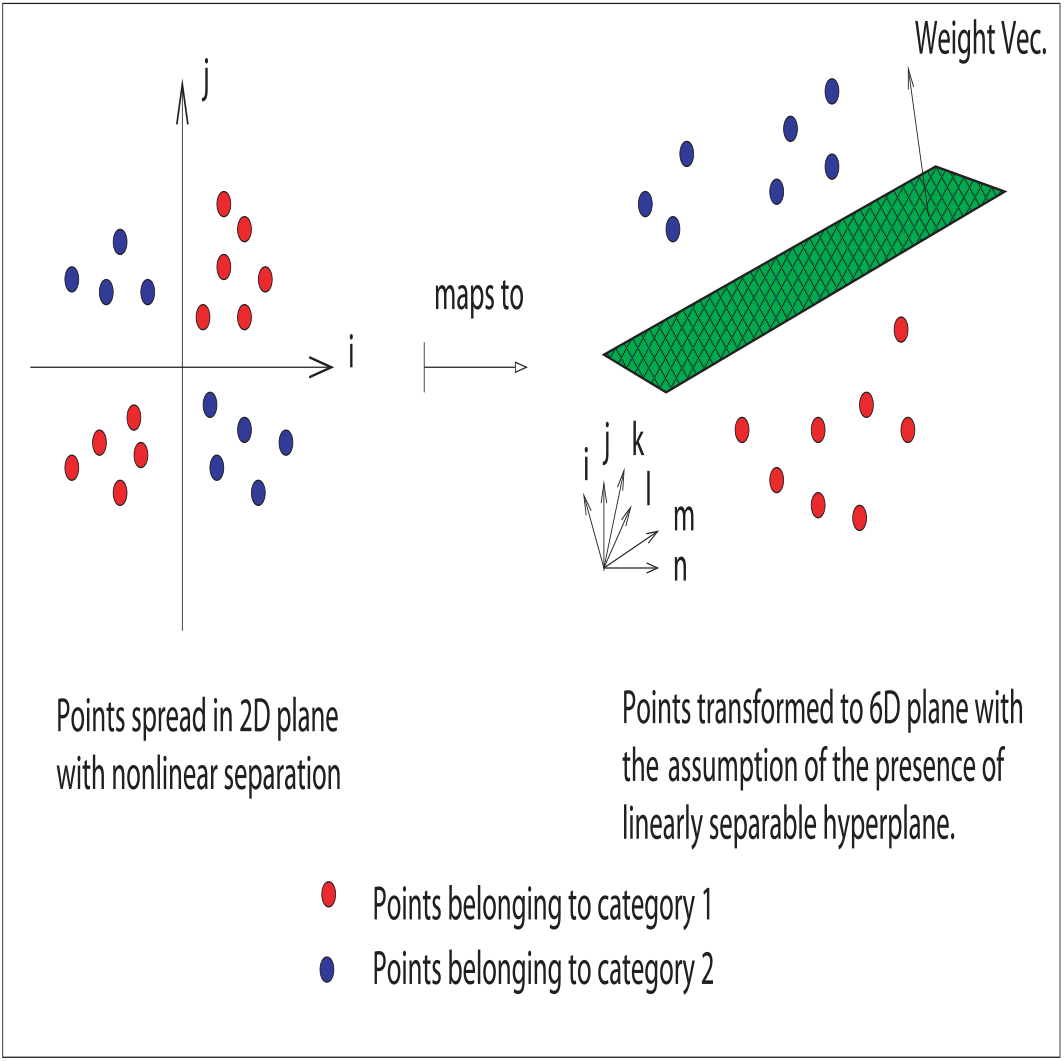
A geometrical interpretation of mapping nonlinearly separable data into higher dimensional space where it is assumed to be linearly separable, subject to the holding of dot product.

Note that the following is the order in which the files should be executed in R, in order, for obtaining the desired results (Note that the code will not be explained here) - • use source("mainScript-1-1.R") with arguments for Dynamic data • source("SVMRank-Results-D.R"), to rank the interactions (again this needs to be done separately for different kinds of SA methods), • use source("Combine-Time-files.R"), if computing indices separately via previous file, • source("Sort-n-Plot-D.R") to sort the interactions. Note that the sorting is chages the interaction ranking in time. Thus • use source("Interaction-Priority-Intime.R") to find the prioritized ranking of each and every interaction over the different time points and • finally use source("Print-Ranking-AND-Interaction-Rank.R") to print individual ranking of the required input factor with other interaction factors.

## 4 Results and Discussion

Results for the 2^*nd*^ order interactions are presented here. The results first discuss the behaviour of interactions across the snapshots of time using the computed sensitivities on fold change measurements per time snapshot. Next the behaviour of the interactions during the time intervals is analysed based on the computed sensitivities on deviations on fold change measurements per time interval. The analysis was done on 12 different sensitivity indices. Results obtained for one of the indices is explained. Next, individual rankings of a few interactions are discussed with respect to the used indices. Finally, the similar rankings for different sensitivity analysis method are made available at the website with contains the code for the same.

### 4.1 2^*nd*^ order interaction rankings at *t_i_* snapshot & Δ*t_i,i_*_+1_ interval

Table 1 (2) shows the 1^*st*^ first 15 and the last 15 ranking of the interactions in the first and the third slot of the 15 rows each, for time snapshots (for time periods between the snapshots). The next two slots contain the ranking behaviour of 1^*st*^ 15 & last 15 interactions at different time snapshots (different time periods for table 2). Note that the interaction order for this second slot of rows is based on the ranking obtained at *t*_1_ (Δ*t*_1,3_) in table 1 (2). First, the ranking for each of the interaction was generated per time snap shot. Column wise, it is observed that any interaction will take up different ranks at the different time snapshots (and time periods). This ranking is based on the sensitivity indices that capture the amount of influence that the combination of factors have on the signaling pathway. To have a better view of how the individual interaction works, the rankings of a particular interaction across the different time snapshots (and time periods) were extracted from the entire set of rankings of all interactions per time snapshot (ans time periods). Both tables 1 and 2 show a glimpse of a few 2^*nd*^ order interactions out of the total of 71 genes. The total number of combinations are 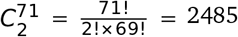.

**Table 1.** 1^*st*^ and 3^*rd*^ set of 15 rows - Ranking of 1^*st*^ 15 & last 15 interactions at different time snapshots. 2^*nd*^ and 4^*th*^ set of 15 rows - Ranking behaviour of 1st 15 & last 15 interactions at different time snapshots. Note that the interaction order for this 2^*nd*^ and 4^*th*^ slot of rows is based on the ranking obtained at *t*_1_. SA - HSICrbf

| RANKING @ $t_1$ USING HSIC - RBF | | | | | |
| --- | --- | --- | --- | --- | --- |
| Rank @ - | $t_1$ | $t_3$ | $t_6$ | $t_{12}$ | $t_{24}$ |
| 1 | SEN2WNT1 | DVL1TCF7L1 | DIXDC1FSHB | FGF4FZD2 | CTNNBIP1FZD7 |
| 2 | SEN2SFRP1 | FZD7TLE1 | DIXDC1NLK | DVL1TLE2 | CTNNBIP1WNT3 |
| 3 | SEN2WNT3 | EP300WNT3 | DIXDC1FZD6 | FZD7LEF1 | CTNNBIP1DAAM1 |
| 4 | FRZBSFRP1 | SFRP1WNT3 | DIXDC1GSK3A | LRP6SEN2 | JUNSRFP4 |
| 5 | FRZBGSK3A | TCF7L1WNT3 | FRZBPPP2CA | PYGO1WNT4 | TWNT1 |
| 6 | DKK1GSK3A | CSNK1A1FOSL1 | DIXDC1PPP2CA | FZD8WNT1 | WIF1WNT1 |
| 7 | SEN2T | NKD1WNT1 | DIXDC1FBXW11 | DVL1DVL2 | FZD1JUN |
| 8 | SEN2WIF1 | CCND3WNT1 | FGF4PPP2CA | FZD2TLE2 | MYCSFRP1 |
| 9 | FRZBFZD2 | KREMEN1WNT3A | DIXDC1WNT2B | SFRP1T | CTNNBIP1SFRP1 |
| 10 | FRZBPYGO1 | EP300WNT2B | DIXDC1FOXN1 | BTRCPYGO1 | FRZBFZD2 |
| 11 | FRZBJUN | TLE1WNT3 | DIXDC1DKK1 | DIXDC1GSK3B | AXIN1WIF1 |
| 12 | FRZBLEF1 | FZD1FZD8 | DIXDC1WNT1 | PYGO1SLC9A3R1 | CTNNBIP1WNT1 |
| 13 | FRZBPORCN | FZD7PPP2R1A | DIXDC1TCF7 | BCL9LEF1 | CTNNBIP1FZD8 |
| 14 | SEN2TLE1 | FZD8KREMEN1 | FGF4WNT2B | CTNNBIP1FOSL1 | CTBP1WNT3 |
| 15 | DKK1WNT1 | FZD6KREMEN1 | DIXDC1LRP6 | CTNNBIP1WNT2B | MYCPYGO1 |
| 2<sup>nd</sup> Ord. Interact. | Rank @ - $t_1$ | $t_3$ | $t_6$ | $t_{12}$ | $t_{24}$ |
| SEN2WNT1 | 1 | 887 | 42 | 16 | 407 |
| SEN2SFRP1 | 2 | 844 | 564 | 1691 | 202 |
| SEN2WNT3 | 3 | 461 | 861 | 992 | 751 |
| FRZBSFRP1 | 4 | 1416 | 1440 | 1205 | 217 |
| FRZBGSK3A | 5 | 1482 | 90 | 1544 | 403 |
| DKK1GSK3A | 6 | 2313 | 689 | 1907 | 1362 |
| SEN2T | 7 | 1140 | 116 | 126 | 1083 |
| SEN2WIF1 | 8 | 1471 | 547 | 2184 | 2070 |
| FRZBFZD2 | 9 | 930 | 923 | 955 | 10 |
| FRZBPYGO1 | 10 | 2291 | 243 | 346 | 113 |
| FRZBJUN | 11 | 2412 | 644 | 507 | 409 |
| FRZBLEF1 | 12 | 629 | 987 | 1090 | 1958 |
| FRZBPORCN | 13 | 1981 | 800 | 1938 | 85 |
| SEN2TLE1 | 14 | 105 | 1586 | 1853 | 339 |
| DKK1WNT1 | 15 | 2415 | 1260 | 2300 | 1302 |
| Rank @ - | $t_1$ | $t_3$ | $t_6$ | $t_{12}$ | $t_{24}$ |
| 2471 | CCND1SEN2 | CXXC4SLC9A3R1 | AXIN1SFRP4 | DAAM1GSK3A | KREMEN1PPP2CA |
| 2472 | LRP6SEN2 | AXIN1DVL1 | LRP5SFRP1 | CCND2WNT3 | CTBP2TCF7L1 |
| 2473 | FOXN1SEN2 | LRP6NKD1 | CTBP2FGF4 | SFRP1TCF7L1 | AESFZD5 |
| 2474 | CCND2SEN2 | SFRP4WNT5A | DKK1LEF1 | CCND3CSNK1D | NKD1TLE2 |
| 2475 | CCND1FRZB | FRZBSEN2 | APCTLE1 | FOXN1FZD8 | BCL9PYGO1 |
| 2476 | BTRCSEN2 | FOSL1WNT1 | BTRCWNT5A | FBXW2LRP6 | CCND2LEF1 |
| 2477 | PORCSEN2 | BCL9FBXW4 | BCL9WNT3 | PPP2CAWNT3A | CXXC4GSK3B |
| 2478 | AXIN1SEN2 | FRZBWNT1 | AESCTBP1 | DIXDC1TLE1 | DIXDC1GSK3B |
| 2479 | LEF1SEN2 | CTBP1TCF7 | CCND1SLC9A3R1 | FZD7TCF7L1 | TLE2WNT2B |
| 2480 | MYCSEN2 | SFRP4WIF1 | CCND1CXXC4 | CCND3CSNK2A1 | CSNK1DCSNK2A1 |
| 2481 | GSK3ASEN2 | CTBP2SFRP4 | GSK3APORCN | CCND1FOXN1 | SFRP1WNT3 |
| 2482 | CSNK1A1SEN2 | SFRP4TCF7L1 | AESAPC | WIF1WNT2 | DIXDC1LEF1 |
| 2483 | AESSEN2 | FRZBWIF1 | NLKPORCN | CSNK2A1WNT3A | CSNK1G1WNT3A |
| 2484 | JUNSEN2 | CXXC4NKD1 | CSNK1G1FRAT1 | LRP6TCF7 | SFRP1WNT5A |
| 2485 | PYGO1SEN2 | SFRP4TLE1 | LEF1SLC9A3R1 | LRP5PORCN | SEN2WNT2 |
| 2<sup>nd</sup> Ord. Interact. | Rank @ - $t_1$ | $t_3$ | $t_6$ | $t_{12}$ | $t_{24}$ |
| CCND1SEN2 | 2471 | 555 | 1824 | 53 | 1640 |
| LRP6SEN2 | 2472 | 2268 | 877 | 4 | 659 |
| FOXN1SEN2 | 2473 | 427 | 1964 | 197 | 1578 |
| CCND2SEN2 | 2474 | 1527 | 1866 | 597 | 1670 |
| CCND1FRZB | 2475 | 1944 | 2196 | 2146 | 1400 |
| BTRCSEN2 | 2476 | 1848 | 2112 | 49 | 1484 |
| PORCSEN2 | 2477 | 1934 | 1963 | 400 | 1275 |
| AXIN1SEN2 | 2478 | 1724 | 2427 | 565 | 815 |
| LEF1SEN2 | 2479 | 2006 | 2172 | 188 | 1682 |
| MYCSEN2 | 2480 | 1021 | 2246 | 366 | 302 |
| GSK3ASEN2 | 2481 | 1798 | 1323 | 250 | 2261 |
| CSNK1A1SEN2 | 2482 | 989 | 2158 | 994 | 247 |
| AESSEN2 | 2483 | 115 | 2078 | 269 | 1239 |
| JUNSEN2 | 2484 | 1385 | 2300 | 1590 | 364 |
| PYGO1SEN2 | 2485 | 2126 | 1227 | 94 | 1945 |

**Table 2.** 1*^st^* and 3*^rd^* set of 15 rows - Ranking of 1*^st^* 15 & last 15 interactions at different time intervals. 2*^nd^* and 4*^th^* set of 15 rows - Ranking behaviour of 1st 15 & last 15 interactions at different time intervals. Note that the interaction order for the 2*^nd^* and 4*^th^* slot of rows is based on the ranking obtained at Δt_1,3_. SA - HSICrbf

| RANKING @ $\Delta t_{i,i+1}$ USING HSIC - RBF | | | | | |
| --- | --- | --- | --- | --- | --- |
| Rank @ - | $\Delta t_{1,3}$ | $\Delta t_{3,6}$ | $\Delta t_{6,12}$ | $\Delta t_{12,24}$ | |
| 1 | CSNK1A1SEN2 | CSNK1G1CTNNB1 | GSK3BTCF7 | CSNK1A1WNT2B |  |
| 2 | FZD5SFRP1 | DIXDC1EP300 | GSK3ATCF7 | TWNT3 |  |
| 3 | FZD5CSNK2A1 | DKK1EP300 | FZD2TCF7 | TWIF1 |  |
| 4 | FZD5CSNK1G1 | DIXDC1FRZB | FZD7TCF7 | LRP5SFRP1 |  |
| 5 | PYGO1SEN2 | FZD8NKD1 | FZD6NLK | JUNSRFP4 |  |
| 6 | PYGO1WNT5A | CSNK1G1EP300 | FZD7NKD1 | AESWIF1 |  |
| 7 | FZD5SEN2 | CTBP2CTNNB1 | FZD8NLK | TLE2WNT3 |  |
| 8 | FZD5FBXW2 | PITX2FBXW4 | FZD7SEN2 | CSNK1A1DKK1 |  |
| 9 | CSNK1A1CSNK1G1 | DIXDC1WNT1 | KREMEN1WNT2B | AESFZD6 |  |
| 10 | FZD5FZD7 | GSK3BNKD1 | FZD7MYC | AESDAAM1 |  |
| 11 | FZD5LRP6 | DIXDC1NKD1 | FZD6SEN2 | CSNK1A1FBXW11 |  |
| 12 | FZD5SFRP4 | DIXDC1FBXW4 | FZD7WNT3 | FRZBSFRP4 |  |
| 13 | FZD5LEF1 | DIXDC1DVL1 | FZD7SFRP4 | TLE2WIF1 |  |
| 14 | PYGO1FBXW4 | CSNK1G1CTBP1 | FZD7PPP2CA | CSNK1A1DVL1 |  |
| 15 | FZD5FGF4 | DIXDC1NLK | GSK3ASEN2 | RHOUWIF1 |  |
| 2<sup>nd</sup> Ord. Interact. | Rank @ - $\Delta t_{1,3}$ | $t_{3,6}$ | $t_{6,12}$ | $t_{12,24}$ | |
| CSNK1A1SEN2 | 1 | 1971 | 2108 | 98 |  |
| FZD5SFRP1 | 2 | 1461 | 1499 | 1726 |  |
| FZD5CSNK2A1 | 3 | 1099 | 646 | 1647 |  |
| FZD5CSNK1G1 | 4 | 2430 | 1410 | 2054 |  |
| PYGO1SEN2 | 5 | 2467 | 752 | 453 |  |
| PYGO1WNT5A | 6 | 479 | 506 | 1847 |  |
| FZD5SEN2 | 7 | 1110 | 2167 | 1402 |  |
| FZD5FBXW2 | 8 | 921 | 2298 | 1953 |  |
| CSNK1A1CSNK1G1 | 9 | 1546 | 1340 | 302 |  |
| FZD5FZD7 | 10 | 2424 | 509 | 2303 |  |
| FZD5LRP6 | 11 | 2235 | 1633 | 1593 |  |
| FZD5SFRP4 | 12 | 1326 | 2012 | 926 |  |
| FZD5LEF1 | 13 | 1838 | 758 | 937 |  |
| PYGO1FBXW4 | 14 | 245 | 1608 | 571 |  |
| FZD5FGF4 | 15 | 2225 | 1399 | 2305 |  |
| Rank @ - | $\Delta t_{1,3}$ | $\Delta t_{3,6}$ | $\Delta t_{6,12}$ | $\Delta t_{12,24}$ | |
| 2471 | FZD1WNT2B | CSNK2A1RHOU | FOSL1T | CTBP1DKK1 |  |
| 2472 | SEN2SLC9A3R1 | SEN2SFRP1 | CTBP2DVL2 | CTBP2WNT5A |  |
| 2473 | CSNK1G1MYC | JUNWNT1 | FZD6WNT3A | CSNK1G1CSNK2A1 |  |
| 2474 | NKD1WNT3 | TCF7L1WNT3 | DVL1FZD8 | FZD8WNT5A |  |
| 2475 | FGF4WNT3 | CCND3WNT1 | FGF4TLE2 | DVL2TCF7 |  |
| 2476 | FZD2PYGO1 | BCL9PYGO1 | EP300TLE1 | DIXDC1TLE1 |  |
| 2477 | FZD7PORCN | NKD1PORCN | CSNK1G1FSHB | DKK1WNT3A |  |
| 2478 | SFRP4WNT2 | NLKTLE2 | CSNK1A1SFRP4 | JUNRHOU |  |
| 2479 | LEF1PORCN | FRZBFZD6 | CTBP2PYGO1 | AESPPP2CA |  |
| 2480 | FGF4RHOU | PORCNWNT2B | DVL1GSK3A | KREMEN1WNT5A |  |
| 2481 | CSNK1G1CTNNBIP1 | FRAT1WIF1 | FOSL1PPP2CA | CTNNB1FZD8 |  |
| 2482 | SFRP4SLC9A3R1 | CTNNBIP1EP300 | LEF1SLC9A3R1 | CTBP1WNT4 |  |
| 2483 | CCND3CSNK1A1 | SEN2TCF7L1 | BCL9WNT5A | FBXW11TCF7L1 |  |
| 2484 | CSNK1G1WNT3 | APCDVL1 | FOSL1WNT1 | APCNLK |  |
| 2485 | PITX2PYGO1 | DVL1JUN | CSNK1A1FRZB | CSNK1G1WNT3A |  |
| 2<sup>nd</sup> Ord. Interact. | Rank @ - $\Delta t_{1,3}$ | $t_{3,6}$ | $t_{6,12}$ | $t_{12,24}$ | |
| FZD1WNT2B | 2471 | 694 | 516 | 191 |  |
| SEN2SLC9A3R1 | 2472 | 576 | 1199 | 981 |  |
| CSNK1G1MYC | 2473 | 370 | 484 | 638 |  |
| NKD1WNT3 | 2474 | 1329 | 2101 | 656 |  |
| FGF4WNT3 | 2475 | 1912 | 214 | 449 |  |
| FZD2PYGO1 | 2476 | 1027 | 608 | 1467 |  |
| FZD7PORCN | 2477 | 145 | 133 | 2132 |  |
| SFRP4WNT2 | 2478 | 351 | 1692 | 619 |  |
| LEF1PORCN | 2479 | 158 | 878 | 1778 |  |
| FGF4RHOU | 2480 | 587 | 2109 | 726 |  |
| CSNK1G1CTNNBIP1 | 2481 | 185 | 1545 | 618 |  |
| SFRP4SLC9A3R1 | 2482 | 498 | 1266 | 1106 |  |
| CCND3CSNK1A1 | 2483 | 784 | 324 | 427 |  |
| CSNK1G1WNT3 | 2484 | 2446 | 225 | 1872 |  |
| PITX2PYGO1 | 2485 | 144 | 369 | 2304 |  |

What we find is that the interactions or combinations are ranked but they do-not explicitly state the direct or indirect dependencies between the combinations. For example, we find that *FZD*7-*LEF*1 have an extremely high ranking of 3 at time snap-shot *t*_12_ in table 1. There might not be a direct link between these two but the prioritised list indicates their high ranking in the Wnt pathway at a certain stage and affecting these two in wet lab experiment at this stage would reveal a possible insights into the connectivity or influence these two have on the pathway. Thus unexplored biological hypothesis could be tested using these rankings. If we look at the *SENP*2-*WNT*1 combination and its ranking over time snapshots in table 1, we find that its influence on the pathway is extremely high except at time points *t*_3_ and *t*_24_. This suggests that the combination of *SENP*2-*WNT*1 might be playing an a highly influencing role in the pathway, either directly or indirectly. Recently, it has been shown by Du *et al*.^69^, that *SENP*2, a member of the family SUMOylation-related proteins does have increased level of expression in colorectal cancer cell lines. Thus, studying the ranking of a combination at various time snap shots can facilitate in understanding the influence of the combination as well as testing the unexplored biological hypothesis and developing target based therapeutic drugs that can interven the pathway.

In table 2, we observe similar rankings but instead of considering the prioritization at time points, we consider ranking at time periods. These are of much more greater importance that the rankings at the time points. This is attributed to the fact that the deviation in fold changes in between the recorded fold changes at consecutive time points indicate how much a particular combination is influencing the pathway in that particular period of time. Currently, the proposed insilico pipeline takes into account the data that has been recorded at long intervals of time (see Gujral and MacBeath^17^). But recordings at shorter time intervals will give a more detailed ranking list for the different time periods. *FZD*7-*LRP*6 combination shows an extremely high ranking in the first period of *t*_1,3_. But its ranking goes down in the second period before it steadily rises in ranking in the later periods. This suggests that in the initial period the combination of *FZD*7-*LRP*6 is extremely crucial when the *WNT*3*A* is been injected into the cell and the signaling has gained momentum. During the next period, the ranking significantly drops to a very low value indicating that the combination at the receptor level is not that effective. But this reverses in the next two periods. These changing rankings over time periods shows in which time period, a combination might be effective in the pathway and help the biologists to intervene the pathway at that particular period of time.

Gujral and MacBeath^17^ proposes a system wide analysis of the WNT pathway in which the analysis of *CTBP*1-*MYC* has been presented. It was observed that *CTBP*1 plays a positive role in the WNT pathway as a transcription factor in the early phases of the pathway and is correlated with the time dependent profile of *MYC*. The current pipeline confirms the wet lab experiments by indicating the changing ranking in time. For rankings in time-points • *CTBP*2*MYC* 401 352 433 1765 2024 • *CTBP*1*MYC* 411 2011 580 683 1227, it was found that *CTBP*1 and *MYC* have similar affect on the pathway. Initially the ranking is high but goes down significantly at the second timesnap shot before showing a decreasing peaked behaviour. Similarly, *CTBP*2 (though not reported in Gujral and MacBeath^17^) shows that a similar ranking pattern as the *CTBP*1-*MYC*. These findings point to the efficacy of the proposed pipeline. A further analysis of rankings in time periods show a completely different profile - • *CTBP*2*MYC* 1848 941 2037 1692 • *CTBP*1*MYC* 980 2321 2387 1558. These rankings over time periods give a different picture mainly due to the large gaps in time snapshot recordings. As stated earlier, the significance might be much improved if the duration between the time points at which the *WNT*3*A* is being injected is shortened. In general, a system wide analysis is now possible with a pipeline as developed in this work, which prioritises the interactions or combinations in a pathway over time snapshots and period of time.

### 4.2 Analysis of WNT-FZD variants at *t_i_* snapshot & Δ*t_i,i_*_+1_ interval

It is widely known that the family of *WNT* and *FZD* interact at the cell surface to instigate the pathway. Hsieh *et al*.^70^ conducted one of the earliest biochemical characterisation of the *WNT*-*FZD* interactions. The most recent wet lab study to map the particular combinations of *WNT*-*FZD* was done by Dijksterhuis *et al*.^71^. Albeit the different cell lines used for finding the combinations of *WNT*-*FZD*, we would like to know if the combinations generated by the proposed pipeline in this work have some match with the existing wet lab tested combinations. The existing problem with the combinations of *WNT*-*FZD* is that there is a large range of combinations of the order of 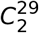 and the mapping of these combinations have not been done extensively. The reasons for this are purely biological in nature as unknown specific *WNT* binding activities and lack of binding assays for full length receptors (Dijksterhuis *et al*.^71^). Nevertheless, despite these hinderances, advances in wet lab have been made to map these combinations. The major problem here is the amount to time needed to search for these specific bindings and from a biological standpoint, it is not easy to find which combination to search for first, given the vast combinatorial space. We test the pipeline to generate ranks for a few *WNT*-*FZD* interactions and a prioritised list has been provided which contains the changing ranks at different time points and time periods. For data set generated from the cell lines in Gujral and MacBeath^17^, a limited number of *WNT* and *FZD* family members were profiled as the *WNT*3*A* molecule was injected into the serum starved cells. Tables 3 and 4 represent the rankings at different time points while tables 5 and 6 represent the rankings at different time periods, for the different *WNT*-*FZD* interactions.

**Table 3.** 2*^nd^* order interaction ranking for FZD-WNT variants using HSIC-rbf for different intervals of time. Total number of interactions were 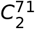.

| RANKING @ $t_i$ FOR FZD-WNT FAMILY INTERACTION | | | | | |
| --- | --- | --- | --- | --- | --- |
| $2^{nd}$ order intr. | $t_1$ | $t_3$ | $t_6$ | $t_{12}$ | $t_{24}$ |
| FZD7WNT1 | 55 | 1790 | 81 | 1241 | 817 |
| FZD5WNT1 | 58 | 2346 | 161 | 1928 | 131 |
| FZD1WNT1 | 124 | 2181 | 157 | 1473 | 195 |
| FZD5WNT3 | 164 | 913 | 1139 | 1633 | 192 |
| FZD7WNT3 | 168 | 493 | 1216 | 508 | 274 |
| FZD1WNT3 | 181 | 1834 | 909 | 831 | 681 |
| FZD6WNT1 | 267 | 803 | 1938 | 299 | 1394 |
| FZD6WNT3 | 494 | 16 | 2436 | 1371 | 776 |
| FZD8WNT3 | 500 | 127 | 1621 | 2029 | 2318 |
| FZD1WNT2B | 693 | 277 | 105 | 2424 | 1609 |
| FZD1WNT4 | 748 | 2153 | 297 | 2423 | 797 |
| FZD1WNT5A | 805 | 1924 | 307 | 1120 | 165 |
| FZD7WNT2 | 815 | 173 | 1242 | 84 | 1788 |
| FZD1WNT2 | 838 | 222 | 1540 | 235 | 1256 |
| FZD1WNT3A | 881 | 482 | 2298 | 2417 | 1337 |
| FZD7WNT5A | 887 | 1403 | 41 | 1466 | 1629 |
| FZD7WNT3A | 910 | 1757 | 705 | 2105 | 390 |
| FZD7WNT4 | 926 | 1925 | 694 | 949 | 1046 |
| FZD5WNT3A | 981 | 822 | 1946 | 2243 | 468 |
| FZD7WNT2B | 1038 | 328 | 29 | 2174 | 1849 |
| FZD5WNT5A | 1059 | 1553 | 543 | 1566 | 193 |

**Table 4.** 2*^nd^* order interaction ranking for FZD-WNT variants using HSIC-rbf for different intervals of time. Total number of interactions were 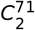.

| RANKING @ $t_i$ FOR FZD-WNT FAMILY INTERACTION | | | | | |
| --- | --- | --- | --- | --- | --- |
| FZD5WNT2 | 1157 | 1319 | 2227 | 1535 | 2036 |
| FZD5WNT4 | 1196 | 514 | 621 | 2391 | 2362 |
| FZD2WNT1 | 1208 | 1049 | 26 | 1665 | 398 |
| FZD5WNT2B | 1212 | 1377 | 94 | 1359 | 1753 |
| FZD6WNT2 | 1262 | 649 | 1471 | 648 | 275 |
| FZD8WNT4 | 1296 | 810 | 1566 | 1162 | 942 |
| FZD8WNT5A | 1316 | 73 | 334 | 1596 | 676 |
| FZD8WNT2B | 1319 | 756 | 109 | 427 | 2166 |
| FZD6WNT5A | 1320 | 1231 | 1375 | 2168 | 445 |
| FZD6WNT3A | 1364 | 102 | 2388 | 2192 | 891 |
| FZD8WNT2 | 1403 | 2424 | 952 | 968 | 772 |
| FZD8WNT3A | 1422 | 706 | 1145 | 1628 | 1499 |
| FZD6WNT4 | 1445 | 946 | 1846 | 1188 | 1696 |
| FZD6WNT2B | 1568 | 205 | 1183 | 1538 | 1922 |
| FZD2WNT3 | 1679 | 1758 | 958 | 675 | 270 |
| FZD2WNT2 | 2083 | 1807 | 758 | 357 | 1192 |
| FZD2WNT2B | 2177 | 1528 | 120 | 1316 | 763 |
| FZD2WNT4 | 2280 | 1372 | 600 | 1988 | 1835 |
| FZD2WNT3A | 2301 | 1552 | 1327 | 1302 | 1709 |
| FZD2WNT5A | 2304 | 1101 | 632 | 1964 | 190 |
| FZD8WNT1 | 147 | 748 | 173 | 6 | 1085 |

**Table 5.** 2*^nd^* order interaction ranking for FZD-WNT variants using HSIC-rbf for different intervals of time. Total number of interactions were 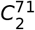.

| RANKING @ $\Delta t_{i,i+1}$ FOR FZD-WNT INTERACTION | | | | | |
| --- | --- | --- | --- | --- | --- |
| 2<sup>nd</sup> order intr. | $\Delta t_{1,3}$ | $t_{3,6}$ | $t_{6,12}$ | $t_{12,24}$ | |
| FZD5WNT2 | 26 | 897 | 1803 | 1450 |  |
| FZD5WNT4 | 49 | 463 | 945 | 2348 |  |
| FZD5WNT5A | 136 | 579 | 1528 | 2296 |  |
| FZD5WNT1 | 564 | 617 | 2284 | 1432 |  |
| FZD5WNT2B | 803 | 1896 | 808 | 1089 |  |
| FZD5WNT3A | 902 | 2287 | 1744 | 1093 |  |
| FZD6WNT5A | 1213 | 1165 | 162 | 1497 |  |
| FZD8WNT5A | 1316 | 41 | 294 | 2474 |  |
| FZD5WNT3 | 1463 | 1358 | 1232 | 1333 |  |
| FZD8WNT2 | 1499 | 186 | 1282 | 1890 |  |
| FZD7WNT5A | 1553 | 149 | 79 | 2181 |  |
| FZD1WNT5A | 1602 | 146 | 839 | 2217 |  |
| FZD7WNT4 | 1724 | 287 | 537 | 677 |  |
| FZD6WNT4 | 1786 | 1018 | 766 | 602 |  |
| FZD8WNT4 | 1803 | 124 | 1666 | 1968 |  |
| FZD1WNT4 | 1825 | 58 | 1445 | 1186 |  |
| FZD2WNT4 | 1871 | 434 | 2352 | 2388 |  |
| FZD2WNT2 | 1875 | 435 | 1451 | 1562 |  |
| FZD2WNT5A | 1908 | 420 | 397 | 1428 |  |
| FZD7WNT2 | 2041 | 299 | 326 | 811 |  |
| FZD6WNT1 | 2059 | 425 | 183 | 667 |  |

**Table 6.** 2*^nd^* order interaction ranking for FZD-WNT variants using HSIC-rbf for different intervals of time. Total number of interactions were 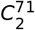.

| 2<sup>nd</sup> order intr. | $\Delta t_{1,3}$ | $t_{3,6}$ | $t_{6,12}$ | $t_{12,24}$ |
| --- | --- | --- | --- | --- |
| FZD8WNT1 | 2094 | 42 | 260 | 493 |
| FZD2WNT2B | 2101 | 1563 | 210 | 281 |
| FZD8WNT3A | 2111 | 426 | 1622 | 1988 |
| FZD7WNT3A | 2121 | 514 | 735 | 968 |
| FZD1WNT3A | 2170 | 611 | 1772 | 1213 |
| FZD2WNT1 | 2188 | 893 | 281 | 1938 |
| FZD1WNT1 | 2221 | 62 | 1237 | 1995 |
| FZD1WNT2 | 2231 | 408 | 2069 | 1664 |
| FZD6WNT3 | 2258 | 1929 | 66 | 1499 |
| FZD6WNT2B | 2341 | 2315 | 132 | 1965 |
| FZD7WNT3 | 2351 | 1322 | 12 | 678 |
| FZD8WNT2B | 2375 | 572 | 59 | 660 |
| FZD2WNT3A | 2379 | 1495 | 2310 | 1820 |
| FZD8WNT3 | 2381 | 2032 | 110 | 2309 |
| FZD7WNT1 | 2385 | 83 | 50 | 452 |
| FZD6WNT2 | 2396 | 1368 | 565 | 1119 |
| FZD7WNT2B | 2402 | 874 | 21 | 1740 |
| FZD2WNT3 | 2443 | 2319 | 54 | 1361 |
| FZD1WNT3 | 2453 | 962 | 193 | 444 |
| FZD6WNT3A | 2465 | 1127 | 2473 | 1208 |
| FZD1WNT2B | 2471 | 694 | 516 | 191 |

Dijksterhuis *et al*.^71^ conclude in their wet lab findings that *WNT* — 3*A/* 4/ 5*A* functionally bind to *FZD* — 2/ 4/ 5. Of the recorded rankings of *FZD*2 with the above *WNT*s, the combination of *FZD*2-*WNT* — 4/ 5*A* showed a moderately high rank of around 600 in 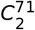 combinations, at time point *t*_6_. Additionally, the pipeline also generated higher ranking for interactions *FZD*2-*WNT* — 2/ 2*B/* 3 at other time points in the later stage of the pathway (table 4). *FZD*5 also showed confirmatory results with *WNT* — 3*A/* 4/ 5 (tables 3 and 4). Rankings of these interactions are relatively moderate/weak but show promise in the fact that molecules do bind during the signaling process. Apart from these interactions, other recorded rankings for *FZD*5 that showed high affinity for variants of WNTs were *FZD*5-*WNT* — 1/ 2*B/* 3 (the higer ranks in the aforementioned tables). *FZD*5-*WNT*2 did not show that high ranking in comparison to their other counterparts. Similarly, *FZD*7 showed high binding affinity to *WNT* − 1/2/2*B/*3/5*A/* at different time points and lower or weaker affinity for *WNT* − 3*A/*4. *FZD*8 showed high binding affinity to *WNT* − 1/2*B/*3/5*A* at different time points and lower or weaker affinity for *WNT* − 2/3*A/*4. *FZD*1 showed high binding affinity to *WNT* − 1/2/2*B/*3/3*A/*4/5*A* at different time points. Finally, *FZD*6 showed strong affinity at singular time points with *WNT* 1/2/2*B/*3/3*A/*5*A* and overall relatively moderate or weak affinity to *WNT* 3*A/* 4/ 5*A*. Similar confirmatory results of binding have been found in wet lab tests in Ueno *et al*.^72^ and Vincan *et al*.^73^. Finally, similar interpretations can be made for the *FZD*-*WNT* interactions for the different durations or time periods as has been shown in tables 5 and 6.

The appendix contains a list of rankings for 2*^n^d* order combinations for different factors at different time points and time durations. These contain the unexplored biological hypotheses for Wnt signaling pathway.

## Conclusion

COMPUTATIONAL - A pipeline has been developed to prioritise an *n^th^* order combination of factors that affect a signaling pathway. It takes into account the sensitivity indices computed from variance based (SOBOL) and density-kernel based (HSIC) methods to estimate the influence of each factor or combination of factors. These are then fed as feature vectors into a powerful support vector ranking algorithm that produces a ranked list of the interactions/combinations. The pipeline is aimed to (1) reveal important combinations that form unexplored biological hypotheses (2) study the behaviour of these combinations in time (3) reduce the cost in time/energy for the biologists by providing a prioritized list in a vast combinatorial search space (4) reveal target sites for therapeutic interventions and (5) facilitate in conducting system wide analysis via observation of changing rankings in time.

BIOLOGICAL - The pipeline provides a prioritised list of important 2^*nd*^ order combinations of a range of family of genes involved in the Wnt pathway. More specifically, it reveals the various unexplored *FZD*-*WNT* combinations that have been untested till now in the pathway. Due to paucity of space, an extensive prioritised list has been provided in the appendix for wet lab testing.

## Acknowledgement

This work was conducted independently over a period of four years and the author is grateful to the generous support of Mrs. Rita Sinha and Mr. Prabhat Sinha for the financial help they provided during these years.

## Appendix

### 6.1 Sensitivity indices

Given the range of estimators available for testing the sensitivity, it might be useful to list a few which are going to be employed in this research study. Also, a brief introduction into the fundamentals of the derivation of the three main indicies has been provided.

#### 6.1.1 Variance based indices

The variance based indices as proposed by Sobol’^21^ prove a theorem that an integrable function can be decomposed into summands of different dimensions. Also, a Monte Carlo algorithm is used to estimate the sensitivity of a function apropos arbitrary group of variables. It is assumed that a model denoted by function *u* = *ƒ*(*X*), *X* = (*X*_1_, *x*_2_, *…, X_n_*), is defined in a unit *n*-dimensional cube *K^n^* with *u* as the scalar output. The requirement of the problem is to find the sensitivity of function *ƒ* (*x*) with respect to different variables. If *u*^*^ **=** *ƒ* (*x*^*^) is the required solution, then the sensitivity of *u*^*^ apropos *x_k_* is estimated via the partial derivative (*∂u/∂X_k_*)_*x*=*x*\*_. This approach is the local sensitivity. In global sensitivity, the input *x* **=** *x*^*^ is not specified. This implies that the model *ƒ*(*x*) lies inside the cube and the sensitivity indices are regarded as tools for studying the model instead of the solution. Detailed technical aspects with examples can be found in Homma and Saltelli^33^ and Sobol^74^.

Let a group of indices *i*_1_, *i*_2_,…, *i_s_* exist, where 1 ≤ *i*_1_ <…< *i_s_* ≤ *n* and 1 ≤ *s* ≤ *n*. Then the notation for sum over all different groups of indices is -

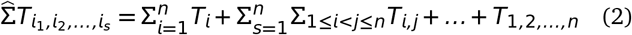

Then the representation of *ƒ*(*x*) using equation 2 in the form -

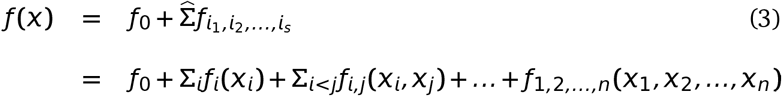

is called ANOVA-decomposition from Archer *et al*.^46^ or expansion into summands of different dimensions, if *ƒ*_0_ is a constant and integrals of the summands *ƒ*_*i*_1_, *i*_2_,…,*i*_s__ with respect to their own variables are zero, i.e,

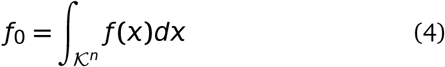

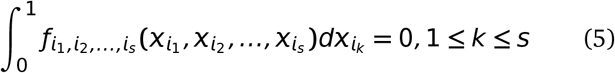

It follows from equation 4 that all summands on the right hand side are orthogonal, i.e if at least one of the indices in *i*_1_, *i*_2_, *…, i_s_* and *j*_1_, *j*_2_, *…, j_l_* is not repeated i.e

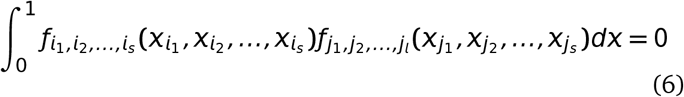

Sobol’^21^ proves a theorem stating that there is an existence of a unique expansion of equation 4 for any *ƒ*(*x*) integrable in *K^n^*. In brief, this implies that for each of the indices as well as a group of indices, integrating equation 4 yields the following –

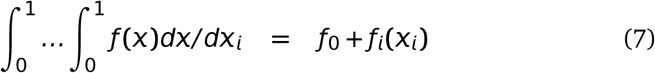

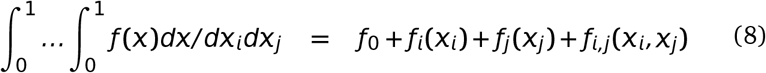

were, *dx/dx_i_* is ∏_∀*k*∈{1,…,*n*};*i*∉ *k*_ dx_*k*_ and *dx/dx_i_dx_j_* is ∏_∀*k*∈{1,…,*n*};*i*∉ *k*_ dx_k_. For higher orders of grouped indices, similar computations follow. The computation of any summand *ƒ_i_*_1_, *i*2, *…,i_s_* (*x_i_*_1_, *x_i_*_2_, *…, X_is_*) is reduced to an integral in the cube *K^n^*. The last summand *ƒ*_1,2,*…,n*_(*x*_1_, *x*_2_, *…, X_n_*) is *ƒ* (*x***) –** *ƒ*_0_ from equation 4. Homma and Saltelli^33^ stresses that use of Sobol sensitivity indices does not require evaluation of any *ƒ_i_*_1, *i*2, *…,is*_ (*x_i_*_1_, *x_i_*_2_, *…, X_is_*) nor the knowledge of the form of *ƒ*(*x*) which might well be represented by a computational model i.e a function whose value is only obtained as the output of a computer program.

Finally, assuming that *ƒ*(*x*) is square integrable, i.e *ƒ*(*x*) ∈ *L*_2_, then all of *ƒ_i_*_1, *i*2, *…,is*_ (*x_i_*_1_, *x_i_*_2_, *…, X_is_*) ∈ *L*_2_. Then the following constants

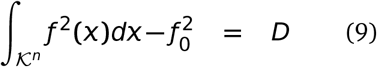

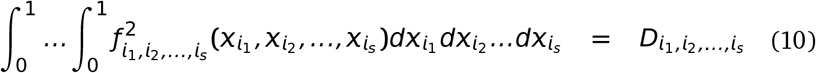

are termed as variances. Squaring equation 4, integrating over and using the orthogonality property in equation 6, *D* evaluates to -

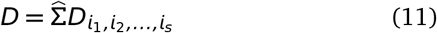

Then the global sensitivity estimates is defined as –

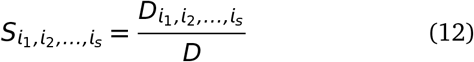

It follows from equations 11 and 12 that

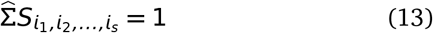

Clearly, all sensitivity indices are non-negative, i.e an index *S_i_*_1, *i*2, *…,is*_ = 0 if and only if *ƒ_i_*_1, *i*2, *…,is*_ ≡ 0. The true potential of Sobol indices is observed when variables *x*_1_, *x*_2_, *…, x_n_* are divided into *m* different groups with *y*_1_, *y*_2_, *…, y_m_* such that *m < n*. Then *ƒ*(*x*) ≡ *ƒ*(*y*_1_, *y*_2_, …, *y_m_*). All properties remain the same for the computation of sensitivity indices with the fact that integration with respect to *y_k_* means integration with respect to all the *x_i_*’s in *y_k_*. Details of these computations with examples can be found in Sobol^74^. Variations and improvements over Sobol indices have already been stated in section 2.1.

#### 6.1.2 Density based indices

As discussed before, the issue with variance based methods is the high computational cost incurred due to the number of interactions among the variables. This further requires the use of screening methods to filter out redundant or unwanted factors that might not have significant impact on the output. Recent work by Da Veiga^57^ proposes a new class of sensitivity indicies which are a special case of density based indicies Borgonovo^54^. These indicies can handle multivariate variables easily and relies on density ratio estimation. Key points from Da Veiga^57^ are mentioned below.

Considering the similar notation in previous section, *ƒ*: *R^n^* → *R* (*u* **=** *ƒ*(*x*)) is assumed to be continuous. It is also assumed that *x_k_* has a known distribution and are independent. Baucells and Borgonovo^75^ state that a function which measures the similarity between the distribution of *U* and that of *U|X_k_* can define the impact of *x_k_* on *U*. Thus the impact is defined as –

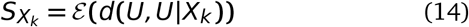

were *d*(⋅, ⋅) is a dissimilarity measure between two random variables. Here *d* can take various forms as long as it satisfies the criteria of a dissimilarity measure. Csiszár *et al*.^60^’s f-divergence between *U* and *U |X_k_* when all input random variables are considered to be absolutely continuous with respect to Lebesgue measure on *R* is formulated as –

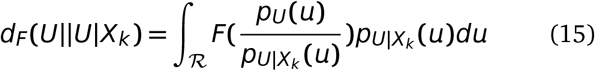

were *F* is a convex function such that *F*(1) = 0 and *p_U_* and *pU |x_k_* are the probability distribution functions of *U* and *U| X_k_*. Standard choices of *F* include Kullback-Leibler divergence *F*(*t*) = –log_*e*_(*t*), Hellinger distance **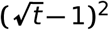**, Total variation distance *F*(*t***) =** |*t* − 1|, Pearson *χ*^2^ divergence *F*(*t***) =** *t*^2^ − 1 and Neyman *χ*^2^ divergence *F*(*t*) = (1 − *t*^2^)*/t*. Substituting equation 15 in equation 14, gives the following sensitivity index -

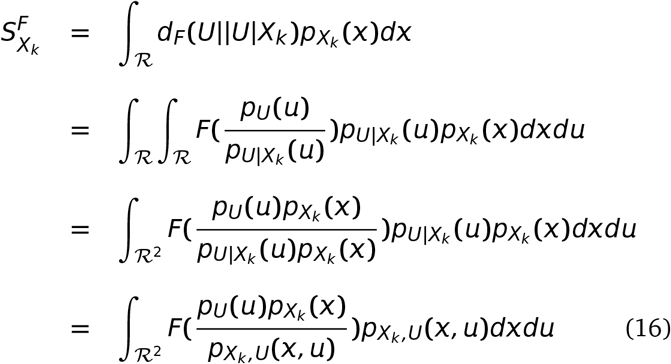

were *px_k_* and *px_k_,y* are the probability distribution functions of *x_k_* and (*x_k_, U*), respectively. Csiszár *et al*.^60^ f-divergences imply that these indices are positive and equate to 0 when *U* and *x_k_* are independent. Also, given the formulation of 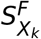, it is invariant under any smooth and uniquely invertible transformation of the variables *x_k_* and *U* (Kraskov *et al*.^69^). This has an advantage over Sobol sensitivity indices which are invariant under linear transformations.

By substituting the different formulations of *F* in equation 16, Da Veiga^57^’s work claims to be the first in establishing the link that previously proposed sensitivity indices are actually special cases of more general indices defined through Csiszár *et al*.^60^’s f-divergence. Then equation 16 changes to estimation of ratio between the joint density of (*x_k_, U*) and the marginals, i.e -

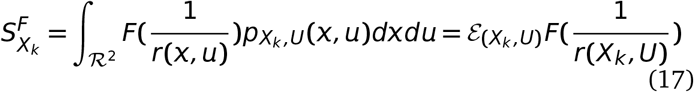

were, *r*(*x, y*) = (*p_Xk,U_* (*x, u*))/ (*p_U_* (*u*)*p_Xk_* (*x*)). Multivariate extensions of the same are also possible under the same formulation.

Finally, given two random vectors *x* ∈ *𝓡^p^* and *Y* ∈ *𝓡^q^*, the dependence measure quantifies the dependence between with the property that the measure equates to 0 if and only if *x* and *Y* are independent. These measures carry deep links (Sejdinovic *et al*.^77^) with distances between embeddings of distributions to reproducing kernel Hilbert spaces (RHKS) and here the related Hilbert-Schmidt independence criterion (HSIC by Gretton *et al*.^59^) is explained.

In a very brief manner from an extremely simple introduction by Daumé III^78^ -”We first defined a field, which is a space that supports the usual operations of addition, subtraction, multiplication and division. We imposed an ordering on the field and described what it means for a field to be complete. We then defined vector spaces over fields, which are spaces that interact in a friendly way with their associated fields. We defined complete vector spaces and extended them to Banach spaces by adding a norm. Banach spaces were then extended to Hilbert spaces with the addition of a dot product.” Mathematically, a Hilbert space *𝓗* with elements *r, s* ∈ *𝓗* has dot product 〈*r, s*〉_*𝓗*_ and *r* • *s*. When *𝓗* is a vector space over a field *𝓕*, then the dot product is an element in *𝓕*. The product 〈*r, s*〉_*𝓗*_ follows the below mentioned properties when *r, s, t* ∈ *𝓗* and for all *a* ∈ *𝓕* -

- Associative: (*ar*) • *s* = *a*(*r* • *s*)
- Commutative: *r* • *s* = *s* • *r*
- Distributive: *r* • (*s* **+** *t*) = *r* • *s* **+** *r* • *t*

Given a complete vector space *V* with a dot product 〈 •, • 〉, the norm on *V* defined by ||*r*|| *_V_* = 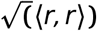 makes this space into a Banach space and therefore into a full Hilbert space.

A reproducing kernel Hilbert space (RKHS) builds on a Hilbert space *𝓗* and requires all Dirac evaluation functionals in *𝓗* are bounded and continuous (on implies the other). Assuming *𝓗* is the *L*_2_ space of functions from *x* to *R* for some measurable, *x* is a vector and *g* a function which maps from this *x*. For an element *x* ∈ *x*, a Dirac evaluation functional at x is a functional *δ_x_* ∈*𝓗* such that *δ_x_*(*g***) =** *g*(*x*). For the case of real numbers, X. is a vector and *g* a function which maps from this vector space to *R*. Then *δ_x_* is simply a function which maps *g* to the value *g* has at *x*. Thus, *δ_x_* is a function from (*𝓡^n^* ↦ *𝓡*) into *𝓡*.

The requirement of Dirac evaluation functions basically means (via the Riesz^79^ representation theorem) if *ϕ* is a bounded linear functional (conditions satisfied by the Dirac evaluation functionals) on a Hilbert space *𝓗*, then there is a unique vector *ℓ* in *𝓗* such that *ϕg* =〈 *g, ℓ* 〉_*𝓗*_ for all *ℓ*∈ *H*. Translating this theorem back into Dirac evaluation functionals, for each *δ_x_* there is a unique vector *k_x_* in *𝓗* such that *δ_x_g* = *g*(*x*) = 〈 *g, k_x_* 〉 _*H*_. The reproducing kernel *K* for *𝓗* is then defined as: *K*(*x, x*′) = 〈 *k_X_, k_X_*〉, were *k_X_* and *k_X_*′ are unique representatives of *δ_X_* and *δ_X_*′. The main property of interest is 〈*g*, *K*(*X, X*^I^) 〉 _*H*_ = *g* (*X* Furthermore, *k_x_* is defined to be a function *y* ↦ *K*(*X, y*) and thus the reproducibility is given by 〈*K*(*X*, •), *K*(*y*, •)〉_*𝓗*_ = *K*(*X, y*).

Basically, the distance measures between two vectors represent the degree of closeness among them. This degree of closeness is computed on the basis of the discriminative patterns inherent in the vectors. Since these patterns are used implicitly in the distance metric, a question that arises is, how to use these distance metric for decoding purposes?

The kernel formulation as proposed by Aizerman *et al*.^61^, is a solution to our problem mentioned above. For simplicity, we consider the labels of examples as binary in nature. Let **x**_*i*_ ∈*𝓡^n^*, be the set of *n* feature values with corresponding category of the example label (*y_i_*) in data set *D*. Then the data points can be mapped to a higher dimensional space *𝓗* by the transformation *ϕ*:

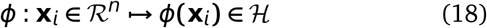

This *𝓗* is the *Hilbert Space* which is a strict inner product space, along with the property of completeness as well as separability. The inner product formulation of a space helps in discriminating the location of a data point w.r.t a separating hyperplane in *𝓗*. This is achieved by the evaluation of the inner product between the normal vector representing the hyperplane along with the vectorial representation of a data point in *𝓗*. Thus, the idea behind equation (18) is that even if the data points are nonlinearly clustered in space *R^n^*, the transformation spreads the data points into *𝓗*, such that they can be linearly separated in its range in *𝓗*.

Often, the evaluation of dot product in higher dimensional spaces is computationally expensive. To avoid incurring this cost, the concept of kernels in employed. The trick is to formulate kernel functions that depend on a pair of data points in the space *𝓡^n^*, under the assumption that its evaluation is equivalent to a dot product in the higher dimensional space. This is given as:

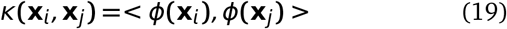

Two advantages become immediately apparent. First, the evaluation of such kernel functions in lower dimensional space is computationally less expensive than evaluating the dot product in higher dimensional space. Secondly, it relieves the burden of searching an appropriate transformation that may map the data points in *𝓡^n^* to *𝓗*. Instead, all computations regarding discrimination of location of data points in higher dimensional space involves evaluation of the kernel functions in lower dimension. The matrix containing these kernel evaluations is referred to as the *kernel* matrix. With a cell in the kernel matrix containing a kernel evaluation between a pair of data points, the kernel matrix is square in nature.

As an example in practical applications, once the kernel has been computed, a pattern analysis algorithm uses the kernel function to evaluate and predict the nature of the new example using the general formula:

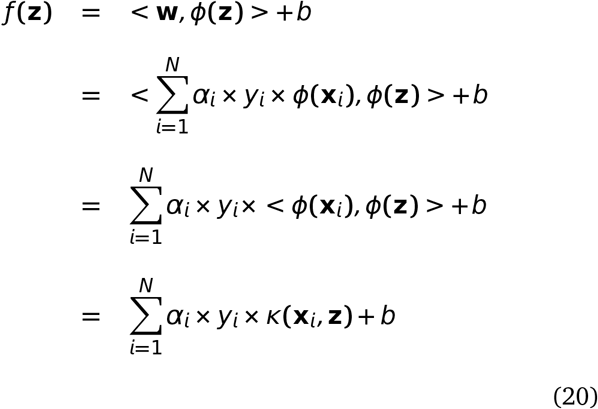

where **w** defines the hyperplane as some linear combination of training basis vectors, **z** is the test data point, *y_i_* the class label for training point **x**_*i*_, *α_i_* and *b* are the constants. Various transformations to the kernel function can be employed, based on the properties a kernel must satisfy. Interested readers are referred to Taylor and Cristianini^80^ for description of these properties in detail.

The Hilbert-Schmidt independence criterion (HSIC) proposed by Gretton *et al*.^59^ is based on kernel approach for finding dependences and on cross-covariance operators in RKHS. Let *X* ∈ *X* have a distribution *P_X_* and consider a RKHS *A* of functions *X* → *𝓡* with kernel *k_X_* and dot product 〈 •, • 〉_*A*_. Similarly, Let *U* ∈ *Y* have a distribution *P_Y_* and consider a RKHS *B* of functions *U* →*𝓡* with kernel *k_B_* and dot product, 〈 •, • 〉_*B*_. Then the cross-covariance operator *C_X,U_* associated with the joint distribution *P_XU_* of (*X, U*) is the linear operator *B* → *A* defined for every *a* ∈ *A* and *b* ∈ *B* as -

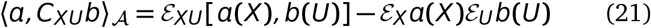

The cross-covariance operator generalizes the covariance matrix by representing higher order correlations between *X* and *U* through nonlinear kernels. For every linear operator *C*: *B* → *A* and provided the sum converges, the Hilbert-Schmidt norm of *C* is given by -

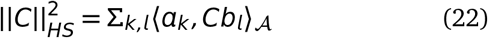

were *a_k_* and *b_l_* are orthonormal bases of *A* and *B*, respectively. The HSIC criterion is then defined as the Hilbert-Schmidt norm of cross-covariance operator –

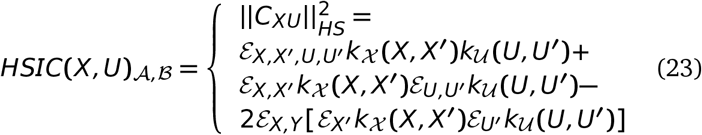

were the equality in terms of kernels is proved in Gretton *et al*.^59^. Finally, assuming (*X_i_, U_i_*) (*i* **=** 1, 2, *…, n*) is a sample of the random vector (*X, U*) and denote *K_X_* and *K_U_* the Gram matrices with entries *K_X_* (*i, j*) = *k_X_* (*X_i_, X_j_*) and *K_U_* (*i, j*) = *k_U_* (*U_i_, U_j_*). Gretton *et al*.^59^ proposes the following estimator for *HSIC_n_*(*X, U*)_*A,B*_ –

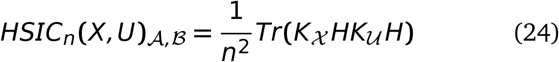

were *H* is the centering matrix such that *H*(*i,j*) = *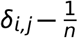*. Then *HSIC_n_*(*X, U*)_*A,B*_ can be expressed as -

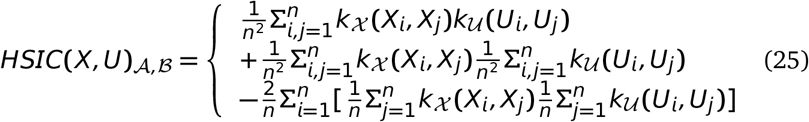

Finally, Da Veiga^57^ proposes the sensitivity index based on distance correlation as -

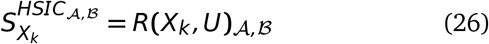

were the kernel based distance correlation is given by –

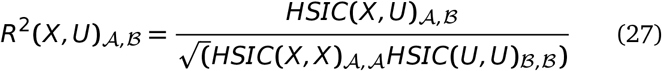

were kernels inducing *A* and *B* are to be chosen within a universal class of kernels. Similar multivariate formulation for equation 24 are possible.

#### 6.1.3 Choice of sensitivity indices

The SENSITIVITY PACKAGE (Faivre *et al*.^81^ and Iooss and Lemaître^22^) in R langauge provides a range of functions to compute the indices and the following indices will be taken into account for addressing the posed questions in this manuscript.

1. sensiFdiv - conducts a density-based sensitivity analysis where the impact of an input variable is defined in terms of dissimilarity between the original output density function and the output density function when the input variable is fixed. The dissimilarity between density functions is measured with Csiszar f-divergences. Estimation is performed through kernel density estimation and the function kde of the package ks. (Borgonovo^54^, Da Veiga^57^)
2. sensiHSIC - conducts a sensitivity analysis where the impact of an input variable is defined in terms of the distance between the input/output joint probability distribution and the product of their marginals when they are embedded in a Reproducing Kernel Hilbert Space (RKHS). This distance corresponds to HSIC proposed by Gretton *et al*.^59^ and serves as a dependence measure between random variables.
3. soboljansen - implements the Monte Carlo estimation of the Sobol indices for both first-order and total indices at the same time (all together 2p indices), at a total cost of (p+2) × n model evaluations. These are called the Jansen estimators. (Jansen^49^ and Saltelli *et al*.^41^)
4. sobol2002 - implements the Monte Carlo estimation of the Sobol indices for both first-order and total indices at the same time (all together 2p indices), at a total cost of (p+2) × n model evaluations. These are called the Saltelli estimators. This estimator suffers from a conditioning problem when estimating the variances behind the indices computations. This can seriously affect the Sobol indices estimates in case of largely non-centered output. To avoid this effect, you have to center the model output before applying “sobol2002”. Functions “soboljansen” and “sobolmartinez” do not suffer from this problem. (Saltelli^35^)
5. sobol2007 - implements the Monte Carlo estimation of the Sobol indices for both first-order and total indices at the same time (all together 2p indices), at a total cost of (p+2) × n model evaluations. These are called the Mauntz estimators. (Saltelli and Annoni^48^)
6. sobolmartinez - implements the Monte Carlo estimation of the Sobol indices for both first-order and total indices using correlation coefficients-based formulas, at a total cost of (p+ 2) × n model evaluations. These are called the Martinez estimators.
7. sobol - implements the Monte Carlo estimation of the Sobol sensitivity indices. Allows the estimation of the indices of the variance decomposition up to a given order, at a total cost of (N + 1) × n where N is the number of indices to estimate. (Sobol’^21^)

### 7 Optimization and Support Vector Machines

Aspects of SVMs from Sinha^75^ are reproduced for completeness.

#### 7.1 Optimization Problems

##### 7.1.1 Introduction

The main focus in this section is optimization problems, the concept of Lagrange multipliers and KKT conditions, which will be later used to explain the details about the SVMs.

##### 7.1.2 Mathematical Formulation

Optimization problems arise in almost every area of engineering. The goal is to achieve an almost perfect and efficient result, while carrying out certain procedures of optimization. Our main source of reference on this topic derives from Bletzinger^83^. We will be using notations used in Bletzinger^83^. In mathematical terms the general form of optimization problem can be represented as:

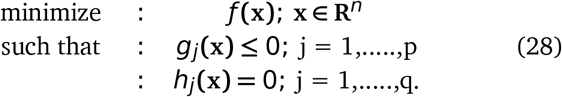

where *ƒ, g_j_* and *h_j_* are the objective function, equality constraints and inequality constraints. Generally, the number of constraints is less than the number of variables used to formulate the optimization problem. For a problem to be linear, both the constraints and the objective function need to be linear. Quadratic problems require only the objective function to be quadratic, while the constraints remain linear in formulation. Besides these, if any one of the functions is nonlinear, then the problem becomes nonlinear in nature. A graphical view of the types of the problems can be seen in fig. 8).

**Fig. 8.**
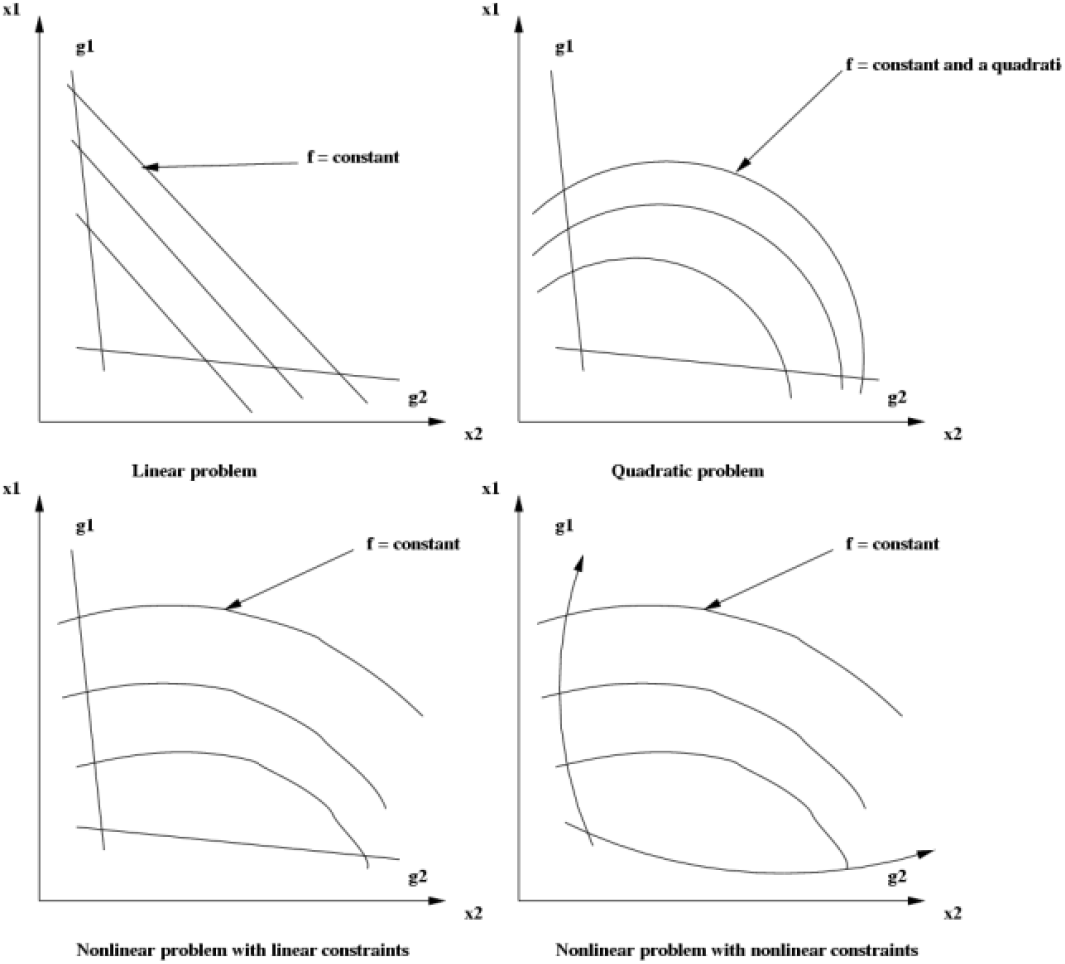
Kinds of optimization problems.

##### 7.1.3 Lagrange Multipliers

In unconstrained optimization problems, where the first order derivatives are assumed continuous, the solution is found by solving:

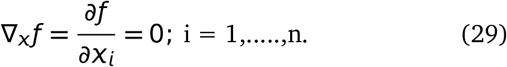

where *ƒ* is a function of *X*. Since most of the optimization problems are constrained, the concept of Lagrange multipliers is introduced in order to solve the problem. Thus, the Lagrangian formulation, for Eqn. 28 becomes:

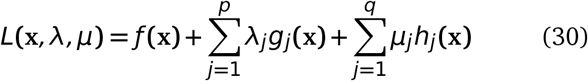

where *L* is the Lagrangian, *λ* and *µ* are the vectors of the Lagrange multipliers for inequality and equality constraints, respectively.

Next comes the solving of the Lagrangian. We try to derive a solution in terms of variables used and show that the final solution achieved by Equ. 28 and Eqn. 30 remains the same. For the sake of derivation, we assume that each of the vectors *x*, *λ* and *µ* have a single element and also there exists a single optimal solution. We will then generalize the solution to vectors containing various elements. Let *x*^*^, *λ*^*^ and *µ*^*^ be the optimal solution for the Lagrangian. Let *x*^!^ be the optimal solution for *ƒ*(*x*). To begin with, our Lagrangian has the form:

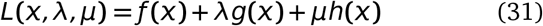

###### Derivation

- **Step 1:** Differentiate the Lagrangian in Eqn 31 w.r.t *x* and equate it to zero.

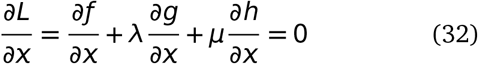
- **Step 2:** Find *x* in terms of *λ* and *µ*, such that *x* **=** *x*(*λ, µ*).
- **Step 3:** Differentiate the Lagrangian in Eqn. 31 w.r.t *λ* and equate it to zero.

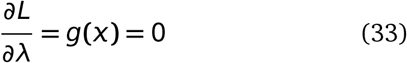
- **Step 4:** Differentiate the Lagrangian in Eqn 31 w.r.t *µ* and equate it to zero.

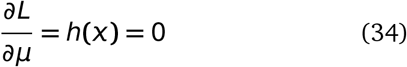
- **Step 5:** Substitute *x*(*λ, µ*) in Equ. 33 and Eqn. 34 to get two equations in two unknowns *λ* and *µ* and solve to get the optimal values.

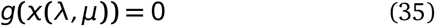

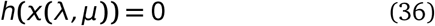 Let *λ*^*^, *µ*^*^ be the solution. Substituting these in *x* = *x*(*λ, µ*), we get *x*^*^.
- **Step 6:** Combining Eqn. 33 and Eqn. 34 in Eqn. 31, along with *λ*^*^, *µ*^*^ and *x*^*^, we have:

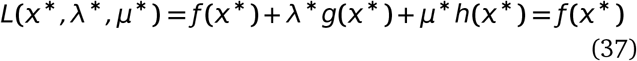 Since, it is assumed that there exist only one optimal solution we have:

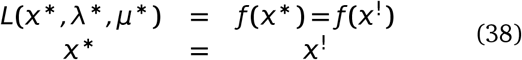

Lastly, since *g*(*x*) in Eqn. 33 is a inequality constraint, we have:

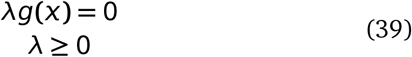

##### 7.1.4 Dual Functions

For sake of simplicity, let us for a moment ignore the equality constraint. Then the Lagrangian becomes:

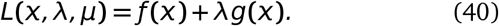

It is sometimes easy to transform the Lagrangian into a simpler form, in order to find an optimal solution. We can represent the Lagrangian as a Dual function in such a manner that the optimal solution defined as minimum of *L*(*x, λ*^*^) w.r.t *x* where *λ* **=** *λ*^*^, can be represented as the maximum of dual function *D*(*λ*) w.r.t *λ*. For a given *λ*, the dual is evaluated by finding the minimum of *L*(*x, λ*) w.r.t *x*. Thus to find the optimal point we evaluate:

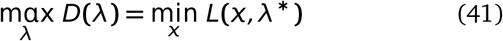

So the basic steps to solve the dual problem are as follows: **Step 1:** Minimize *L*(*x, λ*) w.r.t *x*, and find *x* in terms of *λ*. **Step 2:** Substitute *x*(*λ*) in *L* s.t. *D*(*λ***) =** *L*(*x*(*λ*), *λ*). **Step 3:** Maximize *D*(*λ*) w.r.t *λ*.

##### 7.1.5 Karush Kuhn Tucker Conditions

The derivation in the last part (Eqn. 31 to Eqn. 39)gives us a set of equations that need to be evaluated along with the consideration of constraints present. These set of equations and constraints in terms of the Lagrangian, form the Karush Kuhn Tucker Conditions. We give here the generalized KKT conditions and explain the necessary details

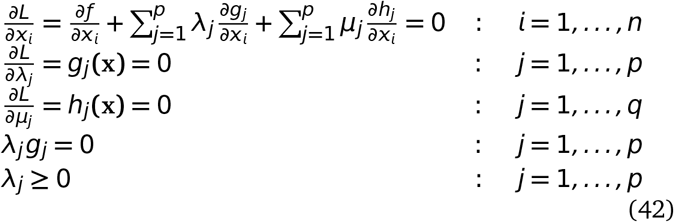

where *L* is Eqn. 30.

The KKT conditions specify a few points which are as follows:

1. The first line states that the linear combination of objective and constraint gradients vanishes.
2. A prerequisite of the KKT conditions is that the gradients of the constraints must be continuous (evident from second and third lines in Eqn. 42).
3. The last two lines in Eqn. 42 state that at optimum either the constraints are active or the constrains are inactive.

#### 7.2 Support Vector Machines

Armed with the knowledge of optimization problems and concept of Lagrange multipliers, we now delve into the workings of support vector machines. Burges^84^ provides a good introduction to SVMs and is our main reference. Interested readers should refer to Cristianini and Shawe-Taylor^85^, Schölkopf and Smola^86^ and Vapnik and Vapnik^87^ for detailed references.

##### 7.2.1 Separable Case

Let us suppose that we are presented with a data set that is linearly separable. We assume that there are *m* examples of data in the format {**x**_*i*_, *y_i_*}, s.t. **x**_*i*_ ∈ **R**^*n*^; *i* = 1, …, *m*, where *y_i_* ∈ {−1, 1} is the corresponding true label of **x**_*i*_. We also suppose there is an existence of a linear hyperplane in the *n* dimensional space that separates the positively labeled data from the negatively labeled data. Let this separating hyperplane be given by

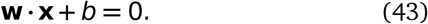

where, **w** is the normal vector to the hyperplane and |*b*| / ‖**w**‖ is the shortest perpendicular distance of the hyperplane to the origin. ||**w**|| is the Euclidean norm of **w**. The *margin* of a hyperplane is then defined as the minimum of the distance of the positively and negatively labeled examples, to the hyperplane. For the linear case, the SVM searches for the hyperplane with largest margin. We now have three conditions, based on the location of an example **x**_*i*_ w.r.t the hyperplane:

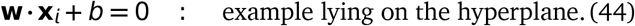

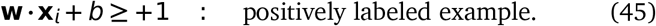

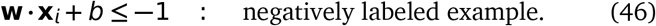

Combining the equality and the two inequalities we have:

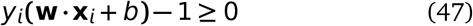

Since the SVMs search for the largest margin, we now try to find a mathematical expression of the margin. Considering the examples that satisfy equality in Eqn. 45, the distance of the closest positive example can be expressed as |1 − *b*| / ||**w**||. Similarly, considering the negative examples that satisfy equality in Eqn. 46, the distance of the closest negative example can be expressed as | −1 −*b*|/||**w**||. On summation of the two shortest distances, we get the margin of the hyperplane as 2/||**w**||. Since the labels are {−1, 1}, no example lies inside the hyperplanes representing the margin in this case. Taking into account that the SVM searches for the largest margin, we can say that it can be achieved by minimizing ||**w**||^2^, subject to the constraints in Eqn. 47. Examples lying on the hyperplanes of the margins are termed *support vectors*, as their removal would change the margin and thus the solution. Figure 9 represents the conceptual points about separating hyperplanes.

**Fig. 9.**
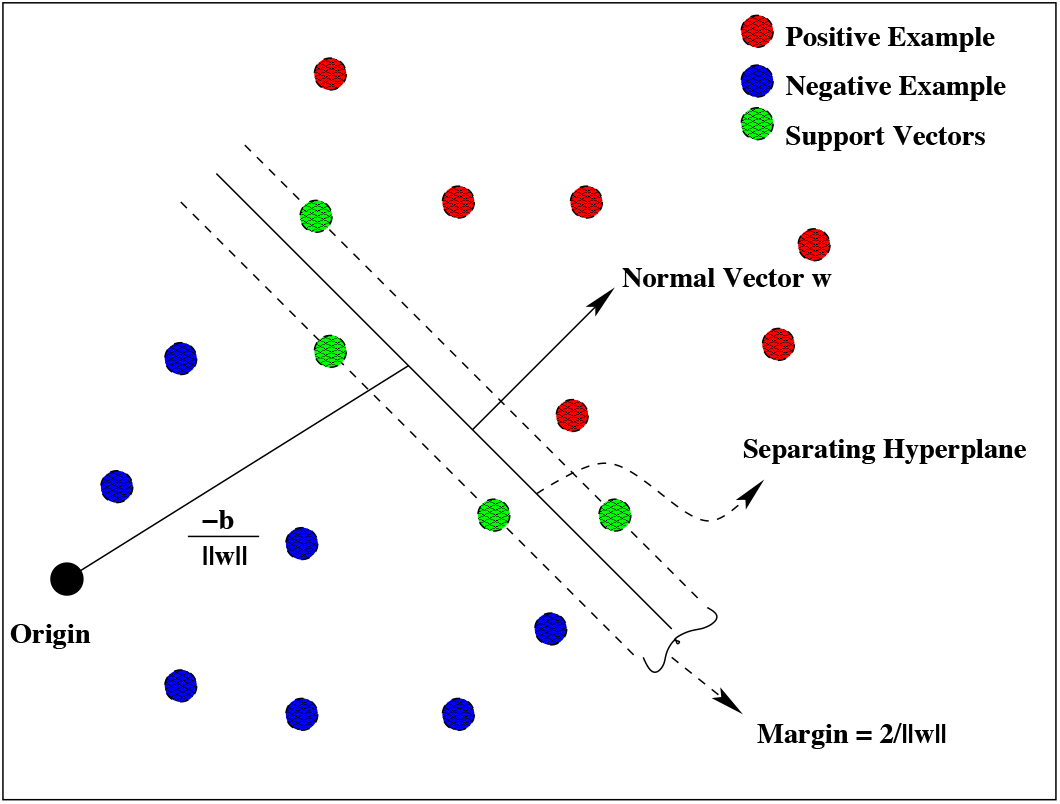
Linear hyperplane for separable data. Adapted from Burges (A Tutorial on Support Vector Machines)

#### 7.3 Lagrangian Representation: Separable Case

Clearly, the previous paragraph shows that finding the margin is a problem of optimization as the goal is to minimize ||**w**||^2^ subject to constraints in Eqn. 47. Employing the ideas of Chapter 3, the Lagrangian for the above problem, is:

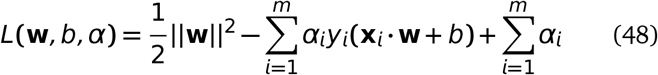

where 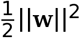 is the objective function, *α* is the Lagrangian multiplier and the Eqn. 47 is the inequality constraint. Since the minimization of the objective function is required, we employ the ideas of the derivation of KKT conditions (Eqn. 42) to Eqn. 48. In short, we would require the *L*(**w**, *b, α*) to be minimized w.r.t **w** and *b* and also require its derivative w.r.t all *α_i_*’s to vanish. Thus the KKT conditions take the form:

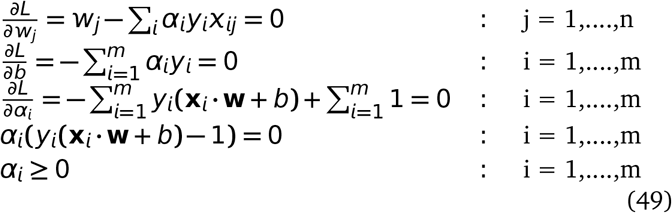

Thus solving the SVMs is equivalent to solving the KKT conditions. While **w** is determined by the training set, *b* can be found by solving the penultimate equation in Eqn. 49 for which *α_i_* ≠ 0. Also note that examples that have *α_i_* ≠ 0 form the set of support vectors.

The dual problem for the same Lagrangian is:

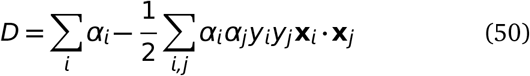

Solving for Eqn. 50 requires maximization of *D* w.r.t *α_i_*, subject to second line of Eqn. 49 and positivity of *α_i_*, with the solution given by first line of Eqn. 49.

To classify or predict the label of a new example **x**_*new*_, the SVM has to evaluate (**x**_*new*_ • **w** + *b*) and check the sign of the evaluated value. A positive sign would lead to assignment of a +1 label and a negative sign to −1.

#### 7.4 Nonseparable Case

For many classification problems, the data present is nonseparable. To extend the idea to nonseparable case, some amount of cost is added, which takes care of particular cases of examples. This is achieved by introducing slack in the constraints Eqn. 45 and Eqn. 46 (Burges^84^, Vapnik and Vapnik^87^). The equations then becomes

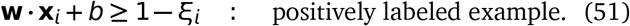

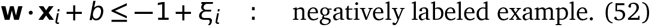

For an error to occur, the *ξ_i_* value must exceed unity. To take care of the cost of errors, a penalty is introduced which changes the objective function from ||**w**||^2^ / 2 to ||**w**||^2^ / 2 **+** *C*(Σ_*i*_ *ξ_i_*)^*k*^. Thus Σ*i ξ_i_* represents the upper bound on the training error. For quadratic problems, *k* can be 1 or 2.

#### 7.5 Lagrangian Representation: Nonseparable Case

Since the formulation of the Lagrangian and its dual follow the same procedure, as mentioned before, we only mention the equations. The Lagrangian for nonlinear nonseparable case is:

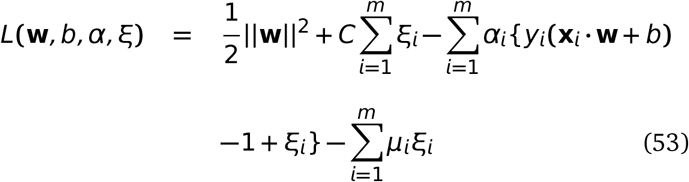

The corresponding KKT conditions are:

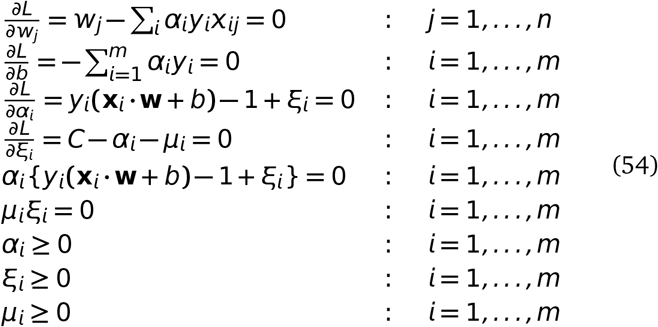

The dual formulation *k* = 1 for the Lagrangian just discussed is:

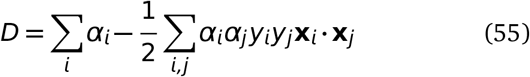

All the previous conditions remain same, except that the Lagrangian multiplier *α_i_* now has a upper bound of value *C*. The solution for the dual is given by 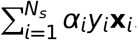. *N_s_* is the number of support vectors. Figure 10 depicts the nonseparable case.

**Fig. 10.**
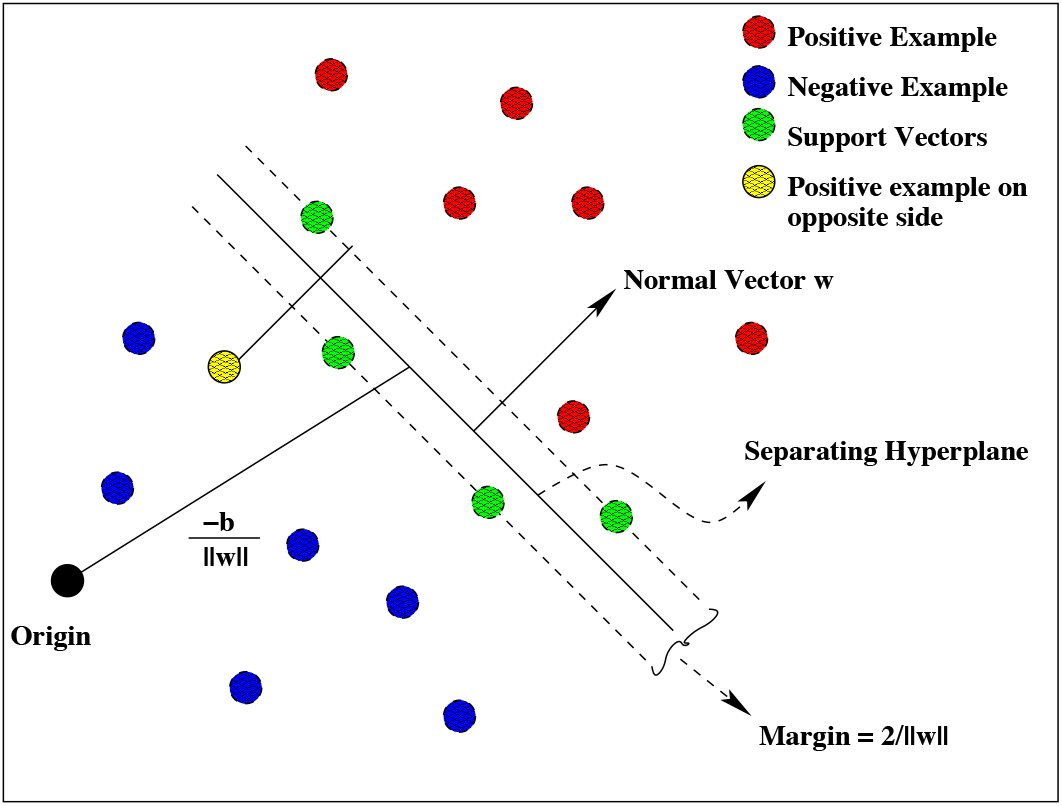
Linear hyperplane for nonseparable data. Adapted from Burges (A Tutorial on Support Vector Machines)

#### 7.6 Kernels and Space Dimensionality Transformation

The above cases were for linear separating hyperplanes. In order to generalize for nonlinear cases, Boser et.al^88^ employed the idea of Aizerman^61^ as follows; Since the Dual in Eqn. 55 and its corresponding constraint equations employ the dot product of the examples, **x**_*i*_ • **x**_*j*_, it was proposed to map the data in a higher dimensional space using a function *ϕ* s.t. the algorithm would depend only on dot products in the higher space. Next, the existence of a function called *kernel*, dependent on **x**_*i*_ and **x**_*j*_, was assumed s.t. the value reported by the kernel was equal to the value resulting from the dot product in the higher space. A mathematical representation of the above concept is –

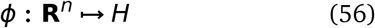

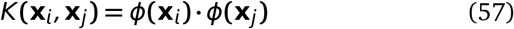

where *H* is a higher dimensional space.

This technique drastically reduces the amount of work required while dealing with nonlinear separating hyperplanes, concerning search for appropriate *ϕ*. Instead, one only works with *K*(**x**_*i*,_ **x**_*j*_), in place of **x**_*i*_ • **x**_*j*_. For classification purpose, where the sign of the function (**x**_*new*_ • **w** + *b*) is evaluated, the formulation employing kernels become:

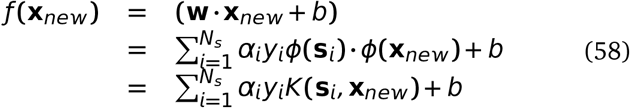

where **s**_*i*_ are the support vectors.

## WNT family 2*^nd^* order interaction ranking at different time points

SENP2WNT1 1 887 42 16 407 • SENP2WNT3 3 461 861 992 751 • DKK1WNT1 15 2415 1260 2300 1302 • FRZBWNT1 28 2478 63 528 144 • FRZBWNT3 29 2155 630 1213 153 • DKK1WNT3 31 2040 2041 906 1998 • SENP2WNT2B 32 848 45 572 529 • SENP2WNT3A 38 546 1146 583 964 • SENP2WNT5A 40 203 97 630 1124 • SENP2WNT4 45 830 492 2308 2211 • FZD7WNT1 55 1790 81 1241 817 • FZD5WNT1 58 2346 161 1928 131 • FRZBWNT4 76 1273 155 2047 1647 • PPP2CAWNT1 85 849 2018 439 513 • SENP2WNT2 86 1136 863 824 2485 • TCF7L1WNT1 87 1846 233 128 356 • RHOUWNT1 89 2193 151 2203 693 • SLC9A3R1WNT1 90 2281 101 931 397 • LRP5WNT1 92 2215 508 2133 863 • APCWNT1 104 2384 80 365 29 • FRZBWNT3A 114 1233 1049 1780 145 • NLKWNT1 115 485 1014 560 43 • FZD1WNT1 124 2181 157 1473 195 • CXXC4WNT1 125 2455 113 338 1558 • TCF7L1WNT3 126 5 2233 859 133 • LRP5WNT3 127 1716 1011 2369 708 • PPP2R1AWNT1 129 619 566 660 582 • APCWNT3 130 1907 1291 1399 86 • FRZBWNT2 131 520 796 579 1608 • FRZBWNT5A 135 2354 235 1726 518 • TLE2WNT1 136 2411 598 35 293 • FOSL1WNT1 143 2476 55 325 61 • FZD8WNT1 147 748 173 6 1085 • CSNK1DWNT1 150 2436 568 2316 1021 • FRZBWNT2B 158 1756 27 1589 157 • FZD5WNT3 164 913 1139 1633 192 • FZD7WNT3 168 493 1216 508 274 • GSK3BWNT1 172 1939 54 894 1897 • FZD1WNT3 181 1834 909 831 681 • SFRP4WNT1 185 2470 52 646 3 • FRAT1WNT1 200 2131 592 709 440 • DIXDC1WNT1 202 2172 12 516 2000 • TCF7WNT1 207 1677 1463 249 669 • FBXW11WNT1 223 327 787 1926 1139 • TLE2WNT3 226 902 2154 1496 381 • KREMEN1WNT1 228 2123 242 2436 824 • DAAM1WNT1 231 1245 965 102 2453 • FOSL1WNT3 233 796 526 227 218 • WNT2BWNT3 242 861 2191 139 249 • CTBP1WNT1 245 1792 34 32 159 • CSNK2A1WNT1 251 1172 793 2070 532 • FRAT1WNT3 252 2372 1439 1524 1058 • FZD6WNT1 267 803 1938 299 1394 • CCND3WNT1 273 8 405 2253 1037 • PPP2R1AWNT3 282 312 2409 1755 429 • CSNK1DWNT3 289 2103 2262 2354 898 • EP300WNT1 297 717 204 488 536 • FBXW11WNT3 299 678 969 1600 1237 • FBXW2WNT1 308 2031 785 476 696 • DKK1WNT4 312 1968 1719 185 208 • DKK1WNT3A 317 1408 2321 2430 2283 • TLE1WNT1 321 340 67 1559 173 • CSNK2A1WNT3 322 70 2143 2019 788 • DIXDC1WNT3 331 847 332 755 1294 • NKD1WNT1 338 7 260 1267 1212 • RHOUWNT3 343 1376 1052 341 1189 • WNT2WNT3 355 882 1069 408 231 • TCF7WNT3 368 620 1195 1838 1044 • DKK1WNT2 370 1201 2098 702 1287 • GSK3BWNT3 378 1722 837 1523 1019 • PITX2WNT1 380 1706 86 2064 1007 • DKK1WNT5A 382 2269 1464 2219 1479 • NLKWNT3 387 107 1795 380 158 • CTNNBIP1WNT1 400 2292 184 857 12 • DKK1WNT2B 406 1982 1099 1683 2383 • SLC9A3R1WNT3 410 1726 1400 2298 321 • EP300WNT3 419 3 2055 1083 348 • FBXW2WNT3 422 961 2381 1618 1994 • CXXC4WNT3 441 1930 1555 2452 1726 • PPP2CAWNT3 469 292 1807 1884 350 • FSHBWNT1 471 2394 761 928 234 • CTNNB1WNT1 488 2246 361 2364 759 • FZD6WNT3 494 16 2436 1371 776 • FZD8WNT3 500 127 1621 2029 2318 • CTBP1WNT3 503 2015 1703 2207 14 • FSHBWNT3 506 1255 2009 378 100 • CCND3WNT3 509 658 1636 716 1762 • CTBP2WNT1 510 783 128 628 621 • DAAM1WNT3 516 1500 1191 1583 1611 • FBXW4WNT1 530 2256 581 534 411 • SFRP4WNT3 533 2437 792 1779 51 • KREMEN1WNT3 545 2051 2108 158 219 • CCND2WNT1 573 2125 187 623 1535 • NKD1WNT3 591 53 2430 1271 288 • LRP6WNT1 597 1916 1184 1684 584 • FBXW4WNT3 607 900 1984 988 278 • FOSL1WNT2 626 549 933 2130 738 • CTNNB1WNT3 627 1276 1376 1310 710 • TWNT3 635 2026 2044 2135 40 • CTNNBIP1WNT3 639 1093 1486 1383 2 • FBXW11WNT4 653 1660 1080 587 1321 • DVL2WNT3 654 858 2395 1288 420 • TCF7L1WNT4 663 1031 2032 2406 836 • DVL1WNT1 672 393 239 191 756 • APCWNT2 680 243 1022 908 910 • TCF7L1WNT2 683 1190 1364 2060 755 • PITX2WNT3 685 1224 1466 1378 1473 • CTBP2WNT3 686 1061 1455 2025 95 • MYCWNT1 687 2365 1041 1603 96 • TCF7L1WNT5A 688 229 702 1670 209 • FZD1WNT2B 693 277 105 2424 1609 • APCWNT3A 701 1398 1674 2409 614 • FBXW11WNT2 703 1195 2438 200 1676 • CSNK1G1WNT3 719 1555 2460 1456 520 • FOSL1WNT5A 728 306 188 943 183 • DVL2WNT1 734 776 606 594 423 • FZD1WNT4 748 2153 297 2423 797 • APCWNT2B 752 1568 319 2415 1418 • APCWNT4 754 694 1992 833 2221 • AXIN1WNT1 761 1943 285 465 1099 • FOSL1WNT3A 784 691 1053 1439 1054 • LRP6WNT3 794 914 2466 1842 597 • TWNT1 797 2080 708 1595 5 • TCF7L1WNT3A 804 1237 1263 1513 1523 • FZD1WNT5A 805 1924 307 1120 165 • TLE1WNT3 810 11 1645 2057 740 • TCF7L1WNT2B 812 128 237 1311 2407 • FZD7WNT2 815 173 1242 84 1788 • FBXW11WNT3A 820 727 2251 1940 1879 • FZD1WNT2 838 222 1540 235 1256 • FOSL1WNT4 840 765 478 2323 1564 • RHOUWNT2 843 1535 1661 153 687 • APCWNT5A 847 2159 391 2305 106 • FRAT1WNT5A 849 1874 355 805 54 • CSNK2A1WNT3A 855 1073 1724 2483 1352 • TLE2WNT2B 868 2258 250 1056 2479 • CSNK2A1WNT2 872 1619 2226 1598 1005 • FGF4WNT1 874 1889 75 136 1793 • FZD1WNT3A 881 482 2298 2417 1337 • RHOUWNT5A 886 2070 531 2311 1579 • FZD7WNT5A 887 1403 41 1466 1629 • TLE2WNT3A 888 904 1097 1079 914 • CSNK2A1WNT4 890 833 2340 1904 1760 • RHOUWNT4 895 1168 2265 287 1482 • DIXDC1WNT4 908 1106 88 2394 2170 • FZD7WNT3A 910 1757 705 2105 390 • FOSL1WNT2B 917 558 30 749 1087 • FZD7WNT4 926 1925 694 949 1046 • CSNK1DWNT2 928 265 1934 204 389 • BTRCWNT1 930 412 1923 67 2004 • CSNK1DWNT4 931 1219 940 1605 2099 • GSK3BWNT4 936 1475 995 686 1480 • FBXW11WNT5A 939 227 1476 41 1067 • WIF1WNT3 944 370 959 34 1056 • CSNK1G1WNT1 946 2370 448 2034 829 • BTRCWNT3 951 1282 2153 1924 1934 • CSNK2A1WNT2B 954 245 954 1852 79 • WNT3AWNT4 968 99 1955 1344 2222 • AXIN1WNT3 969 1665 901 1257 1144 • PYGO1WNT1 975 1549 441 1906 1617 • PPP2R1AWNT5A 980 2288 679 76 957 • FZD5WNT3A 981 822 1946 2243 468 • DVL1WNT3 982 356 1398 1133 562 • WIF1WNT1 985 1244 36 149 6 • LRP5WNT3A 986 492 1258 2231 695 • FRAT1WNT4 991 122 623 1882 1507 • WNT2WNT4 995 303 1232 1818 1089 • DIXDC1WNT5A 996 1710 53 2116 2389 • JUNWNT1 1000 985 328 270 35 • LEF1WNT3 1003 1605 1381 2349 560 • PORCNWNT1 1004 1747 510 924 2388 • WNT2BWNT3A 1005 208 2331 715 978 • RHOUWNT3A 1010 288 2335 2221 1452 • FBXW11WNT2B 1015 460 1352 958 2373 • FRAT1WNT2 1019 1653 2179 1746 1786 • CSNK2A1WNT5A 1024 2374 1580 830 470 • DIXDC1WNT3A 1030 1286 170 1420 2065 • DIXDC1WNT2B 1036 1494 9 1095 1365 • FZD7WNT2B 1038 328 29 2174 1849 • LRP5WNT5A 1040 2397 972 1967 289 • FRAT1WNT2B 1046 2107 443 563 1361 • FRAT1WNT3A 1054 1951 1739 1029 1486 • LRP5WNT2 1056 1504 1803 223 959 • FZD5WNT5A 1059 1553 543 1566 193 • CSNK1DWNT5A 1062 2396 611 1794 986 • GSK3BWNT2 1070 879 1720 401 2431 • DIXDC1WNT2 1079 43 199 938 1421 • RHOUWNT2B 1082 453 293 150 1678 • TLE2WNT2 1087 655 2168 1263 1088 • PORCNWNT3 1088 1604 2052 1315 2298 • TCF7WNT5A 1112 151 2019 1176 1463 • TLE2WNT5A 1115 2032 399 1225 214 • NLKWNT2 1117 134 2272 921 2026 • GSK3BWNT3A 1118 1234 2188 1036 1495 • TCF7WNT2 1131 1699 1307 434 1972 • AESWNT1 1134 631 1063 1182 299 • PPP2R1AWNT2B 1135 1299 999 171 1731 • TLE2WNT4 1147 1020 1085 1528 1368 • MYCWNT3 1149 660 1692 1973 26 • FZD5WNT2 1157 1319 2227 1535 2036 • CSNK1DWNT3A 1166 1179 2161 2402 2009 • WNT2BWNT5A 1182 1580 2180 1430 1253 • EP300WNT4 1185 1341 1068 2427 1537 • CXXC4WNT2 1188 172 740 2353 2314 • NLKWNT3A 1189 1579 2341 926 1298 • FZD5WNT4 1196 514 621 2391 2362 • FSHBWNT2 1202 57 2398 319 1727 • CXXC4WNT4 1207 2245 880 2412 1427 • FZD2WNT1 1208 1049 26 1665 398 • FZD5WNT2B 1212 1377 94 1359 1753 • KREMEN1WNT2B 1216 468 82 1553 1925 • PPP2CAWNT2 1218 483 2176 1246 2111 • LRP5WNT4 1222 117 980 1057 2210 • PPP2R1AWNT2 1229 257 2305 615 478 • TCF7WNT3A 1231 1488 2137 1901 1157 • SFRP1WNT1 1235 1861 180 1151 1734 • EP300WNT3A 1236 669 1550 948 2080 • TCF7WNT4 1237 2128 1970 69 2139 • CSNK1DWNT2B 1241 1254 628 1737 1467 • PPP2CAWNT5A 1246 1216 801 671 728 • CSNK1A1WNT3 1248 133 1215 492 66 • FBXW2WNT2 1252 742 1450 456 1982 • WNT2WNT3A 1257 268 912 542 1646 • CTBP1WNT2 1259 574 1388 1619 1324 • FZD6WNT2 1262 649 1471 648 275 • LRP5WNT2B 1264 718 715 2456 1063 • FGF4WNT3 1270 1484 1404 2463 415 • WNT3AWNT5A 1271 1264 343 1443 2322 • WNT2WNT5A 1272 704 485 209 480 • DAAM1WNT2 1273 1740 2231 2100 1122 • CXXC4WNT5A 1275 1858 295 348 1886 • WNT2BWNT4 1281 2223 2276 168 2063 • KREMEN1WNT2 1284 78 871 1146 1347 • FBXW4WNT2 1285 1404 1507 1107 1290 • PITX2WNT3A 1286 517 615 1340 1630 • SLC9A3R1WNT2 1290 121 1717 1136 268 • GSK3BWNT2B 1293 1228 195 1142 1496 • PPP2R1AWNT4 1294 1166 1499 2045 1006 • FZD8WNT4 1296 810 1566 1162 942 • CTBP1WNT5A 1297 2043 716 1282 71 • KREMEN1WNT4 1299 556 1637 636 841 • BCL9WNT3 1300 975 2477 1374 1197 • CXXC4WNT3A 1304 2295 843 2361 1637 • NLKWNT4 1310 1356 2223 2367 152 • FZD8WNT5A 1316 73 334 1596 676 • FBXW4WNT2B 1317 223 650 557 783 • FZD8WNT2B 1319 756 109 427 2166 • FZD6WNT5A 1320 1231 1375 2168 445 • CCND1WNT1 1325 101 302 1480 2234 • WNT2WNT2B 1331 674 296 510 2029 • FBXW2WNT4 1332 1315 2385 870 1326 • PPP2R1AWNT3A 1335 818 1303 2113 1901 • FBXW4WNT4 1342 1042 1537 2013 1382 • FSHBWNT5A 1344 1383 1747 1738 148 • CCND3WNT2B 1347 1229 1193 88 1172 • SLC9A3R1WNT5A 1351 1732 652 1388 176 • EP300WNT2B 1352 10 688 475 2340 • WNT4WNT5A 1360 1936 822 1122 18 • EP300WNT2 1362 2143 1950 1715 1723 • FZD6WNT3A 1364 102 2388 2192 891 • CXXC4WNT2B 1365 1998 150 1706 1503 • PPP2CAWNT3A 1367 576 1326 2477 1116 • PPP2CAWNT4 1371 1799 2106 1702 2195 • SLC9A3R1WNT2B 1383 743 108 1537 1339 • SFRP4WNT2 1387 1947 1288 814 363 • PITX2WNT2 1390 1561 1819 483 742 • DAAM1WNT3A 1393 503 1430 1929 1471 • PPP2CAWNT2B 1399 1928 601 1165 1405 • FZD8WNT2 1403 2424 952 968 772 • KREMEN1WNT3A 1404 9 1367 2407 1478 • SLC9A3R1WNT4 1405 600 1878 491 605 • CSNK1A1WNT1 1407 463 603 564 146 • CCND3WNT5A 1411 280 1588 1899 503 • FBXW2WNT3A 1412 978 948 2419 2460 • FBXW2WNT2B 1421 333 376 1464 1225 • FZD8WNT3A 1422 706 1145 1628 1499 • CCND2WNT3 1432 19 1181 2472 2100 • FOXN1WNT1 1436 1099 1465 1327 358 • AESWNT3 1443 170 2054 1694 1166 • FZD6WNT4 1445 946 1846 1188 1696 • GSK3BWNT5A 1449 1554 386 1077 1882 • PITX2WNT2B 1451 21 160 364 1741 • LEF1WNT1 1453 2312 553 1004 1293 • DAAM1WNT2B 1465 375 555 1597 2074 • SFRP4WNT5A 1467 2474 112 865 575 • BCL9WNT1 1469 2265 573 267 599 • DAAM1WNT4 1472 751 1839 1201 1391 • TCF7WNT2B 1475 274 1699 1748 665 • SFRP4WNT4 1487 1371 1088 1156 1504 • FSHBWNT3A 1490 689 1505 2355 1698 • NLKWNT2B 1491 954 175 1376 1707 • PITX2WNT4 1494 407 784 89 1020 • TLE1WNT2 1499 1502 1074 1215 2110 • NKD1WNT2 1502 646 1639 297 2133 • CCND3WNT2 1506 1859 2166 549 1904 • FOXN1WNT3 1507 819 1546 2280 758 • NLKWNT5A 1511 1685 1318 1652 1976 • FSHBWNT4 1517 1564 1969 888 1917 • CTBP1WNT4 1519 782 1140 1663 1770 • PYGO1WNT3 1526 262 1638 318 285 • FBXW4WNT3A 1527 607 2312 2278 586 • TLE1WNT5A 1529 1631 281 940 279 • CTBP1WNT2B 1535 929 22 1459 1319 • FBXW4WNT5A 1539 97 984 1118 481 • SLC9A3R1WNT3A 1544 2182 955 50 870 • CCND3WNT3A 1546 259 2413 2291 1320 • KREMEN1WNT5A 1555 1961 726 2426 963 • CTNNBIP1WNT4 1560 1314 550 1346 405 • EP300WNT5A 1562 1523 577 1584 657 • TWNT2 1566 109 2442 70 2333 • FZD6WNT2B 1568 205 1183 1538 1922 • CTNNB1WNT5A 1582 1741 322 2324 230 • FBXW2WNT5A 1588 1262 1210 1312 615 • SFRP4WNT3A 1602 2242 937 936 1813 • GSK3AWNT3 1604 577 1759 1992 1888 • CCND3WNT4 1607 1498 1262 141 1228 • CTNNB1WNT3A 1612 710 2260 2309 2032 • SFRP1WNT3 1613 4 974 2122 2481 • TLE1WNT4 1623 850 925 421 1525 • FSHBWNT2B 1640 2393 1443 879 2041 • WNT1WNT3 1649 820 1341 1767 581 • DAAM1WNT5A 1653 430 135 1063 1687 • TLE1WNT2B 1656 282 119 739 2178 • TLE1WNT3A 1670 1340 234 1389 2469 • NKD1WNT2B 1672 62 1363 2348 1833 • CTNNBIP1WNT3A 1678 2054 1975 1349 298 • FZD2WNT3 1679 1758 958 675 270 • CCND1WNT3 1682 347 1746 2140 1721 • SFRP4WNT2B 1704 2366 85 108 2173 • PITX2WNT5A 1717 1347 440 293 75 • AESWNT4 1722 1434 1595 1017 51 • JUNWNT3 1738 32 1721 2458 54 • LRP6WNT5A 1750 474 274 1446 943 • DVL1WNT2 1752 1472 2235 1521 862 • CTBP2WNT2 1757 641 1838 2418 1151 • DVL2WNT2 1759 1831 2419 590 2464 • CTBP2WNT3A 1763 925 2380 2194 602 • CTBP1WNT3A 1764 1567 2205 414 1009 • AXIN1WNT3A 1773 1277 2237 721 1448 • WIF1WNT2B 1776 433 31 619 1024 • NKD1WNT3A 1778 2297 1972 402 886 • CTNNBIP1WNT2 1785 209 820 317 334 • NKD1WNT4 1790 1674 1886 1065 2104 • CTNNB1WNT4 1795 2184 1328 2003 1014 • CSNK1G1WNT2 1801 298 2024 440 1754 • DVL1WNT5A 1811 917 522 775 317 • DVL2WNT4 1816 1395 737 1351 2351 • NKD1WNT5A 1817 91 917 2134 1836 • CSNK1G1WNT3A 1818 1737 2035 1000 2483 • DVL2WNT5A 1820 1069 839 1342 360 • BTRCWNT4 1821 984 2367 195 977 • CCND2WNT2 1823 1014 1591 1234 2197 • TWNT4 1829 1967 1533 1012 1377 • WIF1WNT2 1838 1688 885 2482 1930 • CTNNB1WNT2 1847 712 1698 902 2345 • CTBP2WNT4 1858 1373 939 1832 1978 • CSNK1G1WNT2B 1863 2086 977 295 2031 • AXIN1WNT2 1865 696 1359 399 734 • GSK3AWNT1 1867 2239 1343 1167 1455 • AESWNT2 1872 935 1402 127 1599 • LRP6WNT2 1875 305 1700 1357 1823 • CSNK1G1WNT4 1888 988 1445 2041 2287 • CTNNBIP1WNT2B 1897 1428 171 15 1113 • LEF1WNT2 1904 242 1428 515 661 • DVL2WNT3A 1912 1636 860 1996 749 • LRP6WNT4 1926 829 1310 576 1952 • CSNK1G1WNT5A 1929 2413 2042 2091 668 • CCND2WNT5A 1932 1348 1126 1650 2156 • AXIN1WNT4 1934 800 1403 2431 2096 • LRP6WNT2B 1937 1540 450 2227 885 • TWNT3A 1938 581 2369 1829 1470 • FGF4WNT2 1947 83 1647 1474 1069 • DVL2WNT2B 1959 737 765 1667 382 • BTRCWNT2 1982 901 2315 394 2112 • PORCNWNT5A 1984 2267 869 437 2316 • CSNK1A1WNT2 1986 2382 750 612 297 • DVL1WNT3A 1999 1507 685 268 1552 • WNT1WNT2B 2004 180 2097 1016 1328 • AXIN1WNT5A 2006 1266 1106 330 1195 • LRP6WNT3A 2007 1576 810 1177 2429 • PORCNWNT3A 2008 240 2323 914 1650 • AESWNT3A 2010 1259 1940 2345 573 • CCND1WNT2 2014 894 1655 1444 777 • TWNT5A 2015 1222 1712 854 444 • CTBP2WNT2B 2016 1908 185 1931 537 • CTNNB1WNT2B 2023 773 591 1587 2299 • TWNT2B 2025 790 735 1578 2192 • BTRCWNT3A 2034 1180 1755 1997 2401 • FOXN1WNT4 2039 2192 1366 154 1140 • DVL1WNT2B 2045 314 127 989 1104 • LEF1WNT4 2049 2033 1587 351 1194 • SFRP1WNT2 2055 300 855 555 1871 • AESWNT2B 2058 86 549 1783 1330 • AESWNT5A 2062 1767 1221 243 1517 • CTBP2WNT5A 2065 175 639 1291 1722 • CTNNBIP1WNT5A 2068 1367 444 1392 169 • LEF1WNT2B 2074 982 47 917 1556 • FOXN1WNT5A 2079 1108 2378 230 2011 • FZD2WNT2 2083 1807 758 357 1192 • PYGO1WNT2 2085 771 1593 152 2205 • PORCNWNT2 2088 524 1813 419 1951 • CCND2WNT3A 2104 1426 2360 2389 2136 • FGF4WNT5A 2108 738 143 1139 2033 • SFRP1WNT5A 2113 2217 482 1725 2484 • WNT3WNT4 2136 880 1153 774 636 • WIF1WNT3A 2154 1574 1083 263 1350 • MYCWNT5A 2156 1327 1547 1639 606 • WIF1WNT4 2157 1267 507 2276 2272 • BTRCWNT2B 2158 940 1401 1076 1565 • SFRP1WNT3A 2160 837 338 1425 705 • JUNWNT2 2161 507 1458 1010 435 • BTRCWNT5A 2162 922 2476 1862 2327 • CSNK1A1WNT2B 2166 1664 605 2001 91 • AXIN1WNT2B 2167 547 115 1108 828 • GSK3AWNT4 2168 1760 1694 2376 741 • MYCWNT2 2170 1508 1861 530 1595 • LEF1WNT5A 2171 1122 1029 1552 1490 • WNT1WNT2 2172 211 1462 424 2387 • FGF4WNT2B 2173 342 14 526 2404 • FZD2WNT2B 2177 1528 120 1316 763 • MYCWNT2B 2179 246 1240 1232 541 • WNT3WNT3A 2185 747 1980 2053 1068 • FGF4WNT3A 2186 442 818 1769 798 • WNT1WNT5A 2189 1529 1716 277 739 • WNT3WNT5A 2207 1343 404 2380 2091 • BCL9WNT2 2211 873 1683 379 1635 • LEF1WNT3A 2216 326 2238 1714 1103 • SFRP1WNT4 2219 1238 669 2173 1848 • BCL9WNT4 2220 2084 458 1953 1187 • SFRP1WNT2B 2229 36 129 1356 1589 • MYCWNT4 2232 1307 1459 2139 825 • FOXN1WNT3A 2234 831 2124 1984 1242 • FGF4WNT4 2242 2146 406 2399 1145 • FOXN1WNT2B 2247 138 900 2138 2180 • DVL1WNT4 2252 969 763 2413 1332 • CSNK1A1WNT5A 2258 1642 1162 2450 220 • CCND2WNT4 2259 1083 811 291 116 • CCND2WNT2B 2261 671 337 519 1961 • PORCNWNT4 2267 730 1399 57 1576 • MYCWNT3A 2271 942 858 1820 196 • FZD2WNT4 2280 1372 600 1988 1835 • PORCNWNT2B 2283 22 467 120 1931 • CSNK1A1WNT4 2284 1378 1999 1440 1789 • WIF1WNT5A 2285 2175 271 1233 1649 • JUNWNT3A 2288 467 2002 1880 800 • FOXN1WNT2 2296 1026 2362 178 1335 • FZD2WNT3A 2301 1552 1327 1302 1709 • FZD2WNT5A 2304 1101 632 1964 190 • PYGO1WNT3A 2307 1074 1660 1025 883 • GSK3AWNT3A 2309 960 1709 1688 1633 • BCL9WNT3A 2313 1496 1687 2061 2379 • PYGO1WNT4 2315 418 1523 5 1307 • PYGO1WNT2B 2318 828 559 697 1076 • JUNWNT5A 2319 1401 659 1922 98 • CCND1WNT4 2320 1816 1238 1510 1232 • CCND1WNT3A 2326 1545 2375 2343 1937 • GSK3AWNT2 2331 1649 2282 2149 2130 • BCL9WNT2B 2336 996 305 2038 2467 • BCL9WNT5A 2340 1646 1252 2072 918 • WNT1WNT3A 2342 2273 1670 1604 2458 • JUNWNT4 2348 2102 1434 2382 1141 • CCND1WNT5A 2352 534 662 1659 2470 • CSNK1A1WNT3A 2364 1562 1765 1428 126 • CCND1WNT2B 2374 1440 228 112 612 • GSK3AWNT5A 2376 497 1129 1893 1295 • JUNWNT2B 2380 158 211 829 765 • WNT1WNT4 2401 1754 2443 550 1577 • GSK3AWNT2B 2419 30 579 157 1380 • PYGO1WNT5A 2424 1394 663 1089 1428

## WNT family 2^*nd*^ order interaction ranking at different durations

PYGO1WNT5A 6 479 506 1847 • FZD5WNT2 26 897 1803 1450 • FZD5WNT4 49 463 945 2348 • PYGO1WNT4 61 772 1971 2417 • WNT3WNT5A 70 37 1676 2306 • CSNK1A1WNT1 104 2388 2169 1071 • CSNK1A1WNT5A 121 1315 1937 250 • CSNK1A1WNT4 132 647 1320 1520 • FZD5WNT5A 136 579 1528 2296 • PYGO1WNT2 149 1948 1153 1710 • RHOUWNT5A 152 573 590 607 • DIXDC1WNT5A 164 30 442 1764 • JUNWNT5A 194 1515 1716 38 • RHOUWNT4 197 291 556 561 • WNT2BWNT5A 210 252 2084 1578 • RHOUWNT2 225 523 975 192 • WNT3AWNT5A 226 626 348 2302 • PORCNWNT2 231 1928 1576 2289 • PORCNWNT4 246 937 1873 1945 • CTNNB1WNT5A 263 1836 1168 1325 • TWNT5A 292 957 2456 501 • AESWNT5A 312 1576 2187 837 • CTNNBIP1WNT5A 314 1953 1124 1473 • PORCNWNT5A 318 1854 601 2286 • KREMEN1WNT5A 333 974 861 2480 • CTBP1WNT5A 336 2412 1868 2056 • CTNNB1WNT4 352 2035 2388 2457 • BCL9WNT5A 354 2155 2483 2036 • WNT3WNT4 359 35 1564 735 • JUNWNT4 360 1840 1467 499 • AXIN1WNT5A 364 365 667 2107 • CSNK1A1WNT3A 377 1466 618 1354 • WNT2BWNT4 419 508 724 953 • CCND2WNT5A 434 1921 1981 2260 • CTNNB1WNT2 456 1376 1741 1976 • CSNK1DWNT5A 459 321 1554 2342 • FOXN1WNT5A 463 1016 1983 951 • PYGO1WNT3A 474 1475 2256 1142 • APCWNT5A 477 2063 1686 1488 • PYGO1WNT2B 492 1709 1209 151 • FRAT1WNT5A 505 552 862 1990 • CCND2WNT4 511 1056 1461 2029 • MYCWNT5A 512 1161 2412 1803 • CCND1WNT5A 513 929 412 1509 • PORCNWNT1 514 1393 2078 1575 • DIXDC1WNT2 522 244 1647 350 • CSNK1A1WNT2 530 1102 1682 933 • BTRCWNT5A 543 1541 1671 1336 • CTBP1WNT4 559 1416 1094 2482 • WIF1WNT5A 562 241 1836 2013 • FZD5WNT1 564 617 2284 1432 • DIXDC1WNT4 572 24 695 1067 • CTBP2WNT5A 582 1794 2172 2472 • WNT1WNT5A 586 824 2028 1181 • FRAT1WNT4 612 480 2242 1290 • SLC9A3R1WNT5A 618 682 503 2219 • DVL1WNT5A 621 1645 414 1031 • FSHBWNT5A 626 349 704 1725 • WNT2WNT5A 632 490 787 2323 • GSK3AWNT5A 639 286 389 1426 • PYGO1WNT1 661 357 2102 2333 • WNT1WNT4 664 1793 1311 1171 • FOSL1WNT5A 691 231 1864 1751 • WNT3AWNT4 698 345 944 1678 • RHOUWNT2B 702 956 366 70 • DIXDC1WNT1 715 9 630 762 • TWNT1 722 581 2112 562 • MYCWNT4 724 747 1019 1836 • EP300WNT5A 740 2090 964 2422 • AESWNT4 741 1309 1577 127 • NKD1WNT5A 743 1956 1516 2413 • CTNNB1WNT1 753 2052 2379 1931 • FSHBWNT4 764 334 1644 932 • BCL9WNT4 774 986 1966 318 • AXIN1WNT4 780 637 1845 753 • CCND1WNT4 790 1019 1280 1348 • RHOUWNT1 793 292 383 1153 • FOXN1WNT4 796 443 2171 2136 • FZD5WNT2B 803 1896 808 1089 • JUNWNT2 806 1513 1721 2168 • TWNT4 809 641 2074 1472 • CSNK1DWNT4 826 652 2258 1393 • CTBP1WNT3A 836 1562 453 1951 • FBXW4WNT5A 838 1157 1034 672 • LRP5WNT5A 844 208 1780 235 • DVL1WNT4 851 700 1149 335 • BCL9WNT2 860 912 1668 2129 • KREMEN1WNT4 865 925 771 2238 • CTBP1WNT2 874 2304 1571 2070 • DVL2WNT5A 897 2254 614 2395 • FZD5WNT3A 902 2287 1744 1093 • FOXN1WNT2 903 896 1559 1435 • SLC9A3R1WNT4 908 396 864 1374 • FRAT1WNT2 912 844 2252 1162 • WNT4WNT5A 918 1751 636 2255 • CTNNBIP1WNT2 923 2463 1938 1198 • JUNWNT1 933 2473 471 1197 • WNT3WNT3A 942 702 251 1479 • CTNNBIP1WNT4 944 1567 1139 1286 • AXIN1WNT1 951 675 1625 1503 • PYGO1WNT3 962 1103 890 899 • CSNK1A1WNT2B 965 1719 1267 1 • CTNNB1WNT3A 972 1044 1853 2353 • PORCNWNT2B 974 2480 384 333 • WIF1WNT4 975 418 1233 1469 • TLE1WNT5A 978 366 1065 1624 • RHOUWNT3A 986 1501 2264 632 • FRZBWNT5A 988 1687 1593 944 • APCWNT4 992 2228 485 2391 • NKD1WNT4 997 2444 1011 1173 • RHOUWNT3 998 2240 91 177 • TLE2WNT5A 999 836 1152 165 • BTRCWNT4 1013 1399 1658 1837 • FRAT1WNT1 1020 560 2253 1320 • AESWNT1 1032 1313 1599 1475 • AESWNT2 1046 2317 1197 144 • FOXN1WNT1 1057 781 2320 1665 • FOSL1WNT4 1060 307 751 1447 • MYCWNT2 1069 1185 1231 707 • AXIN1WNT2 1080 513 330 1454 • TWNT2 1089 1037 1225 120 • CTNNB1WNT2B 1093 1297 937 40 • CCND3WNT5A 1105 2445 1048 1923 • PPP2CAWNT5A 1111 1029 2392 2262 • CTBP1WNT1 1115 1666 2324 1591 • TCF7WNT5A 1117 1585 2010 1400 • CCND2WNT2 1123 1374 1105 1955 • DKK1WNT5A 1129 264 2270 1893 • KREMEN1WNT2 1131 2236 2139 2001 • CXXC4WNT5A 1144 2318 1259 1771 • CCND1WNT1 1145 780 933 842 • WNT1WNT2 1148 1017 1739 2431 • GSK3AWNT4 1156 175 1638 738 • PORCNWNT3A 1170 949 2304 2150 • CTBP2WNT4 1186 767 1115 1598 • BCL9WNT1 1195 1992 2441 441 • BTRCWNT2 1205 2284 1301 2128 • CCND1WNT2 1206 1426 661 1671 • FSHBWNT1 1212 267 2260 438 • FZD6WNT5A 1213 1165 162 1497 • CSNK1DWNT2 1214 743 1921 269 • GSK3AWNT1 1216 265 748 846 • KREMEN1WNT1 1220 1014 436 2169 • FBXW2WNT5A 1225 640 1973 2204 • CSNK1G1WNT5A 1226 43 2095 2273 • CTNNBIP1WNT2B 1235 2153 643 2149 • JUNWNT2B 1239 2419 1453 173 • DAAM1WNT5A 1240 1005 1023 1698 • CCND2WNT1 1247 1285 941 1590 • SLC9A3R1WNT2 1262 734 538 1539 • GSK3AWNT2 1275 599 2042 1294 • MYCWNT1 1289 858 1769 1193 • FSHBWNT2 1294 683 543 890 • WNT2WNT4 1296 1087 2457 988 • SFRP1WNT5A 1314 166 1969 2159 • FZD8WNT5A 1316 41 294 2474 • FRZBWNT4 1319 782 1017 2153 • NLKWNT5A 1322 752 1421 2135 • FBXW11WNT5A 1329 716 1071 1970 • PITX2WNT5A 1330 68 638 1701 • CCND2WNT3A 1338 639 736 2105 • GSK3BWNT5A 1347 137 577 443 • CSNK1A1WNT3 1353 1657 612 299 • PPP2R1AWNT5A 1366 431 652 734 • LRP5WNT4 1372 93 1930 1936 • WNT2BWNT3A 1380 891 1055 528 • JUNWNT3A 1383 1151 1532 436 • LRP5WNT2 1390 281 2453 531 • CTBP2WNT2 1392 1215 1877 2407 • FRAT1WNT3A 1396 904 1581 1276 • CSNK2A1WNT5A 1398 1041 399 1917 • TLE1WNT4 1406 285 919 1214 • EP300WNT4 1410 2335 1713 1722 • CSNK1DWNT2B 1416 1031 2243 166 • FSHBWNT3A 1424 938 1732 535 • CTNNBIP1WNT3A 1426 1654 2153 1204 • BCL9WNT3A 1433 890 1435 1866 • CCND3WNT4 1443 1320 1926 1251 • DVL2WNT2 1444 1024 1211 1425 • CTNNB1WNT3 1445 1280 572 941 • DVL1WNT2 1450 2429 1323 288 • BTRCWNT1 1452 592 2337 1797 • DVL2WNT4 1461 1972 2269 1877 • FZD5WNT3 1463 1358 1232 1333 • NKD1WNT2 1466 1883 2047 813 • PORCNWNT3 1467 1946 518 2227 • FOXN1WNT3A 1468 1877 2114 1002 • DIXDC1WNT2B 1477 430 227 1581 • CSNK1DWNT1 1484 230 2083 2108 • TWNT3A 1492 1298 1850 330 • FZD8WNT2 1499 186 1282 1890 • FSHBWNT2B 1501 807 698 1487 • CTBP1WNT2B 1513 2083 2422 873 • AESWNT2B 1514 2339 730 30 • TWNT2B 1526 753 855 23 • DIXDC1WNT3 1537 529 206 1341 • APCWNT2 1549 1597 1758 1012 • CTNNBIP1WNT1 1550 2172 702 798 • FRZBWNT3A 1551 1847 1490 1659 • FZD7WNT5A 1553 149 79 2181 • TCF7WNT4 1555 1556 768 2469 • APCWNT3A 1561 1125 798 2294 • TLE2WNT4 1567 601 1484 205 • DIXDC1WNT3A 1572 350 1688 371 • WIF1WNT1 1573 424 1806 349 • AESWNT3A 1576 1763 2024 987 • FOSL1WNT3A 1578 543 1535 2158 • TCF7L1WNT5A 1580 102 876 2243 • DVL1WNT1 1584 2331 432 608 • MYCWNT3A 1588 994 658 2022 • APCWNT1 1590 654 2423 2420 • CSNK1DWNT3A 1599 822 1032 2186 • FZD1WNT5A 1602 146 839 2217 • WIF1WNT2 1619 462 1463 1709 • AXIN1WNT2B 1622 2431 915 2251 • FBXW4WNT4 1623 1131 2212 1683 • CTBP2WNT1 1636 894 942 906 • LEF1WNT5A 1641 191 1058 2236 • CCND1WNT3A 1642 2089 2378 1191 • CCND3WNT1 1643 2475 1077 1825 • FOSL1WNT1 1649 219 2484 1766 • FRZBWNT2 1650 1048 2016 1306 • AESWNT3 1659 1807 644 179 • CCND2WNT2B 1668 2309 1214 900 • FOSL1WNT2 1669 352 1773 1856 • SLC9A3R1WNT1 1670 515 377 419 • LRP6WNT5A 1675 398 1103 2319 • EP300WNT2 1679 1721 1859 1456 • SLC9A3R1WNT2B 1682 1233 498 1145 • BCL9WNT2B 1684 1400 693 597 • TLE2WNT2 1690 1505 2444 1493 • FGF4WNT5A 1700 333 1481 1660 • CXXC4WNT2 1704 1662 1466 2095 • FRZBWNT1 1711 723 2443 1099 • CXXC4WNT4 1721 1920 1163 1215 • FZD7WNT4 1724 287 537 677 • KREMEN1WNT3A 1726 1378 1777 2072 • DAAM1WNT4 1728 2004 1543 1390 • CCND2WNT3 1729 1779 1166 2468 • DKK1WNT4 1735 238 1492 2470 • AXIN1WNT3 1743 2241 533 1753 • FRAT1WNT3 1746 2136 815 960 • CCND3WNT2B 1749 2421 795 925 • CSNK2A1WNT4 1750 1154 1349 2099 • GSK3BWNT4 1754 169 1073 41 • FBXW4WNT1 1756 878 2089 1557 • LRP5WNT1 1758 84 846 1925 • FSHBWNT3 1763 2171 1426 1275 • BTRCWNT2B 1765 2022 1506 229 • FBXW4WNT2 1773 2134 1457 1612 • KREMEN1WNT3 1774 2380 159 359 • CCND3WNT2 1775 1758 1562 2196 • PPP2CAWNT4 1780 1490 2076 1897 • JUNWNT3 1784 898 2061 278 • FZD6WNT4 1786 1018 766 602 • NKD1WNT1 1789 1745 1218 1188 • WNT2WNT2B 1790 1543 558 85 • GSK3BWNT1 1791 89 177 1029 • PITX2WNT4 1792 55 1862 1854 • NLKWNT4 1796 1464 858 1724 • TLE1WNT2 1799 530 1521 2037 • WNT1WNT3A 1801 2191 1287 1627 • FZD8WNT4 1803 124 1666 1968 • CTBP1WNT3 1807 1978 1503 2206 • CSNK1G1WNT4 1811 94 1494 1530 • FZD1WNT4 1825 58 1445 1186 • CSNK2A1WNT2 1827 1033 604 2455 • FRAT1WNT2B 1833 2138 743 247 • WNT2BWNT3 1835 1711 1525 2019 • EP300WNT1 1837 1126 2394 1560 • NKD1WNT2B 1842 2410 344 513 • TWNT3 1851 1363 435 2 • AXIN1WNT3A 1853 2007 1288 703 • CXXC4WNT3A 1862 892 1144 1747 • MYCWNT2B 1863 1145 1092 348 • FZD2WNT4 1871 434 2352 2388 • DVL2WNT1 1873 2442 2007 1914 • FZD2WNT2 1875 435 1451 1562 • TLE1WNT3A 1877 720 1230 2350 • FBXW2WNT4 1878 1000 1474 1107 • CCND1WNT2B 1884 2243 71 1303 • FOXN1WNT2B 1889 1798 717 111 • PPP2CAWNT2 1894 1303 1953 1704 • TCF7WNT2 1899 1519 539 2272 • FBXW11WNT4 1901 588 1687 2330 • FGF4WNT4 1907 138 2057 802 • FZD2WNT5A 1908 420 397 1428 • GSK3AWNT3A 1911 1577 2383 1721 • BTRCWNT3A 1916 2286 1590 2017 • DKK1WNT1 1918 201 2303 1299 • SFRP1WNT2 1919 283 917 1518 • SFRP1WNT4 1922 151 1496 986 • WNT2WNT3A 1925 585 2348 2098 • SLC9A3R1WNT3 1926 2418 253 1793 • CTBP2WNT2B 1932 1644 980 1138 • PPP2R1AWNT4 1942 1592 1888 1148 • DVL1WNT2B 1945 1922 360 2047 • DKK1WNT2 1947 395 510 2366 • KREMEN1WNT2B 1948 1583 9 181 • FOSL1WNT2B 1949 768 962 185 • CXXC4WNT2B 1953 2398 863 1553 • BCL9WNT3 1954 2115 883 713 • WIF1WNT3A 1955 927 2262 426 • PPP2R1AWNT3A 1961 818 1181 1969 • CSNK2A1WNT3A 1966 1268 2435 496 • GSK3AWNT2B 1972 2158 88 1351 • FOXN1WNT3 1973 1083 108 480 • PITX2WNT2 1976 189 1208 1702 • TLE2WNT3A 1977 2064 1916 525 • APCWNT2B 1982 1256 1309 599 • TCF7L1WNT4 1986 275 2043 2178 • CTBP2WNT3A 1989 1412 1452 2078 • CTNNBIP1WNT3 1991 2433 64 903 • TCF7WNT2B 1996 2298 1797 514 • LEF1WNT4 1998 162 2293 945 • SENP2WNT5A 2000 483 1550 1943 • CCND1WNT3 2001 1043 169 1814 • MYCWNT3 2004 1356 328 201 • DKK1WNT2B 2008 649 2135 264 • TCF7WNT1 2010 745 1110 2109 • PITX2WNT3 2015 524 58 589 • FBXW2WNT3A 2018 1969 841 2402 • TLE2WNT1 2020 1558 1478 311 • FBXW2WNT1 2026 1723 1128 966 • FBXW11WNT1 2028 570 1439 446 • CXXC4WNT1 2030 1697 1755 1614 • WNT1WNT2B 2034 1965 935 347 • DVL1WNT3A 2037 1242 2272 401 • FZD7WNT2 2041 299 326 811 • FGF4WNT1 2047 114 1212 569 • DVL2WNT3A 2050 1927 1344 2068 • SFRP4WNT5A 2054 123 504 588 • FZD6WNT1 2059 425 183 667 • WNT2WNT3 2060 2271 63 924 • DAAM1WNT1 2061 1112 832 1039 • CSNK2A1WNT1 2065 918 240 1260 • NLKWNT2 2069 381 893 1677 • NLKWNT1 2073 800 2033 1511 • PPP2CAWNT3A 2075 1835 1372 1989 • EP300WNT2B 2078 1506 782 1185 • EP300WNT3A 2086 1431 2317 1869 • SFRP1WNT3 2088 1627 631 1256 • FOSL1WNT3 2090 810 525 1056 • PPP2R1AWNT2 2092 499 616 2337 • FZD8WNT1 2094 42 260 493 • LRP6WNT3 2098 1336 111 1219 • FZD2WNT2B 2101 1563 210 281 • NKD1WNT3A 2103 1612 1039 1315 • CSNK1G1WNT2 2109 403 2282 1356 • SFRP1WNT3A 2110 436 1867 2123 • FZD8WNT3A 2111 426 1622 1988 • NLKWNT2B 2120 2098 587 399 • FZD7WNT3A 2121 514 735 968 • WIF1WNT2B 2123 489 720 1817 • SFRP4WNT3A 2127 506 1029 448 • TCF7L1WNT3A 2133 555 1460 1569 • GSK3BWNT2 2137 210 1106 91 • LEF1WNT2 2138 374 2350 1210 • FBXW2WNT2 2154 2353 1824 2104 • FZD1WNT3A 2170 611 1772 1213 • CSNK1G1WNT2B 2174 972 930 596 • TLE1WNT2B 2179 604 401 240 • LRP6WNT4 2183 310 1369 1948 • CSNK2A1WNT2B 2185 1900 83 748 • FBXW2WNT2B 2186 2375 732 346 • FZD2WNT1 2188 893 281 1938 • PPP2CAWNT2B 2190 1500 1434 86 • CSNK1DWNT3 2191 1636 274 500 • LEF1WNT1 2192 107 2238 490 • SFRP1WNT1 2193 142 1681 1072 • PPP2R1AWNT1 2196 958 641 980 • TCF7WNT3 2197 2031 1446 457 • LRP5WNT2B 2200 676 1251 26 • SFRP1WNT2B 2201 670 593 799 • WIF1WNT3 2204 1548 246 792 • TCF7L1WNT1 2213 127 1840 1900 • DVL2WNT3 2216 1491 81 630 • TLE1WNT1 2218 259 1223 1601 • FBXW11WNT2 2219 1443 1707 2257 • FRZBWNT2B 2220 1455 1809 16 • FZD1WNT1 2221 62 1237 1995 • SLC9A3R1WNT3A 2222 1114 1980 712 • FRZBWNT3 2224 1874 2455 435 • PITX2WNT3A 2225 215 1750 1554 • TCF7L1WNT2 2227 204 976 2152 • FZD1WNT2 2231 408 2069 1664 • NLKWNT3A 2236 2001 920 1919 • PPP2CAWNT1 2238 1737 2216 1735 • DAAM1WNT2 2243 1601 2266 414 • GSK3BWNT2B 2244 415 165 254 • TLE1WNT3 2245 2057 181 1864 • PITX2WNT1 2249 33 467 1382 • LEF1WNT3 2250 873 140 2240 • FBXW4WNT3 2256 1182 288 1250 • CTBP2WNT3 2257 2165 664 2335 • FZD6WNT3 2258 1929 66 1499 • GSK3BWNT3 2263 995 70 194 • DVL1WNT3 2264 2326 33 1784 • TCF7WNT3A 2268 1292 694 648 • FGF4WNT2 2269 378 1406 2130 • WNT1WNT3 2270 1591 340 946 • CSNK1G1WNT1 2272 132 1495 2233 • LEF1WNT3A 2273 712 2168 505 • FBXW4WNT3A 2274 748 1785 1584 • PPP2R1AWNT2B 2277 805 1121 1075 • TLE2WNT3 2280 2014 297 7 • CCND3WNT3A 2284 889 1281 2308 • SFRP4WNT3 2285 2333 888 644 • NLKWNT3 2287 2373 712 1982 • DVL2WNT2B 2301 1775 535 418 • DAAM1WNT3A 2302 1962 2405 1195 • LRP6WNT2B 2309 947 802 598 • LRP5WNT3A 2311 375 2117 1541 • DKK1WNT3 2312 2229 2060 1265 • FBXW11WNT3A 2313 1616 1148 2216 • GSK3BWNT3A 2314 522 1207 659 • TLE2WNT2B 2316 2278 398 17 • APCWNT3 2332 1469 1276 801 • BTRCWNT3 2335 1698 466 685 • CCND3WNT3 2337 1831 1134 759 • FBXW2WNT3 2338 1293 530 2124 • DAAM1WNT2B 2339 2283 265 2053 • FZD6WNT2B 2341 2315 132 1965 • DKK1WNT3A 2343 943 903 2477 LEF1WNT2B 2345 363 85 582 • CSNK2A1WNT3 2348 2151 127 2090 • SFRP4WNT1 2349 136 2142 567 • FZD7WNT3 2351 1322 12 678 • SFRP4WNT4 2353 250 2131 702 • PITX2WNT2B 2358 343 92 44 • FBXW11WNT2B 2360 2259 2111 773 • LRP6WNT2 2361 382 1338 1062 • FZD8WNT2B 2375 572 59 660 • FZD2WNT3A 2379 1495 2310 1820 • PPP2R1AWNT3 2380 2201 187 972 • FZD8WNT3 2381 2032 110 2309 • GSK3AWNT3 2383 1228 38 2370 • FZD7WNT1 2385 83 50 452 • TCF7L1WNT3 2391 2474 349 1490 • SENP2WNT1 2393 505 2173 361 • LRP6WNT1 2394 200 2465 2185 • FGF4WNT3A 2395 679 1962 1597 • FZD6WNT2 2396 1368 565 1119 • CXXC4WNT3 2400 1300 1015 2060 • EP300WNT3 2401 1642 514 1372 • FZD7WNT2B 2402 874 21 1740 • SENP2WNT2 2411 562 1345 1102 • FBXW4WNT2B 2413 1945 368 534 • FGF4WNT2B 2419 512 350 1577 • TCF7L1WNT2B 2423 871 519 485 • LRP6WNT3A 2424 610 665 2145 • SENP2WNT4 2426 347 1616 585 • PPP2CAWNT3 2428 2139 668 1203 • LRP5WNT3 2430 2267 198 356 • SFRP4WNT2B 2431 584 949 2071 • FZD2WNT3 2443 2319 54 1361 • DAAM1WNT3 2444 1494 120 2032 • CSNK1G1WNT3A 2451 632 1291 2485 • FZD1WNT3 2453 962 193 444 • SENP2WNT3 2454 2123 2420 1444 • SENP2WNT3A 2456 1372 1273 693 • FBXW11WNT3 2458 1759 1695 1850 • FZD6WNT3A 2465 1127 2473 1208 • SENP2WNT2B 2469 857 2068 2393 • FZD1WNT2B 2471 694 516 191 • NKD1WNT3 2474 1329 2101 656 • FGF4WNT3 2475 1912 214 449 • SFRP4WNT2 2478 351 1692 619 • CSNK1G1WNT3 2484 2446 225 1872

## FZD family 2*^nd^* order interaction ranking at different time points

• FRZBFZD2 9 930 923 955 10 • FZD7GSK3A 20 579 23 1892 1742 • FZD8GSK3A 42 772 122 323 1605 • FZD1JUN 43 1296 1287 2106 7 • FZD1GSK3A 47 1007 212 758 315 • FZD7PYGO1 54 1940 166 939 413 • FZD7WNT1 55 1790 81 1241 817 • FZD5WNT1 58 2346 161 1928 131 • FRZBFZD6 64 1953 44 2054 610 • FZD5GSK3A 72 311 163 109 1477 • FZD6GSK3A 75 452 2230 1009 1378 • DKK1FZD6 80 2163 2122 690 2355 • FZD7PORCN 83 188 673 445 170 • FZD7MYC 84 1160 43 2336 1274 • FRZBFZD8 88 2113 1353 1131 373 • FZD5JUN 94 741 1033 1110 2013 • FZD1PYGO1 95 56 308 2171 1220 • DKK1FZD2 97 2073 2046 30 2377 • FZD7WIF1 98 2037 1274 231 2303 • FZD5FOXN1 99 2379 89 45 1767 • FZD1FZD2 109 395 1277 763 1502 • FZD5CCND1 111 2278 1115 2385 105 • FZD1MYC 112 1291 756 266 2412 • FZD1PORCN 113 415 734 2401 273 • FZD1WNT1 124 2181 157 1473 195 • FZD1LEF1 142 1598 1219 544 2103 • FBXW11FZD2 144 353 808 187 2385 • FZD8WNT1 147 748 173 6 1085 • FZD1SFRP1 152 1100 1368 1476 136 • FRAT1FZD2 153 2202 2185 87 2073 • FZD5WIF1 154 1984 1110 2468 853 • FOSL1FZD2 159 854 367 208 989 • FZD5WNT3 164 913 1139 1633 192 • FZD8LEF1 167 120 718 1679 1343 • FZD7WNT3 168 493 1216 508 274 • FZD5PYGO1 170 1711 454 1243 1406 • FRZBFZD1 173 1890 727 46 879 • FZD6JUN 179 1265 2243 1910 1013 • FZD1WNT3 181 1834 909 831 681 • FZD5CSNK1A1 182 977 538 1672 2352 • APCFZD2 187 843 2115 567 1048 • DIXDC1FZD2 196 338 347 396 674 • FZD6PYGO1 208 331 1966 1269 896 • CSNK2A1FZD2 210 703 1468 1486 1004 • FZD7LEF1 212 1024 1136 3 1355 • FZD1WIF1 215 1929 2149 1822 1338 • FZD7SFRP1 220 443 1425 2132 323 • FZD8PYGO1 224 1063 280 398 1663 • FZD5LEF1 229 90 1331 1966 310 • CSNK1DFZD2 237 1453 2461 971 1474 • FZD7JUN 241 1223 1279 2162 1665 • CCND3FZD2 258 1731 2304 1894 2278 • FZD8JUN 259 1289 852 284 1451 • DKK1FZD8 262 2433 1479 2290 2224 • FZD1T 264 458 883 863 789 • FZD6WNT1 267 803 1938 299 1394 • FZD5MYC 274 2249 398 1323 2160 • CXXC4FZD2 275 1573 1211 1103 792 • FZD8WIF1 278 1843 1623 370 1266 • FZD6WIF1 284 2285 2173 1157 1072 • FZD5SFRP1 286 1217 1244 1784 166 • FZD5FZD2 288 1781 979 766 1114 • FZD5CCND2 298 905 477 1577 134 • FZD5PORCN 301 1354 2004 1414 53 • FZD5DVL1 302 1491 1084 852 396 • FZD1NKD1 306 392 291 658 2038 • FRZBFZD7 311 2241 1529 811 635 • FZD6MYC 319 1119 2250 1669 1280 • FZD5T 320 159 683 1644 1082 • FSHBFZD2 337 1393 2426 986 720 • FZD8MYC 344 381 587 1297 2253 • FZD7LRP6 360 1120 223 1920 571 • FZD5FGF4 362 93 2072 1655 359 • EP300FZD2 374 1467 1879 574 2363 • FBXW2FZD2 376 40 1175 1125 1759 • FZD6SFRP1 377 142 1983 1734 809 • FZD8PORCN 379 450 464 2190 504 • FZD7NKD1 388 948 169 589 2320 • FZD7T 394 1162 457 1879 1750 • FZD1LRP6 396 401 465 2398 607 • FZD5CTBP2 409 1583 1435 593 1263 • DKK1FZD1 427 2034 1539 1005 1775 • FZD5NKD1 432 1438 618 1324 745 • CTBP1FZD2 442 1207 802 1754 1411 • FZD6PORCN 448 551 1335 1252 55 • CTNNBIP1FZD2 450 1481 548 286 143 • CTNNB1FZD2 451 509 1424 2326 1744 • FZD6LEF1 452 348 2470 482 1240 • DKK1FZD7 455 2188 2058 1511 1816 • CTBP2FZD2 456 645 1801 899 1689 • FZD8SFRP1 465 817 1725 1806 463 • FZD7TLE1 467 2 2068 1675 1468 • FZD5LRP6 470 572 584 581 1064 • FZD8LRP6 475 1171 473 2017 1739 • APCFZD6 480 2229 189 480 971 • FZD6WNT3 494 16 2436 1371 776 • FZD5DVL2 496 675 889 334 2223 • FZD8WNT3 500 127 1621 2029 2318 • FZD1SFRP4 504 1714 1619 2234 237 • DAAM1FZD2 512 1433 2391 720 952 • FZD1TLE1 520 1923 2420 871 1351 • FZD5DAAM1 534 692 262 1426 94 • FZD5TLE1 537 1320 2069 1695 205 • FZD5CTNNBIP1 554 1661 1513 2275 2203 • FZD8T 555 719 427 1119 1692 • CSNK2A1FZD6 556 258 922 1717 188 • CSNK1G1FZD2 563 1738 2268 523 1143 • FZD1FZD6 564 2008 369 904 585 • FZD1FBXW4 570 504 561 2161 1305 • DVL2FZD2 574 367 1449 642 1431 • FZD7PITX2 581 1197 769 2320 494 • FZD6LRP6 582 976 2428 2447 2332 • FZD1PITX2 589 1455 1998 2127 456 • FOSL1FZD8 593 1516 876 199 660 • FZD1NLK 594 1311 269 691 290 • FZD6T 598 92 1558 1495 2392 • FZD1TCF7 599 379 582 2335 2034 • FZD7PPP2R1A 602 13 208 2441 1936 • FZD1PPP2CA 605 1893 306 1913 1892 • FZD5CSNK1G1 609 1137 484 2000 577 • FZD1FZD8 611 12 1979 1451 130 • FZD7KREMEN1 612 792 578 2166 1587 • FZD7FBXW4 614 745 178 2081 2461 • FOSL1FZD6 615 812 83 2065 1424 • APCFZD8 624 2116 2160 1058 316 • FZD1SLC9A3R1 629 47 2134 1932 475 • APCFZD5 637 729 1301 1851 1439 • BTRCFZD2 641 466 2031 1152 1257 • FBXW11FZD6 643 49 1989 1116 2444 • FZD5FBXW4 648 1963 383 2062 485 • FZD8NKD1 649 1827 326 1574 2416 • AXIN1FZD2 655 87 1678 156 2200 • FZD7PPP2CA 662 642 32 247 1583 • CSNK1DFZD8 669 2068 2145 2307 233 • FZD2SFRP1 679 761 1570 1972 243 • FZD5CTNNB1 684 1045 1370 1034 1840 • FZD1WNT2B 693 277 105 2424 1609 • FZD1PPP2R1A 706 1641 1385 1614 395 • FGF4FZD2 708 302 156 1 286 • FZD5CCND3 718 2014 154 2023 814 • FZD7FZD8 723 323 1050 2148 393 • FZD5PITX2 726 1000 2099 1166 640 • FZD7SFRP4 727 2124 1298 1969 799 • FZD1KREMEN1 738 586 1740 1630 474 • FRAT1FZD8 745 2344 1974 212 787 • FZD7TCF7 747 595 38 479 1993 • FZD1WNT4 748 2153 297 2423 797 • DIXDC1FZD6 755 1829 3 2080 944 • FRAT1FZD6 757 1720 268 1629 574 • FZD2PYGO1 759 1301 384 1276 508 • FZD7SLC9A3R1 765 1570 1167 1861 416 • FZD2GSK3A 767 1457 137 352 1358 • FZD5KREMEN1 773 530 513 2347 79 • CSNK2A1FZD8 778 202 2049 1203 628 • FZD5FZD6 781 590 158 1890 851 • FZD8FBXW4 788 1612 209 640 2357 • FZD5SFRP4 795 918 2105 2299 2276 • FZD5CTBP1 799 604 867 2225 2135 • CSNK1DFZD6 800 2060 846 1138 1356 • FZD5FSHB 802 915 144 22 1672 • FZD1WNT5A 805 1924 307 1120 165 • FZD7WNT2 815 173 1242 84 1788 • FBXW11FZD8 817 1577 1454 1362 1544 • AESFZD2 819 1996 1256 1865 1331 • FZD5EP300 824 870 1855 1878 830 • FZD6TLE1 827 1728 1874 2379 1271 • FZD7GSK3B 834 1117 1510 1985 976 • FZD1WNT2 838 222 1540 235 1256 • FZD1LRP5 850 472 1030 2101 1429 • FZD8KREMEN1 860 14 420 2259 1965 • FZD7NLK 867 2314 50 1333 805 • CCND2FZD2 870 249 1470 2363 1493 • FZD5PPP2R1A 879 1118 946 1416 216 • FZD1WNT3A 881 482 2298 2417 1337 • FZD6NKD1 883 1727 1234 586 1554 • FZD7WNT5A 887 1403 41 1466 1629 • FZD5SLC9A3R1 891 71 1393 2010 967 • FOSL1FZD7 897 2059 1663 1956 704 • FZD5PPP2CA 902 2331 240 416 1264 • FZD7WNT3A 910 1757 705 2105 390 • FZD1TLE2 911 602 706 2147 1208 • FZD1GSK3B 920 762 815 358 1875 • FZD7WNT4 926 1925 694 949 1046 • FZD8TLE1 927 998 585 882 1659 • DIXDC1FZD8 934 1325 401 1091 1541 • FZD7LRP5 943 1218 455 2374 2044 • APCFZD7 950 2406 1625 1367 903 • FZD7TLE2 953 1822 1120 105 1033 • FZD6FBXW4 960 1357 1658 1703 1094 • FZD7RHOU 965 287 1405 326 663 • FZD5FBXW2 966 1696 893 1023 225 • FZD5WNT3A 981 822 1946 2243 468 • CXXC4FZD6 987 1430 183 2220 2421 • FZD8SFRP4 1009 1935 1451 1740 2297 • FZD5CXXC4 1028 667 1192 124 52 • FZD5TCF7 1033 1247 253 2030 1030 • FZD7WNT2B 1038 328 29 2174 1849 • FZD5TLE2 1049 1802 556 1329 945 • CSNK1A1FZD2 1052 1903 1799 435 140 • FZD5WNT5A 1059 1553 543 1566 193 • FOSL1FZD1 1060 1458 779 1014 2230 • FZD6SLC9A3R1 1063 1235 2345 2328 930 • APCFZD1 1065 1913 1395 525 1255 • FBXW11FZD7 1069 106 1981 1733 2317 • FZD1RHOU 1071 513 941 2285 564 • DVL1FZD2 1084 1025 1010 44 1733 • FZD5GSK3B 1086 1447 1086 2282 1985 • FZD5NLK 1091 1600 59 670 486 • CXXC4FZD8 1093 2022 1178 1871 2466 • CCND3FZD6 1097 538 426 2012 907 • FZD5CSNK1D 1108 535 462 1402 946 • FZD8NLK 1109 672 351 1612 1890 • FZD6PPP2CA 1119 204 1164 1951 1967 • FZD6NLK 1133 768 2307 1455 1161 • FZD1FZD7 1136 1795 2383 1627 722 • FZD5LRP5 1153 192 586 1943 1404 • FZD5WNT2 1157 1319 2227 1535 2036 • FZD5FRAT1 1169 1414 1574 2145 2069 • FZD6PITX2 1173 889 1905 1477 267 • BCL9FZD2 1178 301 1726 804 2019 • FRAT1FZD7 1180 2178 2330 509 458 • CCND3FZD8 1193 1178 1922 1647 1403 • FZD8GSK3B 1194 1892 795 422 2109 • FZD5WNT4 1196 514 621 2391 2362 • CTBP1FZD6 1204 1904 65 514 291 • FZD8PITX2 1206 618 1598 573 506 • FZD2WNT1 1208 1049 26 1665 398 • EP300FZD8 1211 72 2336 1463 200 • FZD5WNT2B 1212 1377 94 1359 1753 • FZD6GSK3B 1219 778 1762 383 2371 • FOXN1FZD2 1220 267 1806 86 1027 • FZD1TCF7L1 1223 1749 1351 2446 846 • FBXW11FZD1 1228 380 1793 1485 2376 • EP300FZD7 1232 897 2213 1098 1420 • CSNK2A1FZD1 1234 41 1051 2141 736 • FZD8SLC9A3R1 1243 845 1701 374 306 • EP300FZD6 1245 734 362 561 1230 • FZD5CSNK2A1 1249 1463 397 301 2028 • FZD2MYC 1258 1629 344 1180 1530 • FZD6WNT2 1262 649 1471 648 275 • FZD6FZD8 1267 636 2095 1244 1135 • FZD8PPP2CA 1274 1730 77 1712 1876 • FZD5FZD8 1277 194 2337 2086 20 • FZD7TCF7L1 1295 191 695 2479 533 • FZD8WNT4 1296 810 1566 1162 942 • FZD6SFRP4 1303 1485 1286 461 781 • FZD6KREMEN1 1305 15 1729 1520 1281 • FZD5RHOU 1315 1871 1474 706 1373 • FZD8WNT5A 1316 73 334 1596 676 • FZD8WNT2B 1319 756 109 427 2166 • FZD6WNT5A 1320 1231 1375 2168 445 • FZD6PPP2R1A 1324 724 1914 713 545 • FZD8PPP2R1A 1330 247 1007 822 854 • FBXW2FZD6 1333 2127 430 1859 2058 • FZD5DIXDC1 1337 813 2006 835 484 • FZD8TCF7 1338 895 264 1471 2442 • FZD8LRP5 1343 359 806 1481 1805 • DAAM1FZD8 1348 1424 2067 803 1708 • FRAT1FZD1 1363 1707 496 911 847 • FZD6WNT3A 1364 102 2388 2192 891 • FZD6TCF7 1376 1230 675 979 1832 • FZD2JUN 1377 638 680 956 970 • DAAM1FZD6 1378 1011 424 539 2390 • DIXDC1FZD7 1384 1801 540 2370 1842 • FBXW2FZD8 1389 582 1890 1545 2182 • CSNK2A1FZD7 1401 330 2183 2079 753 • FZD8WNT2 1403 2424 952 968 772 • FZD8TLE2 1410 932 595 1421 246 • CSNK1DFZD1 1419 2048 665 1345 469 • FZD8WNT3A 1422 706 1145 1628 1499 • FZD5FZD7 1423 1017 1738 1662 867 • FSHBFZD8 1435 841 2239 2245 482 • FZD5FBXW11 1438 429 363 1179 346 • FZD6WNT4 1445 946 1846 1188 1696 • CTBP1FZD8 1461 1638 1020 717 63 • CSNK1DFZD7 1462 2135 2193 998 843 • FBXW2FZD1 1463 1182 1496 689 2245 • FZD2WIF1 1466 1586 1526 825 1658 • FZD5FZD1 1468 1419 554 1607 1857 • FZD5FOSL1 1478 1298 2157 1051 1919 • CTNNB1FZD8 1482 1645 976 1592 1626 • FSHBFZD6 1484 519 1108 473 784 • FZD2PORCN 1496 64 719 1449 84 • CTNNBIP1FZD6 1500 1518 251 598 212 • CXXC4FZD1 1528 1444 1568 1325 1219 • FZD8RHOU 1542 698 1544 2185 2381 • FZD6LRP5 1545 522 2280 872 1618 • FZD6RHOU 1550 1300 1958 656 160 • FZD5TCF7L1 1558 1584 1067 2129 1440 • DIXDC1FZD1 1559 993 290 841 2434 • CSNK1G1FZD8 1564 2414 2021 1248 2438 • FZD6WNT2B 1568 205 1183 1538 1922 • EP300FZD1 1597 88 1953 512 2076 • DVL2FZD6 1598 1876 627 345 627 • DVL2FZD8 1616 1303 1564 1900 675 • FZD6FZD7 1632 1683 1797 1262 818 • FZD6TLE2 1635 409 2076 987 2273 • CSNK1G1FZD6 1654 2203 1071 446 2437 • CTNNB1FZD6 1665 725 222 1321 283 • AESFZD6 1673 270 841 1209 1510 • DVL1FZD6 1675 1468 270 1935 703 • FZD2WNT3 1679 1758 958 675 270 • FSHBFZD7 1680 1668 1681 278 1222 • CCND1FZD2 1683 225 1228 1975 873 • CTBP1FZD1 1690 1864 881 584 511 • FZD2LEF1 1697 495 570 1078 823 • FZD6TCF7L1 1709 2082 2195 1280 2328 • FBXW2FZD7 1711 136 2013 2357 997 • CTNNB1FZD7 1716 594 957 1431 752 • CTBP2FZD6 1721 431 316 2429 1772 • FZD2LRP6 1725 75 599 133 401 • CCND3FZD7 1730 269 1117 298 1416 • BTRCFZD8 1731 2385 2322 844 433 • FSHBFZD1 1739 1832 1918 613 2095 • CTBP2FZD8 1744 1242 1557 1819 1148 • CXXC4FZD7 1753 2465 1467 1493 2147 • AXIN1FZD6 1809 866 730 449 227 • DVL1FZD8 1828 1406 894 1074 368 • CTNNBIP1FZD8 1832 2351 790 151 13 • CTNNBIP1FZD1 1834 1130 1412 2094 490 • BCL9FZD6 1836 2330 446 222 1969 • FZD8TCF7L1 1844 1057 682 1153 1825 • CCND3FZD1 1878 435 1506 1758 1216 • BTRCFZD6 1880 498 1452 1391 923 • DAAM1FZD1 1882 1693 1677 2032 1570 • FZD2T 1890 909 515 2050 206 • DAAM1FZD7 1898 1748 2199 1494 2064 • CTNNBIP1FZD7 1899 2392 2447 375 1 • CTBP1FZD7 1908 1323 2384 360 38 • AXIN1FZD8 1914 1213 1197 1211 583 • FGF4FZD6 1931 1826 37 1876 347 • FZD5DKK1 1943 1328 258 2084 1010 • FZD2TLE1 1946 1652 664 733 366 • FZD2NKD1 1950 1828 466 1613 735 • FZD2PITX2 1956 1285 1427 1810 264 • CTBP2FZD7 1968 489 2410 2383 1460 • DVL2FZD7 1975 445 1159 782 1288 • CSNK1G1FZD7 1977 1469 1949 2304 2419 • BTRCFZD1 1978 55 1764 1516 2120 • CCND2FZD6 1980 1712 588 2297 1120 • CTNNB1FZD1 1983 1544 1214 1435 1720 • FZD2FBXW4 1990 2132 213 617 284 • FZD2TCF7 1992 1456 48 1531 1301 • CSNK1G1FZD1 2005 868 1744 2068 1620 • AESFZD7 2028 627 2135 2269 1895 • CSNK1A1FZD6 2041 2101 453 607 655 • BCL9FZD8 2070 1621 1354 1347 2199 • CCND2FZD8 2072 1391 1820 934 1913 • CCND1FZD8 2073 197 1777 1202 2324 • CCND2FZD1 2075 536 2294 392 266 • FZD2KREMEN1 2082 736 1122 1373 604 • FZD2WNT2 2083 1807 758 357 1192 • BTRCFZD5 2090 1421 1973 1192 768 • FZD2RHOU 2092 1155 1763 1043 795 • FZD2NLK 2093 1589 198 918 72 • FOXN1FZD8 2099 349 2111 2475 715 • FZD2SLC9A3R1 2102 685 1348 1393 992 • CSNK1A1FZD8 2110 1521 2198 372 329 • FZD2FZD8 2117 2441 371 2004 771 • FZD2FZD6 2122 406 277 781 849 • AXIN1FZD7 2125 1417 1624 1705 1390 • FZD2SFRP4 2130 957 1916 159 362 • AXIN1FZD5 2137 1252 1853 1875 2449 • FZD2LRP5 2144 1214 539 2015 172 • FZD2PPP2R1A 2149 1596 403 1711 19 • AESFZD8 2150 1001 1911 1360 2052 • DVL2FZD1 2151 1762 1522 2159 1710 • FZD2WNT2B 2177 1528 120 1316 763 • BCL9FZD5 2180 970 1736 2137 2169 • FOXN1FZD6 2200 345 1094 18 1018 • AESFZD5 2204 1086 2290 1160 2473 • DVL1FZD1 2215 217 1155 1106 2275 • FGF4FZD8 2217 1519 729 837 2144 • AXIN1FZD1 2218 1719 1789 1081 1050 • AESFZD1 2235 899 1384 276 497 • FZD2TLE2 2238 1686 1532 8 185 • DVL1FZD7 2240 596 2234 546 770 • CTBP2FZD1 2255 237 1247 810 2066 • FZD2GSK3B 2273 1534 1079 2131 953 • FZD2WNT4 2280 1372 600 1988 1835 • BTRCFZD7 2286 521 1631 843 1588 • CCND2FZD7 2290 2053 1495 2183 948 • CCND1ZD6 2297 365 321 1290 2265 • FZD2WNT3A 2301 1552 1327 1302 1709 • FZD5FRZB 2303 506 1000 1185 2343 • FZD2WNT5A 2304 1101 632 1964 190 • CSNK1A1FZD1 2316 1258 678 1429 2296 • FZD2PPP2CA 2322 1776 288 2321 600 • FZD2FZD7 2327 1769 2363 2123 328 • FGF4FZD1 2332 1692 474 1994 1820 • FOXN1FZD7 2333 708 2113 172 590 • BCL9FZD7 2346 654 1757 163 522 • FZD2TCF7L1 2349 479 438 498 746 • CCND1FZD1 2350 526 927 1803 677 • FZD7SENP2 2354 883 2216 1132 1032 • FZD1SENP2 2357 266 1235 165 451 • CCND1FZD7 2381 357 2417 1320 2181 • BCL9FZD1 2396 1764 1144 1214 2193 • FGF4FZD7 2404 1413 1596 678 1137 • FZD5SENP2 2406 1420 1009 1642 25 • CSNK1A1FZD7 2418 1644 1881 1942 588 • FOXN1FZD1 2421 1477 961 410 2347 • FZD6SENP2 2439 2010 1213 2258 1102 • FZD8SENP2 2458 2417 1971 137 880 • FZD2SENP2 2468 1803 2109 27 483

## FZD family 2*^nd^* order interaction ranking at different durations

• FZD5SFRP1 2 1461 1499 1726 • FZD5CSNK2A1 3 1099 646 1647• FZD5CSNK1G1 4 2430 1410 2054 • FZD5SENP2 7 1110 2167 1402 • FZD5FBXW2 8 921 2298 1953 • FZD5FZD7 10 2424 509 2303 • FZD5LRP6 11 2235 1633 1593 • FZD5SFRP4 12 1326 2012 926 • FZD5LEF1 13 1838 758 937 • FZD5FGF4 15 2225 1399 2305 • FZD5DKK1 16 2181 2313 2462 • CSNK1A1FZD7 17 1782 481 315 • FZD5FRZB 19 525 343 963 • FZD5WNT2 26 897 1803 1450 • CSNK1A1FZD2 28 1633 1220 681 • FZD5FZD1 34 1276 2019 1592 • FZD5FBXW11 35 549 421 2435 • FZD5TCF7L1 40 2452 2050 1910 • FZD5FBXW4 44 455 1805 893 • FZD5GSK3B 45 2414 1261 2038 • CSNK1A1FZD1 46 1728 1901 1723 • FZD5WNT4 49 463 945 2348 • FZD5FZD8 52 1963 714 2426 • FZD5CTBP2 55 950 2306 2466 • FZD5LRP5 62 2047 2279 658 • CTNNB1FZD2 67 2280 1700 584 • FZD5NLK 68 580 1551 1916 • FZD5FZD2 69 865 1054 2003 • CTBP1FZD2 74 2285 654 1474 • FZD5CXXC4 75 46 1869 1502 • CSNK1A1FZD8 77 1175 740 1307 • FZD5PPP2R1A 88 786 1605 1535 • FZD5TCF7 89 797 804 1610 • FZD5PITX2 95 2002 1131 1123 • FZD5TLE1 100 2328 1393 1765 • CTNNB1FZD6 112 1818 1028 160 • FZD5FZD6 115 1933 404 1399 • CTBP1FZD7 119 1915 298 1013 • DIXDC1FZD2 124 248 1200 1194 • CSNK1A1FZD6 126 2195 807 59 • FZD5PPP2CA 128 470 2021 1904 • FZD5WNT5A 136 579 1528 2296 • FZD5CTNNB1 145 159 1846 2321 • CTNNB1FZD7 148 1680 1091 636 • DIXDC1FZD1 156 788 1067 875 • CCND2FZD2 158 976 853 1673 • DIXDC1FZD7 160 808 1898 2142 • FZD5FOSL1 161 1882 1878 1981 • FZD5DAAM1 162 623 1933 2030 • FZD5CTNNBIP1 165 367 1828 1297 • CTBP1FZD8 180 1432 524 1741 • CTNNB1FZD8 187 1814 1553 2481 • CTNNB1FZD1 188 1652 2415 2237 • FZD5NKD1 189 184 687 2270 • FZD5TLE2 192 638 1763 853 • CCND2FZD7 201 1487 355 2055 • CCND1FZD2 208 1829 1254 1987 • CTBP1FZD1 211 1730 1533 2325 • AESFZD2 215 2198 957 283 • FZD5DIXDC1 229 1223 1143 1387 • AXIN1FZD1 235 1911 372 1003 • FZD5CCND3 239 500 2250 2312 • CCND1FZD1 243 1665 1328 869 • AESFZD1 244 1247 2121 805 • CTBP1FZD6 248 1726 648 888 • FZD5DVL2 273 884 1602 2092 • AXIN1FZD2 274 806 2054 2209 • CTNNBIP1FZD2 282 2411 2052 852 • FZD5EP300 285 178 2138 2349 • DIXDC1FZD8 289 1371 1680 1150 • FZD5CCND1 290 662 1582 2197 • AXIN1FZD7 294 2384 1364 2039 • BCL9FZD7 295 1430 560 103 • FOXN1FZD2 297 1966 1646 867 • AXIN1FZD8 303 2211 2129 1804 • DIXDC1FZD6 307 306 1736 789 • BCL9FZD1 325 2268 2333 1070 • FOXN1FZD7 330 1323 822 159 • FRAT1FZD2 341 991 827 1603 • CCND1FZD7 342 2364 1725 1680 • FSHBFZD7 344 1579 700 1863 • AESFZD7 362 1498 826 219 • FRAT1FZD7 365 1202 859 128 • CTNNBIP1FZD1 366 1561 626 1011 • FZD5FOXN1 367 692 1610 1098 • CSNK1DFZD2 369 713 2070 2278 • AESFZD8 374 1261 1517 984 • CSNK1DFZD7 375 1402 1158 304 • BCL9FZD2 382 1168 2194 1053 • FRAT1FZD6 384 1528 1125 117 • CCND2FZD1 385 1394 2006 1651 • FOXN1FZD1 387 2250 1660 2453 • CCND1FZD6 391 1977 2285 472 • FZD5DVL1 394 519 1754 757 • CTNNBIP1FZD8 395 2178 2009 1761 • DVL1FZD2 401 1384 2105 368 • FOXN1FZD6 403 2438 1335 154 • CTNNBIP1FZD7 405 1822 2011 553 • FSHBFZD2 409 593 2049 416 • FZD5SLC9A3R1 416 1265 1813 2020 • CTNNBIP1FZD6 417 2224 1238 1168 • FZD5GSK3A 425 1872 1099 2300 • FSHBFZD6 426 751 986 1441 • AESFZD6 436 2159 1823 9 • FRAT1FZD1 437 1348 1383 328 • CCND2FZD6 449 1274 905 2002 • CSNK1DFZD1 455 1799 2017 2459 • FSHBFZD1 460 1685 2175 904 • FZD8SENP2 471 379 60 287 • FRAT1FZD8 472 1150 1269 2102 • BTRCFZD2 473 1964 1861 1661 • APCFZD2 478 1569 673 2040 • FZD5CSNK1D 479 1258 2066 1283 • BCL9FZD6 483 1588 1151 819 • FZD6SENP2 489 2405 11 807 • CCND2FZD8 502 1460 961 1556 • FZD5CCND2 503 364 2025 2324 • BCL9FZD8 518 2303 1437 2112 • CSNK1DFZD6 527 879 1440 341 • DVL1FZD1 536 833 365 478 • CTBP2FZD7 537 1003 653 2162 • BTRCFZD7 538 1253 769 115 • AXIN1FZD6 548 1590 2058 1477 • DVL1FZD7 553 1319 1866 725 • FOSL1FZD2 556 387 1963 1305 • FZD5WNT1 564 617 2284 1432 • CTBP2FZD2 565 2255 1910 1395 • CSNK1DFZD8 574 1600 1580 830 • APCFZD7 577 1380 609 1639 • FZD1SENP2 584 432 438 184 • FSHBFZD8 588 2024 1235 974 • BTRCFZD1 591 2102 1342 378 • BTRCFZD6 592 2234 1304 447 • APCFZD1 594 1194 1594 719 • FZD5WIF1 601 2106 2092 889 • FOXN1FZD8 603 2174 1325 680 • CTBP2FZD6 607 687 946 722 • FOSL1FZD8 615 2028 1334 2416 • FZD5FRAT1 617 775 1210 1579 • FOSL1FZD1 623 2192 2234 2429 • CCND1FZD8 624 2184 2229 1829 • FZD6SFRP4 629 975 175 866 • DVL1FZD6 656 2292 1875 1835 • APCFZD8 658 1045 586 2449 • FZD5MYC 663 1173 1799 2341 • DVL2FZD7 668 1234 1558 1896 • CTBP2FZD1 682 2167 2126 939 • FZD2SENP2 695 517 197 542 • FOSL1FZD7 701 1497 571 956 • FRZBFZD2 703 1026 1634 864 • DVL2FZD2 705 2448 2088 1209 • DVL1FZD8 706 1259 2474 1084 • FZD7SENP2 711 1423 8 978 • EP300FZD7 712 1722 1222 1112 • DVL2FZD1 716 2344 764 1635 • FZD5FSHB 725 907 1565 1267 • FZD8SFRP4 733 724 57 382 • BTRCFZD8 757 1850 1900 2125 • APCFZD6 762 1232 971 886 • DKK1FZD7 768 1260 400 1489 • FZD8LRP6 769 559 100 1434 • EP300FZD2 771 1914 1968 1715 • FZD5WNT2B 803 1896 808 1089 • FZD5KREMEN1 816 400 1793 1629 • FOSL1FZD6 821 496 1541 155 • FZD5CTBP1 830 482 1072 2252 • FRZBFZD7 840 1841 1262 54 • DKK1FZD2 847 422 1093 2231 • CTBP2FZD8 853 1864 562 1777 • FZD1SFRP4 855 540 224 69 • EP300FZD1 870 1788 2373 1380 • DKK1FZD1 878 1602 2200 1413 • CCND3FZD7 887 2176 918 1559 • FZD5WNT3A 902 2287 1744 1093 • FZD2SFRP4 906 1061 155 433 • CCND3FZD2 915 2276 1922 1516 • FZD5PYGO1 925 1205 2156 2193 • EP300FZD8 928 1330 2211 1463 • DAAM1FZD7 929 2356 1357 2280 • CCND3FZD1 935 1032 697 1852 • DVL2FZD8 947 2072 1486 2389 • FZD6LRP6 948 1770 103 1889 • DAAM1FZD2 960 1263 1986 731 • DVL2FZD6 964 2383 1752 550 • FZD7SFRP4 977 1976 13 2198 • FRZBFZD1 993 1273 2458 1570 • FRZBFZD6 1004 2479 1448 33 • FZD5JUN 1007 582 2426 770 • FZD5PORCN 1011 863 1856 2460 • CXXC4FZD7 1014 1077 497 1688 • FZD6LEF1 1022 2293 474 828 • CXXC4FZD2 1028 2056 984 1378 • DAAM1FZD1 1035 1360 2312 537 • FZD1LRP6 1048 1052 1727 1498 • DKK1FZD8 1055 487 781 2256 • FZD8LEF1 1058 979 170 1720 • FZD5RHOU 1067 840 1063 992 • FRZBFZD8 1072 1944 1069 1774 • CCND3FZD8 1073 1174 889 2334 • FBXW2FZD7 1083 1310 490 1643 • EP300FZD6 1086 2130 2137 502 • FZD8PPP2R1A 1114 95 449 699 • FZD8SFRP1 1120 1668 932 915 • FZD6PPP2R1A 1133 602 154 533 • CXXC4FZD1 1135 1620 1747 429 • FZD6FZD7 1139 2312 747 1021 • FBXW2FZD1 1143 1827 1722 1636 • CSNK2A1FZD7 1155 2055 1770 1999 • FZD7LRP6 1168 1580 48 1918 • FZD6TCF7L1 1171 1172 117 1637 • FZD8TCF7L1 1176 684 280 1818 • FZD5T 1182 616 1738 1523 • FZD7LEF1 1190 2179 218 1175 • FZD6SFRP1 1200 1454 464 1946 • FZD1PPP2R1A 1203 117 1936 1143 • DAAM1FZD6 1209 908 1386 1104 • FZD6WNT5A 1213 1165 162 1497 • FZD1LEF1 1229 717 2113 594 • FZD1FZD6 1237 202 901 81 • CCND3FZD6 1238 1913 2023 800 • FZD1FZD2 1253 272 2255 1649 • FBXW11FZD2 1255 2352 940 957 • FZD6GSK3B 1258 1116 219 524 • FBXW11FZD7 1266 919 785 1176 • FZD8PITX2 1269 1080 1147 257 • FZD1FZD7 1270 1960 1026 313 • DKK1FZD6 1280 466 737 494 • FZD7TCF7L1 1281 1530 43 2004 • CXXC4FZD6 1295 2379 1036 1174 • CXXC4FZD8 1298 1572 929 1816 • FZD6PPP2CA 1300 1658 102 704 • DAAM1FZD8 1302 1342 1491 1239 • FBXW2FZD2 1308 2003 1620 2229 • FZD7SFRP1 1311 1796 136 2362 • FBXW11FZD1 1315 2039 2147 1846 • FZD8WNT5A 1316 41 294 2474 • FZD1TCF7L1 1318 666 1114 2157 • FZD6NLK 1334 502 5 1465 • FZD6PITX2 1336 2260 759 574 • FGF4FZD2 1342 390 1808 2223 • FZD7PPP2R1A 1360 120 49 2284 • FZD8GSK3B 1364 669 1088 1409 • FZD2LEF1 1374 980 1198 1799 • CSNK2A1FZD1 1401 1825 635 2361 • FBXW2FZD6 1402 2050 1206 1026 • FGF4FZD1 1403 491 1193 2191 • FZD1GSK3B 1407 1411 909 1947 • FZD8PPP2CA 1415 75 361 1857 • FZD2LRP6 1425 1786 358 1962 • FBXW2FZD8 1427 1762 1246 1902 • FZD7NLK 1429 81 16 2203 • FZD8TCF7 1432 125 40 2379 • FZD7GSK3B 1441 1830 896 276 • FGF4FZD7 1448 1364 1513 737 • FZD1PITX2 1453 1340 1893 2113 • FZD6LRP5 1455 2426 266 61 • FZD5WNT3 1463 1358 1232 1333 • FZD1NLK 1465 168 39 767 • CSNK2A1FZD2 1474 1941 1292 2005 • FBXW11FZD8 1479 2266 741 1524 • FZD2PPP2R1A 1485 934 255 1571 • FZD2FZD7 1490 1766 2161 1371 • FZD8WNT2 1499 186 1282 1890 • FZD1SFRP1 1508 1682 2215 410 • FBXW11FZD6 1528 2449 546 305 • FZD7PITX2 1538 2059 624 1421 • FZD5CSNK1A1 1542 554 996 1886 • FZD8TLE2 1552 88 873 280 • FZD7WNT5A 1553 149 79 2181 • FZD6FZD8 1556 2044 1249 1580 • CSNK1G1FZD1 1566 1748 2400 2170 • FZD2TCF7L1 1570 815 250 1785 • CSNK1G1FZD2 1571 451 1774 2404 • CSNK2A1FZD8 1579 2037 1358 2211 • FZD8NLK 1582 25 7 2409 • FZD6FBXW4 1585 371 263 460 • FZD7PPP2CA 1587 182 14 1295 • CSNK1G1FZD8 1594 1744 1112 2264 • FZD1WNT5A 1602 146 839 2217 • FZD2GSK3B 1631 2365 1927 927 • FZD8TLE1 1635 277 375 1244 • FZD1FZD8 1644 2389 1699 2415 • FGF4FZD8 1646 1716 1242 1626 • FZD1TLE1 1648 755 1742 822 • FZD6TLE1 1654 2272 220 1091 • FZD2FZD8 1658 2362 2203 2011 • CSNK1G1FZD6 1661 274 174 620 • CSNK1G1FZD7 1667 900 1146 1049 • FZD2PITX2 1672 1397 1250 1205 • FZD2PPP2CA 1680 645 836 2232 • FZD2TCF7 1688 1571 3 1092 • FZD7TCF7 1693 196 4 774 • FGF4FZD6 1705 476 1717 2000 • FZD7TLE2 1712 313 167 227 • FZD7WNT4 1724 287 537 677 • FZD7FZD8 1731 2461 1202 1934 • FZD6TLE2 1739 837 231 67 • FZD6TCF7 1745 982 18 1066 • CSNK2A1FZD6 1747 1724 691 971 • FZD7FBXW4 1767 92 252 1337 • FZD1PPP2CA 1768 71 1303 897 • FZD2TLE2 1777 564 1064 75 • FZD2SFRP1 1785 1479 761 696 • FZD6WNT4 1786 1018 766 602 • FZD8WNT4 1803 124 1666 1968 • FZD1TCF7 1808 177 34 1044 • FZD2TLE1 1810 1739 380 1200 • FZD2FZD6 1812 1086 1790 661 • FZD2NLK 1821 457 75 2155 • FZD1WNT4 1825 58 1445 1186 • FZD1FBXW4 1846 28 2332 579 • FZD7LRP5 1860 1934 153 132 • FZD1LRP5 1864 1768 619 1043 • FZD2WNT4 1871 434 2352 2388 • FZD6SLC9A3R1 1872 1006 443 1501 • FZD2WNT2 1875 435 1451 1562 • FZD1NKD1 1902 66 331 99 • FZD2WNT5A 1908 420 397 1428 • FZD7TLE1 1914 998 318 2340 • FZD8FBXW4 1915 56 733 1379 • FZD7GSK3A 1917 1254 286 1385 • FZD1GSK3A 1920 711 1256 510 • FZD6GSK3A 1921 1362 686 1207 • FZD1TLE2 1934 222 868 1024 • FZD7NKD1 1956 16 6 1152 • FZD8LRP5 1971 1856 448 543 • FZD8NKD1 1975 5 62 2028 • FZD6KREMEN1 2005 821 821 665 • FZD2WIF1 2012 1862 869 310 • FZD8GSK3A 2017 556 1104 1105 • FZD1WIF1 2033 685 1579 319 • FZD7SLC9A3R1 2036 344 387 2443 • FZD7WNT2 2041 299 326 811 • FZD6WIF1 2051 1250 216 1136 • FZD6WNT1 2059 425 183 667 • FZD8WIF1 2076 733 271 1410 • FZD8SLC9A3R1 2084 284 1885 621 • FZD7KREMEN1 2087 113 226 743 • FZD8WNT1 2094 42 260 493 • FZD2MYC 2095 504 90 1849 • FZD2NKD1 2096 444 144 1548 • FZD2WNT2B 2101 1563 210 281 • FZD8WNT3A 2111 426 1622 1988 • FZD7WNT3A 2121 514 735 968 • FZD7MYC 2134 193 10 1455 • FZD6T 2149 744 156 62 • FZD2LRP5 2158 2327 810 1124 • FZD6NKD1 2163 411 121 1319 • FZD7JUN 2164 163 44 373 • FZD1JUN 2169 195 806 679 • FZD1WNT3A 2170 611 1772 1213 • FZD1KREMEN1 2176 139 1959 2097 • FZD2SLC9A3R1 2182 1076 1706 782 • FZD8MYC 2187 52 42 917 • FZD2WNT1 2188 893 281 1938 • FZD1SLC9A3R1 2199 485 1615 1645 • FZD2FBXW4 2207 380 1382 1860 • FZD2GSK3A 2210 1244 1350 1156 • FZD1WNT1 2221 62 1237 1995 • BTRCFZD5 2223 1551 1508 1745 • FZD1WNT2 2231 408 2069 1664 • FZD6MYC 2246 1060 56 580 • FZD6PORCN 2251 795 228 1960 • FZD6WNT3 2258 1929 66 1499 • FZD8PYGO1 2278 115 293 1546 • FZD2RHOU 2279 1266 2294 993 • FZD2T 2294 477 123 1046 • FZD2JUN 2308 697 285 1347 • FZD6RHOU 2315 1635 1162 273 • FZD1T 2320 253 1374 1594 • FZD6JUN 2321 648 161 307 • FZD8RHOU 2322 401 2213 2115 • FZD8KREMEN1 2323 40 1914 2126 • FZD8T 2328 82 73 754 • BCL9FZD5 2331 1094 1013 2077 • FZD6WNT2B 2341 2315 132 1965 • FZD7WNT3 2351 1322 12 678 • FZD8JUN 2365 72 334 1773 • FZD1MYC 2366 67 906 1366 • FZD8WNT2B 2375 572 59 660 • FZD2WNT3A 2379 1495 2310 1820 • FZD8WNT3 2381 2032 110 2309 • FZD7WNT1 2385 83 50 452 • AXIN1FZD5 2388 1177 327 1418 • FZD6WNT2 2396 1368 565 1119 • FZD7WNT2B 2402 874 21 1740 • FZD8PORCN 2405 143 394 2439 • AESFZD5 2410 2183 199 21 • FZD7PYGO1 2412 213 278 1057 • APCFZD5 2414 1853 1429 1032 • FZD2KREMEN1 2416 1248 997 1944 • FZD7WIF1 2421 533 93 578 • FZD1RHOU 2438 464 1890 1891 • FZD1PORCN 2439 153 1710 497 • FZD2WNT3 2443 2319 54 1361 • FZD7T 2446 288 32 717 • FZD7RHOU 2449 569 657 122 • FZD1PYGO1 2450 181 2133 1563 • FZD6PYGO1 2452 1028 269 591 • FZD1WNT3 2453 962 193 444 • FZD2PORCN 2464 1204 276 1676 • FZD6WNT3A 2465 1127 2473 1208 • FZD1WNT2B 2471 694 516 191 • FZD2PYGO1 2476 1027 608 1467 • FZD7PORCN 2477 145 133 2132

## SFRP family 2*^nd^* order interaction ranking at different time points

• SENP2SFRP1 2 844 564 1691 202 • FRZBSFRP1 4 1416 1440 1205 217 • SENP2SFRP4 19 2375 620 1713 2313 • DKK1SFRP1 35 2383 2293 1403 427 • APCSFRP1 39 2431 1945 2009 455 • LRP5SFRP1 70 1751 2472 2201 601 • GSK3BSFRP1 71 141 1336 969 1276 • DIXDC1SFRP1 100 1886 468 2322 622 • RHOUSFRP1 108 1587 1788 698 527 • FRZBSFRP4 121 775 1514 1484 524 • FZD1SFRP1 152 1100 1368 1476 136 • CXXC4SFRP1 166 187 1411 236 2207 • SFRP4WNT1 185 2470 52 646 36 • FRAT1SFRP1 197 2157 394 1335 505 • FBXW11SFRP1 198 24 2408 2108 1265 • FZD7SFRP1 220 443 1425 2132 323 • FOSL1SFRP1 230 261 1567 1911 223 • NLKSFRP1 238 1695 2056 680 1011 • PPP2CASFRP1 244 124 2088 397 434 • DKK1SFRP4 254 111 1607 121 737 • CSNK2A1SFRP1 263 329 2229 1960 46 • EP300SFRP1 266 1639 2453 2169 254 • PITX2SFRP1 276 2019 2086 61 1942 • FZD5SFRP1 286 1217 1244 1784 166 • KREMEN1SFRP1 339 1791 2291 1948 878 • CSNK1DSFRP1 348 2025 1956 990 568 • DAAM1SFRP1 354 285 1441 2433 1728 • PPP2R1ASFRP1 359 545 1723 1265 2124 • FZD6SFRP1 377 142 1983 1734 809 • CCND3SFRP1 383 135 1562 418 304 • SFRP4WIF1 386 2480 968 1186 1410 • FBXW2SFRP1 408 1144 738 1637 1436 • FSHBSFRP1 426 1241 1620 304 161 • CTBP1SFRP1 437 529 1309 2368 210 • CSNK1G1SFRP1 444 1208 2459 1950 2354 • FZD8SFRP1 465 817 1725 1806 463 • FOSL1SFRP4 481 1096 1226 1645 2115 • FZD1SFRP4 504 1714 1619 2234 237 • CTNNB1SFRP1 527 1687 1208 869 811 • SFRP4WNT3 533 2437 792 1779 51 • NKD1SFRP1 535 962 2184 1355 314 • DVL1SFRP1 540 54 1646 551 616 • CTBP2SFRP1 547 1725 2169 219 370 • SFRP4T 565 2458 390 848 548 • RHOUSFRP4 579 855 1825 1380 1539 • APCSFRP4 580 647 2303 474 489 • LRP6SFRP1 618 1431 2292 2236 1664 • DIXDC1SFRP4 636 2189 1156 2103 2240 • CTNNBIP1SFRP1 668 1836 2084 1830 9 • FZD2SFRP1 679 761 1570 1972 243 • AESSFRP1 682 39 1841 1685 670 • FBXW11SFRP4 690 1997 1896 1609 1818 • CSNK1DSFRP4 713 1697 2107 441 2374 • FZD7SFRP4 727 2124 1298 1969 799 • LRP5SFRP4 750 250 1791 2197 1036 • AXIN1SFRP1 751 1338 1727 780 437 • CSNK2A1SFRP4 760 2335 2102 1722 1199 • DVL2SFRP1 793 405 963 329 535 • FZD5SFRP4 795 918 2105 2299 2276 • FRAT1SFRP4 806 1275 2203 217 2007 • GSK3BSFRP4 841 1659 2082 812 2350 • EP300SFRP4 898 2274 2449 2439 1821 • PPP2R1ASFRP4 903 1887 2325 1461 1681 • CCND3SFRP4 913 2364 2027 183 1422 • NLKSFRP4 916 2109 1948 1498 1764 • BTRCSFRP1 940 527 2178 627 1234 • PORCNSFRP1 977 1946 2200 724 2255 • MYCSFRP1 983 337 2374 2270 8 • CCND1SFRP1 1007 273 975 1536 2145 • FZD8SFRP4 1009 1935 1451 1740 2297 • PYGO1SFRP1 1029 1279 2092 1230 572 • FBXW2SFRP4 1032 1388 1682 580 1973 • CCND2SFRP1 1050 2090 1329 1294 820 • FGF4SFRP1 1061 1272 721 629 909 • CXXC4SFRP4 1066 825 2382 2362 1745 • PPP2CASFRP4 1077 2092 1844 718 1109 • CTBP1SFRP4 1138 238 1431 443 1725 • PITX2SFRP4 1140 1072 2373 78 1500 • LEF1SFRP1 1142 2459 2269 2006 813 • BCL9SFRP1 1146 1071 2271 1073 1766 • SFRP4FBXW4 1164 2456 121 2465 1126 • KREMEN1SFRP4 1179 956 1913 180 2039 • SFRP4TLE1 1200 2485 918 674 82 • SFRP1WNT1 1235 1861 180 1151 1734 • FSHBSFRP4 1239 2168 2096 164 1260 • DAAM1SFRP4 1266 2240 1111 1404 1022 • FZD6SFRP4 1303 1485 1286 461 781 • CSNK1A1SFRP1 1321 236 1421 1504 88 • SFRP4SLC9A3R1 1349 2447 991 1502 1040 • SFRP1WIF1 1358 1877 1241 2388 2158 • SFRP4WNT2 1387 1947 1288 814 363 • CTNNB1SFRP4 1395 1332 2424 1045 1933 • SFRP4WNT5A 1467 2474 112 865 575 • SFRP4TCF7 1471 2462 68 1217 648 • DVL2SFRP4 1483 1464 1656 1251 1188 • SFRP4WNT4 1487 1371 1088 1156 1504 • JUNSFRP1 1492 548 1921 404 24 • FOXN1SFRP1 1513 424 1702 791 65 • NKD1SFRP4 1574 2420 1652 638 2238 • CTBP2SFRP4 1585 2481 1446 1218 1679 • SFRP4WNT3A 1602 2242 937 936 1813 • SFRP1WNT3 1613 4 974 2122 2481 • CTNNBIP1SFRP4 1636 1302 2377 64 150 • GSK3ASFRP1 1639 1655 1864 1546 686 • CSNK1G1SFRP4 1646 1536 2182 90 531 • SFRP4TLE2 1696 1917 1338 1026 344 • SFRP4WNT2B 1704 2366 85 108 2173 • AXIN1SFRP4 1729 1199 2471 190 954 • LRP6SFRP4 1735 1149 2141 1515 1684 • SFRP1FBXW4 1765 1442 327 2117 2300 • CCND2SFRP4 1771 1215 1994 1548 1509 • SFRP4TCF7L1 1839 2482 1253 2414 1701 • CCND1SFRP4 1859 1897 2376 1787 1272 • SFRP1T 1879 1964 248 9 2027 • PYGO1SFRP4 1903 1513 1133 314 1975 • DVL1SFRP4 1905 2085 2274 2119 1834 • PORCNSFRP4 1916 1883 2165 19 1661 • BTRCSFRP4 1918 2210 1487 996 928 • MYCSFRP4 1966 1960 1676 122 1206 • LEF1SFRP4 1973 488 2093 62 2081 • SFRP1SFRP4 1987 1462 1469 1747 2270 • BCL9SFRP4 2048 951 2252 950 1112 • SFRP1TLE1 2050 1972 1708 524 1322 • SFRP1WNT2 2055 300 855 555 1871 • AESSFRP4 2081 1869 2339 842 1527 • CSNK1A1SFRP4 2111 1733 1076 1030 240 • SFRP1WNT5A 2113 2217 482 1725 2484 • FZD2SFRP4 2130 957 1916 159 362• JUNSFRP4 2135 1701 2070 26 4 • SFRP1TCF7 2152 239 57 2228 1797 • SFRP1WNT3A 2160 837 338 1425 705 • SFRP1SLC9A3R1 2178 100 1037 2392 2244 • FGF4SFRP4 2191 665 1105 1582 1074 • FOXN1SFRP4 2202 2211 1833 495 694 • SFRP1WNT4 2219 1238 669 2173 1848 • SFRP1WNT2B 2229 36 129 1356 1589 • SFRP1TLE2 2295 1370 835 1469 1900 • GSK3ASFRP4 2311 396 2118 547 1146 • SFRP1TCF7L1 2324 1770 505 2473 2174

## SFRP family 2*^nd^* order interaction ranking at different durations

• FZD5SFRP1 2 1461 1499 1726 • FZD5SFRP4 12 1326 2012 926 • CSNK1A1SFRP4 22 1465 2478 64 • RHOUSFRP4 27 1514 113 56 • CSNK1A1SFRP1 29 2068 1414 412 • PYGO1SFRP4 30 1398 569 24 • CTBP1SFRP4 39 1623 2330 324 • CTNNB1SFRP4 42 1294 329 233 • JUNSFRP4 54 2392 1043 5 • AESSFRP4 58 2200 1366 35 • DIXDC1SFRP4 60 677 196 1682 • PYGO1SFRP1 63 2345 2393 464 • FRAT1SFRP4 73 1120 1996 197 • KREMEN1SFRP4 84 1106 376 238 • FOXN1SFRP4 91 1544 520 102 • CCND2SFRP4 92 1257 2205 1634 • BCL9SFRP4 103 1549 703 850 • AXIN1SFRP4 106 2009 596 794 • FSHBSFRP4 107 829 1792 1363 • JUNSFRP1 113 1873 1613 83 • CTNNB1SFRP1 116 1555 2034 1448 • MYCSFRP4 117 1132 2120 568 • CTNNBIP1SFRP4 118 2202 465 1096 • CSNK1DSFRP4 129 2425 1161 106 • RHOUSFRP1 134 2054 770 57 • DVL1SFRP4 139 2096 336 815 • CCND1SFRP4 143 2081 310 431 • BTRCSFRP4 154 2256 1252 129 • APCSFRP4 166 1128 2091 549 • GSK3ASFRP4 185 2456 305 1669 • CTBP1SFRP1 204 1504 956 918 • FOSL1SFRP4 207 1214 617 145 • FRAT1SFRP1 217 1957 2451 402 • PORCNSFRP4 236 1924 470 329 • FOXN1SFRP1 247 2189 2311 413 • DIXDC1SFRP1 277 785 947 2427 • CTBP2SFRP4 279 2071 1243 724 • CCND2SFRP1 280 728 2251 2291 • PORCNSFRP1 296 1810 1702 1045 • NKD1SFRP4 310 1209 1781 623 • AXIN1SFRP1 315 2423 2110 2194 • AESSFRP1 329 1881 2274 82 • BCL9SFRP1 334 1270 2340 785 • EP300SFRP4 340 1667 396 1466 • LRP5SFRP4 353 1295 651 32 • CTNNBIP1SFRP1 388 2213 912 1768 • DVL2SFRP4 389 2219 342 797 • CSNK1DSFRP1 393 1904 1904 566 • KREMEN1SFRP1 423 2308 854 1492 • MYCSFRP1 424 1986 1798 2190 • BTRCSFRP1 446 2140 1327 929 • FRZBSFRP4 454 2249 2397 12 • CCND1SFRP1 462 1755 671 1480 • CXXC4SFRP4 468 1377 2195 1270 • FSHBSFRP1 507 1553 1795 860 • DKK1SFRP4 528 1109 1951 207 • DVL1SFRP1 534 1486 1549 2079 • PPP2CASFRP4 549 2262 1511 112 • CTBP2SFRP1 552 2314 2448 1880 • DVL2SFRP1 585 1866 926 1739 • GSK3ASFRP1 596 1987 536 2228 • FZD6SFRP4 629 975 175 866 • CCND3SFRP4 640 1797 346 142 • FOSL1SFRP1 660 1092 1934 1005 • EP300SFRP1 669 1217 817 2045 • DAAM1SFRP4 680 2107 427 1760 • APCSFRP1 688 1189 1002 631 • PITX2SFRP4 692 327 238 172 • FZD8SFRP4 733 724 57 382 • GSK3BSFRP4 749 1476 124 19 • NKD1SFRP1 752 1200 1546 1007 • LRP5SFRP1 759 792 2433 4 • NLKSFRP4 770 1452 1123 629 • FRZBSFRP1 791 1894 1083 53 • FZD1SFRP4 855 540 224 69 • FBXW11SFRP4 863 2160 1731 110 • FBXW2SFRP4 879 2244 1420 919 • CCND3SFRP1 883 1870 1830 820 • FZD2SFRP4 906 1061 155 433 • CSNK2A1SFRP4 920 1753 89 336 • SFRP1SFRP4 927 594 718 481 • FGF4SFRP4 950 935 755 1309 • FZD7SFRP4 977 1976 13 2198 • CXXC4SFRP1 983 1521 1800 1742 • DKK1SFRP1 996 1367 1679 617 • DAAM1SFRP1 1021 2277 923 2371 • PPP2R1ASFRP4 1036 2026 1395 840 • CSNK1G1SFRP4 1106 1420 1176 482 • FZD8SFRP1 1120 1668 932 915 • NLKSFRP1 1140 2087 1009 935 • CSNK2A1SFRP1 1192 1386 978 1074 • PPP2CASFRP1 1193 2137 2035 610 • LEF1SFRP4 1194 392 672 536 • FZD6SFRP1 1200 1454 464 1946 • PITX2SFRP1 1243 438 887 779 • LRP6SFRP4 1248 699 1730 475 • FBXW2SFRP1 1260 2005 1228 1481 • FBXW11SFRP1 1274 2040 2020 1623 • FZD7SFRP1 1311 1796 136 2362 • SFRP1WNT5A 1314 166 1969 2159 • SFRP1TCF7L1 1371 690 722 1330 • PPP2R1ASFRP1 1420 1757 2063 2189 • GSK3BSFRP1 1475 719 555 29 • FZD1SFRP1 1508 1682 2215 410 • SFRP1TLE2 1544 133 1995 210 • LEF1SFRP1 1560 1015 1939 964 • FGF4SFRP1 1586 1507 637 2314 • LRP6SFRP1 1647 2049 2209 1483 • CSNK1G1SFRP1 1663 2391 2305 2057 • SENP2SFRP4 1707 1833 1070 1264 • FZD2SFRP1 1785 1479 761 696 • SFRP1FBXW4 1840 39 1050 1794 • SFRP4TCF7L1 1912 456 408 187 • SFRP1WNT2 1919 283 917 1518 • SFRP1WNT4 1922 151 1496 986 • SFRP1TCF7 1943 236 272 829 • SFRP1SLC9A3R1 2009 391 1990 977 • SFRP4WNT5A 2054 123 504 588 • SENP2SFRP1 2056 2472 1883 808 • SFRP4FBXW4 2068 119 952 695 • SFRP1WNT3 2088 1627 631 1256 • SFRP1TLE1 2105 742 1998 1081 • SFRP1WNT3A 2110 436 1867 2123 • SFRP4WNT3A 2127 506 1029 448 • SFRP4TLE2 2166 341 1984 182 • SFRP1WNT1 2193 142 1681 1072 • SFRP1WNT2B 2201 670 593 799 • SFRP4TCF7 2240 157 460 360 • SFRP4TLE1 2247 488 1838 645 • SFRP4WNT3 2285 2333 888 644 • SFRP4T 2340 203 2380 139 • SFRP1WIF1 2346 701 2184 2375 • SFRP4WNT1 2349 136 2142 567 • SFRP4WNT4 2353 250 2131 702 • SFRP1T 2376 100 794 325 • SFRP4WIF1 2406 635 2164 1775 • SFRP4WNT2B 2431 584 949 2071 • SFRP4WNT2 2478 351 1692 619 • SFRP4SLC9A3R1 2482 498 1266 1106

## GSK family 2*^nd^* order interaction ranking at different time points

• FRZBGSK3A 5 1482 90 1544 403 • DKK1GSK3A 6 2313 689 1907 1362 • FZD7GSK3A 20 579 23 1892 1742 • APCGSK3A 30 1591 124 1708 913 • FZD8GSK3A 42 772 122 323 1605 • FBXW11GSK3A 44 334 938 111 2301 • FZD1GSK3A 47 1007 212 758 315 • CSNK2A1GSK3A 53 991 746 2186 1207 • GSK3BSFRP1 71 141 1336 969 1276 • FZD5GSK3A 72 311 163 109 1477 • FOSL1GSK3A 73 1202 62 1115 461 • CXXC4GSK3A 74 147 114 415 2259 • FZD6GSK3A 75 452 2230 1009 1378 • GSK3BPYGO1 91 972 292 2229 539 • CCND3GSK3A 105 1200 670 115 2399 • FRZBGSK3B 107 2165 470 1773 1518 • CSNK1DGSK3A 119 933 261 1319 680 • DAAM1GSK3A 132 1331 534 2471 1771 • GSK3BJUN 133 1510 851 1724 1168 • DIXDC1GSK3A 151 260 4 2044 2448 • FRAT1GSK3A 161 1129 194 2043 1807 • GSK3BWNT1 172 1939 54 894 1897 • GSK3BMYC 190 752 429 1812 2427 • GSK3BWIF1 211 1840 1988 260 1125 • EP300GSK3A 218 1350 521 1506 794 • CTNNBIP1GSK3A 225 766 35 856 376 • FBXW2GSK3A 236 1675 375 1472 2146 • CTBP1GSK3A 253 1551 49 305 861 • DKK1GSK3B 279 2389 1937 2198 2201 • GSK3BLEF1 293 52 1003 2222 1566 • FSHBGSK3A 303 307 1173 487 641 • CTBP2GSK3A 309 1143 325 1219 1866 • DVL2GSK3A 353 1898 1103 74 987 • GSK3BWNT3 378 1722 837 1523 1019 • GSK3BPORCN 412 1010 314 1646 327 • GSK3BLRP6 417 1620 533 1789 75 • CTNNB1GSK3A 447 1713 304 1640 186 • AXIN1GSK3A 458 1443 527 255 342 • CSNK1G1GSK3A 462 113 1893 300 2462 • GSK3BT 473 1531 762 602 2452 • CCND2GSK3A 498 662 676 1762 1632 • AESGSK3A 501 178 692 1730 595 • DVL1GSK3A 529 2072 131 1370 2367 • BTRCGSK3A 538 827 1786 1889 2366 • FGF4GSK3A 576 1670 28 1651 1038 • GSK3BNKD1 578 753 463 226 1494 • FBXW11GSK3B 660 1365 1025 2246 1093 • APCGSK3B 700 2255 1553 1445 525 • CSNK1A1GSK3A 742 1261 483 2020 377 • GSK3BTLE1 763 1812 2324 909 1453 • FZD2GSK3A 767 1457 137 352 1358 • BCL9GSK3A 790 1599 412 1164 1409 • GSK3BPITX2 809 84 1650 1140 845 • GSK3BSLC9A3R1 818 362 1641 166 2356 • FZD7GSK3B 834 1117 1510 1985 976 • GSK3BSFRP4 841 1659 2082 812 2350 • GSK3BFBXW4 858 1983 854 238 2279 • CSNK2A1GSK3B 875 1115 1899 1701 1128 • GSK3BTCF7 878 1131 66 306 1964 • FOXN1GSK3A 880 1274 2450 272 994 • FZD1GSK3B 920 762 815 358 1875 • GSK3BWNT4 936 1475 995 686 1480 • GSK3BNLK 974 297 637 1190 2042 • FOSL1GSK3B 976 2305 872 313 123 • GSK3BKREMEN1 998 161 2138 489 330 • CCND1GSK3A 1041 184 312 957 1447 • FRAT1GSK3B 1058 2294 1754 880 859 • GSK3BWNT2 1070 879 1720 401 2431 • FZD5GSK3B 1086 1447 1086 2282 1985 • EP300GSK3B 1107 276 1935 622 1949 • GSK3BWNT3A 1118 1234 2188 1036 1495 • GSK3APYGO1 1121 411 1993 1052 1717 • DIXDC1GSK3B 1127 1352 252 11 2478 • GSK3BPPP2CA 1130 2453 152 819 2206 • CSNK1DGSK3B 1160 2452 2448 2268 1830 • FZD8GSK3B 1194 1892 795 422 2109 • GSK3BPPP2R1A 1205 313 728 1562 596 • FZD6GSK3B 1219 778 1762 383 2371 • GSK3BTLE2 1283 1525 1976 146 1277 • GSK3BLRP5 1288 496 636 558 2128 • GSK3BWNT2B 1293 1228 195 1142 1496 • FBXW2GSK3B 1314 459 2279 1395 804 • CCND3GSK3B 1318 687 2317 2284 1327 • CTBP1GSK3B 1350 2212 2128 229 1039 • CXXC4GSK3B 1394 2112 1849 1278 2477 • GSK3BRHOU 1441 1854 551 1689 1121 • GSK3BWNT5A 1449 1554 386 1077 1882 • GSK3AMYC 1464 2169 1538 745 2288 • FSHBGSK3B 1516 2167 2254 2206 498 • GSK3AJUN 1554 233 2047 609 869 • GSK3ALEF1 1567 784 1281 637 2242 • DAAM1GSK3B 1575 846 1151 332 2252 • GSK3AWNT3 1604 577 1759 1992 1888 • GSK3ASFRP1 1639 1655 1864 1546 686 • GSK3AWIF1 1693 1515 2204 2342 1317 • GSK3BTCF7L1 1710 1105 699 1763 2191 • CSNK1G1GSK3B 1712 2466 1919 1793 2138 • CTNNB1GSK3B 1826 1785 697 2126 1553 • GSK3APORCN 1854 278 2481 1352 1575 • DVL2GSK3B 1855 1533 1172 1986 1289 • GSK3AWNT1 1867 2239 1343 1167 1455 • GSK3ALRP6 1901 653 817 1279 1852 • CTNNBIP1GSK3B 1913 2300 1689 1339 343 • CTBP2GSK3B 2017 81 1472 1886 1677 • BTRCGSK3B 2018 1359 1685 2167 2344 • CCND2GSK3B 2022 1922 2091 257 1962 • AXIN1GSK3B 2059 1490 1194 161 864 • BCL9GSK3B 2060 1306 2119 264 2409 • GSK3AWNT4 2168 1760 1694 2376 741 • GSK3AFBXW4 2184 2369 1679 2432 198 • GSK3ANKD1 2196 2045 766 1834 1874 • DVL1GSK3B 2209 123 1378 390 2445 • GSK3APITX2 2212 1865 1735 742 1466 • GSK3AT 2223 254 2114 2377 2411 • FGF4GSK3B 2233 1622 1058 1255 2247 • FZD2GSK3B 2273 1534 1079 2131 953 • CCND1GSK3B 2274 426 1308 142 162 • AESGSK3B 2300 1473 2469 1728 1878 • CSNK1A1GSK3B 2302 2005 1321 575 816 • GSK3AWNT3A 2309 960 1709 1688 1633 • GSK3ASFRP4 2311 396 2118 547 1146 • GSK3APPP2CA 2328 2050 798 1503 2012 • GSK3AWNT2 2331 1649 2282 2149 2130 • FOXN1GSK3B 2334 731 1382 2059 2219 • GSK3ATLE1 2339 1999 1386 1134 438 • GSK3ATCF7 2365 2110 1061 1796 1180 • GSK3ANLK 2372 1405 1371 765 1071 • GSK3AWNT5A 2376 497 1129 1893 1295 • GSK3AKREMEN1 2399 1248 1868 1771 1025 • GSK3APPP2R1A 2410 1478 1731 1326 850 • GSK3AWNT2B 2419 30 579 157 1380 • GSK3BSENP2 2422 1441 1885 42 991 • GSK3AGSK3B 2423 1949 2342 559 1966 • GSK3ATCF7L1 2426 2078 2284 1423 1585 • GSK3ATLE2 2433 807 1873 2022 702 • GSK3ALRP5 2437 881 2025 2172 993 • GSK3ASLC9A3R1 2438 814 2207 1792 2257 • GSK3ARHOU 2449 2162 2332 1308 2417 • GSK3ASENP2 2481 1798 1323 250 2261

## GSK family 2*^nd^* order interaction ranking at different durations

• FZD5GSK3B 45 2414 1261 2038 • GSK3ASENP2 99 968 15 1795 • CSNK1A1GSK3B 147 1715 991 1042 • GSK3ASFRP4 185 2456 305 1669 • CSNK1A1GSK3A 193 2094 680 308 • DIXDC1GSK3B 213 953 1928 113 • AESGSK3B 216 2187 924 892 • CTNNB1GSK3B 223 1278 1084 1491 • CTBP1GSK3B 301 1140 845 627 • CCND2GSK3B 380 940 936 687 • AXIN1GSK3B 390 2351 2353 34 • FZD5GSK3A 425 1872 1099 2300 • GSK3ALRP6 438 1251 109 1690 • CTNNBIP1GSK3B 444 1820 1196 146 • FRAT1GSK3B 453 1714 1154 2336 • GSK3BSENP2 457 591 105 58 • CSNK1DGSK3B 461 1046 1145 1236 • FSHBGSK3B 494 2432 1630 272 • GSK3ALEF1 496 2065 598 1728 • GSK3AGSK3B 498 1661 1485 114 • CCND1GSK3B 515 1842 1961 907 • GSK3APPP2R1A 519 439 367 950 • BTRCGSK3B 532 1819 1185 1233 • APCGSK3B 555 1188 386 1681 • BCL9GSK3B 558 2323 1604 795 • DVL1GSK3B 589 1158 2048 107 • GSK3ASFRP1 596 1987 536 2228 • GSK3ATCF7L1 611 1629 304 1867 • FOSL1GSK3B 616 1480 995 947 • FOXN1GSK3B 631 1917 1346 2226 • GSK3AWNT5A 639 286 389 1426 • CTBP2GSK3B 673 2082 1321 175 • GSK3APITX2 720 2295 2155 868 • CTNNB1GSK3A 747 1408 1689 267 • GSK3BSFRP4 749 1476 124 19 • EP300GSK3B 781 1586 1952 393 • GSK3ANLK 783 237 19 1929 • GSK3ATLE2 807 428 599 408 • GSK3APPP2CA 813 338 235 772 • CTBP1GSK3A 858 1560 381 476 • GSK3AFBXW4 899 164 1170 2074 • GSK3BLRP6 938 750 67 224 • GSK3ATLE1 940 1135 1107 1028 • FRZBGSK3B 949 1834 663 601 • DKK1GSK3B 971 1405 685 411 • DVL2GSK3B 981 2135 494 1079 • AESGSK3A 1000 2310 1753 290 • DIXDC1GSK3A 1012 440 2018 1279 • GSK3ATCF7 1068 427 2 1339 • GSK3BLEF1 1077 1203 676 604 • FSHBGSK3A 1084 1493 1880 1832 • AXIN1GSK3A 1085 2434 1529 1935 • CXXC4GSK3B 1096 1542 750 239 • GSK3ALRP5 1100 2100 719 195 • FRAT1GSK3A 1108 1566 2178 440 • CCND2GSK3A 1113 2359 1066 1712 • CCND1GSK3A 1126 1122 2370 652 • GSK3BTCF7L1 1137 603 444 1542 • GSK3AWNT4 1156 175 1638 738 • CCND3GSK3B 1162 933 1903 1127 • DAAM1GSK3B 1180 1564 1293 158 • GSK3AWNT1 1216 265 748 846 • FGF4GSK3B 1219 1066 2096 383 • GSK3ASLC9A3R1 1224 650 1184 895 • BCL9GSK3A 1241 2399 1908 778 • FZD6GSK3B 1258 1116 219 524 • GSK3ANKD1 1259 76 68 1377 • CSNK1DGSK3A 1263 1943 1524 202 • GSK3AWNT2 1275 599 2042 1294 • GSK3BPPP2R1A 1282 98 207 171 • FOXN1GSK3A 1290 1079 1909 149 • FBXW11GSK3B 1291 1302 973 877 • BTRCGSK3A 1326 2447 1621 487 • GSK3AWIF1 1328 831 457 2307 • GSK3AJUN 1340 305 431 467 • CTNNBIP1GSK3A 1346 2143 597 2138 • GSK3BWNT5A 1347 137 577 443 • FZD8GSK3B 1364 669 1088 1409 • CSNK2A1GSK3B 1391 2110 204 2258 • FBXW2GSK3B 1405 1923 1075 670 • FZD1GSK3B 1407 1411 909 1947 • FZD7GSK3B 1441 1830 896 276 • CTBP2GSK3A 1456 1718 1611 1020 • GSK3BSFRP1 1475 719 555 29 • GSK3BPITX2 1495 1525 559 669 • GSK3BNLK 1506 105 30 52 • FOSL1GSK3A 1533 1137 1522 1087 • GSK3BTLE2 1548 179 545 1574 • DKK1GSK3A 1581 888 987 231 • GSK3BPPP2CA 1583 87 314 2018 • GSK3AT 1603 578 468 293 • FRZBGSK3A 1629 2038 1392 397 • FZD2GSK3B 1631 2365 1927 927 • APCGSK3A 1639 1670 885 1587 • DVL1GSK3A 1662 1851 2480 366 • GSK3AKREMEN1 1665 478 1159 2034 • FBXW2GSK3A 1691 1749 1062 415 • GSK3AMYC 1692 330 22 831 • GSK3BTCF7 1723 69 1 870 • GSK3BWNT4 1754 169 1073 41 • GSK3BWNT1 1791 89 177 1029 • GSK3BFBXW4 1800 20 776 261 • GSK3BNKD1 1818 10 131 765 • CSNK1G1GSK3B 1831 1162 1272 504 • DAAM1GSK3A 1834 1477 2225 2425 • DVL2GSK3A 1839 2227 2307 1621 • EP300GSK3A 1845 990 1373 495 • GSK3BTLE1 1850 535 894 552 • GSK3BLRP5 1885 804 186 2352 • GSK3BSLC9A3R1 1903 317 552 93 • GSK3AWNT3A 1911 1577 2383 1721 • FZD7GSK3A 1917 1254 286 1385 • FZD1GSK3A 1920 711 1256 510 • FZD6GSK3A 1921 1362 686 1207 • GSK3AWNT2B 1972 2158 88 1351 • FBXW11GSK3A 1974 1299 820 720 • GSK3BWIF1 1992 526 426 77 • FZD8GSK3A 2017 556 1104 1105 • GSK3BT 2080 111 191 573 • GSK3ARHOU 2085 867 1814 710 • CXXC4GSK3A 2100 1621 529 979 • GSK3BRHOU 2126 356 570 673 • GSK3BKREMEN1 2135 235 1116 2381 • GSK3BWNT2 2137 210 1106 91 • FGF4GSK3A 2145 1055 677 982 • GSK3BJUN 2155 135 80 2298 • FZD2GSK3A 2210 1244 1350 1156 • CCND3GSK3A 2212 2016 2125 715 • CSNK2A1GSK3A 2233 1998 951 1370 • GSK3BWNT2B 2244 415 165 254 • GSK3BWNT3 2263 995 70 194 • GSK3BWNT3A 2314 522 1207 659 • CSNK1G1GSK3A 2330 852 1284 174 • GSK3BPYGO1 2347 148 455 2250 • GSK3BMYC 2382 70 53 403 • GSK3AWNT3 2383 1228 38 2370 • GSK3BPORCN 2407 97 439 727 • GSK3APORCN 2433 354 628 470 • GSK3APYGO1 2463 486 640 686

## CTBP family 2*^nd^* order interaction ranking at different time points

• APCCTBP2 217 2410 2039 826 1884 • CTBP1WNT1 245 1792 34 32 159 • CTBP1GSK3A 253 1551 49 305 861 • CTBP2JUN 285 567 1396 425 1806 • CTBP2GSK3A 309 1143 325 1219 1866 • CTBP1LEF1 346 1389 725 273 464 • CTBP1FOXN1 350 2271 149 322 1550 • CTBP1FGF4 392 931 1118 1816 248 • CTBP2MYC 401 352 433 1765 2024 • CTBP1JUN 405 2160 878 1365 1747 • FZD5CTBP2 409 1583 1435 593 1263 • CTBP1MYC 411 2011 580 683 1227 • CSNK1DCTBP2 416 2400 1630 1236 1660 • CTBP1WIF1 436 2350 1924 342 810 • CTBP1SFRP1 437 529 1309 2368 210 • CTBP1FZD2 442 1207 802 1754 1411 • CTBP2FZD2 456 645 1801 899 1689 • CSNK2A1CTBP2 464 2130 2347 1916 1914 • CTBP2FOXN1 486 469 329 1586 868 • CTBP1PORCN 492 1654 421 1369 62 • CTBP1WNT3 503 2015 1703 2207 14 • CTBP1DVL1 505 2071 1305 1593 221 • CTBP1PYGO1 507 860 255 403 114 • CTBP2WNT1 510 783 128 628 621 • APCCTBP1 519 1607 2015 2093 2342 • CTBP2SFRP1 547 1725 2169 219 370 • CTBP1LRP6 577 1035 693 1874 487 • CTBP2PYGO1 600 1633 626 1657 1210 • CCND3CTBP2 652 1909 1180 1441 812 • CTBP2FGF4 661 1497 2473 1109 502 • CTBP1T 667 1524 944 983 2233 • CTBP2WNT3 686 1061 1455 2025 95 • CTBP1DVL2 697 2343 537 847 1393 • CTBP2LEF1 739 447 388 1183 2049 • CTBP2DVL1 743 903 1334 1411 1095 • CSNK2A1CTBP1 768 221 1236 2213 1885 • CSNK1DCTBP1 775 417 2320 923 1107 • FZD5CTBP1 799 604 867 2225 2135 • CTBP1CTBP2 814 1616 1572 2405 2071 • CTBP2WIF1 830 1880 1101 705 2053 • CTBP1DAAM1 851 1204 393 972 64 • CTBP1TLE1 852 1966 1640 1284 691 • CTBP1FSHB 857 1027 231 840 2418 • CTBP1CTNNBIP1 859 967 1536 2217 1607 • CTBP2PORCN 862 553 827 1008 147 • CTBP1CTNNB1 909 1082 807 2164 764 • CTBP1NKD1 938 2166 487 1839 128 • CTBP1FBXW4 945 2064 203 1615 626 • CTBP2LRP6 978 296 967 1199 510 • CSNK1G1CTBP2 1096 2302 1432 1979 2054 • CTBP2DVL2 1102 505 1438 1756 1779 • CTBP1PITX2 1113 1194 1741 220 417 • CTBP2CTNNBIP1 1125 1358 1830 1795 1877 • CTBP1SFRP4 1138 238 1431 443 1725 • CTBP2T 1139 2293 192 2125 634 • AESCTBP2 1167 177 985 1059 899 • CCND3CTBP1 1176 981 2210 764 1497 • CTBP1FZD6 1204 1904 65 514 291 • CTBP1KREMEN1 1221 1835 1127 1687 760 • CTBP1NLK 1227 2323 100 1055 499 • CTBP2CTNNB1 1254 190 2411 2375 2025 • CTBP1WNT2 1259 574 1388 1619 1324 • CTBP1WNT5A 1297 2043 716 1282 71 • CTBP1DIXDC1 1301 1866 1166 2467 191 • AXIN1CTBP2 1306 1987 1043 1989 1636 • CTBP1EP300 1323 1830 1669 2089 2228 • CTBP1GSK3B 1350 2212 2128 229 1039 • CTBP2NKD1 1359 2213 246 1547 1342 • CTBP1SLC9A3R1 1416 1779 2465 1802 384 • CTBP1FBXW2 1418 1640 780 1833 156 • CTBP1PPP2CA 1440 1663 58 1033 1002 • CTBP2FBXW4 1447 234 229 2422 1542 • CTBP1FZD8 1461 1638 1020 717 63 • CTBP1CXXC4 1477 862 2033 1554 101 • CTBP2FSHB 1479 780 509 244 1248 • CTBP1WNT4 1519 782 1140 1663 1770 • CTBP2TLE1 1522 614 1925 2028 1016 • CTBP1PPP2R1A 1523 1092 743 1727 77 • CTBP1LRP5 1531 446 449 463 1226 • CTBP1WNT2B 1535 929 22 1459 1319 • CTBP1TCF7 1552 2479 368 2112 802 • CTBP2DAAM1 1556 456 136 28 530 • CCND1CTBP2 1580 983 775 2218 713 • BTRCCTBP2 1584 834 2062 800 1610 • CTBP2SFRP4 1585 2481 1446 1218 1679 • CTBP1TLE2 1605 1851 1089 2208 1960 • AXIN1CTBP1 1610 934 1728 712 1414 • CSNK1G1CTBP1 1622 1952 1490 1266 1765 • CCND2CTBP2 1629 1344 2162 1885 1831 • CTBP2PPP2CA 1638 864 153 1565 950 • CSNK1A1CTBP1 1645 1243 1897 2365 1533 • CTBP2FBXW2 1668 1173 772 2115 731 • CTBP1FZD1 1690 1864 881 584 511 • CTBP2FZD6 1721 431 316 2429 1772 • CTBP1FOSL1 1728 657 2244 2331 2269 • BCL9CTBP2 1737 1797 2123 2120 1159 • CTBP1FRAT1 1742 1825 943 952 1003 • CSNK1A1CTBP2 1743 720 1722 2230 519 • CTBP2FZD8 1744 1242 1557 1819 1148 • CTBP1FBXW11 1748 1868 340 177 292 • CTBP2NLK 1751 387 518 608 345 • CTBP2EP300 1754 916 1019 973 1203 • CTBP2WNT2 1757 641 1838 2418 1151 • CTBP2WNT3A 1763 925 2380 2194 602 • CTBP1WNT3A 1764 1567 2205 414 1009 • CTBP2TCF7 1782 863 191 2366 352 • CTBP2PPP2R1A 1789 1337 201 2200 688 • CTBP2KREMEN1 1794 1627 1046 2114 1582 • CTBP2PITX2 1799 884 612 215 890 • CTBP2SLC9A3R1 1804 419 753 2360 2305 • CTBP2RHOU 1810 1465 1107 984 793 • CTBP2CXXC4 1815 562 1942 809 625 • BTRCCTBP1 1833 1142 1498 331 1205 • AESCTBP1 1837 2190 2478 216 1514 • CCND2CTBP1 1840 1755 1657 1092 353 • CTBP2WNT4 1858 1373 939 1832 1978 • CTBP1RHOU 1889 2017 895 423 1357 • CTBP1TCF7L1 1891 1743 517 1031 1628 • CTBP1FZD7 1908 1323 2384 360 38 • CTBP2TLE2 1921 2220 2037 1638 2097 • CTBP2LRP5 1927 1375 1073 2237 1652 • CTBP2FZD7 1968 489 2410 2383 1460 • CTBP2WNT2B 2016 1908 185 1931 537 • CTBP2GSK3B 2017 81 1472 1886 1677 • CTBP2FBXW11 2026 740 92 2069 576 • CTBP2FOSL1 2033 2156 1904 2438 1233 • CTBP2FRAT1 2052 2377 2129 1145 2267 • CTBP2DIXDC1 2061 1052 1927 1085 338 • CTBP2WNT5A 2065 175 639 1291 1722 • BCL9CTBP1 2165 1690 1603 59 1829 • CTBP2FZD1 2255 237 1247 810 2066 • CTBP2TCF7L1 2278 2283 1810 1881 2472 • CTBP1DKK1 2335 1590 96 1298 207 • CTBP2DKK1 2351 1993 310 1261 966 • CCND1CTBP1 2361 1995 970 1366 1147 • CTBP1FRZB 2412 575 1772 1692 961 • CTBP2FRZB 2453 2226 1112 261 1838 • CTBP1SENP2 2465 1852 1205 1526 180 • CTBP2SENP2 2469 256 2301 447 419

## CTBP family 2*^nd^* order interaction ranking at different durations

• CTBP1SENP2 21 2029 2136 101 • CTBP1SFRP4 39 1623 2330 324 • FZD5CTBP2 55 950 2306 2466 • CTBP1FZD2 74 2285 654 1474 • CSNK1A1CTBP2 94 2403 2329 586 • CTBP2SENP2 108 1288 2071 1540 • CTBP1FZD7 119 1915 298 1013 • CTBP1TCF7L1 120 1967 1053 2317 • CTBP1FBXW2 125 2182 403 2046 • CTBP1LRP6 144 1761 1537 2048 • CTBP1LEF1 169 1659 1592 518 • CTBP1PPP2R1A 172 1593 1458 1512 • CTBP1FBXW11 176 2347 1757 1223 • CTBP1FZD8 180 1432 524 1741 • CTBP1PPP2CA 199 1626 1662 1216 • CTBP1SFRP1 204 1504 956 918 • CTBP1FZD1 211 1730 1533 2325 • CTBP1FGF4 220 1510 773 985 • CTBP1CXXC4 227 1252 2359 1404 • CTBP1PITX2 234 1334 591 1287 • CTBP1FZD6 248 1726 648 888 • CTBP2SFRP4 279 2071 1243 724 • CTBP1GSK3B 301 1140 845 627 • CTBP1TLE2 306 2008 1500 1135 • CTBP1WNT5A 336 2412 1868 2056 • CTBP1NLK 350 945 632 2454 • CTBP1FRZB 356 1950 1380 1736 • CTBP1DKK1 358 1389 1779 2471 • CTBP1DAAM1 381 2341 1018 396 • CTBP1LRP5 404 1696 1606 1169 • CTBP2LEF1 440 1906 2104 2082 • CTBP1EP300 442 776 1756 1324 • CTBP2LRP6 464 2076 1751 2354 • CTBP1CTBP2 470 1605 1587 2432 • CTBP2FZD7 537 1003 653 2162 • CTBP2FGF4 547 2330 2188 1756 • CTBP2SFRP1 552 2314 2448 1880 • CTBP1WNT4 559 1416 1094 2482 • CTBP2FZD2 565 2255 1910 1395 • CTBP2WNT5A 582 1794 2172 2472 • CTBP2TCF7L1 604 2378 2429 1184 • CTBP2FZD6 607 687 946 722 • CTBP1NKD1 614 868 1804 2419 • CTBP1TCF7 630 985 1120 2343 • CTBP2PPP2R1A 641 1672 2371 2245 • CTBP2DAAM1 670 1036 1567 1047 • CTBP1DVL2 671 981 2176 2380 • CTBP2GSK3B 673 2082 1321 175 • CTBP2FBXW2 677 1634 1487 556 • CTBP1TLE1 679 1071 856 1714 • CTBP2FZD1 682 2167 2126 939 • CTBP1WIF1 694 1806 2029 871 • CTBP1FBXW4 718 1496 1920 1249 • CTBP2PITX2 731 1910 1331 1292 • CTBP1DVL1 760 1255 745 473 • CTBP2DKK1 766 2099 2191 2007 • CTBP1SLC9A3R1 802 999 634 2180 • CTBP2FBXW11 823 740 707 1076 • FZD5CTBP1 830 482 1072 2252 • CTBP1FRAT1 835 2306 1816 1986 • CTBP1WNT3A 836 1562 453 1951 • CTBP2FZD8 853 1864 562 1777 • CTBP1GSK3A 858 1560 381 476 • CTBP1WNT2 874 2304 1571 2070 • AXIN1CTBP2 876 1471 625 1807 • CCND2CTBP2 916 1898 1771 317 • CTBP2TCF7 936 1305 254 1468 • CTBP2NLK 941 926 200 841 • BCL9CTBP2 961 1574 1703 740 • CTBP1MYC 980 2321 2387 1558 • CTBP2PPP2CA 984 835 2336 1644 • CTBP1FOSL1 1006 1673 1333 2452 • CTBP2CXXC4 1015 91 983 380 • CTBP2FRZB 1030 766 1187 728 • CTBP1FOXN1 1037 1936 2015 2383 • CTBP2TLE1 1079 2019 2081 1050 • AESCTBP2 1097 1785 779 256 • CCND1CTBP2 1098 2128 243 2268 • CTBP1WNT1 1115 1666 2324 1591 • BTRCCTBP2 1124 577 1940 2295 • CTBP2FBXW4 1127 644 2127 1996 • CSNK1DCTBP2 1146 668 1464 746 • CTBP1KREMEN1 1150 2058 1177 2448 • CTBP2WNT4 1186 767 1115 1598 • CTBP2TLE2 1188 1129 1300 560 • CTBP2DVL2 1245 459 2472 2451 • CSNK1A1CTBP1 1246 1429 902 1161 • CTBP1FSHB 1252 2124 2369 296 • CTBP1T 1264 1456 2210 1278 • CTBP2LRP5 1287 1991 2454 572 • CTBP2NKD1 1301 971 566 1801 • CTBP1JUN 1327 2302 2385 973 • APCCTBP2 1382 1196 1913 1885 • CTBP2DVL1 1385 550 1538 459 • CTBP1CTNNBIP1 1386 630 1030 564 • CTBP2WNT2 1392 1215 1877 2407 • CTBP2EP300 1400 53 2462 1268 • CSNK1DCTBP1 1454 104 1129 775 • CTBP2GSK3A 1456 1718 1611 1020 • CTBP2WIF1 1510 2164 2440 1125 • CTBP1WNT2B 1513 2083 2422 873 • CTBP1CTNNB1 1520 413 1847 2265 • CTBP2SLC9A3R1 1527 931 1173 1132 • BCL9CTBP1 1536 402 2119 1928 • CCND1CTBP1 1539 377 301 1875 • CTBP1DIXDC1 1577 1350 1127 1313 • CTBP2WNT1 1636 894 942 906 • CTBP2FOXN1 1651 774 1241 1515 • CTBP2FOSL1 1676 2237 1734 2205 • CCND2CTBP1 1695 730 765 2089 • CTBP2T 1713 1458 1833 512 • CTBP2FRAT1 1716 951 1762 913 • CTBP2FSHB 1719 1656 2014 880 • AESCTBP1 1751 689 97 1252 • CTBP1RHOU 1797 2093 1390 706 • CTBP1WNT3 1807 1978 1503 2206 • AXIN1CTBP1 1816 79 237 1824 • APCCTBP1 1828 600 1912 2101 • CTBP2KREMEN1 1836 722 921 2080 • CTBP2MYC 1848 941 2037 1692 • CTBP1PORCN 1859 1801 2027 1718 • BTRCCTBP1 1868 826 1046 1696 • CCND3CTBP2 1909 2357 573 1338 • CTBP1PYGO1 1927 1984 1268 1537 • CTBP2WNT2B 1932 1644 980 1138 • CSNK1G1CTBP1 1951 14 1441 2465 • CTBP2DIXDC1 1978 1166 1221 1254 • CTBP2WNT3A 1989 1412 1452 2078 • CTBP2PORCN 2115 882 1946 1796 • CTBP2CTNNBIP1 2139 433 2261 375 • CCND3CTBP1 2168 834 639 1958 • CSNK1G1CTBP2 2184 242 934 1964 • CTBP2JUN 2194 1401 1889 876 • CTBP2WNT3 2257 2165 664 2335 • CSNK2A1CTBP2 2276 2203 176 1359 • CTBP2CTNNB1 2325 7 2464 2111 • CTBP2PYGO1 2350 634 2479 2214 • CTBP2RHOU 2427 1791 2246 1453 • CSNK2A1CTBP1 2432 492 118 2346

## CCND family 2*^nd^* order interaction ranking at different time points

• APCCCND2 59 1759 1426 785 603 • APCCCND1 81 2340 1295 1390 21 • CCND3GSK3A 105 1200 670 115 2399 • FZD5CCND1 111 2278 1115 2385 105 • CCND3FOXN1 232 1038 1711 2442 2397 • CCND3PYGO1 234 580 1064 391 1756 • CCND3JUN 247 746 2338 290 318 • CCND3FZD2 258 1731 2304 1894 2278 • CCND3WNT1 273 8 405 2253 1037 • CCND3CSNK1A1 283 1159 919 1318 1776 • FZD5CCND2 298 905 477 1577 134 • CCND3MYC 323 1673 1154 1534 1580 • CCND3FGF4 342 966 2040 1448 1101 • CCND3SFRP1 383 135 1562 418 304 • CCND3WIF1 403 570 2101 620 1854 • CCND3PORCN 453 1324 2418 1961 611 • CCND2GSK3A 498 662 676 1762 1632 • CCND3WNT3 509 658 1636 716 1762 • CCND3LEF1 514 713 2295 722 2402 • CCND3DVL1 518 1028 1931 288 2010 • CCND2CSNK1A1 526 1742 1616 995 1586 • AESCCND1 528 1970 2090 2314 119 • CCND3T 531 564 890 2073 726 • CCND3CSNK1G1 552 1787 1374 2464 965 • CCND2WNT1 573 2125 187 623 1535 • APCCCND3 610 1689 125 1193 204 • CCND3DVL2 621 104 2358 2223 1751 • BTRCCCND1 634 397 2366 2273 303 • CCND3CTBP2 652 1909 1180 1441 812 • AXIN1CCND1 692 2087 2328 411 22 • CCND3NKD1 695 836 1294 327 1557 • CCND3CTNNBIP1 696 1777 2264 937 1705 • BTRCCCND2 707 793 2289 106 135 • CCND3LRP6 709 1526 2401 2373 875 • FZD5CCND3 718 2014 154 2023 814 • CCND3CTNNB1 774 539 2253 1813 1136 • CCND2PYGO1 776 1873 1534 1184 1529 • CCND3TLE1 791 112 2125 40 2430 • CCND2PORCN 837 2407 755 302 139 • CCND3FSHB 842 1814 887 753 49 • CCND2FZD2 870 249 1470 2363 1493 • AXIN1CCND2 893 1509 1259 1046 1061 • CCND2FOXN1 899 557 69 970 1389 • CCND3FBXW4 905 61 1333 2205 1979 • CCND3SFRP4 913 2364 2027 183 1422 • CCND3DAAM1 921 351 545 757 2311 • BCL9CCND1 922 2093 1048 1093 257 • CCND1JUN 942 1578 797 2143 617 • CCND3PITX2 958 422 1062 521 855 • CCND1SFRP1 1007 273 975 1536 2145 • CCND2DVL1 1013 821 1822 1143 2046 • AESCCND2 1021 220 1190 68 1752 • CCND2WIF1 1027 1657 2400 2066 1084 • CCND2MYC 1037 1196 536 2283 1238 • CCND1GSK3A 1041 184 312 957 1447 • CCND2JUN 1043 1643 1473 2329 559 • CCND2SFRP1 1050 2090 1329 1294 820 • CCND3KREMEN1 1073 382 1142 2396 2003 • CCND3FZD6 1097 538 426 2012 907 • CCND3PPP2CA 1104 315 1299 1817 454 • CCND3FBXW2 1143 391 1957 1745 495 • CCND3CTBP1 1176 981 2210 764 1497 • CCND3SLC9A3R1 1191 140 1985 1823 1152 • CCND2FGF4 1192 1506 2350 2156 1092 • CCND3FZD8 1193 1178 1922 1647 1403 • CCND3NLK 1250 1613 1808 2301 1401 • CCND2LEF1 1265 896 1282 2018 2476 • CCND3GSK3B 1318 687 2317 2284 1327 • CCND1WNT1 1325 101 302 1480 2234 • CCND3CXXC4 1327 757 1413 626 2088 • CCND3WNT2B 1347 1229 1193 88 1172 • CCND3LRP5 1373 1582 2403 1571 2006 • CCND1WIF1 1380 626 932 919 215 • CCND3EP300 1381 673 1600 256 2235 • CCND3DIXDC1 1392 1878 1604 1621 1810 • CCND3TCF7 1396 462 964 2257 1674 • CCND3WNT5A 1411 280 1588 1899 503 • CCND1FGF4 1425 432 1590 2042 2131 • CCND3PPP2R1A 1431 886 1168 1704 2030 • CCND2WNT3 1432 19 1181 2472 2100 • CCND3CSNK1D 1470 1437 594 2474 2122 • CCND1CSNK1A1 1480 994 546 1800 281 • CCND2CTNNBIP1 1493 867 1734 95 1017 • CCND2LRP6 1495 1336 1275 1432 2216 • CCND3WNT2 1506 1859 2166 549 1904 • CCND1DVL1 1512 29 1947 174 1150 • CCND3CSNK2A1 1532 1436 1800 2480 1522 • CCND3WNT3A 1546 259 2413 2291 1320 • CCND1FOXN1 1577 1635 472 2481 1738 • CCND1CTBP2 1580 983 775 2218 713 • CCND2T 1581 709 1549 1760 1445 • CCND1PYGO1 1587 150 442 1044 1118 • CCND1MYC 1593 1459 823 2312 1218 • CCND2CSNK1G1 1594 947 523 954 2229 • AXIN1CCND3 1603 1901 857 621 673 • CCND3WNT4 1607 1498 1262 141 1228 • CCND3RHOU 1625 1030 1821 2011 426 • BCL9CCND2 1628 1965 479 2352 351 • CCND2CTBP2 1629 1344 2162 1885 1831 • CCND3TLE2 1644 1435 2034 801 2188 • CCND3FRAT1 1655 1381 2452 1382 1947 • CCND3FOSL1 1660 1862 1704 731 915 • CCND3FBXW11 1666 2158 1977 1020 1916 • CCND1WNT3 1682 347 1746 2140 1721 • CCND1FZD2 1683 225 1228 1975 873 • CCND2DVL2 1686 373 1488 874 889 • CCND1LEF1 1695 528 1116 2333 761 • CCND2NKD1 1706 2469 754 1175 2114 • CCND3FZD7 1730 269 1117 298 1416 • CCND2SLC9A3R1 1745 1718 1856 130 2161 • CCND2FBXW4 1767 1915 911 2404 2400 • CCND2SFRP4 1771 1215 1994 1548 1509 • CCND2PPP2CA 1781 936 445 493 1247 • CCND2CTNNB1 1806 89 1447 767 1077 • CCND2WNT2 1823 1014 1591 1234 2197 • CCND2CTBP1 1840 1755 1657 1092 353 • CCND1SFRP4 1859 1897 2376 1787 1272 • CCND1NKD1 1864 1316 722 649 888 • CCND1CCND2 1871 449 1460 265 1469 • CCND1DVL2 1877 754 821 518 1340 • CCND3FZD1 1878 435 1506 1758 1216 • CCND2KREMEN1 1884 943 717 1631 2370 • CCND2PPP2R1A 1911 791 831 681 2198 • CCND2FSHB 1915 464 583 522 1729 • CCND2DAAM1 1928 1211 1960 966 1811 • CCND2WNT5A 1932 1348 1126 1650 2156 • BTRCCCND3 1933 760 2147 1071 555 • CCND2TCF7 1945 808 360 2118 1781 • CCND2NLK 1949 2136 389 1550 102 • CCND1CSNK1G1 1951 1991 1021 1807 644 • CCND2PITX2 1960 1278 1204 23 2123 • CCND1T 1967 1789 803 2272 1547 • CCND2CSNK2A1 1969 2326 833 2344 1165 • CCND2FZD6 1980 1712 588 2297 1120 • AESCCND3 1991 399 567 1622 2209 • CCND1PORCN 1993 185 671 2124 2326 • CCND2TLE1 1998 2030 1826 849 748 • CCND3TCF7L1 2001 1103 2434 1915 2168 • CCND2CXXC4 2002 2029 870 486 2085 • CCND1WNT2 2014 894 1655 1444 777 • CCND2GSK3B 2022 1922 2091 257 1962 • CCND1LRP6 2040 213 848 2107 1675 • CCND2EP300 2046 763 1895 2155 2243 • CCND1CTNNBIP1 2063 650 2011 1364 549 • CCND2FZD8 2072 1391 1820 934 1913 • CCND1FZD8 2073 197 1777 1202 2324 • CCND2FZD1 2075 536 2294 392 266 • CCND2DIXDC1 2076 354 2026 802 969 • CCND1PITX2 2078 149 2077 735 1369 • CCND1TLE1 2080 610 1516 1419 1246 • CCND2TCF7L1 2091 1628 1322 1549 1902 • CCND2LRP5 2100 635 1585 787 2425 • CCND2WNT3A 2104 1426 2360 2389 2136 • CCND1SLC9A3R1 2107 51 2479 929 1476 • CCND2FBXW2 2114 1187 747 1069 973 • CCND2FRAT1 2116 963 1202 618 1763 • CCND2CSNK1D 2126 781 982 2434 1955 • CCND1CXXC4 2128 928 2480 784 1367 • CCND2CCND3 2129 319 604 1210 1622 • CCND1CTNNB1 2132 769 904 1491 2217 • CCND2FBXW11 2140 295 214 1912 2465 • CCND2RHOU 2146 372 2299 595 2410 • CCND1EP300 2148 801 1267 464 1127 • BCL9CCND3 2159 2140 519 1909 471 • CCND1FBXW4 2193 31 370 1957 1245 • CCND1PPP2CA 2203 252 226 58 1990 • CCND1DAAM1 2237 77 653 818 1850 • CCND1CSNK2A1 2248 18 1442 1690 920 • CCND3DKK1 2253 1058 704 2078 2295 • CCND2WNT4 2259 1083 811 291 116 • CCND2WNT2B 2261 671 337 519 1961 • CCND1FSHB 2262 1906 64 1835 488 • CCND1GSK3B 2274 426 1308 142 162 • CCND2FZD7 2290 2053 1495 2183 948 • CCND1FZD6 2297 365 321 1290 2265 • CCND1WNT4 2320 1816 1238 1510 1232 • CCND2FOSL1 2323 465 2094 1304 1614 • CCND1WNT3A 2326 1545 2375 2343 1937 • CCND3FRZB 2337 1895 1212 633 988 • CCND1FZD1 2350 526 927 1803 677 • CCND1WNT5A 2352 534 662 1659 2470 • CCND1LRP5 2358 1295 1123 2240 2106 • CCND1KREMEN1 2360 893 1206 1945 1521 • CCND1CTBP1 2361 1995 970 1366 1147 • CCND2TLE2 2368 1333 1276 2175 1910 • CCND1FOSL1 2369 1647 2468 768 857 • CCND1PPP2R1A 2370 219 341 585 1489 • CCND1FBXW2 2373 404 504 1522 1862 • CCND1WNT2B 2374 1440 228 112 612 • CCND1CCND3 2375 1090 419 779 1571 • CCND1NLK 2379 890 698 467 2218 • CCND1FZD7 2381 357 2417 1320 2181 • CCND1FRAT1 2385 1310 1832 1509 1995 • CCND1TCF7 2393 682 220 1588 2165 • CCND1CSNK1D 2400 1450 1362 1941 1325 • CCND1TCF7L1 2402 552 1394 554 1794 • CCND1DIXDC1 2408 1658 1605 386 1536 • CCND2DKK1 2417 1387 141 657 1043 • CCND1FBXW11 2429 226 635 886 949 • CCND1RHOU 2431 853 993 777 1300 • CCND1TLE2 2434 1503 1743 1501 551 • CCND3SENP2 2444 216 2181 644 858 • CCND1DKK1 2459 999 562 2437 1091 • CCND2FRZB 2461 764 2310 1854 1963 • CCND1SENP2 2471 555 1824 53 1640 • CCND2SENP2 2474 1527 1866 597 1670 • CCND1FRZB 2475 1944 2196 2146 1400

## CCND family 2*^nd^* order interaction ranking at different durations

• CCND1SENP2 72 2154 95 1956 • CCND2SENP2 76 1613 649 1811 • CCND2SFRP4 92 1257 2205 1634 • CCND2CSNK1G1 101 1271 965 2063 • CCND2FGF4 109 1594 1473 2139 • CCND2LRP6 142 1867 1089 1165 • CCND1SFRP4 143 2081 310 431 • CCND2FZD2 158 976 853 1673 • CCND1LEF1 174 1206 627 1769 • CCND2LEF1 181 1741 1886 2376 • CCND1LRP6 196 2120 789 861 • CCND2FZD7 201 1487 355 2055 • CCND1FZD2 208 1829 1254 1987 • CCND1CSNK1G1 218 1708 595 2066 • CCND2FBXW2 221 1706 1603 688 • FZD5CCND3 239 500 2250 2312 • CCND1FZD1 243 1665 1328 869 • CCND1FGF4 257 1852 1982 1920 • CCND3SENP2 258 1754 684 793 • CCND1FBXW11 262 756 373 2151 • CCND1FBXW2 266 1705 528 1905 • CCND2SFRP1 280 728 2251 2291 • FZD5CCND1 290 662 1582 2197 • CCND2PITX2 309 1527 1076 1178 • CCND1PPP2R1A 323 779 429 1222 • CCND1CSNK2A1 331 1085 1003 2220 • CCND2TLE2 335 1683 1217 193 • CCND2CSNK2A1 338 1607 1087 2365 • CCND1FZD7 342 2364 1725 1680 • CCND2FBXW11 348 1409 1037 1403 • CCND1DAAM1 373 698 1895 1691 • CCND2GSK3B 380 940 936 687 • CCND2FZD1 385 1394 2006 1651 • CCND1FZD6 391 1977 2285 472 • CCND2TCF7L1 406 1089 1879 1271 • CCND1TLE2 421 575 689 492 • CCND2CXXC4 429 225 1057 1887 • CCND2WNT5A 434 1921 1981 2260 • CCND1TCF7L1 441 1604 784 2355 • CCND2DAAM1 443 2428 938 1532 • CCND2FZD6 449 1274 905 2002 • CCND2PPP2R1A 450 1267 1045 1619 • CCND1SFRP1 462 1755 671 1480 • CCND2NLK 488 620 1623 787 • CCND2FZD8 502 1460 961 1556 • FZD5CCND2 503 364 2025 2324 • CCND1PITX2 510 1529 2470 825 • CCND2WNT4 511 1056 1461 2029 • CCND1WNT5A 513 929 412 1509 • CCND1GSK3B 515 1842 1961 907 • CCND1CXXC4 517 57 353 1055 • CCND2CCND3 541 1808 1628 1833 • CCND2DVL2 546 2129 1402 2373 • CCND1NLK 560 1435 147 2179 • CCND2DKK1 573 1366 341 639 • CCND2PPP2CA 602 2451 1178 883 • CCND1CCND3 609 642 317 2367 • CCND1PPP2CA 610 757 829 872 • CCND2TLE1 622 1192 1552 1528 • CCND1FZD8 624 2184 2229 1829 • CCND2LRP5 637 1710 1691 914 • CCND3SFRP4 640 1797 346 142 • CCND1FBXW4 654 557 1488 1657 • BCL9CCND3 655 631 2391 1263 • CCND3LRP6 657 1637 1801 1284 • AESCCND3 665 624 1842 908 • CCND2FRZB 666 2340 715 381 • CCND2FBXW4 678 2355 950 1443 • CCND2NKD1 704 1023 1456 1830 • CCND1TLE1 710 959 1079 1895 • CCND1TCF7 732 877 61 1750 • CCND2EP300 767 362 1942 1063 • CCND2TCF7 777 1675 194 2285 • CCND3CSNK1G1 782 1996 600 2456 • CCND1DVL2 784 759 513 2147 • CCND1WNT4 790 1019 1280 1348 • CCND1DKK1 798 2216 158 2134 • CCND1FRZB 818 568 213 1770 • AXIN1CCND3 827 547 422 2259 • CCND1EP300 834 32 1389 2387 • CCND3PPP2R1A 871 963 1863 2279 • CCND3SFRP1 883 1870 1830 820 • CCND3FZD7 887 2176 918 1559 • CCND2FOSL1 888 1383 1994 2006 • CCND3LEF1 890 2173 1733 1157 • CCND1LRP5 896 1699 1454 1308 • CCND3CSNK2A1 907 2390 1999 1752 • CCND3FZD2 915 2276 1922 1516 • CCND2CTBP2 916 1898 1771 317 • CCND2WIF1 931 1671 2434 2161 • CCND3TCF7L1 934 966 542 2363 • CCND3FZD1 935 1032 697 1852 • CCND3FGF4 946 1382 2231 1289 • CCND2T 956 1974 2308 136 • BTRCCCND3 987 407 2224 1485 • CCND1FSHB 995 2060 292 756 • CCND2DVL1 1019 691 1001 498 • CCND2SLC9A3R1 1041 1954 1409 1551 • CCND2MYC 1051 1983 2026 2384 • AXIN1CCND1 1052 1090 842 2122 • CCND3FBXW2 1053 1740 699 616 • CCND1NKD1 1059 368 184 649 • CCND1SLC9A3R1 1063 1638 1008 1513 • CCND3FZD8 1073 1174 889 2334 • CCND1MYC 1074 661 295 1870 • CCND1CTBP2 1098 2128 243 2268 • CCND1DVL1 1101 1117 2192 745 • CCND1FRAT1 1102 2073 611 936 • CCND3WNT5A 1105 2445 1048 1923 • CCND3FBXW11 1107 2455 655 1749 • CCND2GSK3A 1113 2359 1066 1712 • CCND2FSHB 1122 1565 2174 507 • CCND2WNT2 1123 1374 1105 1955 • BCL9CCND1 1125 1646 709 1420 • CCND1GSK3A 1126 1122 2370 652 • CCND1FOSL1 1138 2116 299 1844 • CCND1WNT1 1145 780 933 842 • CCND2CSNK1D 1153 1545 2331 1065 • CCND3PPP2CA 1161 2020 603 387 • CCND3GSK3B 1162 933 1903 1127 • APCCCND3 1163 2300 2199 2447 • CCND3PITX2 1191 1453 1505 626 • CCND3NLK 1197 2101 512 1808 • CCND2CTNNBIP1 1204 2387 739 1293 • CCND1WNT2 1206 1426 661 1671 • BCL9CCND2 1215 528 2079 1384 • CCND3TLE2 1227 2157 1447 1277 • AXIN1CCND2 1236 167 258 1622 • CCND3FZD6 1238 1913 2023 800 • CCND2WNT1 1247 1285 941 1590 • CCND3CXXC4 1261 696 363 1059 • CCND3DKK1 1276 1720 1012 2345 • CCND3TCF7 1286 1916 563 2086 • CCND1FOXN1 1292 655 1299 1192 • CCND2FRAT1 1320 1238 1317 1375 • CCND2FOXN1 1331 749 621 817 • CCND2WNT3A 1338 639 736 2105 • CCND1CSNK1D 1344 910 711 1586 • CCND2DIXDC1 1348 960 1081 1257 • CCND2KREMEN1 1349 2188 895 1855 • CCND3DAAM1 1357 1459 847 1699 • CCND1WIF1 1359 1632 454 905 • CCND2JUN 1369 2112 1854 998 • CCND1CCND2 1375 545 150 1972 • CCND1KREMEN1 1378 2027 2366 2247 • AESCCND2 1379 764 308 39 • CCND3FRZB 1408 1639 146 865 • CCND3DVL2 1411 1771 1359 2356 • CCND3WNT4 1443 1320 1926 1251 • AESCCND1 1460 2180 2202 268 • CCND1CTNNBIP1 1481 732 907 1843 • BTRCCCND1 1504 1198 1669 777 • CCND3NKD1 1516 961 268 2339 • CCND3FBXW4 1519 2144 943 2049 • CCND1JUN 1529 622 275 1641 • CCND3EP300 1534 399 2425 1246 • CCND1CTBP1 1539 377 301 1875 • CCND1DIXDC1 1541 1067 2190 334 • CCND3TLE1 1589 1937 1844 1235 • CCND3LRP5 1595 895 1137 367 • CCND3FOSL1 1604 2232 967 1607 • CCND3SLC9A3R1 1621 1428 1573 2269 • CCND2CTNNB1 1630 131 1431 1126 • CCND1T 1634 563 356 206 • CCND1WNT3A 1642 2089 2378 1191 • CCND3WNT1 1643 2475 1077 1825 • CCND2WNT2B 1668 2309 1214 900 • CCND2CTBP1 1695 730 765 2089 • CCND1PORCN 1701 851 728 2297 • APCCCND1 1717 1281 931 2187 • CCND1CTNNB1 1720 99 447 952 • CCND2WNT3 1729 1779 1166 2468 • CCND3FSHB 1741 1492 809 370 • CCND3WNT2B 1749 2421 795 925 • CCND3DVL1 1759 1097 1109 948 • CCND3WNT2 1775 1758 1562 2196 • BTRCCCND2 1804 643 478 1120 • CCND2RHOU 1813 2023 1827 790 • CCND1RHOU 1844 2161 2467 332 • CCND3FOXN1 1883 2408 1180 1082 • CCND1WNT2B 1884 2243 71 1303 • CCND3CTBP2 1909 2357 573 1338 • CCND3WIF1 1913 1736 1204 1182 • CCND3CTNNBIP1 1967 1221 1314 541 • CCND3FRAT1 1980 1462 511 1034 • CCND3T 1983 1811 738 764 • CCND3CSNK1D 1999 1095 521 1128 • CCND1WNT3 2001 1043 169 1814 • CCND3MYC 2025 1643 696 1095 • CCND3KREMEN1 2035 2281 1915 2276 • CCND3PORCN 2040 1979 2259 2392 • CCND1PYGO1 2046 1240 1667 1141 • CCND2PORCN 2064 1318 2132 690 • APCCCND2 2102 1187 2198 1979 • CCND1CSNK1A1 2161 1178 143 1544 • CCND3CTBP1 2168 834 639 1958 • CCND3RHOU 2177 1164 2003 682 • CCND3GSK3A 2212 2016 2125 715 • CCND3CTNNB1 2230 468 1398 1589 • CCND3DIXDC1 2267 1518 1253 1006 • CCND2PYGO1 2271 2048 2103 766 • CCND3WNT3A 2284 889 1281 2308 • CCND3WNT3 2337 1831 1134 759 • CCND3JUN 2355 825 553 423 • CCND2CSNK1A1 2359 729 483 1131 • CCND3PYGO1 2418 1961 857 1762 • CCND3CSNK1A1 2483 784 324 427

## CTNNB family 2*^nd^* order interaction ranking at different time points

• CTNNBIP1GSK3A 225 766 35 856 376 • CTNNB1FOXN1 351 2270 300 1168 938 • CTNNB1FGF4 352 1427 1780 1693 56 • CTNNB1LEF1 366 912 1713 2014 1252 • CTNNB1JUN 390 798 2100 1934 2260 • CTNNBIP1WNT1 400 2292 184 857 12 • CTNNB1PORCN 418 1811 357 792 111 • APCCTNNBIP1 420 2177 2464 1656 2002 • CTNNB1MYC 425 2028 576 884 2422 • CTNNB1PYGO1 429 1059 524 1962 194 • CSNK2A1CTNNBIP1 443 1454 2357 1998 2359 • CSNK2A1CTNNB1 445 289 2406 1207 1462 • CTNNB1WIF1 446 1651 1633 1664 1209 • CTNNB1GSK3A 447 1713 304 1640 186 • CTNNBIP1FZD2 450 1481 548 286 143 • CTNNB1FZD2 451 509 1424 2326 1744 • APCCTNNB1 457 1988 862 1686 412 • CTNNB1WNT1 488 2246 361 2364 759 • CTNNBIP1FGF4 499 176 1680 409 41 • CSNK1DCTNNBIP1 525 603 2412 2121 1065 • CTNNB1SFRP1 527 1687 1208 869 811 • FZD5CTNNBIP1 554 1661 1513 2275 2203 • CTNNBIP1WIF1 557 2419 2287 1606 392 • CTNNBIP1DVL1 571 2450 1347 1104 76 • CTNNBIP1LEF1 585 481 850 1154 142 • CTNNB1WNT3 627 1276 1376 1310 710 • CTNNBIP1JUN 632 1390 1614 1543 558 • CTNNBIP1WNT3 639 1093 1486 1383 2 • CSNK1DCTNNB1 657 1962 1032 114 2231 • CTNNBIP1FOXN1 658 2083 186 1729 309 • CTNNBIP1SFRP1 668 1836 2084 1830 9 • FZD5CTNNB1 684 1045 1370 1034 1840 • CCND3CTNNBIP1 696 1777 2264 937 1705 • CTNNBIP1PYGO1 716 1114 358 920 34 • CTNNBIP1MYC 729 1744 597 1905 1432 • CTNNB1DVL1 756 2353 1769 123 1645 • CCND3CTNNB1 774 539 2253 1813 1136 • CTNNBIP1PORCN 822 973 490 52 28 • CTNNB1DVL2 835 2448 674 737 1508 • CTBP1CTNNBIP1 859 967 1536 2217 1607 • CTBP1CTNNB1 909 1082 807 2164 764 • CTNNBIP1DVL2 929 2252 732 1925 1012 • CTNNBIP1LRP6 967 2427 783 1141 222 • CSNK1G1CTNNBIP1 992 1044 2454 1412 1584 • CTNNB1CTNNBIP1 1048 153 1284 289 2329 • CTNNBIP1NKD1 1053 2442 279 451 229 • CTNNB1NKD1 1081 1158 609 134 2117 • CTBP2CTNNBIP1 1125 1358 1830 1795 1877 • CSNK1G1CTNNB1 1128 2228 2386 1001 1297 • CTNNBIP1T 1155 2009 520 442 301 • CTNNB1LRP6 1170 1032 1036 1330 1434 • CTNNB1T 1215 1772 1535 1053 1487 • CTBP2CTNNB1 1254 190 2411 2375 2025 • AXIN1CTNNBIP1 1263 1198 771 1507 2441 • CTNNBIP1FBXW4 1291 2373 378 1902 716 • CTNNB1FSHB 1312 198 387 1295 1654 • CTNNB1TLE1 1322 1891 2061 887 16 • CTNNB1SFRP4 1395 1332 2424 1045 1933 • AXIN1CTNNB1 1413 1167 2074 131 1769 • CTNNB1NLK 1417 1715 168 2102 935 • CTNNBIP1PITX2 1460 346 1541 457 78 • CTNNB1PPP2CA 1473 2067 197 356 1519 • BTRCCTNNBIP1 1474 857 2433 951 1042 • CTNNB1FZD8 1482 1645 976 1592 1626 • CCND2CTNNBIP1 1493 867 1734 95 1017 • CTNNBIP1FZD6 1500 1518 251 598 212 • CTNNB1PITX2 1514 964 1778 1887 332 • AESCTNNB1 1537 1581 2389 285 2177 • CTNNB1DAAM1 1540 1979 812 1641 512 • CTNNB1EP300 1541 785 1982 1620 1926 • AESCTNNBIP1 1548 1283 1569 253 1516 • CSNK1A1CTNNBIP1 1549 2303 1015 652 2236 • CTNNBIP1WNT4 1560 1314 550 1346 405 • CTNNB1FBXW4 1565 2145 856 1617 1790 • CTNNB1WNT5A 1582 1741 322 2324 230 • CTNNB1TCF7 1591 639 417 1616 543 • CTNNB1PPP2R1A 1596 1191 813 839 457 • CTNNBIP1PPP2R1A 1599 1077 624 2338 48 • CTNNB1WNT3A 1612 710 2260 2309 2032 • CTNNBIP1DAAM1 1619 2444 774 444 3 • CTNNBIP1NLK 1620 2287 476 2210 81 • CTNNBIP1TLE1 1626 2429 930 922 326 • BTRCCTNNB1 1627 1015 1500 1599 1308 • CTNNB1KREMEN1 1634 1397 914 460 1163 • CTNNBIP1SFRP4 1636 1302 2377 64 150 • CTNNBIP1FSHB 1648 1346 91 946 747 • CTNNBIP1CXXC4 1652 1070 1865 1841 17 • CTNNB1FZD6 1665 725 222 1321 283 • CTNNBIP1FBXW2 1669 1569 1031 666 67 • CTNNBIP1WNT3A 1678 2054 1975 1349 298 • CTNNB1FBXW2 1685 1796 897 1173 414 • CTNNBIP1PPP2CA 1692 2310 71 1873 239 • CTNNB1DIXDC1 1705 1345 1684 1744 774 • CTNNB1FZD7 1716 594 957 1431 752 • CTNNBIP1EP300 1733 1669 1485 1433 385 • CTNNBIP1KREMEN1 1741 851 971 991 125 • CTNNB1FRAT1 1747 1321 1664 960 1370 • CTNNB1LRP5 1762 588 1422 2216 833 • CTNNB1FBXW11 1766 1709 460 961 1606 • CTNNBIP1TCF7 1784 2198 118 1087 241 • CTNNBIP1WNT2 1785 209 820 317 334 • CTNNB1RHOU 1791 2316 1141 1121 1345 • CTNNB1WNT4 1795 2184 1328 2003 1014 • CTNNB1CXXC4 1796 119 2186 1636 848 • CCND2CTNNB1 1806 89 1447 767 1077 • CTNNBIP1LRP5 1819 358 459 1946 459 • CTNNB1GSK3B 1826 1785 697 2126 1553 • CTNNBIP1FZD8 1832 2351 790 151 13 • CTNNBIP1FZD1 1834 1130 1412 2094 490 • CTNNB1WNT2 1847 712 1698 902 2345 • CTNNBIP1DIXDC1 1853 2428 1752 393 141 • CTNNBIP1RHOU 1860 2027 1478 654 201 • CTNNBIP1SLC9A3R1 1870 1412 2148 614 203 • CTNNB1SLC9A3R1 1885 755 2270 466 1028 • CTNNBIP1WNT2B 1897 1428 171 15 1113 • CTNNBIP1FZD7 1899 2392 2447 375 1 • BCL9CTNNBIP1 1906 25 1390 569 1796 • CSNK1A1CTNNB1 1907 965 724 1508 1250 • CTNNBIP1GSK3B 1913 2300 1689 1339 343 • CTNNBIP1FBXW11 1930 2021 103 1254 232 • CTNNB1TLE2 1939 1445 1802 2315 1873 • CTNNBIP1FRAT1 1964 2195 701 1401 706 • BCL9CTNNB1 1970 1783 2050 2128 110 • CTNNB1FZD1 1983 1544 1214 1435 1720 • CTNNB1FOSL1 1997 1036 2065 975 1827 • CTNNB1WNT2B 2023 773 591 1587 2299 • CTNNBIP1TLE2 2035 1081 1093 1216 282 • CTNNB1TCF7L1 2043 944 805 381 1449 • CCND1CTNNBIP1 2063 650 2011 1364 549 • CTNNBIP1WNT5A 2068 1367 444 1392 169• CTNNBIP1TCF7L1 2123 1739 1130 1580 43 • CCND1CTNNB1 2132 769 904 1491 2217 • CTNNBIP1FOSL1 2174 664 1571 14 1078 • CTNNBIP1DKK1 2389 1910 84 1601 44 • CTNNB1DKK1 2390 1931 276 1671 839 • CTNNBIP1FRZB 2428 264 2043 2340 121 • CTNNB1FRZB 2436 809 2458 1883 1817 • CTNNBIP1SENP2 2443 2388 1842 1635 57 • CTNNB1SENP2 2454 561 2308 413 164

## CTNNB family 2*^nd^* order interaction ranking at different durations

• CTNNB1SENP2 32 1344 234 546 • CTNNB1SFRP4 42 1294 329 233 • CTNNB1LRP6 65 1355 1570 1033 • CTNNB1FZD2 67 2280 1700 584 • CTNNBIP1SENP2 78 2396 74 624 • CTNNB1FBXW2 81 1596 1956 1568 • CTNNB1FZD6 112 1818 1028 160 • CTNNB1SFRP1 116 1555 2034 1448 • CTNNBIP1SFRP4 118 2202 465 1096 • CTNNB1LEF1 122 1051 2360 1755 • CTNNB1PITX2 138 783 2301 1357 • FZD5CTNNB1 145 159 1846 2321 • CTNNB1FZD7 148 1680 1091 636 • CTNNB1TCF7L1 157 1641 1213 2213 • FZD5CTNNBIP1 165 367 1828 1297 • CTNNB1PPP2R1A 177 1614 1520 1805 • CTNNB1DKK1 184 996 507 2200 • CTNNB1FZD8 187 1814 1553 2481 • CTNNB1FZD1 188 1652 2415 2237 • CTNNB1FGF4 190 1341 1600 357 • CTNNB1CXXC4 191 1404 1597 409 • CTNNB1FBXW11 205 2122 354 1884 • CTNNB1GSK3B 223 1278 1084 1491 • CTNNB1WNT5A 263 1836 1168 1325 • CTNNBIP1FBXW2 269 1311 419 1217 • CTNNB1DAAM1 271 2407 1728 730 • CTNNB1TLE2 278 1523 805 1417 • CTNNBIP1FZD2 282 2411 2052 852 • CTNNBIP1PPP2R1A 288 2462 499 517 • CTNNB1PPP2CA 293 2018 1905 563 • CTNNB1FBXW4 302 1650 1192 391 • CTNNBIP1WNT5A 314 1953 1124 1473 • CTNNB1TLE1 327 1631 2325 1389 • CTNNBIP1LRP6 337 2043 385 2266 • CTNNBIP1FBXW11 347 2394 208 2065 • CTNNB1WNT4 352 2035 2388 2457 • CTNNBIP1LEF1 361 2223 1789 1398 • CTNNBIP1FZD1 366 1561 626 1011 • CTNNB1NLK 368 2367 382 2271 • CTNNBIP1FGF4 378 1733 1547 994 • CTNNBIP1SFRP1 388 2213 912 1768 • CTNNBIP1FZD8 395 2178 2009 1761 • CTNNBIP1FZD7 405 1822 2011 553 • CTNNBIP1DAAM1 408 1800 2361 1526 • CTNNBIP1FZD6 417 2224 1238 1168 • CTNNBIP1GSK3B 444 1820 1196 146 • CTNNB1WNT2 456 1376 1741 1976 • CTNNB1FRZB 469 2053 411 916 • CTNNBIP1TCF7L1 487 2088 417 551 • CTNNB1DVL1 521 2265 2414 119 • CTNNB1EP300 524 970 1979 520 • CTNNB1LRP5 540 1059 1038 1549 • CTNNBIP1TLE2 545 2246 544 164 • CTNNB1DVL2 569 791 1874 1274 • CTNNBIP1PPP2CA 583 2220 620 1248 • CTNNBIP1PITX2 593 1880 1016 595 • CTNNB1TCF7 597 1307 261 1851 • CTNNBIP1TCF7 608 2177 72 530 • CTNNB1SLC9A3R1 644 1272 2461 384 • CTNNBIP1CXXC4 653 2248 247 1201 • CTNNBIP1NLK 683 2034 84 1954 • CTNNBIP1TLE1 696 2282 871 1561 • CTNNB1NKD1 699 1143 547 1588 • CTNNBIP1DKK1 728 1774 138 1121 • CTNNBIP1EP300 734 2482 1341 1772 • CTNNB1GSK3A 747 1408 1689 267 • CTNNB1WNT1 753 2052 2379 1931 • CSNK1A1CTNNBIP1 754 546 1379 47 • CTNNBIP1SLC9A3R1 773 2214 1401 2288 • CTNNB1KREMEN1 805 1213 2343 1534 • CTNNB1FOSL1 814 1160 1020 516 • CTNNB1MYC 825 1712 423 1237 • CTNNB1T 850 2080 1911 1179 • CTNNBIP1DVL2 862 2415 1360 2182 • CTNNBIP1FRZB 889 2221 230 248 • CTNNBIP1FOSL1 893 1816 1014 1118 • CTNNBIP1FBXW4 901 2349 179 969 • CTNNBIP1LRP5 917 2127 430 260 • CTNNBIP1WNT2 923 2463 1938 1198 • CTNNBIP1WNT4 944 1567 1139 1286 • CTNNB1WNT3A 972 1044 1853 2353 • CTNNBIP1FSHB 989 2329 1157 1282 • CTNNB1CTNNBIP1 1038 2117 568 736 • CTNNB1FSHB 1045 1868 1275 204 • CSNK1A1CTNNB1 1047 583 1820 252 • CTNNB1WIF1 1061 1113 1701 363 • CTNNB1WNT2B 1093 1297 937 40 • CTNNBIP1FOXN1 1110 1511 1432 684 • CTNNBIP1NKD1 1116 1275 173 700 • CTNNBIP1FRAT1 1160 2210 814 506 • CTNNBIP1MYC 1181 2409 379 1927 • CCND2CTNNBIP1 1204 2387 739 1293 • CTNNBIP1T 1233 1463 688 282 • CTNNBIP1WNT2B 1235 2153 643 2149 • CTNNB1FOXN1 1244 1392 1316 1149 • AESCTNNBIP1 1254 536 2316 45 • CTNNBIP1JUN 1271 1837 459 640 • CTNNBIP1GSK3A 1346 2143 597 2138 • CTBP1CTNNBIP1 1386 630 1030 564 • CTNNBIP1WIF1 1387 1689 505 732 • CTNNBIP1WNT3A 1426 1654 2153 1204 • CTNNBIP1KREMEN1 1439 2346 1526 826 • CTNNB1WNT3 1445 1280 572 941 • BCL9CTNNBIP1 1471 1068 1332 1504 • CCND1CTNNBIP1 1481 732 907 1843 • CTNNB1FRAT1 1483 1704 1519 2347 • CSNK1DCTNNBIP1 1509 233 1130 153 • CTNNB1JUN 1511 1387 913 1531 • CTBP1CTNNB1 1520 413 1847 2265 • CTNNBIP1DVL1 1530 770 2399 1748 • APCCTNNBIP1 1547 1343 848 909 • CTNNBIP1WNT1 1550 2172 702 798 • CTNNB1DIXDC1 1615 860 2143 216 • CSNK1DCTNNB1 1616 23 1430 1117 • CCND2CTNNB1 1630 131 1431 1126 • AXIN1CTNNBIP1 1683 323 1167 761 • BTRCCTNNBIP1 1708 1020 2323 95 • CCND1CTNNB1 1720 99 447 952 • CTNNB1RHOU 1769 1584 2446 1961 • APCCTNNB1 1772 1467 1313 791 • CTNNBIP1PORCN 1778 2358 622 2244 • AXIN1CTNNB1 1830 171 1675 1611 • CTNNBIP1PYGO1 1855 2368 1443 1085 • CTNNBIP1DIXDC1 1876 1901 1449 1888 • CTNNB1PYGO1 1891 1824 2280 2292 • CTNNBIP1RHOU 1939 1997 1947 221 • BTRCCTNNB1 1962 239 2055 845 • CCND3CTNNBIP1 1967 1221 1314 541 • CTNNBIP1WNT3 1991 2433 64 903 • BCL9CTNNB1 2066 176 2375 511 • AESCTNNB1 2106 229 1219 721 • CTBP2CTNNBIP1 2139 433 2261 375 • CCND3CTNNB1 2230 468 1398 1589 • CTBP2CTNNB1 2325 7 2464 2111 • CSNK1G1CTNNB1 2354 1 2099 824 • CSNK2A1CTNNBIP1 2356 190 851 1388 • CTNNB1PORCN 2377 1611 409 833 • CSNK2A1CTNNB1 2434 314 610 1711 • CSNK1G1CTNNBIP1 2481 185 1545 618

## CSNK family 2*^nd^* order interaction ranking at different time points

• CSNK1DFOXN1 51 2149 868 1245 2365 • CSNK2A1GSK3A 53 991 746 2186 1207 • CSNK2A1PYGO1 82 388 2066 1551 1693 • CSNK2A1FGF4 102 805 2073 2111 89 • CSNK1DGSK3A 119 933 261 1319 680 • CSNK1DPYGO1 128 2403 908 2260 1612 • CSNK1DWNT1 150 2436 568 2316 1021 • CSNK2A1WIF1 177 1227 2281 1676 714 • FZD5CSNK1A1 182 977 538 1672 2352 • CSNK1DPORCN 184 2144 2399 1417 1154 • CSNK2A1FOXN1 205 1624 645 1088 826 • CSNK2A1DVL1 209 1585 2353 1363 1798 • CSNK2A1FZD2 210 703 1468 1486 1004 • CSNK2A1LEF1 214 50 2456 2435 1601 • CSNK2A1PORCN 219 2117 1782 2180 47 • CSNK2A1MYC 222 1837 1012 2176 2384 • APCCSNK1A1 235 2035 684 47 500 • CSNK1DFZD2 237 1453 2461 971 1474 • CSNK1DJUN 246 2221 2422 897 1316 • CSNK1DMYC 250 2298 1695 1460 2125 • CSNK2A1WNT1 251 1172 793 2070 532 • CSNK2A1DVL2 256 369 910 1809 2254 • CSNK1DWIF1 257 2461 2352 910 2094 • CSNK2A1SFRP1 263 329 2229 1960 46 • CSNK2A1JUN 270 1151 2192 2372 1908 • CSNK1DFGF4 281 949 1626 1169 2310 • CCND3CSNK1A1 283 1159 919 1318 1776 • CSNK1DWNT3 289 2103 2262 2354 898 • CSNK2A1WNT3 322 70 2143 2019 788 • CSNK2A1LRP6 325 1002 1245 1483 589 • CSNK1DT 329 1374 2278 1696 1108 • APCCSNK1G1 332 1558 809 2204 260 • CSNK1DDVL1 341 1955 1163 25 1921 • CSNK1DSFRP1 348 2025 1956 990 568 • CSNK1DLRP6 361 722 1380 861 1616 • CSNK1G1FOXN1 385 1700 913 1749 2227 • CSNK1DCTBP2 416 2400 1630 1236 1660 • CSNK2A1CTNNBIP1 443 1454 2357 1998 2359 • CSNK1G1SFRP1 444 1208 2459 1950 2354 • CSNK2A1CTNNB1 445 289 2406 1207 1462 • CSNK1G1GSK3A 462 113 1893 300 2462 • CSNK1DDVL2 463 1400 2348 1808 1158 • CSNK2A1CTBP2 464 2130 2347 1916 1914 • CSNK1G1FGF4 468 707 1862 596 620 • CSNK1DLEF1 474 700 1358 1068 2153 • CSNK2A1NKD1 478 1232 1437 2271 1464 • CSNK2A1T 483 2129 2005 1113 493 • CSNK1DCSNK1G1 511 2199 1554 2226 1697 • CSNK1DCTNNBIP1 525 603 2412 2121 1065 • CCND2CSNK1A1 526 1742 1616 995 1586 • AXIN1CSNK1A1 551 1603 1908 2244 2014 • CCND3CSNK1G1 552 1787 1374 2464 965 • CSNK2A1FZD6 556 258 922 1717 188 • CSNK1G1FZD2 563 1738 2268 523 1143 • CSNK1DFSHB 583 659 345 48 2258 • CSNK1G1MYC 588 2186 782 1172 2293 • CSNK1DNKD1 596 2438 640 2215 1868 • CSNK1G1JUN 608 1606 2217 1296 1456 • FZD5CSNK1G1 609 1137 484 2000 577 • CSNK2A1FBXW4 613 1164 1627 533 1031 • CSNK1DEP300 617 1008 1917 760 108 • CSNK2A1TLE1 620 1446 2133 1720 92 • CSNK1DTLE1 630 2120 2311 1500 981 • CSNK1G1PYGO1 631 2315 1941 828 2285 • CSNK1DCTNNB1 657 1962 1032 114 2231 • CSNK2A1KREMEN1 666 683 2023 1467 1639 • CSNK1DFZD8 669 2068 2145 2307 233 • CSNK2A1EP300 675 2133 1563 1980 2151 • CSNK2A1SLC9A3R1 678 33 1965 730 721 • CSNK2A1FSHB 699 384 2053 1936 785 • CSNK1G1WIF1 702 2409 2030 877 1348 • CSNK1DPPP2CA 705 1681 1060 1300 2455 • CSNK2A1DAAM1 711 155 2444 1863 563 • CSNK1DSFRP4 713 1697 2107 441 2374 • CSNK1G1WNT3 719 1555 2460 1456 520 • CSNK1G1DVL1 731 1263 1218 363 806 • CSNK1DSLC9A3R1 732 1705 2431 2144 1415 • CSNK1A1GSK3A 742 1261 483 2020 377 • CSNK1G1LEF1 744 368 2441 2027 2395 • CSNK2A1CXXC4 758 60 2296 2457 124 • CSNK2A1SFRP4 760 2335 2102 1722 1199 • CSNK2A1FBXW2 764 907 2038 2445 74 • CSNK2A1CTBP1 768 221 1236 2213 1885 • CSNK1DCTBP1 775 417 2320 923 1107 • CSNK2A1FZD8 778 202 2049 1203 628 • CSNK1DPITX2 782 1911 992 1850 1613 • CSNK1DFBXW4 787 2387 1096 1415 2089 • CSNK2A1PITX2 796 1154 2421 1959 115 • CSNK1DFBXW2 798 2207 1203 2384 477 • CSNK1DFZD6 800 2060 846 1138 1356 • CSNK2A1PPP2CA 801 423 350 2410 132 • APCCSNK1D 808 2236 514 1814 630 • CSNK1A1JUN 832 321 1169 1777 1201 • BTRCCSNK1A1 846 544 2355 769 2331 • CSNK2A1NLK 848 668 1601 1918 834 • CSNK2A1WNT3A 855 1073 1724 2483 1352 • CSNK1DDAAM1 865 1565 902 1575 167 • CSNK1A1FOXN1 871 624 324 1458 462 • CSNK2A1WNT2 872 1619 2226 1598 1005 • CSNK2A1GSK3B 875 1115 1899 1701 1128 • CSNK2A1WNT4 890 833 2340 1904 1760 • CSNK2A1TCF7 896 324 1552 96 1506 • CSNK2A1PPP2R1A 925 1399 1152 192 631 • CSNK1DWNT2 928 265 1934 204 389 • CSNK1DWNT4 931 1219 940 1605 2099 • CSNK1G1WNT1 946 2370 448 2034 829 • CSNK1DCXXC4 948 688 1059 430 921 • CSNK1DKREMEN1 952 1723 1481 2099 296 • CSNK2A1WNT2B 954 245 954 1852 791 • CSNK1A1PORCN 962 1055 1954 1314 39 • CSNK1G1CTNNBIP1 992 1044 2454 1412 1584 • APCCSNK2A1 1012 2069 723 1914 947 • CSNK1DNLK 1018 2142 921 1903 730 • CSNK1DPPP2R1A 1023 2376 2059 2251 874 • CSNK2A1WNT5A 1024 2374 1580 830 470 • AESCSNK1A1 1025 473 834 816 1909 • CSNK2A1FBXW11 1045 480 849 1377 1270 • CSNK1G1PORCN 1047 1269 1229 2233 698 • CSNK1A1FZD2 1052 1903 1799 435 140 • CSNK1G1LRP6 1055 1146 768 1075 1634 • CSNK2A1LRP5 1057 739 749 1208 678 • CSNK1DWNT5A 1062 2396 611 1794 986 • CSNK1DTCF7 1089 1141 1171 2462 598 • CSNK1G1CTBP2 1096 2302 1432 1979 2054 • FZD5CSNK1D 1108 535 462 1402 946 • CSNK1DDIXDC1 1114 986 1774 866 1627 • CSNK1DLRP5 1116 1902 1792 2466 1667 • CSNK1A1FGF4 1123 386 1581 2142 32 • CSNK2A1DIXDC1 1126 413 1316 1947 58 • CSNK1G1CTNNB1 1128 2228 2386 1001 1297 • CSNK1G1DVL2 1137 815 1654 1678 2291 • AXIN1CSNK1G1 1141 892 1008 354 882 • CSNK1A1PYGO1 1145 366 1753 63 179 • CSNK1DGSK3B 1160 2452 2448 2268 1830 • CSNK1A1MYC 1163 1566 634 1569 177 • CSNK1DWNT3A 1166 1179 2161 2402 2009 • CSNK2A1TLE2 1168 865 2368 1782 2061 • CSNK1DFRAT1 1181 1175 1017 616 1959 • CSNK1DTLE2 1183 2402 2131 591 1755 • CSNK2A1RHOU 1187 1542 1484 2443 819 • CSNK2A1FRAT1 1190 802 2155 1453 837 • CSNK1DFOSL1 1233 1920 2451 1558 1787 • CSNK2A1FZD1 1234 41 1051 2141 736 • CSNK2A1FOSL1 1240 648 1392 541 1441 • CSNK1DWNT2B 1241 1254 628 1737 1467 • CSNK1A1WNT3 1248 133 1215 492 66 • FZD5CSNK2A1 1249 1463 397 301 2028 • CSNK1G1T 1289 1771 1365 1872 593 • CSNK1DFBXW11 1311 1870 777 1384 803 • CSNK1A1SFRP1 1321 236 1421 1504 88 • CSNK1G1FBXW4 1346 2371 1612 1422 2179 • CSNK1G1NKD1 1353 2337 607 2136 2358 • BTRCCSNK1G1 1372 1672 1784 1653 1333 • CSNK1DRHOU 1379 2235 2319 1766 1814 • CSNK2A1TCF7L1 1388 487 1758 1542 1883 • CSNK1DCSNK2A1 1391 1808 1912 1533 2480 • CSNK2A1FZD7 1401 330 2183 2079 753 • CSNK1A1WNT1 1407 463 603 564 146 • CSNK1DFZD1 1419 2048 665 1345 469 • CSNK1A1WIF1 1420 1702 2263 1700 263 • CSNK1G1TLE1 1433 2421 1551 1277 1177 • CSNK1DFZD7 1462 2135 2193 998 843 • CCND3CSNK1D 1470 1437 594 2474 2122 • CCND1CSNK1A1 1480 994 546 1800 281 • CSNK1A1LEF1 1498 1110 1850 1487 775 • CSNK1DTCF7L1 1510 2074 2257 481 1186 • CSNK1A1T 1518 320 1834 389 633 • CSNK1G1EP300 1521 1360 1339 1785 2386 • CCND3CSNK2A1 1532 1436 1800 2480 1522 • CSNK1A1CTNNBIP1 1549 2303 1015 652 2236 • CSNK1A1LRP6 1557 212 1134 632 467 • CSNK1G1FZD8 1564 2414 2021 1248 2438 • BCL9CSNK1A1 1578 1735 942 1732 1184 • CSNK1G1NLK 1589 2174 2209 116 1196 • CCND2CSNK1G1 1594 947 523 954 2229 • CSNK1A1NKD1 1617 1037 707 20 905 • CSNK1G1CTBP1 1622 1952 1490 1266 1765 • CSNK1G1PITX2 1631 599 1796 2088 1461 • CSNK1G1DAAM1 1642 2062 2457 540 757 • CSNK1A1CTBP1 1645 1243 1897 2365 1533 • CSNK1G1SFRP4 1646 1536 2182 90 531 • CSNK1G1FZD6 1654 2203 1071 446 2437 • CSNK1A1DVL2 1657 1926 838 850 766 • CSNK1A1DVL1 1662 2460 2445 893 662 • CSNK1G1FSHB 1676 676 1859 1896 925 • CSNK1G1FBXW11 1687 2289 1332 181 2256 • CSNK1G1GSK3B 1712 2466 1919 1793 2138 • AESCSNK1G1 1715 1221 552 2274 1160 • CSNK1A1CTBP2 1743 720 1722 2230 519 • CSNK1G1FBXW2 1760 1176 1231 2470 2008 • CSNK1A1NLK 1779 2449 532 1171 117 • BCL9CSNK1G1 1787 924 301 1978 439 • CSNK1G1WNT2 1801 298 2024 440 1754 • CSNK1G1LRP5 1803 436 2440 1317 1757 • CSNK1G1WNT3A 1818 1737 2035 1000 2483 • CSNK1G1KREMEN1 1824 1978 1045 1457 1181 • CSNK1A1CSNK1G1 1825 152 1239 1224 305 • CSNK1DDKK1 1843 1698 884 770 1346 • CSNK1A1FSHB 1846 697 227 1674 1512 • CSNK1G1SLC9A3R1 1848 1366 1648 873 1388 • CSNK1G1PPP2R1A 1849 2097 667 511 235 • AESCSNK1D 1851 1193 1278 2036 1569 • CSNK1G1TCF7 1856 1879 1760 1698 1988 • CSNK1G1WNT2B 1863 2086 977 295 2031 • CSNK1G1TLE2 1874 806 1997 2261 1417 • CSNK1G1PPP2CA 1883 2254 1420 2033 1730 • CSNK1G1WNT4 1888 988 1445 2041 2287 • CSNK1A1TLE1 1896 2238 1737 1397 307 • CSNK1G1FRAT1 1900 2023 2484 1024 400 • CSNK1A1CTNNB1 1907 965 724 1508 1250 • CSNK1A1FBXW4 1910 612 949 1336 236 • CSNK1A1PITX2 1922 1721 2316 1856 189 • CSNK1G1DIXDC1 1925 811 1672 1424 2225 • CSNK1G1WNT5A 1929 2413 2042 2091 668 • CCND1CSNK1G1 1951 1991 1021 1807 644 • CSNK1G1FOSL1 1962 1860 1651 513 984 • CSNK1G1CSNK2A1 1963 1806 1266 700 1000 • CSNK1G1CXXC4 1965 322 2318 1921 569 • CCND2CSNK2A1 1969 2326 833 2344 1165 • CSNK1G1FZD7 1977 1469 1949 2304 2419 • CSNK1A1WNT2 1986 2382 750 612 297 • CSNK1A1TCF7 1988 2185 646 2178 45 • AESCSNK2A1 1989 554 1143 2264 2167 • CSNK1G1RHOU 2000 1041 2212 531 835 • CSNK1G1FZD1 2005 868 1744 2068 1620 • CSNK1A1CSNK2A1 2011 2179 1039 1037 552 • AXIN1CSNK1D 2012 428 1128 252 926 • CSNK1A1EP300 2020 2231 1906 2224 1231 • CSNK1A1PPP2CA 2024 1694 254 759 750 • CSNK1A1FZD6 2041 2101 453 607 655 • CSNK1A1DAAM1 2053 1666 1511 2007 73 • CSNK2A1DKK1 2056 1813 1147 1268 460 • AXIN1CSNK2A1 2069 888 1233 1238 1822 • CSNK1A1TLE2 2087 1804 1201 432 448 • CSNK1G1TCF7L1 2097 2094 1978 535 1446 • CSNK1A1FZD8 2110 1521 2198 372 329 • CSNK1A1SFRP4 2111 1733 1076 1030 240 • CSNK1A1CXXC4 2112 767 1530 1556 103 • CCND2CSNK1D 2126 781 982 2434 1955 • CSNK2A1FRZB 2127 859 1805 891 1483 • BTRCCSNK1D 2141 1773 2079 2051 2157 • CSNK1A1WNT2B 2166 1664 605 2001 91 • CSNK1A1PPP2R1A 2195 842 1325 1845 118 • BCL9CSNK2A1 2198 2237 950 2416 2015 • CSNK1A1FBXW2 2206 715 147 2390 69 • BTRCCSNK2A1 2222 1841 617 2082 1844 • CSNK1A1FOSL1 2226 6 1804 406 253 • CSNK1A1KREMEN1 2227 17 622 320 238 • CCND1CSNK2A1 2248 18 1442 1690 920 • CSNK1A1DIXDC1 2250 1734 1767 426 168 • CSNK1A1RHOU 2254 2227 778 501 430 • CSNK1A1WNT5A 2258 1642 1162 2450 220 • BCL9CSNK1D 2281 1823 1453 1126 2433 • CSNK1A1WNT4 2284 1378 1999 1440 1789 • CSNK1A1SLC9A3R1 2291 154 1072 316 955 • CSNK1A1GSK3B 2302 2005 1321 575 816 • CSNK1A1TCF7L1 2312 941 1272 2266 1593 • CSNK1A1FZD1 2316 1258 678 1429 2296 • CSNK1A1LRP5 2341 992 1389 1475 579 • CSNK1A1CSNK1D 2344 235 1188 2310 1548 • CSNK1A1FBXW11 2347 85 709 452 324 • CSNK1A1WNT3A 2364 1562 1765 1428 126 • CSNK1A1FRAT1 2367 1006 2286 2453 972 • CSNK1DFRZB 2386 1933 2394 1299 2183 • CSNK1G1FRZB 2397 231 2222 1939 707 • CCND1CSNK1D 2400 1450 1362 1941 1325 • CSNK2A1SENP2 2409 420 1987 704 367 • CSNK1G1DKK1 2415 2322 847 66 2005 • CSNK1A1FZD7 2418 1644 1881 1942 588 • CSNK1DSENP2 2420 2222 1751 600 1138 • CSNK1A1DKK1 2427 2004 318 2179 349 • CSNK1G1SENP2 2450 2164 1406 147 1311 • CSNK1A1FRZB 2462 560 1884 167 325 • CSNK1A1SENP2 2482 989 2158 994 247

## CSNK family 2*^nd^* order interaction ranking at different durations

• CSNK1A1SENP2 1 1971 2108 98 • FZD5CSNK2A1 3 1099 646 1647 • FZD5CSNK1G1 4 2430 1410 2054 • CSNK1A1FZD7 17 1782 481 315 • CSNK1A1LEF1 18 1554 922 78 • CSNK1A1SFRP4 22 1465 2478 64 • CSNK1A1FBXW11 23 1773 1902 11 • CSNK1A1FZD2 28 1633 1220 681 • CSNK1A1SFRP1 29 2068 1414 412 • CSNK1DSENP2 36 1952 1363 133 • CSNK1A1FBXW2 38 703 1802 503 • CSNK1A1PITX2 41 1284 1271 1602 • CSNK1A1PPP2R1A 43 1484 2382 316 • CSNK1A1FZD1 46 1728 1901 1723 • CSNK1A1LRP5 47 847 1405 1992 • CSNK1A1LRP6 53 1817 1598 162 • CSNK1A1DAAM1 57 1676 1086 94 • CSNK1A1FGF4 59 1625 1965 234 • CSNK1A1DVL2 64 1767 1056 161 • CSNK1A1CSNK2A1 71 2118 843 295 • CSNK1A1FZD8 77 1175 740 1307 • CSNK1A1FRZB 80 2231 2485 1638 • CSNK1A1NLK 83 1190 899 733 • CSNK1A1CTBP2 94 2403 2329 586 • CSNK1A1PPP2CA 96 1063 1339 1596 • CSNK1A1TCF7L1 97 1169 2288 1506 • CCND2CSNK1G1 101 1271 965 2063 • CSNK1A1WNT1 104 2388 2169 1071 • CSNK1A1TLE2 105 598 1624 1838 • CSNK1A1WNT5A 121 1315 1937 250 • AESCSNK1G1 123 2086 911 156 • CSNK1A1FZD6 126 2195 807 59 • CSNK1DSFRP4 129 2425 1161 106 • CSNK1A1TLE1 130 1353 1227 84 • CSNK1A1WNT4 132 647 1320 1520 • CSNK1A1KREMEN1 133 1876 867 1730 • CSNK1A1GSK3B 147 1715 991 1042 • CSNK1A1SLC9A3R1 150 1509 1068 455 • CSNK1A1CXXC4 151 449 1661 352 • CSNK1A1WIF1 153 1118 1172 20 • AXIN1CSNK2A1 171 657 2004 2117 • CSNK1DLRP6 178 2422 2036 1245 • CSNK1A1DKK1 179 1283 1312 8 • CSNK1DCSNK2A1 182 828 1302 1350 • CSNK1DLEF1 183 1241 1782 1242 • CSNK1A1GSK3A 193 2094 680 308 • CCND1CSNK1G1 218 1708 595 2066 • BCL9CSNK1G1 230 2247 1677 1582 • CSNK1A1NKD1 232 1630 713 614 • CSNK1A1EP300 249 511 2314 51 • CSNK1A1FRAT1 251 1050 1935 1921 • CSNK1A1FOXN1 255 1457 1786 1352 • CSNK1A1DIXDC1 256 1532 756 243 • CSNK1A1FBXW4 259 1108 1636 369 • CSNK1DFGF4 283 1220 1489 211 • CSNK1A1TCF7 286 864 818 1113 • AXIN1CSNK1G1 305 1001 1049 1873 • BCL9CSNK2A1 308 1231 860 1064 • CSNK1DCSNK1G1 319 777 2005 1828 • CCND1CSNK2A1 331 1085 1003 2220 • CCND2CSNK2A1 338 1607 1087 2365 • AESCSNK2A1 339 1702 2257 548 • CSNK1A1DVL1 349 1115 1140 14 • CSNK1DFZD2 369 713 2070 2278 • CSNK1DPPP2R1A 370 551 2152 1035 • CSNK1DFZD7 375 1402 1158 304 • CSNK1A1WNT3A 377 1466 618 1354 • CSNK1DPITX2 379 1885 1169 1840 • CSNK1DSFRP1 393 1904 1904 566 • CSNK1A1CSNK1D 396 1557 1748 1376 • APCCSNK1G1 407 1533 2295 2081 • BTRCCSNK2A1 418 707 1108 2421 • CSNK1DTCF7L1 435 1239 1609 851 • CSNK1DFZD1 455 1799 2017 2459 • CSNK1DWNT5A 459 321 1554 2342 • CSNK1DGSK3B 461 1046 1145 1236 • CSNK1DFBXW11 475 359 866 1396 • CSNK1DFBXW2 476 614 2148 477 • FZD5CSNK1D 479 1258 2066 1283 • BTRCCSNK1G1 486 2324 1504 891 • CSNK1DDAAM1 504 608 2116 18 • CSNK1DCXXC4 523 205 816 178 • CSNK1DFZD6 527 879 1440 341 • CSNK1A1WNT2 530 1102 1682 933 • APCCSNK2A1 563 2142 670 2441 • CSNK1DFZD8 574 1600 1580 830 • CSNK1DNLK 575 258 452 1458 • CSNK1DTLE2 619 336 2442 2173 • CSNK2A1SENP2 627 2257 51 508 • CSNK1DFBXW4 642 324 2008 878 • CSNK1A1FOSL1 643 417 2401 1600 • CSNK1A1MYC 649 942 775 355 • CSNK1A1FSHB 650 2289 2124 100 • CSNK1DDKK1 651 2372 211 1806 • CSNK1DTCF7 685 571 45 1780 • CSNK1DFRZB 730 329 1534 2192 • CSNK1A1CTNNBIP1 754 546 1379 47 • CSNK1DPPP2CA 755 680 333 1508 • CCND3CSNK1G1 782 1996 600 2456 • CSNK1DTLE1 799 983 2430 1224 • CSNK1A1T 812 2469 2182 2091 • CSNK1DWNT4 826 652 2258 1393 • CSNK1G1SENP2 837 911 393 1058 • CSNK1DNKD1 854 339 1656 1419 • CSNK1DLRP5 881 1849 2141 1881 • CCND3CSNK2A1 907 2390 1999 1752 • CSNK2A1SFRP4 920 1753 89 336 • CSNK1A1JUN 959 1599 2466 1950 • CSNK1A1WNT2B 965 1719 1267 1 • CSNK1A1RHOU 979 1008 1539 2275 • CSNK2A1LRP6 1026 1470 98 2088 • CSNK1DDVL2 1034 393 2341 1288 • CSNK1DEP300 1042 112 1944 27 • CSNK1A1CTNNB1 1047 583 1820 252 • AXIN1CSNK1D 1049 1610 762 344 • CSNK1DFOSL1 1064 1413 2460 1440 • CSNK1DKREMEN1 1082 472 1561 2075 • CSNK1G1SFRP4 1106 1420 1176 482 • CSNK1DCTBP2 1146 668 1464 746 • CCND2CSNK1D 1153 1545 2331 1065 • CSNK2A1FZD7 1155 2055 1770 1999 • CSNK2A1LEF1 1158 2275 800 991 • CSNK2A1SFRP1 1192 1386 978 1074 • CSNK1DDVL1 1210 223 1767 76 • CSNK1DWNT2 1214 743 1921 269 • CSNK1DFSHB 1221 2012 994 96 • CSNK1G1WNT5A 1226 43 2095 2273 • CSNK1A1CTBP1 1246 1429 902 1161 • CSNK1DGSK3A 1263 1943 1524 202 • CSNK1DSLC9A3R1 1285 955 1614 385 • BCL9CSNK1D 1288 2470 2263 1788 • CSNK2A1PPP2R1A 1305 674 221 2299 • AESCSNK1D 1332 2377 1540 1364 • CSNK1DMYC 1333 509 1450 768 • CSNK1DWIF1 1339 817 1007 109 • CCND1CSNK1D 1344 910 711 1586 • CSNK1A1WNT3 1353 1657 612 299 • CSNK1A1PYGO1 1363 2212 1835 2401 • CSNK1DFRAT1 1384 672 2240 1789 • CSNK2A1GSK3B 1391 2110 204 2258 • CSNK1G1LRP6 1393 854 491 1903 • CSNK2A1FBXW2 1394 1844 202 2410 • CSNK2A1WNT5A 1398 1041 399 1917 • CSNK2A1FGF4 1399 2294 746 881 • CSNK2A1FZD1 1401 1825 635 2361 • CSNK1DWNT2B 1416 1031 2243 166 • CSNK2A1TCF7L1 1437 2465 190 1536 • CSNK1G1FGF4 1438 771 2389 995 • BTRCCSNK1D 1446 1622 1821 1763 • CSNK1DCTBP1 1454 104 1129 775 • CSNK2A1NLK 1472 1449 29 2287 • CSNK2A1FZD2 1474 1941 1292 2005 • CSNK1G1LEF1 1480 1973 1465 2428 • CSNK1DWNT1 1484 230 2083 2108 • CSNK1G1PPP2R1A 1488 197 1326 1100 • CSNK1DCTNNBIP1 1509 233 1130 153 • CSNK2A1PPP2CA 1518 619 267 2368 • CSNK1DJUN 1522 271 2419 2328 • CSNK2A1PITX2 1523 1649 1674 1862 • CSNK1G1FBXW11 1531 73 160 2357 • FZD5CSNK1A1 1542 554 996 1886 • CSNK1G1TCF7L1 1565 1183 1483 2156 • CSNK1G1FZD1 1566 1748 2400 2170 • CSNK1G1FZD2 1571 451 1774 2404 • CSNK2A1FZD8 1579 2037 1358 2211 • CSNK1G1FZD8 1594 1744 1112 2264 • CSNK1DWNT3A 1599 822 1032 2186 • CSNK1DCTNNB1 1616 23 1430 1117 • CSNK2A1FRZB 1627 606 296 1080 • CSNK2A1TLE2 1628 2440 335 1090 • CSNK1G1FZD6 1661 274 174 620 • CSNK1G1SFRP1 1663 2391 2305 2057 • CSNK1G1FZD7 1667 900 1146 1049 • BCL9CSNK1A1 1694 629 690 1183 • CSNK1G1NLK 1696 63 141 2085 • APCCSNK1D 1698 899 1960 1653 • CSNK1G1PITX2 1702 1448 1560 1974 • CSNK1DFOXN1 1703 503 1239 2067 • CSNK2A1FBXW11 1706 326 116 2242 • CSNK2A1DKK1 1733 2269 106 2234 • CSNK1DDIXDC1 1742 2290 2300 389 • CSNK2A1FZD6 1747 1724 691 971 • CSNK2A1WNT4 1750 1154 1349 2099 • CSNK1G1PPP2CA 1783 226 1618 1998 • CSNK2A1CXXC4 1787 77 25 555 • CSNK1G1FRZB 1809 121 1645 1060 • CSNK1G1WNT4 1811 94 1494 1530 • CSNK2A1DAAM1 1814 1765 592 1345 • CSNK2A1WNT2 1827 1033 604 2455 • CSNK1G1GSK3B 1831 1162 1272 504 • CSNK2A1TCF7 1832 1441 24 1462 • CSNK1G1FBXW2 1861 261 1641 2144 • CSNK1G1DKK1 1866 1746 1205 962 • CSNK2A1FBXW4 1869 967 462 999 • CSNK1G1CSNK2A1 1895 268 852 2473 • CSNK1G1LRP5 1897 2166 1470 1460 • CSNK2A1NKD1 1936 450 96 2316 • CSNK1G1CTBP1 1951 14 1441 2465 • CSNK1DT 1963 497 1424 297 • CSNK2A1LRP5 1964 2348 437 1617 • CSNK2A1WNT3A 1966 1268 2435 496 • CSNK2A1FSHB 1993 2036 345 683 • CCND3CSNK1D 1999 1095 521 1128 • CSNK1A1PORCN 2002 1439 1884 509 • CSNK2A1TLE1 2014 1734 554 1041 • CSNK1G1TCF7 2016 255 69 2411 • CSNK1G1NKD1 2022 110 540 922 • CSNK1G1FBXW4 2024 36 1118 2084 • AXIN1CSNK1A1 2031 386 17 633 • APCCSNK1A1 2042 1909 825 246 • CSNK1G1CXXC4 2044 54 928 958 • CSNK2A1WNT1 2065 918 240 1260 • CSNK2A1DVL2 2074 465 222 2408 • CSNK1G1DVL2 2093 106 1858 1451 • CSNK1DRHOU 2108 1237 1711 1809 • CSNK1G1WNT2 2109 403 2282 1356 • CSNK2A1SLC9A3R1 2118 989 517 697 • CSNK1G1DVL1 2124 116 1698 266 • CSNK2A1FRAT1 2128 2186 325 615 • CSNK2A1WIF1 2147 2010 315 1225 • CSNK1G1DAAM1 2152 254 1033 781 • CSNK2A1EP300 2153 129 339 814 • CSNK2A1DVL1 2157 671 1174 222 • CSNK1G1EP300 2159 6 2447 1971 • CSNK1G1DIXDC1 2160 2361 1945 776 • CCND1CSNK1A1 2161 1178 143 1544 • CSNK2A1FOXN1 2162 194 303 742 • CSNK1G1WNT2B 2174 972 930 596 • CSNK1G1TLE2 2175 152 1378 131 • CSNK1G1CTBP2 2184 242 934 1964 • CSNK2A1WNT2B 2185 1900 83 748 • CSNK1G1WIF1 2189 658 2418 1298 • CSNK1DWNT3 2191 1636 274 500 • CSNK1DPYGO1 2208 397 2398 2140 • CSNK1G1RHOU 2229 484 2001 463 • CSNK2A1GSK3A 2233 1998 951 1370 • BTRCCSNK1A1 2259 973 726 1019 • CSNK1G1WNT1 2272 132 1495 2233 • CSNK2A1CTBP2 2276 2203 176 1359 • CSNK2A1MYC 2282 1995 115 2438 • CSNK1G1T 2288 319 2302 887 • AESCSNK1A1 2289 516 233 1993 • CSNK2A1RHOU 2292 2471 669 806 • CSNK1G1FOSL1 2293 1939 1078 2174 • CSNK1G1SLC9A3R1 2295 295 1709 1108 • CSNK1DPORCN 2296 447 2386 1606 • CSNK1G1KREMEN1 2304 297 1548 2059 • CSNK2A1KREMEN1 2305 988 683 1787 • CSNK1G1GSK3A 2330 852 1284 174 • CSNK2A1WNT3 2348 2151 127 2090 • CSNK1G1CTNNB1 2354 1 2099 824 • CSNK2A1CTNNBIP1 2356 190 851 1388 • CCND2CSNK1A1 2359 729 483 1131 • CSNK1G1FSHB 2364 952 2477 1114 • CSNK2A1FOSL1 2367 763 273 2202 • CSNK1G1FOXN1 2369 372 972 2116 • CSNK2A1JUN 2371 651 332 911 • CSNK1G1FRAT1 2373 355 710 2061 • CSNK2A1T 2392 1226 142 1521 • CSNK1G1PORCN 2398 251 1978 1687 • CSNK2A1PORCN 2403 1104 302 2403 • CSNK1G1JUN 2408 304 1459 2433 • CSNK1G1PYGO1 2420 103 1683 2461 • CSNK2A1CTBP1 2432 492 118 2346 • CSNK2A1CTNNB1 2434 314 610 1711 • CSNK2A1DIXDC1 2440 1388 1376 554 • CSNK1G1WNT3A 2451 632 1291 2485 • CSNK1G1TLE1 2457 471 1132 1909 • CSNK2A1PYGO1 2470 2316 463 1258 • CSNK1G1MYC 2473 370 484 638 • CSNK1G1CTNNBIP1 2481 185 1545 618 • CCND3CSNK1A1 2483 784 324 427 • CSNK1G1WNT3 2484 2446 225 1872

## DVL family 2*^nd^* order interaction ranking at different time points

• DKK1DVL1 36 2426 2175 75 1642 • DKK1DVL2 96 2214 2220 1654 1932 • CSNK2A1DVL1 209 1585 2353 1363 1798 • APCDVL1 239 2352 1209 203 1920 • CSNK2A1DVL2 256 369 910 1809 2254 • FZD5DVL1 302 1491 1084 852 396 • APCDVL2 307 2041 994 189 1402 • DIXDC1DVL1 328 2055 176 1035 671 • CSNK1DDVL1 341 1955 1163 25 1921 • DVL2GSK3A 353 1898 1103 74 987 • DIXDC1DVL2 421 491 72 1144 1286 • CXXC4DVL1 438 2280 1280 1239 1312 • DVL2JUN 440 486 1575 179 1845 • CSNK1DDVL2 463 1400 2348 1808 1158 • CXXC4DVL2 487 1004 1342 794 406 • FZD5DVL2 496 675 889 334 2223 • CTBP1DVL1 505 2071 1305 1593 221 • CCND3DVL1 518 1028 1931 288 2010 • DVL1GSK3A 529 2072 131 1370 2367 • DVL2FOXN1 532 1451 372 2378 1828 • DVL2FGF4 536 797 2255 1082 642 • DVL1SFRP1 540 54 1646 551 616 • DVL2PORCN 544 167 791 592 99 • DVL1FOXN1 566 196 130 91 2098 • CTNNBIP1DVL1 571 2450 1347 1104 76 • DVL2FZD2 574 367 1449 642 1431 • DAAM1DVL1 606 1479 1456 1949 1318 • DVL2LEF1 616 283 2132 895 1110 • DVL2WIF1 619 408 892 308 372 • CCND3DVL2 621 104 2358 2223 1751 • DVL2MYC 622 625 739 2421 2302 • DVL1FGF4 633 1184 2150 1149 1117 • DAAM1DVL2 638 1112 1597 275 2304 • DVL1JUN 644 46 2029 1394 844 • DVL2WNT3 654 858 2395 1288 420 • DVL1WNT1 672 393 239 191 756 • DVL2PYGO1 673 23 1297 2193 647 • CTBP1DVL2 697 2343 537 847 1393 • DVL1PYGO1 730 201 335 77 1131 • CSNK1G1DVL1 731 1263 1218 363 806 • DVL2WNT1 734 776 606 594 423 • CTBP2DVL1 743 903 1334 1411 1095 • CTNNB1DVL1 756 2353 1769 123 1645 • DVL2SFRP1 793 405 963 329 535 • CTNNB1DVL2 835 2448 674 737 1508 • DVL1WIF1 876 1034 1444 1753 990 • DVL1LEF1 906 398 752 941 261 • CTNNBIP1DVL2 929 2252 732 1925 1012 • DVL1WNT3 982 356 1398 1133 562 • AESDVL1 997 438 2201 175 1178 • CCND2DVL1 1013 821 1822 1143 2046 • AXIN1DVL1 1042 2472 1812 751 1105 • DVL1FZD2 1084 1025 1010 44 1733 • DVL1MYC 1094 199 380 933 1986 • CTBP2DVL2 1102 505 1438 1756 1779 • DVL1PORCN 1105 726 525 294 68 • CSNK1G1DVL2 1137 815 1654 1678 2291 • DVL2LRP6 1158 454 629 1462 933 • DVL2T 1159 735 853 953 692 • DVL2NKD1 1251 1717 174 529 2232 • AESDVL2 1282 779 2189 2469 1839 • AXIN1DVL2 1307 1493 1264 242 1425 • DVL2TLE1 1415 621 1556 472 2163 • BTRCDVL1 1424 749 2190 99 983 • DVL1T 1454 1548 643 1283 1666 • DVL2SFRP4 1483 1464 1656 1251 1188 • DVL1FBXW4 1503 157 207 2177 1837 • CCND1DVL1 1512 29 1947 174 1150 • DVL1LRP6 1515 1132 289 81 443 • BCL9DVL1 1551 2000 2283 1032 1860 • DVL2PPP2CA 1561 2052 365 1741 295 • DVL1DVL2 1583 1768 164 7 1563 • BTRCDVL2 1590 1309 1200 2359 2378 • DVL2FZD6 1598 1876 627 345 627 • DVL2KREMEN1 1601 2002 1968 1697 1694 • DVL2FZD8 1616 1303 1564 1900 675 • DVL2PITX2 1618 1268 1268 517 1716 • DVL1NKD1 1624 693 382 582 690 • CSNK1A1DVL2 1657 1926 838 850 766 • DVL2FBXW2 1659 908 1104 2239 786 • CSNK1A1DVL1 1662 2460 2445 893 662 • DVL2SLC9A3R1 1667 920 2390 634 1688 • DVL1FZD6 1675 1468 270 1935 703 • DVL2FBXW4 1681 945 569 693 1604 • DVL1FSHB 1684 1018 217 321 2436 • CCND2DVL2 1686 373 1488 874 889 • BCL9DVL2 1689 1667 649 71 2060 • DVL2NLK 1703 1094 1871 1334 650 • DVL1TLE1 1713 1127 770 1275 1371 • DVL1SLC9A3R1 1714 927 1898 1309 331 • DVL2FSHB 1732 110 602 1161 2330 • DVL2TCF7 1734 1838 1090 1468 151 • DVL1WNT2 1752 1472 2235 1521 862 • DVL2WNT2 1759 1831 2419 590 2464 • DVL1NLK 1761 728 126 570 980 • DVL1PITX2 1775 1680 703 169 1314 • DVL2PPP2R1A 1777 1353 830 144 333 • DVL1TCF7 1780 2282 165 1096 2050 • DVL1KREMEN1 1788 171 989 496 109 • DVL1PPP2R1A 1793 143 804 650 272 • DVL1WNT5A 1811 917 522 775 317 • DVL2WNT4 1816 1395 737 1351 2351 • DVL2WNT5A 1820 1069 839 1342 360 • DVL1FZD8 1828 1406 894 1074 368 • DVL2LRP5 1830 2243 1300 1328 718 • DVL1LRP5 1845 632 244 965 1911 • DVL2GSK3B 1855 1533 1172 1986 1289 • DVL2EP300 1876 804 1361 225 1262 • CCND1DVL2 1877 754 821 518 1340 • DVL1SFRP4 1905 2085 2274 2119 1834 • DVL2FRAT1 1909 1003 1131 1305 1887 • DVL2WNT3A 1912 1636 860 1996 749 • DVL1PPP2CA 1923 1134 224 778 2067 • DVL1FBXW2 1944 478 267 504 2051 • DVL2RHOU 1955 615 2364 2454 767 • DVL2WNT2B 1959 737 765 1667 382 • DVL2TLE2 1971 714 764 33 1119 • DVL2FOSL1 1974 130 2142 143 2176 • DVL2FZD7 1975 445 1159 782 1288 • DVL1WNT3A 1999 1507 685 268 1552 • DVL1EP300 2031 68 786 210 1568 • DVL1WNT2B 2045 314 127 989 1104 • DVL1RHOU 2047 1888 1035 237 929 • DVL2TCF7L1 2098 103 1781 382 2121 • DVL1FOSL1 2109 425 1196 395 547 • DVL1FBXW11 2121 79 206 1361 632 • DVL2FZD1 2151 1762 1522 2159 1710 • DVL1FRAT1 2164 1593 1161 1413 1531 • DVL2FBXW11 2181 1392 373 1482 1223 • DVL1GSK3B 2209 123 1378 390 2445 • DVL1FZD1 2215 217 1155 1106 2275 • DVL1FZD7 2240 596 2234 546 770 • DVL1WNT4 2252 969 763 2413 1332 • DVL1TCF7L1 2298 1 1306 1930 2016 • DVL1TLE2 2310 1369 219 2 1648 • DVL1FRZB 2414 1080 1528 1465 956 • DVL2FRZB 2416 1342 1545 1206 1164 • DVL1SENP2 2463 1087 1092 756 1560 • DVL2SENP2 2467 502 1920 51 998

## DVL family 2*^nd^* order interaction ranking at different durations

• CSNK1A1DVL2 64 1767 1056 161 • DVL1SENP2 98 1004 94 2405 • DVL1SFRP4 139 2096 336 815 • DVL2SENP2 167 2190 236 570 • FZD5DVL2 273 884 1602 2092 • DVL1LRP6 291 2342 613 1496 • DIXDC1DVL2 332 22 2072 1010 • CSNK1A1DVL1 349 1115 1140 14 • DVL2SFRP4 389 2219 342 797 • FZD5DVL1 394 519 1754 757 • DVL1FZD2 401 1384 2105 368 • DVL2LRP6 414 2343 580 2390 • DVL1PPP2R1A 482 875 287 2103 • DVL1FBXW2 500 1199 101 1826 • DVL1FBXW11 509 2454 134 1697 • CTNNB1DVL1 521 2265 2414 119 • DVL1LEF1 525 1727 1766 666 • DVL1SFRP1 534 1486 1549 2079 • DVL1FZD1 536 833 365 478 • CCND2DVL2 546 2129 1402 2373 • DVL1TCF7L1 551 1938 1080 1023 • DVL1FZD7 553 1319 1866 725 • DVL1FGF4 557 688 1289 635 • CTNNB1DVL2 569 791 1874 1274 • DVL2SFRP1 585 1866 926 1739 • DVL1GSK3B 589 1158 2048 107 • AXIN1DVL2 595 232 1694 2378 • DVL1WNT5A 621 1645 414 1031 • DVL2FGF4 647 1438 2197 1975 • DVL1FZD6 656 2292 1875 1835 • DVL2FZD7 668 1234 1558 1896 • CTBP1DVL2 671 981 2176 2380 • DVL1PITX2 693 1674 998 758 • DVL2FZD2 705 2448 2088 1209 • DVL1FZD8 706 1259 2474 1084 • DVL2LEF1 709 1195 1764 1997 • DVL2FZD1 716 2344 764 1635 • DVL2PPP2R1A 756 2169 1396 2360 • CTBP1DVL1 760 1255 745 473 • AESDVL2 765 2385 2404 31 • CCND1DVL2 784 759 513 2147 • DVL2FBXW2 811 2297 1179 1703 • DVL1TLE2 819 1932 575 200 • DVL1PPP2CA 820 1888 482 1018 • DVL2FBXW11 842 1955 125 1822 • DVL1WNT4 851 700 1149 335 • DIXDC1DVL1 861 13 1837 1321 • CTNNBIP1DVL2 862 2415 1360 2182 • DVL2TCF7L1 867 1191 716 2293 • DVL1DVL2 869 1354 2363 1538 • DVL1NLK 873 565 46 1129 • DVL2PITX2 884 1703 2281 1608 • DVL2WNT5A 897 2254 614 2395 • DVL1TCF7 905 1899 37 1262 • BCL9DVL2 910 1047 1987 2358 • BTRCDVL2 932 247 1384 1323 • DVL2FZD8 947 2072 1486 2389 • DVL2PPP2CA 955 841 1096 2248 • DVL1TLE1 963 1316 1113 2026 • DVL2FZD6 964 2383 1752 550 • DVL2GSK3B 981 2135 494 1079 • CCND2DVL1 1019 691 1001 498 • CSNK1DDVL2 1034 393 2341 1288 • DVL1FRZB 1040 1149 128 258 • DVL1LRP5 1056 1893 1097 92 • BTRCDVL1 1065 1229 1678 137 • APCDVL2 1081 2402 1596 896 • DVL2NLK 1099 1982 77 1564 • CCND1DVL1 1101 1117 2192 745 • DVL1NKD1 1103 861 86 755 • AXIN1DVL1 1130 520 1006 1326 • DVL1FBXW4 1183 928 1839 1685 • BCL9DVL1 1187 809 1412 611 • CSNK1DDVL1 1210 223 1767 76 • CTBP2DVL2 1245 459 2472 2451 • DVL2FBXW4 1249 1578 2356 1232 • DVL2TLE2 1272 1379 2241 170 • DVL1EP300 1293 1186 1745 294 • AESDVL1 1307 618 2309 36 • DVL2LRP5 1312 1738 1791 1253 • DVL1MYC 1321 789 241 943 • DVL1WIF1 1343 2150 1183 587 • DVL2TCF7 1354 2270 36 2475 • DVL1FOSL1 1367 2253 561 613 • DVL2FRZB 1373 838 495 237 • DVL2FOSL1 1381 1391 1514 1613 • CTBP2DVL1 1385 550 1538 459 • CCND3DVL2 1411 1771 1359 2356 • DVL1SLC9A3R1 1434 2126 1387 1640 • DVL2WNT2 1444 1024 1211 1425 • DVL2EP300 1449 320 1194 2021 • DVL1WNT2 1450 2429 1323 288 • DVL2NKD1 1457 901 232 2025 • DVL2WNT4 1461 1972 2269 1877 • DVL2SLC9A3R1 1502 1534 2244 1543 • DVL2TLE1 1507 1663 1855 1892 • CTNNBIP1DVL1 1530 770 2399 1748 • DVL1WNT1 1584 2331 432 608 • CXXC4DVL2 1600 1013 2228 1915 • DVL1FSHB 1610 1942 589 1429 • DKK1DVL2 1611 64 1815 392 • DVL1KREMEN1 1625 2313 1370 694 • DVL1GSK3A 1662 1851 2480 366 • APCDVL1 1699 2484 679 843 • DVL1T 1710 1070 229 259 • DAAM1DVL2 1737 1219 1735 1973 • CCND3DVL1 1759 1097 1109 948 • DVL2FSHB 1805 1279 1810 928 • DVL2T 1820 1415 881 786 • DVL2GSK3A 1839 2227 2307 1621 • DVL2WNT1 1873 2442 2007 1914 • DVL2WIF1 1881 1989 1640 2207 • DVL1WNT2B 1945 1922 360 2047 • DVL1FRAT1 1950 1994 289 430 • DVL1FOXN1 1952 1335 137 180 • DVL2MYC 1965 1121 248 2406 • DVL2FRAT1 1985 2196 1255 1963 • CXXC4DVL1 2007 2105 1337 1707 • DVL2KREMEN1 2011 987 2158 634 • DKK1DVL1 2013 150 1436 150 • DVL2FOXN1 2027 1424 767 2282 • DVL1WNT3A 2037 1242 2272 401 • DVL2RHOU 2049 2079 1322 386 • DVL2WNT3A 2050 1927 1344 2068 • DAAM1DVL1 2063 922 2154 858 • CSNK2A1DVL2 2074 465 222 2408 • CSNK1G1DVL2 2093 106 1858 1451 • DVL2JUN 2114 738 496 859 • DVL1JUN 2122 2485 290 196 • CSNK1G1DVL1 2124 116 1698 266 • CSNK2A1DVL1 2157 671 1174 222 • DVL1PORCN 2195 1153 650 321 • DVL1RHOU 2214 997 1411 188 • DVL2WNT3 2216 1491 81 630 • DVL2PYGO1 2255 2464 1397 2051 • DVL2PORCN 2261 1163 882 2437 • DVL1WNT3 2264 2326 33 1784 • DVL2WNT2B 2301 1775 535 418 • DVL1PYGO1 2455 2145 1035 747

## LRP family 2*^nd^* order interaction ranking at different time points

• FRZBLRP6 26 1067 123 2202 294 • DKK1LRP6 50 2347 2208 669 2353 • LRP5SFRP1 70 1751 2472 2201 601 • LRP5WNT1 92 2215 508 2133 863 • LRP5WNT3 127 1716 1011 2369 708 • FRZBLRP5 175 1511 309 1019 446 • LRP5MYC 193 2244 516 1739 1889 • LRP5PYGO1 213 1489 641 978 450 • APCLRP6 305 1753 934 905 689 • FRAT1LRP6 316 2224 616 1259 181 • CSNK2A1LRP6 325 1002 1245 1483 589 • FOSL1LRP6 326 2095 111 834 60 • DKK1LRP5 345 1339 1690 2288 782 • FZD7LRP6 360 1120 223 1920 571 • CSNK1DLRP6 361 722 1380 861 1616 • LRP5WIF1 363 2440 1312 682 2463 • LRP5PORCN 367 1016 1851 2485 149 • LRP5T 371 2049 1900 1049 556 • EP300LRP6 389 1329 1023 1963 1080 • FBXW11LRP6 395 540 1589 2330 1992 • FZD1LRP6 396 401 465 2398 607 • LRP5LRP6 415 189 920 2035 1881 • GSK3BLRP6 417 1620 533 1789 75 • LRP5NKD1 434 2307 981 878 1433 • FZD5LRP6 470 572 584 581 1064 • FZD8LRP6 475 1171 473 2017 1739 • CXXC4LRP6 477 2408 488 2279 1944 • DIXDC1LRP6 513 2038 15 2214 724 • LRP6PYGO1 521 455 1293 901 1927 • FBXW2LRP6 524 501 1001 2476 645 • LRP5TLE1 542 1594 2248 1301 807 • DAAM1LRP6 569 451 1078 896 1713 • CTBP1LRP6 577 1035 693 1874 487 • FZD6LRP6 582 976 2428 2447 2332 • LRP6WNT1 597 1916 1184 1684 584 • LRP6SFRP1 618 1431 2292 2236 1664 • KREMEN1LRP6 676 490 557 1100 528 • LRP5FBXW4 691 2170 794 1716 895 • CCND3LRP6 709 1526 2401 2373 875 • LRP6PORCN 724 1056 1149 186 319 • LRP5PITX2 725 1896 966 1719 999 • LRP5PPP2CA 733 2161 410 2286 27 • LRP5SFRP4 750 250 1791 2197 1036 • FSHBLRP6 762 1626 2215 1479 780 • LRP6WNT3 794 914 2466 1842 597 • LRP6WIF1 816 2266 2372 790 1515 • LRP6MYC 821 750 1055 2400 2432 • FZD1LRP5 850 472 1030 2101 1429 • APCLRP5 861 1079 1027 2277 496 • FOSL1LRP5 866 168 79 1292 1856 • LRP5SLC9A3R1 884 1152 2349 450 594 • LRP5NLK 904 2464 530 1258 565 • FZD7LRP5 943 1218 455 2374 2044 • FBXW11LRP5 949 1556 512 1200 2375 • LRP5PPP2R1A 961 2368 1644 677 380 • CTNNBIP1LRP6 967 2427 783 1141 222 • CTBP2LRP6 978 296 967 1199 510 • LRP5WNT3A 986 492 1258 2231 695 • LRP5TCF7 1026 2443 1579 113 340 • LRP5WNT5A 1040 2397 972 1967 289 • CSNK1G1LRP6 1055 1146 768 1075 1634 • LRP5WNT2 1056 1504 1803 223 959 • CSNK2A1LRP5 1057 739 749 1208 678 • BTRCLRP6 1078 910 1773 1573 2148 • LRP5TLE2 1092 1148 2423 1857 1718 • CSNK1DLRP5 1116 1902 1792 2466 1667 • FRAT1LRP5 1144 1050 814 2267 2102 • AESLRP6 1150 799 560 2420 922 • FZD5LRP5 1153 192 586 1943 1404 • DVL2LRP6 1158 454 629 1462 933 • CTNNB1LRP6 1170 1032 1036 1330 1434 • LRP5WNT4 1222 117 980 1057 2210 • LRP5WNT2B 1264 718 715 2456 1063 • LRP6T 1287 1571 1875 1821 1115 • GSK3BLRP5 1288 496 636 558 2128 • DIXDC1LRP5 1336 1029 25 610 337 • FZD8LRP5 1343 359 806 1481 1805 • CXXC4LRP5 1366 290 1223 1561 1169 • LRP5RHOU 1370 1239 1337 2381 2137 • CCND3LRP5 1373 1582 2403 1571 2006 • LEF1LRP6 1386 1053 1040 1564 107 • LRP6NKD1 1409 2473 486 1270 2118 • AXIN1LRP6 1428 2138 844 543 1567 • FBXW2LRP5 1437 997 1863 2075 1869 • KREMEN1LRP5 1444 1575 1243 2077 1282 • BCL9LRP6 1485 1054 408 2095 546 • CCND2LRP6 1495 1336 1275 1432 2216 • EP300LRP5 1497 2122 898 813 1815 • DVL1LRP6 1515 1132 289 81 443 • CTBP1LRP5 1531 446 449 463 1226 • LRP6FBXW4 1534 2334 1448 2256 1561 • FZD6LRP5 1545 522 2280 872 1618 • DAAM1LRP5 1547 403 2139 1837 1778 • CSNK1A1LRP6 1557 212 1134 632 467 • FOXN1LRP6 1571 1157 1599 714 1217 • FSHBLRP5 1600 410 2266 2455 557 • FGF4LRP6 1606 2200 172 1750 1412 • LRP6PITX2 1633 28 1379 527 1193 • LRP5TCF7L1 1691 1313 2302 1624 744 • LRP6SLC9A3R1 1698 1023 816 1306 2190 • JUNLRP6 1702 195 1477 2262 80 • FZD2LRP6 1725 75 599 133 401 • LRP6SFRP4 1735 1149 2141 1515 1684 • AXIN1LRP5 1740 628 396 56 2062 • LRP6WNT5A 1750 474 274 1446 943 • LRP6TLE1 1756 1212 1504 282 934 • CTNNB1LRP5 1762 588 1422 2216 833 • LRP6PPP2CA 1792 2445 575 2334 2084 • LRP6RHOU 1797 2253 2218 1375 1549 • CSNK1G1LRP5 1803 436 2440 1317 1757 • LRP6TCF7 1808 148 435 2484 2349 • CTNNBIP1LRP5 1819 358 459 1946 459 • DVL2LRP5 1830 2243 1300 1328 718 • DVL1LRP5 1845 632 244 965 1911 • LRP6WNT2 1875 305 1700 1357 1823 • LRP6NLK 1894 1539 1768 823 1980 • GSK3ALRP6 1901 653 817 1279 1852 • LRP6PPP2R1A 1924 116 134 1987 658 • LRP6WNT4 1926 829 1310 576 1952 • CTBP2LRP5 1927 1375 1073 2237 1652 • LRP6WNT2B 1937 1540 450 2227 885 • LRP6WNT3A 2007 1576 810 1177 2429 • LRP6TCF7L1 2037 955 1634 1396 428 • CCND1LRP6 2040 213 848 2107 1675 • LRP6TLE2 2064 389 528 913 127 • CCND2LRP5 2100 635 1585 787 2425 • AESLRP5 2101 816 1066 1634 1106 • FZD2LRP5 2144 1214 539 2015 172 • JUNLRP5 2183 711 511 1505 271 • LEF1LRP5 2190 457 1311 2428 1598 • BTRCLRP5 2197 613 1577 807 1615 • FGF4LRP5 2201 617 216 1981 1182 • BCL9LRP5 2317 1121 1199 2083 2075 • FOXN1LRP5 2325 1075 2045 883 1498 • CSNK1A1LRP5 2341 992 1389 1475 579 • CCND1LRP5 2358 1295 1123 2240 2106 • LRP5SENP2 2377 2301 2463 959 1894 • GSK3ALRP5 2437 881 2025 2172 993 • LRP6SENP2 2472 2268 877 4 659

## LRP family 2*^nd^* order interaction ranking at different durations

• FZD5LRP6 11 2235 1633 1593 • CSNK1A1LRP5 47 847 1405 1992 • CSNK1A1LRP6 53 1817 1598 162 • FZD5LRP5 62 2047 2279 658 • CTNNB1LRP6 65 1355 1570 1033 • DIXDC1LRP6 85 553 531 1924 • JUNLRP6 127 1970 2214 1322 • AXIN1LRP6 141 1869 1295 2249 • CCND2LRP6 142 1867 1089 1165 • CTBP1LRP6 144 1761 1537 2048 • CSNK1DLRP6 178 2422 2036 1245 • BCL9LRP6 186 1742 1832 930 • CCND1LRP6 196 2120 789 861 • LRP5SENP2 206 917 1530 167 • FRAT1LRP6 212 2458 1138 1616 • FSHBLRP6 214 1700 2217 1675 • KREMEN1LRP6 270 2242 731 1327 • AESLRP6 281 2208 1584 89 • DVL1LRP6 291 2342 613 1496 • CTNNBIP1LRP6 337 2043 385 2266 • LRP5SFRP4 353 1295 651 32 • FOXN1LRP6 357 1414 574 975 • CTBP1LRP5 404 1696 1606 1169 • DVL2LRP6 414 2343 580 2390 • BTRCLRP6 415 1655 1510 1103 • FOSL1LRP6 433 1422 803 1800 • GSK3ALRP6 438 1251 109 1690 • CTBP2LRP6 464 2076 1751 2354 • DIXDC1LRP5 481 1410 666 48 • APCLRP6 495 1624 1270 1708 • CTNNB1LRP5 540 1059 1038 1549 • JUNLRP5 598 1053 1817 625 • EP300LRP6 600 1582 461 2318 • CCND2LRP5 637 1710 1691 914 • CCND3LRP6 657 1637 1801 1284 • FRAT1LRP5 674 1865 875 1876 • LRP5PPP2R1A 689 60 1512 148 • FRZBLRP6 708 2084 2208 437 • LRP5LRP6 721 853 424 545 • LRP5SFRP1 759 792 2433 4 • LRP6SENP2 761 328 970 961 • FZD8LRP6 769 559 100 1434 • LRP5TCF7L1 778 1338 1381 1572 • DKK1LRP6 785 859 799 1334 • DAAM1LRP6 829 2252 508 1853 • CXXC4LRP6 843 1473 1918 1202 • LRP5WNT5A 844 208 1780 235 • AXIN1LRP5 848 2156 1997 372 • KREMEN1LRP5 857 1608 886 1416 • CSNK1DLRP5 881 1849 2141 1881 • LRP5NLK 882 45 701 263 • CCND1LRP5 896 1699 1454 1308 • LRP5PITX2 911 1312 1643 628 • FOXN1LRP5 914 1857 1759 2221 • CTNNBIP1LRP5 917 2127 430 260 • AESLRP5 926 1442 2000 1369 • GSK3BLRP6 938 750 67 224 • FZD6LRP6 948 1770 103 1889 • FSHBLRP5 952 1365 1371 90 • FBXW2LRP6 991 2386 2358 2167 • BCL9LRP5 1009 1999 1670 857 • CSNK2A1LRP6 1026 1470 98 2088 • FZD1LRP6 1048 1052 1727 1498 • DVL1LRP5 1056 1893 1097 92 • LRP5TLE2 1091 234 880 394 • GSK3ALRP5 1100 2100 719 195 • FOSL1LRP5 1142 746 1182 838 • LRP5TCF7 1164 130 392 306 • FBXW11LRP6 1166 2404 2237 2412 • FZD7LRP6 1168 1580 48 1918 • APCLRP5 1169 1547 2286 374 • BTRCLRP5 1178 1552 2179 1437 • LRP6SFRP4 1248 699 1730 475 • FGF4LRP6 1256 708 523 2215 • LRP5PPP2CA 1267 31 420 1882 • EP300LRP5 1277 1296 2432 242 • CTBP2LRP5 1287 1991 2454 572 • DVL2LRP5 1312 1738 1791 1253 • FRZBLRP5 1341 1693 1285 2463 • LRP5FBXW4 1368 140 2283 72 • LRP5WNT4 1372 93 1930 1936 • LRP5WNT2 1390 281 2453 531 • CSNK1G1LRP6 1393 854 491 1903 • LRP5TLE1 1413 621 1572 65 • FZD2LRP6 1425 1786 358 1962 • LEF1LRP6 1447 686 874 2008 • FZD6LRP5 1455 2426 266 61 • CXXC4LRP5 1562 1012 2193 612 • CCND3LRP5 1595 895 1137 367 • LRP6TCF7L1 1598 2125 2438 2450 • LRP6PPP2R1A 1607 469 706 1522 • LRP6SFRP1 1647 2049 2209 1483 • DAAM1LRP5 1652 1951 582 342 • LRP5WIF1 1671 790 2032 125 • LRP6WNT5A 1675 398 1103 2319 • DKK1LRP5 1727 1155 2339 1470 • LRP5NKD1 1730 34 244 212 • LRP5MYC 1738 174 185 60 • LRP6TCF7 1753 227 428 1980 • FBXW2LRP5 1757 1184 2450 2012 • LRP5WNT1 1758 84 846 1925 • FGF4LRP5 1795 1054 1958 967 • LRP6NLK 1824 243 164 2023 • FBXW11LRP5 1829 1690 1566 862 • FZD7LRP5 1860 1934 153 132 • FZD1LRP5 1864 1768 619 1043 • LRP5SLC9A3R1 1880 421 2427 701 • GSK3BLRP5 1885 804 186 2352 • LRP5T 1890 224 1788 1439 • LRP6TLE1 1893 769 791 818 • CSNK1G1LRP5 1897 2166 1470 1460 • LRP6PITX2 1959 2000 844 1719 • CSNK2A1LRP5 1964 2348 437 1617 • LEF1LRP5 1969 1503 2227 744 • FZD8LRP5 1971 1856 448 543 • LRP5PORCN 1984 290 2357 323 • LRP5RHOU 2003 494 1829 1930 • LRP6PPP2CA 2058 454 2390 2064 • LRP6WNT3 2098 1336 111 1219 • LRP6TLE2 2129 458 2377 209 • FZD2LRP5 2158 2327 810 1124 • LRP6SLC9A3R1 2167 532 1726 1212 • LRP6WNT4 2183 310 1369 1948 • LRP5WNT2B 2200 676 1251 26 • LRP6MYC 2228 409 1277 1706 • LRP6FBXW4 2242 160 777 1686 • LRP6PORCN 2265 296 1469 1667 • LRP6WIF1 2290 1197 2396 1412 • LRP6WNT2B 2309 947 802 598 • LRP5WNT3A 2311 375 2117 1541 • LRP6T 2336 156 2030 168 • LRP6NKD1 2357 29 2140 2369 • LRP6WNT2 2361 382 1338 1062 • LRP6PYGO1 2384 481 1190 1226 • LRP6WNT1 2394 200 2465 2185 • LRP6RHOU 2397 737 2044 592 • LRP6WNT3A 2424 610 665 2145 • LRP5WNT3 2430 2267 198 356 • LRP5PYGO1 2441 361 774 902

## FBXW family 2*^nd^* order interaction ranking at different time points

• SENP2FBXW4 34 308 225 1062 1175 • FBXW11GSK3A 44 334 938 111 2301 • FBXW11FGF4 46 824 1340 1194 1737 • FRZBFBXW4 68 2391 495 1781 171 • FBXW2JUN 110 1284 2462 545 1631 • FBXW11WIF1 117 510 1930 350 1724 • FBXW11FOXN1 120 1501 346 224 2249 • DKK1FBXW4 137 2358 1667 1492 2268 • FBXW11FZD2 144 353 808 187 2385 • FBXW11PYGO1 149 224 840 631 2346 • FBXW11JUN 156 332 1707 1135 2405 • FBXW11LEF1 160 568 1907 1517 2072 • FBXW11PORCN 178 63 1087 2181 637 • FBXW11MYC 192 979 687 684 1419 • FBXW11SFRP1 198 24 2408 2108 1265 • FBXW11WNT1 223 327 787 1926 1139 • FBXW2GSK3A 236 1675 375 1472 2146 • FBXW2FOXN1 269 531 558 1827 2134 • FBXW11WNT3 299 678 969 1600 1237 • FBXW2WNT1 308 2031 785 476 696 • FBXW11T 336 550 2163 1752 1999 • FBXW2PORCN 356 1763 2387 2110 1285 • FBXW2PYGO1 357 241 1492 1080 1715 • DKK1FBXW2 365 2065 1349 1007 1968 • FBXW2FZD2 376 40 1175 1125 1759 • FBXW2MYC 381 1630 1319 641 1292 • DKK1FBXW11 391 2423 1179 1124 2382 • FBXW11LRP6 395 540 1589 2330 1992 • FBXW2LEF1 398 681 2288 1974 2456 • FBXW2SFRP1 408 1144 738 1637 1436 • FBXW2WNT3 422 961 2381 1618 1994 • FBXW11TLE1 430 304 1816 1742 1435 • FBXW2T 485 228 1903 1163 901 • FBXW4WIF1 493 215 1748 1341 553 • RHOUFBXW4 502 2333 498 815 2068 • FBXW2LRP6 524 501 1001 2476 645 • FBXW4WNT1 530 2256 581 534 411 • FOSL1FBXW4 541 2233 215 1847 276 • FBXW2FGF4 548 2118 1771 2071 1465 • CXXC4FBXW4 559 2056 500 1147 656 • FBXW11SLC9A3R1 560 663 1910 685 1185 • APCFBXW2 562 2434 767 710 154 • APCFBXW4 567 1833 257 2238 402 • FBXW11NKD1 568 721 1583 999 2423 • FZD1FBXW4 570 504 561 2161 1305 • FBXW4T 592 622 681 1260 1597 • FBXW11PITX2 601 630 1775 747 1803 • DIXDC1FBXW4 603 1793 51 2313 1843 • FBXW4WNT3 607 900 1984 988 278 • CSNK2A1FBXW4 613 1164 1627 533 1031 • FZD7FBXW4 614 745 178 2081 2461 • FBXW2WIF1 625 1013 1649 2248 2208 • FBXW11FZD6 643 49 1989 1116 2444 • FBXW11KREMEN1 647 343 1611 2008 2172 • FZD5FBXW4 648 1963 383 2062 485 • FBXW11FSHB 651 1546 1867 1011 2263 • FBXW11WNT4 653 1660 1080 587 1321 • FBXW11NLK 659 182 1790 881 174 • FBXW11GSK3B 660 1365 1025 2246 1093 • FBXW11PPP2CA 664 317 1608 2254 2394 • FBXW11TCF7 670 1470 907 1860 743 • FBXW11SFRP4 690 1997 1896 1609 1818 • LRP5FBXW4 691 2170 794 1716 895 • FBXW11WNT2 703 1195 2438 200 1676 • FBXW2NKD1 714 856 1397 2319 335 • PPP2R1AFBXW4 722 1133 1095 2327 1957 • FBXW11FBXW4 736 310 962 1759 2364 • CSNK2A1FBXW2 764 907 2038 2445 74 • CSNK1DFBXW4 787 2387 1096 1415 2089 • FZD8FBXW4 788 1612 209 640 2357 • CSNK1DFBXW2 798 2207 1203 2384 477 • FBXW11PPP2R1A 807 789 2356 1137 447 • FBXW11FBXW2 811 20 2224 405 587 • CXXC4FBXW2 813 2183 535 536 697 • FBXW11FZD8 817 1577 1454 1362 1544 • FBXW11WNT3A 820 727 2251 1940 1879 • DIXDC1FBXW2 829 1425 39 1735 683 • FRAT1FBXW4 831 2047 491 468 653 • FBXW2FBXW4 856 583 2000 889 2175 • GSK3BFBXW4 858 1983 854 238 2279 • CCND3FBXW4 905 61 1333 2205 1979 • KREMEN1FBXW4 907 1905 333 340 881 • FBXW2PITX2 924 394 1742 1060 1621 • FBXW11FRAT1 932 1177 1783 245 1651 • FBXW11WNT5A 939 227 1476 41 1067 • CTBP1FBXW4 945 2064 203 1615 626 • FBXW11LRP5 949 1556 512 1200 2375 • NLKFBXW4 959 139 2167 1709 1870 • FZD6FBXW4 960 1357 1658 1703 1094 • FZD5FBXW2 966 1696 893 1023 225 • FBXW4TLE1 970 1784 1901 1274 1623 • FBXW2TLE1 979 656 760 1673 300 • EP300FBXW2 994 1076 589 1127 580 • EP300FBXW4 1008 1461 781 2265 1673 • PPP2CAFBXW4 1011 684 825 1625 1891 • FBXW11WNT2B 1015 460 1352 958 2373 • NKD1FBXW4 1016 541 502 1955 1381 • DAAM1FBXW4 1020 1522 272 692 1792 • FBXW2SFRP4 1032 1388 1682 580 1973 • CSNK2A1FBXW11 1045 480 849 1377 1270 • FBXW11TLE2 1067 1205 1951 2341 2338 • FBXW11FZD7 1069 106 1981 1733 2317 • FBXW2KREMEN1 1083 542 1986 1452 2414 • APCFBXW11 1090 2348 447 744 1026 • FBXW2SLC9A3R1 1099 1656 1475 635 1363 • FBXW4SLC9A3R1 1101 651 2159 2052 432 • FBXW11RHOU 1110 378 2144 1995 2294 • CCND3FBXW2 1143 391 1957 1745 495 • FBXW2FSHB 1161 118 571 611 1029 • SFRP4FBXW4 1164 2456 121 2465 1126 • FSHBFBXW4 1165 1304 990 976 2101 • FBXW11FOSL1 1175 661 1044 867 1802 • DAAM1FBXW2 1184 476 1344 1054 1680 • FBXW2NLK 1210 533 541 2016 1407 • FBXW11FZD1 1228 380 1793 1485 2376 • FBXW2WNT2 1252 742 1450 456 1982 • PITX2FBXW4 1280 1818 415 1204 1714 • FBXW4WNT2 1285 1404 1507 1107 1290 • CTNNBIP1FBXW4 1291 2373 378 1902 716 • FBXW2PPP2CA 1298 76 469 359 1581 • CSNK1DFBXW11 1311 1870 777 1384 803 • FBXW11TCF7L1 1313 232 2467 240 1749 • FBXW2GSK3B 1314 459 2279 1395 804 • FBXW4WNT2B 1317 223 650 557 783 • FBXW2WNT4 1332 1315 2385 870 1326 • FBXW2FZD6 1333 2127 430 1859 2058 • FBXW2TCF7 1340 471 935 2092 1162 • FBXW4WNT4 1342 1042 1537 2013 1382 • CSNK1G1FBXW4 1346 2371 1612 1422 2179 • FBXW4TCF7 1356 2088 960 726 1047 • FBXW2FZD8 1389 582 1890 1545 2182 • DIXDC1FBXW11 1400 871 7 1337 1602 • FBXW2PPP2R1A 1408 1022 1320 1039 523 • FBXW2WNT3A 1412 978 948 2419 2460 • CTBP1FBXW2 1418 1640 780 1833 156 • FBXW2WNT2B 1421 333 376 1464 1225 • FBXW2LRP5 1437 997 1863 2075 1869 • FZD5FBXW11 1438 429 363 1179 346 • CTBP2FBXW4 1447 234 229 2422 1542 • FBXW2FZD1 1463 1182 1496 689 2245 • FBXW2RHOU 1501 921 1894 310 1049 • DVL1FBXW4 1503 157 207 2177 1837 • EP300FBXW11 1505 37 741 1666 1426 • FBXW2TLE2 1508 2264 2140 2306 2372 • FBXW4WNT3A 1527 607 2312 2278 586 • LRP6FBXW4 1534 2334 1448 2256 1561 • FBXW4WNT5A 1539 97 984 1118 481 • CTNNB1FBXW4 1565 2145 856 1617 1790 • FBXW2FOSL1 1570 336 2085 1111 838 • CXXC4FBXW11 1572 2137 385 1398 1704 • AXIN1FBXW4 1573 1495 929 1150 1183 • FBXW2WNT5A 1588 1262 1210 1312 615 • FBXW2FRAT1 1641 1422 2017 1002 2184 • DVL2FBXW2 1659 908 1104 2239 786 • AESFBXW4 1663 2036 1157 97 2185 • CCND3FBXW11 1666 2158 1977 1020 1916 • CTBP2FBXW2 1668 1173 772 2115 731 • CTNNBIP1FBXW2 1669 1569 1031 666 67 • FBXW4TCF7L1 1671 1210 2407 1048 1190 • FBXW4TLE2 1677 1048 1543 1129 1442 • DVL2FBXW4 1681 945 569 693 1604 • CTNNB1FBXW2 1685 1796 897 1173 414 • CSNK1G1FBXW11 1687 2289 1332 181 2256 • LEF1FBXW4 1700 1559 773 2157 1543 • AXIN1FBXW2 1708 1181 2206 752 269 • FBXW2FZD7 1711 136 2013 2357 997 • DAAM1FBXW11 1718 1019 1182 1581 1528 • CTBP1FBXW11 1748 1868 340 177 292 • CSNK1G1FBXW2 1760 1176 1231 2470 2008 • SFRP1FBXW4 1765 1442 327 2117 2300 • CTNNB1FBXW11 1766 1709 460 961 1606 • CCND2FBXW4 1767 1915 911 2404 2400 • BTRCFBXW2 1768 48 1686 1888 1559 • PORCNFBXW4 1774 1885 1114 932 1800 • AESFBXW2 1783 926 230 1372 1624 • BTRCFBXW4 1807 1429 2081 520 2360 • FOXN1FBXW4 1861 414 2020 412 1359 • CSNK1A1FBXW4 1910 612 949 1336 236 • FBXW2TCF7L1 1919 1867 1313 890 1846 • CTNNBIP1FBXW11 1930 2021 103 1254 232 • DVL1FBXW2 1944 478 267 504 2051 • FZD2FBXW4 1990 2132 213 617 284 • FGF4FBXW4 1994 1499 132 1094 2001 • CTBP2FBXW11 2026 740 92 2069 576 • JUNFBXW4 2029 511 107 2393 379 • BCL9FBXW4 2038 2477 677 2031 1956 • MYCFBXW4 2071 1863 799 1227 860 • AESFBXW11 2084 786 1246 1608 1398 • CCND2FBXW2 2114 1187 747 1069 973 • PYGO1FBXW4 2115 2139 842 947 1341 • FBXW11FRZB 2118 1065 2214 101 2457 • BCL9FBXW2 2119 1989 436 1944 917 • DVL1FBXW11 2121 79 206 1361 632 • CCND2FBXW11 2140 295 214 1912 2465 • BTRCFBXW11 2176 286 480 1222 769 • DVL2FBXW11 2181 1392 373 1482 1223 • GSK3AFBXW4 2184 2369 1679 2432 198 • CCND1FBXW4 2193 31 370 1957 1245 • CSNK1A1FBXW2 2206 715 147 2390 69 • AXIN1FBXW11 2228 2061 613 458 924 • CSNK1A1FBXW11 2347 85 709 452 324 • CCND1FBXW2 2373 404 504 1522 1862 • FBXW2SENP2 2382 1297 1854 205 155 • FBXW11SENP2 2387 1355 1933 60 308 • FBXW2FRZB 2425 2345 1642 1273 1867 • CCND1FBXW11 2429 226 635 886 949 • BCL9FBXW11 2430 1009 638 1977 2079

## FBXW family 2*^nd^* order interaction ranking at different durations

• FZD5FBXW2 8 921 2298 1953 • PYGO1FBXW4 14 245 1608 571 • CSNK1A1FBXW11 23 1773 1902 11 • FZD5FBXW11 35 549 421 2435 • CSNK1A1FBXW2 38 703 1802 503 • FZD5FBXW4 44 455 1805 893 • CTNNB1FBXW2 81 1596 1956 1568 • CTBP1FBXW2 125 2182 403 2046 • DIXDC1FBXW2 140 18 205 1422 • CTBP1FBXW11 176 2347 1757 1223 • CTNNB1FBXW11 205 2122 354 1884 • AXIN1FBXW11 209 725 557 2148 • CCND2FBXW2 221 1706 1603 688 • DIXDC1FBXW11 233 38 107 1604 • AESFBXW11 253 1993 662 989 • AXIN1FBXW2 254 939 416 2093 • CSNK1A1FBXW4 259 1108 1636 369 • CCND1FBXW11 262 756 373 2151 • PORCNFBXW4 265 373 1258 1078 • CCND1FBXW2 266 1705 528 1905 • CTNNBIP1FBXW2 269 1311 419 1217 • BCL9FBXW11 272 1351 2463 1550 • CTNNB1FBXW4 302 1650 1192 391 • BCL9FBXW2 324 1009 1993 657 • CTNNBIP1FBXW11 347 2394 208 2065 • CCND2FBXW11 348 1409 1037 1403 • RHOUFBXW4 355 331 1480 559 • AESFBXW2 376 758 647 339 • JUNFBXW4 398 812 2338 140 • CSNK1DFBXW11 475 359 866 1396 • CSNK1DFBXW2 476 614 2148 477 • DVL1FBXW2 500 1199 101 1826 • DVL1FBXW11 509 2454 134 1697 • APCFBXW11 542 2199 1851 2444 • AXIN1FBXW4 544 628 760 1344 • FBXW2SENP2 550 2336 993 337 • FRAT1FBXW4 561 376 1783 643 • DIXDC1FBXW4 579 12 391 1036 • APCFBXW2 581 2325 2146 1662 • BTRCFBXW2 605 876 2002 1355 • FBXW11SENP2 606 1861 1887 996 • BTRCFBXW11 625 978 82 1632 • AESFBXW4 628 715 966 442 • BCL9FBXW4 634 1065 1857 2009 • CSNK1DFBXW4 642 324 2008 878 • CCND1FBXW4 654 557 1488 1657 • CTBP2FBXW2 677 1634 1487 556 • CCND2FBXW4 678 2355 950 1443 • CTBP1FBXW4 718 1496 1920 1249 • EP300FBXW11 750 872 425 651 • FSHBFBXW4 763 214 2395 2118 • KREMEN1FBXW4 800 1130 1417 353 • DVL2FBXW2 811 2297 1179 1703 • CTBP2FBXW11 823 740 707 1076 • MYCFBXW4 828 633 1896 976 • EP300FBXW2 831 2218 1637 1576 • FBXW4WNT5A 838 1157 1034 672 • DVL2FBXW11 842 1955 125 1822 • FBXW11SFRP4 863 2160 1731 110 • FBXW2SFRP4 879 2244 1420 919 • GSK3AFBXW4 899 164 1170 2074 • CTNNBIP1FBXW4 901 2349 179 969 • APCFBXW4 904 2149 1575 1957 • DKK1FBXW2 966 256 939 1318 • CXXC4FBXW2 990 2017 2165 2222 • FBXW2LRP6 991 2386 2358 2167 • FOXN1FBXW4 1024 1417 1822 526 • BTRCFBXW4 1029 566 2459 557 • FBXW4TCF7L1 1043 1981 588 1670 • CCND3FBXW2 1053 1740 699 616 • DKK1FBXW11 1070 155 1941 2052 • FBXW2FZD7 1083 1310 490 1643 • FOSL1FBXW4 1087 309 2408 1259 • CCND3FBXW11 1107 2455 655 1749 • CXXC4FBXW11 1109 2013 2059 2042 • FBXW2LEF1 1112 1499 2297 1519 • CTBP2FBXW4 1127 644 2127 1996 • FBXW2FZD1 1143 1827 1722 1636 • FBXW2FGF4 1151 2075 1051 1266 • FBXW11LRP6 1166 2404 2237 2412 • DAAM1FBXW11 1173 1903 172 590 • DVL1FBXW4 1183 928 1839 1685 • FBXW4TCF7 1184 1502 148 1505 • FBXW11FGF4 1198 2217 1578 751 • FBXW4TLE2 1199 2299 1778 483 • EP300FBXW4 1217 1781 1419 1291 • FBXW2WNT5A 1225 640 1973 2204 • FBXW11PPP2R1A 1232 1947 2230 1068 • FBXW11LEF1 1242 2193 1195 479 • DVL2FBXW4 1249 1578 2356 1232 • FBXW11FZD2 1255 2352 940 957 • FBXW2SFRP1 1260 2005 1228 1481 • FBXW11FZD7 1266 919 785 1176 • FBXW11SFRP1 1274 2040 2020 1623 • FBXW11GSK3B 1291 1302 973 877 • FBXW11TCF7L1 1297 1926 1394 2483 • FBXW2FZD2 1308 2003 1620 2229 • DAAM1FBXW2 1313 1369 541 1329 • FBXW11FZD1 1315 2039 2147 1846 • FBXW11WNT5A 1329 716 1071 1970 • FBXW2PPP2R1A 1350 2095 1932 2171 • LRP5FBXW4 1368 140 2283 72 • CSNK2A1FBXW2 1394 1844 202 2410 • FBXW2FZD6 1402 2050 1206 1026 • CXXC4FBXW4 1404 1485 999 2199 • FBXW2GSK3B 1405 1923 1075 670 • FBXW11FBXW2 1409 609 1260 1452 • NKD1FBXW4 1412 1419 2207 1445 • FBXW2TCF7L1 1418 2209 2038 1942 • FBXW2FZD8 1427 1762 1246 1902 • FBXW11PITX2 1469 1756 877 1859 • FRZBFBXW4 1473 1301 1964 55 • FBXW11FZD8 1479 2266 741 1524 • FBXW2PITX2 1482 1375 1290 1823 • FBXW4TLE1 1487 1575 2428 1552 • FBXW2NLK 1489 1243 312 1941 • FBXW2TLE2 1491 1483 1917 1340 • CCND3FBXW4 1519 2144 943 2049 • FBXW11FZD6 1528 2449 546 305 • CSNK1G1FBXW11 1531 73 160 2357 • FBXW11TLE2 1543 881 1400 343 • DKK1FBXW4 1559 147 1776 655 • FBXW11FBXW4 1563 596 1329 1312 • FZD6FBXW4 1585 371 263 460 • FBXW11NKD1 1608 706 1351 2143 • FBXW2TCF7 1612 1804 410 1414 • NLKFBXW4 1620 1752 898 1933 • FBXW4WNT4 1623 1131 2212 1683 • PPP2CAFBXW4 1638 1289 1438 1147 • FBXW11NLK 1645 1512 1040 1133 • FBXW2PPP2CA 1655 1907 2065 255 • FBXW11PPP2CA 1657 1211 1377 1977 • FBXW2FBXW4 1673 1488 2073 1695 • FBXW2GSK3A 1691 1749 1062 415 • CSNK2A1FBXW11 1706 326 116 2242 • FGF4FBXW4 1714 122 1583 769 • DAAM1FBXW4 1736 340 660 1952 • FBXW4WNT1 1756 878 2089 1557 • FBXW2LRP5 1757 1184 2450 2012 • FBXW11FRZB 1764 446 1407 1146 • FZD7FBXW4 1767 92 252 1337 • FBXW4WNT2 1773 2134 1457 1612 • FBXW4WIF1 1781 1421 605 2069 • PITX2FBXW4 1798 8 313 692 • GSK3BFBXW4 1800 20 776 261 • FBXW11TLE1 1815 1035 1729 912 • FBXW11LRP5 1829 1690 1566 862 • SFRP1FBXW4 1840 39 1050 1794 • FZD1FBXW4 1846 28 2332 579 • CSNK1G1FBXW2 1861 261 1641 2144 • CSNK2A1FBXW4 1869 967 462 999 • FBXW2NKD1 1874 429 192 2274 • FBXW2WNT4 1878 1000 1474 1107 • FBXW2FRZB 1887 709 989 1314 • FBXW2FRAT1 1896 2066 1518 637 • FBXW11WNT4 1901 588 1687 2330 • FBXW11TCF7 1904 673 337 1407 • FZD8FBXW4 1915 56 733 1379 • PPP2R1AFBXW4 1924 765 1004 279 • FBXW4SLC9A3R1 1929 1474 2249 2035 • FBXW11MYC 1933 1925 1831 1311 • FBXW2T 1968 2460 1685 522 • FBXW2WIF1 1970 1707 2115 1845 • FBXW11GSK3A 1974 1299 820 720 • FBXW2TLE1 1987 2337 1408 1273 • FBXW4T 2006 1805 488 354 • FBXW2WNT3A 2018 1969 841 2402 • FBXW11WIF1 2019 1290 2160 954 • CSNK1G1FBXW4 2024 36 1118 2084 • FBXW2WNT1 2026 1723 1128 966 • FBXW11WNT1 2028 570 1439 446 • FBXW2SLC9A3R1 2045 1832 1101 1436 • SFRP4FBXW4 2068 119 952 695 • FBXW11FOSL1 2072 2194 892 1566 • FBXW2MYC 2099 1764 476 2322 • FBXW11FOXN1 2107 389 813 2188 • LEF1FBXW4 2112 128 2235 1180 • FBXW2FOXN1 2130 589 2162 1841 • FBXW11T 2148 1905 2218 1737 • FBXW2FOSL1 2150 1859 1765 1457 • FBXW2RHOU 2151 801 1957 760 • FBXW2WNT2 2154 2353 1824 2104 • FBXW2WNT2B 2186 2375 732 346 • FZD2FBXW4 2207 380 1382 1860 • FBXW11JUN 2217 527 1775 1281 • FBXW11WNT2 2219 1443 1707 2257 • LRP6FBXW4 2242 160 777 1686 • FBXW2KREMEN1 2248 1975 615 1983 • FBXW4WNT3 2256 1182 288 1250 • FBXW4WNT3A 2274 748 1785 1584 • FBXW11FRAT1 2281 2204 2130 2146 • SENP2FBXW4 2297 269 1705 1913 • FBXW2JUN 2306 1134 2046 314 • FBXW11WNT3A 2313 1616 1148 2216 • FBXW11FSHB 2318 1373 2232 780 • FBXW2PYGO1 2327 2363 2236 1679 • FBXW2WNT3 2338 1293 530 2124 • FBXW11PORCN 2352 1091 2275 2364 • FBXW11WNT2B 2360 2259 2111 773 • FBXW2FSHB 2363 1812 1515 245 • FBXW2PORCN 2399 727 2452 2301 • FBXW4WNT2B 2413 1945 368 534 • FBXW11PYGO1 2417 1539 1074 2283 • FBXW11SLC9A3R1 2422 1440 1923 884 • FBXW11KREMEN1 2425 920 1433 2436 • FBXW11RHOU 2442 1489 1234 674 • FBXW11WNT3 2458 1759 1695 1850

## TCF family 2*^nd^* order interaction ranking at different time points

• SENP2TCF7 57 1107 117 945 472 • TCF7L1WIF1 60 566 2194 668 1176 • TCF7L1WNT1 87 1846 233 128 356 • FRZBTCF7 101 2467 241 2087 866 • TCF7L1WNT3 126 5 2233 859 133 • SENP2TCF7L1 134 885 1070 292 1861 • TCF7WNT1 207 1677 1463 249 669 • TCF7WIF1 324 1432 2083 2255 1812 • FRZBTCF7L1 349 2309 348 1223 1940 • TCF7WNT3 368 620 1195 1838 1044 • DKK1TCF7 407 2338 1936 1761 1438 • TCF7L1TLE1 489 1111 2064 2460 1758 • DKK1TCF7L1 550 1958 2221 740 1641 • FZD1TCF7 599 379 582 2335 2034 • TCF7L1WNT4 663 1031 2032 2406 836 • APCTCF7 665 1986 177 2281 37 • FBXW11TCF7 670 1470 907 1860 743 • TCF7TLE1 671 1145 1750 1990 1073 • TCF7L1WNT2 683 1190 1364 2060 755 • TCF7L1WNT5A 688 229 702 1670 209 • FZD7TCF7 747 595 38 479 1993 • FOSL1TCF7 789 1761 60 79 1133 • TCF7L1WNT3A 804 1237 1263 1513 1523 • TCF7L1WNT2B 812 128 237 1311 2407 • GSK3BTCF7 878 1131 66 306 1964 • CSNK2A1TCF7 896 324 1552 96 1506 • FRAT1TCF7 915 2075 74 2153 1996 • TCF7L1TLE2 937 1563 1915 2448 1395 • CXXC4TCF7 1001 1287 336 1751 2443 • RHOUTCF7 1002 1326 359 603 1896 • PPP2R1ATCF7 1022 2257 886 2076 941 • LRP5TCF7 1026 2443 1579 113 340 • FZD5TCF7 1033 1247 253 2030 1030 • NLKTCF7 1085 732 1653 1379 937 • CSNK1DTCF7 1089 1141 1171 2462 598 • TCF7WNT5A 1112 151 2019 1176 1463 • TCF7WNT2 1131 1699 1307 434 1972 • APCTCF7L1 1174 1819 1610 1801 974 • SLC9A3R1TCF7 1197 1474 142 836 664 • DIXDC1TCF7 1213 1189 13 1532 1304 • FZD1TCF7L1 1223 1749 1351 2446 846 • TCF7WNT3A 1231 1488 2137 1901 1157 • TCF7WNT4 1237 2128 1970 69 2139 • KREMEN1TCF7 1238 1608 146 1187 2194 • TCF7TLE2 1260 585 1207 1743 2426 • FZD7TCF7L1 1295 191 695 2479 533 • PPP2CATCF7 1309 1817 2014 1805 211 • FBXW11TCF7L1 1313 232 2467 240 1749 • FOSL1TCF7L1 1328 2216 736 1968 2439 • RHOUTCF7L1 1329 2234 788 43 2428 • FZD8TCF7 1338 895 264 1471 2442 • FBXW2TCF7 1340 471 935 2092 1162 • FBXW4TCF7 1356 2088 960 726 1047 • EP300TCF7 1357 1256 437 832 252 • FZD6TCF7 1376 1230 675 979 1832 • CSNK2A1TCF7L1 1388 487 1758 1542 1883 • CCND3TCF7 1396 462 964 2257 1674 • DIXDC1TCF7L1 1402 1671 349 283 1296 • PITX2TCF7 1426 1956 102 868 1214 • FRAT1TCF7L1 1427 2111 879 875 666 • DAAM1TCF7 1439 608 497 2187 1366 • SFRP4TCF7 1471 2462 68 1217 648 • TCF7WNT2B 1475 274 1699 1748 665 • CSNK1DTCF7L1 1510 2074 2257 481 1186 • FSHBTCF7 1543 1260 1113 2024 856 • CTBP1TCF7 1552 2479 368 2112 802 • FZD5TCF7L1 1558 1584 1067 2129 1440 • EP300TCF7L1 1576 1098 2314 1772 1801 • CTNNB1TCF7 1591 639 417 1616 543 • KREMEN1TCF7L1 1609 377 1006 1658 2251 • CXXC4TCF7L1 1611 1104 859 173 90 • TCF7TCF7L1 1647 2259 1383 1530 1655 • FBXW4TCF7L1 1671 1210 2407 1048 1190 • LRP5TCF7L1 1691 1313 2302 1624 744 • FZD6TCF7L1 1709 2082 2195 1280 2328 • GSK3BTCF7L1 1710 1105 699 1763 2191 • PPP2R1ATCF7L1 1720 874 1330 1368 1053 • DVL2TCF7 1734 1838 1090 1468 151 • NLKTCF7L1 1746 59 1592 494 1267 • DVL1TCF7 1780 2282 165 1096 2050 • CTBP2TCF7 1782 863 191 2366 352 • CTNNBIP1TCF7 1784 2198 118 1087 241 • PPP2CATCF7L1 1786 248 744 2289 2143 • LRP6TCF7 1808 148 435 2484 2349 • NKD1TCF7 1813 165 654 1407 2043 • DAAM1TCF7L1 1822 1460 2393 2097 418 • SFRP4TCF7L1 1839 2482 1253 2414 1701 • FZD8TCF7L1 1844 1057 682 1153 1825 • CSNK1G1TCF7 1856 1879 1760 1698 1988 • CTBP1TCF7L1 1891 1743 517 1031 1628 • FSHBTCF7L1 1895 1449 2343 1015 902 • MYCTCF7 1917 1611 713 672 70 • FBXW2TCF7L1 1919 1867 1313 890 1846 • PITX2TCF7L1 1935 1251 873 962 1259 • BTRCTCF7 1942 1126 1508 553 906 • CCND2TCF7 1945 808 360 2118 1781 • SLC9A3R1TCF7L1 1958 953 1560 601 465 • FOXN1TCF7 1976 1780 888 707 808 • FGF4TCF7 1979 777 106 1923 821 • CSNK1A1TCF7 1988 2185 646 2178 45 • FZD2TCF7 1992 1456 48 1531 1301 • CCND3TCF7L1 2001 1103 2434 1915 2168 • LRP6TCF7L1 2037 955 1634 1396 428 • CTNNB1TCF7L1 2043 944 805 381 1449 • BCL9TCF7 2044 1809 133 1660 521 • AXIN1TCF7 2051 2104 1150 798 1600 • TTCF7 2086 1882 864 1982 790 • CCND2TCF7L1 2091 1628 1322 1549 1902 • NKD1TCF7L1 2096 1068 1257 2346 2127 • CSNK1G1TCF7L1 2097 2094 1978 535 1446 • DVL2TCF7L1 2098 103 1781 382 2121 • CTNNBIP1TCF7L1 2123 1739 1130 1580 43 • BTRCTCF7L1 2134 27 1718 1067 2162 • SFRP1TCF7 2152 239 57 2228 1797 • TTCF7L1 2153 2108 1509 2317 651 • LEF1TCF7 2155 1150 503 2459 1258 • AXIN1TCF7L1 2194 1538 1356 497 1783 • BCL9TCF7L1 2213 2196 1230 1013 1060 • AESTCF7L1 2224 634 1176 407 1824 • PYGO1TCF7 2243 1746 1075 981 1059 • PYGO1TCF7L1 2249 1439 2258 1038 2323 • PORCNTCF7 2260 98 906 2211 2312 • MYCTCF7L1 2269 2204 2429 2158 2092 • AESTCF7 2276 162 422 1436 609 • CTBP2TCF7L1 2278 2283 1810 1881 2472 • PORCNTCF7L1 2292 1382 731 1405 1475 • FGF4TCF7L1 2293 1483 733 1849 1055 • DVL1TCF7L1 2298 1 1306 1930 2016 • FOXN1TCF7L1 2305 444 608 743 1132 • LEF1TCF7L1 2306 1102 1559 1237 336 • CSNK1A1TCF7L1 2312 941 1272 2266 1593 • SFRP1TCF7L1 2324 1770 505 2473 2174 • FZD2TCF7L1 2349 479 438 498 746 • JUNTCF7 2362 96 311 1993 516 • GSK3ATCF7 2365 2110 1061 1796 1180 • JUNTCF7L1 2384 1691 1186 2021 1413 • CCND1TCF7 2393 682 220 1588 2165 • CCND1TCF7L1 2402 552 1394 554 1794 • GSK3ATCF7L1 2426 2078 2284 1423 1585

## TCF family 2*^nd^* order interaction ranking at different durations

• FZD5TCF7L1 40 2452 2050 1910 • FZD5TCF7 89 797 804 1610 • PYGO1TCF7 90 656 157 1555 • PYGO1TCF7L1 93 1930 1059 228 • CSNK1A1TCF7L1 97 1169 2288 1506 • CTBP1TCF7L1 120 1967 1053 2317 • RHOUTCF7L1 135 1328 163 836 • CTNNB1TCF7L1 157 1641 1213 2213 • RHOUTCF7 175 410 31 784 • JUNTCF7L1 240 1863 2085 2230 • DIXDC1TCF7L1 250 906 242 1241 • PORCNTCF7 261 1647 279 2440 • AESTCF7L1 276 2332 1834 1985 • PORCNTCF7L1 284 2273 1305 1648 • CSNK1A1TCF7 286 864 818 1113 • TTCF7L1 328 1729 1286 821 • AXIN1TCF7L1 345 1895 744 1140 • FOXN1TCF7L1 371 2468 788 931 • KREMEN1TCF7L1 397 1321 306 1898 • FRAT1TCF7L1 400 1306 583 2015 • CCND2TCF7L1 406 1089 1879 1271 • DIXDC1TCF7 431 47 26 848 • CSNK1DTCF7L1 435 1239 1609 851 • CCND1TCF7L1 441 1604 784 2355 • BCL9TCF7L1 448 2393 1787 2073 • FSHBTCF7L1 451 2374 527 1397 • MYCTCF7L1 480 1678 2094 2166 • SLC9A3R1TCF7L1 484 2420 753 1394 • CTNNBIP1TCF7L1 487 2088 417 551 • AESTCF7 539 739 257 1757 • DVL1TCF7L1 551 1938 1080 1023 • BTRCTCF7L1 566 2170 2239 2208 • JUNTCF7 578 916 434 708 • TTCF7 590 827 78 203 • CTNNB1TCF7 597 1307 261 1851 • CTBP2TCF7L1 604 2378 2429 1184 • CTNNBIP1TCF7 608 2177 72 530 • GSK3ATCF7L1 611 1629 304 1867 • AXIN1TCF7 620 574 357 1827 • CTBP1TCF7 630 985 1120 2343 • FRAT1TCF7 662 678 725 300 • APCTCF7L1 684 1317 2093 2313 • CSNK1DTCF7 685 571 45 1780 • FOSL1TCF7L1 729 1225 793 752 • CCND1TCF7 732 877 61 1750 • FSHBTCF7 736 473 370 1014 • KREMEN1TCF7 742 1064 65 2424 • CCND2TCF7 777 1675 194 2285 • LRP5TCF7L1 778 1338 1381 1572 • NKD1TCF7L1 792 1890 1826 540 • BCL9TCF7 795 1286 352 1779 • APCTCF7 815 1780 1652 1231 • FOXN1TCF7 833 1179 338 1573 • BTRCTCF7 841 1396 217 1525 • SLC9A3R1TCF7 852 811 23 1230 • EP300TCF7L1 864 1193 1005 1984 • DVL2TCF7L1 867 1191 716 2293 • MYCTCF7 877 1694 277 1834 • TCF7TCF7L1 886 1648 1654 2119 • DVL1TCF7 905 1899 37 1262 • CCND3TCF7L1 934 966 542 2363 • CTBP2TCF7 936 1305 254 1468 • FRZBTCF7L1 957 1843 2335 1190 • FOSL1TCF7 969 353 727 2110 • EP300TCF7 985 1105 28 1729 • DKK1TCF7L1 1005 1640 2437 2338 • FBXW4TCF7L1 1043 1981 588 1670 • GSK3ATCF7 1068 427 2 1339 • DAAM1TCF7L1 1095 1524 405 1831 • TCF7WNT5A 1117 1585 2010 1400 • DKK1TCF7 1119 246 2106 1166 • FRZBTCF7 1121 2074 708 469 • GSK3BTCF7L1 1137 603 444 1542 • NKD1TCF7 1152 1171 347 2320 • LRP5TCF7 1164 130 392 306 • FZD6TCF7L1 1171 1172 117 1637 • NLKTCF7L1 1175 1980 1497 1746 • FZD8TCF7L1 1176 684 280 1818 • FBXW4TCF7 1184 1502 148 1505 • PPP2CATCF7L1 1189 2338 1201 1139 • FZD7TCF7L1 1281 1530 43 2004 • CXXC4TCF7L1 1283 1436 674 1495 • CCND3TCF7 1286 1916 563 2086 • FBXW11TCF7L1 1297 1926 1394 2483 • FZD1TCF7L1 1318 666 1114 2157 • DVL2TCF7 1354 2270 36 2475 • SFRP1TCF7L1 1371 690 722 1330 • FGF4TCF7L1 1389 1679 418 1163 • FBXW2TCF7L1 1418 2209 2038 1942 • TCF7TLE2 1428 1516 1010 718 • FZD8TCF7 1432 125 40 2379 • CSNK2A1TCF7L1 1437 2465 190 1536 • PITX2TCF7L1 1451 605 188 729 • PPP2CATCF7 1458 902 870 1478 • TCF7TLE1 1459 2288 1891 609 • DAAM1TCF7 1500 1918 112 1221 • PPP2R1ATCF7L1 1545 1262 645 1812 • TCF7WNT4 1555 1556 768 2469 • CSNK1G1TCF7L1 1565 1183 1483 2156 • FZD2TCF7L1 1570 815 250 1785 • TCF7L1WNT5A 1580 102 876 2243 • CXXC4TCF7 1596 1931 2013 1628 • LRP6TCF7L1 1598 2125 2438 2450 • FBXW2TCF7 1612 1804 410 1414 • LEF1TCF7L1 1617 1010 1423 2114 • FZD2TCF7 1688 1571 3 1092 • FZD7TCF7 1693 196 4 774 • NLKTCF7 1709 1725 534 2218 • GSK3BTCF7 1723 69 1 870 • TCF7L1TLE2 1725 388 2376 1500 • FZD6TCF7 1745 982 18 1066 • LRP6TCF7 1753 227 428 1980 • PITX2TCF7 1779 78 20 1471 • FZD1TCF7 1808 177 34 1044 • CSNK2A1TCF7 1832 1441 24 1462 • FGF4TCF7 1843 90 87 1967 • TCF7WNT2 1899 1519 539 2272 • TCF7WIF1 1900 1887 2186 1009 • FBXW11TCF7 1904 673 337 1407 • SFRP4TCF7L1 1912 456 408 187 • SFRP1TCF7 1943 236 272 829 • TCF7L1WNT4 1986 275 2043 2178 • TCF7WNT2B 1996 2298 1797 514 • PPP2R1ATCF7 1997 760 139 2467 • TCF7WNT1 2010 745 1110 2109 • CSNK1G1TCF7 2016 255 69 2411 • LEF1TCF7 2052 192 27 1037 • TCF7L1TLE1 2081 659 682 938 • SENP2TCF7L1 2082 2483 1906 1734 • TCF7L1WNT3A 2133 555 1460 1569 • TCF7WNT3 2197 2031 1446 457 • TCF7L1WNT1 2213 127 1840 1900 • TCF7L1WNT2 2227 204 976 2152 • SFRP4TCF7 2240 157 460 360 • TCF7L1WIF1 2262 984 1673 539 • TCF7WNT3A 2268 1292 694 648 • SENP2TCF7 2370 567 904 1238 • TCF7L1WNT3 2391 2474 349 1490 • TCF7L1WNT2B 2423 871 519 485

## PPP2 family 2*^nd^* order interaction ranking at different time points

• FRZBPPP2R1A 67 2232 413 806 178 • FRZBPPP2CA 78 2468 5 1286 226 • PPP2CAWNT1 85 849 2018 439 513 • PPP2R1AWNT1 129 619 566 660 582 • PPP2R1APYGO1 189 193 1877 193 1170 • DKK1PPP2R1A 204 2066 2151 2449 1450 • PPP2CASFRP1 244 124 2088 397 434 • PPP2CAPYGO1 271 1161 2001 1293 1221 • PPP2R1AWNT3 282 312 2409 1755 429 • PPP2R1AWIF1 291 1990 1823 1155 2341 • DKK1PPP2CA 296 2422 1285 2212 1236 • PPP2CAWIF1 310 573 1845 2408 2336 • PPP2R1AT 314 58 789 885 1795 • PPP2R1ASFRP1 359 545 1723 1265 2124 • PPP2CAWNT3 469 292 1807 1884 350 • PPP2R1ATLE1 558 294 1423 1061 1167 • FOSL1PPP2R1A 575 1617 298 845 244 • APCPPP2CA 586 2263 452 388 1224 • FZD7PPP2R1A 602 13 208 2441 1936 • FZD1PPP2CA 605 1893 306 1913 1892 • FZD7PPP2CA 662 642 32 247 1583 • FBXW11PPP2CA 664 317 1608 2254 2394 • CSNK1DPPP2CA 705 1681 1060 1300 2455 • FZD1PPP2R1A 706 1641 1385 1614 395 • PPP2CAT 721 1128 2008 2287 422 • PPP2R1AFBXW4 722 1133 1095 2327 1957 • LRP5PPP2CA 733 2161 410 2286 27 • PPP2CATLE1 735 1046 1882 930 492 • FOSL1PPP2CA 746 2176 24 377 1408 • APCPPP2R1A 777 1312 425 667 138 • FRAT1PPP2CA 792 2225 98 1041 1142 • CSNK2A1PPP2CA 801 423 350 2410 132 • FBXW11PPP2R1A 807 789 2356 1137 447 • DIXDC1PPP2CA 869 2380 6 2302 1899 • FZD5PPP2R1A 879 1118 946 1416 216 • FZD5PPP2CA 902 2331 240 416 1264 • PPP2R1ASFRP4 903 1887 2325 1461 1681 • CSNK2A1PPP2R1A 925 1399 1152 192 631 • PPP2CASLC9A3R1 933 477 2003 1825 629 • PPP2R1ASLC9A3R1 941 537 1618 1764 538 • FRAT1PPP2R1A 956 1900 1269 1611 197 • LRP5PPP2R1A 961 2368 1644 677 380 • NLKPPP2R1A 964 1051 2329 2067 570 • PPP2R1AWNT5A 980 2288 679 76 957 • PPP2CAFBXW4 1011 684 825 1625 1891 • PPP2R1ATCF7 1022 2257 886 2076 941 • CSNK1DPPP2R1A 1023 2376 2059 2251 874 • DIXDC1PPP2R1A 1044 434 70 1228 1344 • PPP2CASFRP4 1077 2092 1844 718 1109 • CXXC4PPP2CA 1098 2404 162 82 2214 • CCND3PPP2CA 1104 315 1299 1817 454 • FZD6PPP2CA 1119 204 1164 1951 1967 • EP300PPP2CA 1120 230 287 1381 1923 • NLKPPP2CA 1124 210 1518 1976 1653 • GSK3BPPP2CA 1130 2453 152 819 2206 • PPP2R1AWNT2B 1135 1299 999 171 1731 • CXXC4PPP2R1A 1152 919 377 1602 646 • PPP2R1ATLE2 1195 666 2197 2163 1732 • KREMEN1PPP2CA 1201 1634 423 1797 2471 • GSK3BPPP2R1A 1205 313 728 1562 596 • KREMEN1PPP2R1A 1209 318 658 2293 1386 • EP300PPP2R1A 1217 1206 660 1626 386 • PPP2CAWNT2 1218 483 2176 1246 2111 • PPP2R1AWNT2 1229 257 2305 615 478 • PPP2CAWNT5A 1246 1216 801 671 728 • PPP2CATLE2 1268 1135 1602 279 2289 • FZD8PPP2CA 1274 1730 77 1712 1876 • PPP2R1AWNT4 1294 1166 1499 2045 1006 • FBXW2PPP2CA 1298 76 469 359 1581 • FSHBPPP2CA 1302 1918 1417 659 2290 • PPP2CAPPP2R1A 1308 1614 1705 1191 1156 • PPP2CATCF7 1309 1817 2014 1805 211 • FZD6PPP2R1A 1324 724 1914 713 545 • FZD8PPP2R1A 1330 247 1007 822 854 • PPP2R1AWNT3A 1335 818 1303 2113 1901 • PPP2CAWNT3A 1367 576 1326 2477 1116 • PPP2CAWNT4 1371 1799 2106 1702 2195 • PITX2PPP2CA 1375 1363 190 1770 2187 • PPP2CAWNT2B 1399 1928 601 1165 1405 • FBXW2PPP2R1A 1408 1022 1320 1039 523 • CCND3PPP2R1A 1431 886 1168 1704 2030 • DAAM1PPP2CA 1434 1351 829 37 1457 • CTBP1PPP2CA 1440 1663 58 1033 1002 • PITX2PPP2R1A 1458 1821 202 271 762 • CTNNB1PPP2CA 1473 2067 197 356 1519 • PPP2CARHOU 1488 2115 1869 1130 1284 • CTBP1PPP2R1A 1523 1092 743 1727 77 • DVL2PPP2CA 1561 2052 365 1741 295 • DAAM1PPP2R1A 1592 516 625 1867 701 • CTNNB1PPP2R1A 1596 1191 813 839 457 • CTNNBIP1PPP2R1A 1599 1077 624 2338 48 • FSHBPPP2R1A 1630 475 1889 1497 383 • CTBP2PPP2CA 1638 864 153 1565 950 • NKD1PPP2CA 1651 160 249 1178 2450 • CTNNBIP1PPP2CA 1692 2310 71 1873 239 • PPP2R1ARHOU 1701 623 1843 1576 1644 • PPP2R1ATCF7L1 1720 874 1330 1368 1053 • AESPPP2CA 1772 1246 501 303 1200 • DVL2PPP2R1A 1777 1353 830 144 333 • CCND2PPP2CA 1781 936 445 493 1247 • PPP2CATCF7L1 1786 248 744 2289 2143 • CTBP2PPP2R1A 1789 1337 201 2200 688 • LRP6PPP2CA 1792 2445 575 2334 2084 • DVL1PPP2R1A 1793 143 804 650 272 • BTRCPPP2CA 1800 2191 648 2242 2241 • NKD1PPP2R1A 1805 609 661 1681 87 • BTRCPPP2R1A 1831 677 2435 2196 567 • CSNK1G1PPP2R1A 1849 2097 667 511 235 • AXIN1PPP2CA 1868 2296 506 827 2213 • CSNK1G1PPP2CA 1883 2254 1420 2033 1730 • CCND2PPP2R1A 1911 791 831 681 2198 • AXIN1PPP2R1A 1920 1290 1491 732 394 • DVL1PPP2CA 1923 1134 224 778 2067 • LRP6PPP2R1A 1924 116 134 1987 658 • LEF1PPP2CA 1953 1857 400 797 2090 • PORCNPPP2R1A 2019 1170 1047 213 2393 • CSNK1A1PPP2CA 2024 1694 254 759 750 • JUNPPP2CA 2042 341 99 860 1430 • MYCPPP2R1A 2057 1855 1714 1539 97 • FGF4PPP2CA 2077 2121 8 1529 1855 • BCL9PPP2CA 2094 1602 402 2250 1423 • PORCNPPP2CA 2103 186 275 274 897 • LEF1PPP2R1A 2124 644 1355 2182 466 • FZD2PPP2R1A 2149 1596 403 1711 19 • CSNK1A1PPP2R1A 2195 842 1325 1845 118 • CCND1PPP2CA 2203 252 226 58 1990 • AESPPP2R1A 2210 125 936 1790 453 • MYCPPP2CA 2225 563 499 688 1706 • JUNPPP2R1A 2265 569 614 1908 42 • BCL9PPP2R1A 2275 2457 416 2074 2116 • FGF4PPP2R1A 2287 1618 247 2241 262 • FZD2PPP2CA 2322 1776 288 2321 600 • GSK3APPP2CA 2328 2050 798 1503 2012 • FOXN1PPP2CA 2359 1881 905 1788 1983 • FOXN1PPP2R1A 2363 214 1779 1437 112 • CCND1PPP2R1A 2370 219 341 585 1489 • GSK3APPP2R1A 2410 1478 1731 1326 850 • PPP2CASENP2 2411 1750 2087 1097 369 • PPP2R1ASENP2 2446 2262 1926 696 2266

## PPP2 family 2*^nd^* order interaction ranking at different durations

• CSNK1A1PPP2R1A 43 1484 2382 316 • FZD5PPP2R1A 88 786 1605 1535 • CSNK1A1PPP2CA 96 1063 1339 1596 • FZD5PPP2CA 128 470 2021 1904 • DIXDC1PPP2R1A 146 26 374 1684 • CTBP1PPP2R1A 172 1593 1458 1512 • CTNNB1PPP2R1A 177 1614 1520 1805 • AESPPP2R1A 195 1651 1416 1431 • CTBP1PPP2CA 199 1626 1662 1216 • FOXN1PPP2R1A 222 1346 1375 491 • PORCNPPP2CA 228 1809 2291 1052 • FRAT1PPP2R1A 238 721 1472 1069 • JUNPPP2R1A 252 2251 1749 88 • DIXDC1PPP2CA 268 65 195 1061 • BCL9PPP2R1A 275 773 2409 1630 • CTNNBIP1PPP2R1A 288 2462 499 517 • CTNNB1PPP2CA 293 2018 1905 563 • AXIN1PPP2R1A 320 965 796 2133 • CCND1PPP2R1A 323 779 429 1222 • FSHBPPP2R1A 346 663 2087 1791 • AXIN1PPP2CA 351 923 977 2031 • CSNK1DPPP2R1A 370 551 2152 1035 • PORCNPPP2R1A 402 1111 445 2329 • PPP2CASENP2 413 2453 988 152 • KREMEN1PPP2R1A 445 726 475 1625 • CCND2PPP2R1A 450 1267 1045 1619 • DVL1PPP2R1A 482 875 287 2103 • MYCPPP2R1A 493 1587 1536 1966 • KREMEN1PPP2CA 497 714 549 2445 • JUNPPP2CA 506 1282 2056 421 • GSK3APPP2R1A 519 439 367 950 • FOXN1PPP2CA 529 736 364 1732 • PPP2CASFRP4 549 2262 1511 112 • APCPPP2R1A 554 1481 1501 2010 • FRAT1PPP2CA 570 501 891 1529 • MYCPPP2CA 571 862 2040 1218 • AESPPP2CA 576 1581 1365 2479 • BTRCPPP2R1A 580 1968 2265 1926 • CTNNBIP1PPP2CA 583 2220 620 1248 • CCND2PPP2CA 602 2451 1178 883 • CCND1PPP2CA 610 757 829 872 • FOSL1PPP2R1A 613 212 1052 2311 • CTBP2PPP2R1A 641 1672 2371 2245 • FSHBPPP2CA 659 383 953 406 • BCL9PPP2CA 675 1821 2150 1300 • LRP5PPP2R1A 689 60 1512 148 • NKD1PPP2R1A 714 1210 2247 894 • CSNK1DPPP2CA 755 680 333 1508 • DVL2PPP2R1A 756 2169 1396 2360 • PPP2R1ASENP2 779 2450 402 1310 • FOSL1PPP2CA 789 316 2481 2423 • BTRCPPP2CA 810 1170 2051 1848 • GSK3APPP2CA 813 338 235 772 • DVL1PPP2CA 820 1888 482 1018 • APCPPP2CA 824 1772 1975 2014 • NKD1PPP2CA 868 1795 1696 2267 • CCND3PPP2R1A 871 963 1863 2279 • FRZBPPP2R1A 924 1119 2374 1727 • EP300PPP2R1A 953 2185 1215 2359 • DKK1PPP2R1A 954 260 1796 1911 • DVL2PPP2CA 955 841 1096 2248 • CTBP2PPP2CA 984 835 2336 1644 • CXXC4PPP2R1A 1023 1884 1841 2050 • FRZBPPP2CA 1033 541 2381 2106 • PPP2R1ASFRP4 1036 2026 1395 840 • DAAM1PPP2R1A 1039 850 981 1633 • PPP2CAPPP2R1A 1044 814 2362 965 • PPP2CAWNT5A 1111 1029 2392 2262 • FZD8PPP2R1A 1114 95 449 699 • FZD6PPP2R1A 1133 602 154 533 • NLKPPP2R1A 1154 653 2402 1819 • PITX2PPP2R1A 1159 80 135 1367 • CCND3PPP2CA 1161 2020 603 387 • PPP2CATCF7L1 1189 2338 1201 1139 • PPP2CASFRP1 1193 2137 2035 610 • FZD1PPP2R1A 1203 117 1936 1143 • EP300PPP2CA 1211 2114 1082 1383 • FBXW11PPP2R1A 1232 1947 2230 1068 • LRP5PPP2CA 1267 31 420 1882 • GSK3BPPP2R1A 1282 98 207 171 • FZD6PPP2CA 1300 1658 102 704 • CSNK2A1PPP2R1A 1305 674 221 2299 • DKK1PPP2CA 1309 173 1479 2442 • FGF4PPP2R1A 1323 211 1708 879 • FBXW2PPP2R1A 1350 2095 1932 2171 • FZD7PPP2R1A 1360 120 49 2284 • PPP2R1AWNT5A 1366 431 652 734 • FZD8PPP2CA 1415 75 361 1857 • PPP2R1ASFRP1 1420 1757 2063 2189 • PPP2CATCF7 1458 902 870 1478 • PITX2PPP2CA 1470 21 178 1170 • FZD2PPP2R1A 1485 934 255 1571 • CXXC4PPP2CA 1486 1664 1812 519 • CSNK1G1PPP2R1A 1488 197 1326 1100 • PPP2CATLE2 1494 885 979 1317 • DAAM1PPP2CA 1503 2133 479 874 • CSNK2A1PPP2CA 1518 619 267 2368 • LEF1PPP2R1A 1535 216 1308 1821 • PPP2R1ATCF7L1 1545 1262 645 1812 • GSK3BPPP2CA 1583 87 314 2018 • FZD7PPP2CA 1587 182 14 1295 • PPP2R1ATLE1 1591 1427 1468 1424 • NLKPPP2CA 1593 1246 2045 2100 • LRP6PPP2R1A 1607 469 706 1522 • PPP2CAFBXW4 1638 1289 1438 1147 • FBXW2PPP2CA 1655 1907 2065 255 • FBXW11PPP2CA 1657 1211 1377 1977 • PPP2CASLC9A3R1 1678 796 969 575 • FZD2PPP2CA 1680 645 836 2232 • PPP2CATLE1 1748 761 1876 1408 • PPP2R1ATLE2 1760 1681 772 653 • FZD1PPP2CA 1768 71 1303 897 • PPP2CAWNT4 1780 1490 2076 1897 • LEF1PPP2CA 1782 240 1117 750 • CSNK1G1PPP2CA 1783 226 1618 1998 • FGF4PPP2CA 1870 85 1102 1331 • PPP2CAWNT2 1894 1303 1953 1704 • PPP2R1AFBXW4 1924 765 1004 279 • PPP2R1AWNT4 1942 1592 1888 1148 • PPP2CAWIF1 1958 1618 954 711 • PPP2R1AWNT3A 1961 818 1181 1969 • PPP2R1ASLC9A3R1 1981 1345 1343 1705 • PPP2R1ATCF7 1997 760 139 2467 • LRP6PPP2CA 2058 454 2390 2064 • PPP2CAWNT3A 2075 1835 1372 1989 • PPP2R1AWNT2 2092 499 616 2337 • PPP2R1AWIF1 2117 1735 1784 1514 • PPP2CAWNT2B 2190 1500 1434 86 • PPP2R1AWNT1 2196 958 641 980 • PPP2R1AT 2237 1568 838 1631 • PPP2CAWNT1 2238 1737 2216 1735 • PPP2CAT 2241 1124 721 244 • PPP2R1APYGO1 2260 1339 1970 1154 • PPP2R1AWNT2B 2277 805 1121 1075 • PPP2R1ARHOU 2324 1615 1498 320 • PPP2R1AWNT3 2380 2201 187 972 • PPP2CAWNT3 2428 2139 668 1203 • PPP2CAPYGO1 2467 832 2064 1949 • PPP2CARHOU 2468 1536 1240 2094

## TLE family 2*^nd^* order interaction ranking at different time points

• SENP2TLE1 14 105 1586 1853 339 • FRZBTLE1 52 1914 1270 2055 1385 • SENP2TLE2 65 1060 407 2039 1987 • TLE2WNT1 136 2411 598 35 293 • FRZBTLE2 155 1945 691 980 887 • DKK1TLE1 188 2390 1622 198 975 • TLE2WIF1 199 2339 1706 2150 1057 • TLE2WNT3 226 902 2154 1496 381 • TLE1WNT1 321 340 67 1559 173 • APCTLE1 373 2454 2475 1710 638 • FBXW11TLE1 430 304 1816 1742 1435 • FOSL1TLE1 449 1415 951 694 213 • DKK1TLE2 461 2324 2057 361 2368 • FZD7TLE1 467 2 2068 1675 1468 • TLE1WIF1 476 2001 1609 1519 1740 • TCF7L1TLE1 489 1111 2064 2460 1758 • FZD1TLE1 520 1923 2420 871 1351 • FZD5TLE1 537 1320 2069 1695 205 • FRAT1TLE1 539 1849 2202 347 1273 • LRP5TLE1 542 1594 2248 1301 807 • PPP2R1ATLE1 558 294 1423 1061 1167 • DIXDC1TLE1 590 1921 256 2478 672 • CSNK2A1TLE1 620 1446 2133 1720 92 • CSNK1DTLE1 630 2120 2311 1500 981 • RHOUTLE1 640 2299 2397 773 2325 • TCF7TLE1 671 1145 1750 1990 1073 • NLKTLE1 677 1109 1929 651 479 • PPP2CATLE1 735 1046 1882 930 492 • GSK3BTLE1 763 1812 2324 909 1453 • CXXC4TLE1 769 1601 1273 1848 1735 • CCND3TLE1 791 112 2125 40 2430 • TLE1WNT3 810 11 1645 2057 740 • FZD6TLE1 827 1728 1874 2379 1271 • CTBP1TLE1 852 1966 1640 1284 691 • TLE2WNT2B 868 2258 250 1056 2479 • EP300TLE1 877 1379 1350 2295 1008 • TLE2WNT3A 888 904 1097 1079 914 • FZD1TLE2 911 602 706 2147 1208 • FZD8TLE1 927 998 585 882 1659 • TCF7L1TLE2 937 1563 1915 2448 1395 • APCTLE2 947 1088 1794 1499 1066 • FZD7TLE2 953 1822 1120 105 1033 • KREMEN1TLE1 957 1480 1158 855 387 • NKD1TLE1 963 42 2306 80 1215 • FBXW4TLE1 970 1784 1901 1274 1623 • SLC9A3R1TLE1 971 163 1785 2451 1590 • FBXW2TLE1 979 656 760 1673 300 • FRAT1TLE2 988 1078 926 1991 1592 • PITX2TLE1 993 2325 1561 54 341 • FOSL1TLE2 999 181 1002 2397 842 • DAAM1TLE1 1034 515 2075 728 1690 • FZD5TLE2 1049 1802 556 1329 945 • FBXW11TLE2 1067 1205 1951 2341 2338 • FSHBTLE1 1075 1418 2242 1568 685 • TLE2WNT2 1087 655 2168 1263 1088 • LRP5TLE2 1092 1148 2423 1857 1718 • TLE2WNT5A 1115 2032 399 1225 214 • TLE2WNT4 1147 1020 1085 1528 1368 • CSNK2A1TLE2 1168 865 2368 1782 2061 • CSNK1DTLE2 1183 2402 2131 591 1755 • PPP2R1ATLE2 1195 666 2197 2163 1732 • RHOUTLE2 1199 1280 1606 368 927 • SFRP4TLE1 1200 2485 918 674 82 • TCF7TLE2 1260 585 1207 1743 2426 • PPP2CATLE2 1268 1135 1602 279 2289 • GSK3BTLE2 1283 1525 1976 146 1277 • CTNNB1TLE1 1322 1891 2061 887 16 • DIXDC1TLE2 1345 45 236 1289 1254 • EP300TLE2 1361 1703 2379 571 1893 • FZD8TLE2 1410 932 595 1421 246 • KREMEN1TLE2 1414 787 1165 335 1052 • DVL2TLE1 1415 621 1556 472 2163 • AESTLE1 1429 1226 983 892 1392 • CSNK1G1TLE1 1433 2421 1551 1277 1177 • SLC9A3R1TLE2 1442 156 1137 2195 1492 • CXXC4TLE2 1456 987 1124 1836 2204 • TTLE1 1481 2277 2392 218 544 • NLKTLE2 1489 1752 2309 2199 2083 • TLE1WNT2 1499 1502 1074 1215 2110 • FBXW2TLE2 1508 2264 2140 2306 2372 • AXIN1TLE1 1520 877 1265 500 1669 • CTBP2TLE1 1522 614 1925 2028 1016 • TLE1WNT5A 1529 1631 281 940 279 • DAAM1TLE2 1530 950 2164 1287 2196 • PITX2TLE2 1553 1322 1576 1084 773 • FSHBTLE2 1586 980 1659 562 1619 • CTBP1TLE2 1605 1851 1089 2208 1960 • TLE1TLE2 1614 1089 315 1623 1524 • TLE1WNT4 1623 850 925 421 1525 • CTNNBIP1TLE1 1626 2429 930 922 326 • FZD6TLE2 1635 409 2076 987 2273 • CCND3TLE2 1644 1435 2034 801 2188 • TLE1WNT2B 1656 282 119 739 2178 • TLE1WNT3A 1670 1340 234 1389 2469 • FBXW4TLE2 1677 1048 1543 1129 1442 • SFRP4TLE2 1696 1917 1338 1026 344 • BTRCTLE1 1707 587 1836 110 801 • DVL1TLE1 1713 1127 770 1275 1371 • FGF4TLE1 1719 2134 1271 2109 1668 • LRP6TLE1 1756 1212 1504 282 934 • NKD1TLE2 1798 1615 1189 1112 2474 • JUNTLE1 1814 1948 1902 2090 83 • MYCTLE1 1827 1550 2370 915 916 • PYGO1TLE1 1852 840 2063 944 2119 • PORCNTLE1 1862 1220 1177 662 1841 • CSNK1G1TLE2 1874 806 1997 2261 1417 • CSNK1A1TLE1 1896 2238 1737 1397 307 • CTBP2TLE2 1921 2220 2037 1638 2097 • CTNNB1TLE2 1939 1445 1802 2315 1873 • FZD2TLE1 1946 1652 664 733 366 • DVL2TLE2 1971 714 764 33 1119 • FOXN1TLE1 1995 605 1776 1514 129 • CCND2TLE1 1998 2030 1826 849 748 • LEF1TLE1 2009 1169 1991 499 1513 • CTNNBIP1TLE2 2035 1081 1093 1216 282 • SFRP1TLE1 2050 1972 1708 524 1322 • LRP6TLE2 2064 389 528 913 127 • BCL9TLE1 2066 1775 1024 1105 313 • CCND1TLE1 2080 610 1516 1419 1246 • CSNK1A1TLE2 2087 1804 1201 432 448 • AXIN1TLE2 2131 2076 2127 2386 2055 • MYCTLE2 2143 1572 1809 1264 1488 • TTLE2 2205 1942 1665 1354 1853 • BTRCTLE2 2214 826 1880 376 1953 • FZD2TLE2 2238 1686 1532 8 185 • PYGO1TLE2 2266 597 2346 645 421 • LEF1TLE2 2272 1113 2219 606 932 • FOXN1TLE2 2277 1368 1261 663 1625 • PORCNTLE2 2282 2016 1840 336 1859 • AESTLE2 2294 1927 1745 761 2239 • SFRP1TLE2 2295 1370 835 1469 1900 • DVL1TLE2 2310 1369 219 2 1648 • GSK3ATLE1 2339 1999 1386 1134 438 • JUNTLE2 2345 376 1517 1338 277 • FGF4TLE2 2356 1543 395 104 1872 • CCND2TLE2 2368 1333 1276 2175 1910 • BCL9TLE2 2391 131 1220 214 2369 • GSK3ATLE2 2433 807 1873 2022 702 • CCND1TLE2 2434 1503 1743 1501 551

## TLE family 2*^nd^* order interaction ranking at different durations

• RHOUTLE2 37 538 1122 2277 • RHOUTLE1 56 914 982 28 • FZD5TLE1 100 2328 1393 1765 • CSNK1A1TLE2 105 598 1624 1838 • PYGO1TLE2 110 1446 2384 2164 • CSNK1A1TLE1 130 1353 1227 84 • FZD5TLE2 192 638 1763 853 • PYGO1TLE1 245 1445 1639 698 • CTNNB1TLE2 278 1523 805 1417 • PORCNTLE1 299 2311 1860 1335 • CTBP1TLE2 306 2008 1500 1135 • DIXDC1TLE2 311 74 778 163 • JUNTLE2 322 948 2299 198 • DIXDC1TLE1 326 294 1278 2476 • CTNNB1TLE1 327 1631 2325 1389 • CCND2TLE2 335 1683 1217 193 • PORCNTLE2 372 793 1849 577 • AXIN1TLE2 412 1123 1649 230 • CCND1TLE2 421 575 689 492 • AESTLE2 458 2148 1627 1978 • TTLE2 491 1146 2097 527 • TTLE1 533 1098 1413 199 • CTNNBIP1TLE2 545 2246 544 164 • FRAT1TLE1 567 2230 955 832 • BCL9TLE1 599 2397 1085 1527 • CSNK1DTLE2 619 336 2442 2173 • CCND2TLE1 622 1192 1552 1528 • BCL9TLE2 633 2061 2349 298 • KREMEN1TLE1 635 794 245 558 • FSHBTLE2 645 358 1740 126 • CTBP1TLE1 679 1071 856 1714 • AXIN1TLE1 690 992 1388 816 • CTNNBIP1TLE1 696 2282 871 1561 • CCND1TLE1 710 959 1079 1895 • MYCTLE2 719 539 2267 326 • BTRCTLE2 723 1147 2355 823 • KREMEN1TLE2 737 1823 897 1269 • FRAT1TLE2 746 1570 1544 934 • FOXN1TLE2 758 2416 602 1810 • JUNTLE1 775 1669 1931 622 • CSNK1DTLE1 799 983 2430 1224 • GSK3ATLE2 807 428 599 408 • SLC9A3R1TLE2 817 437 692 124 • DVL1TLE2 819 1932 575 200 • AESTLE1 822 2141 2222 646 • FSHBTLE1 856 731 2354 1743 • APCTLE2 859 1030 1477 812 • MYCTLE1 898 1792 1428 803 • FOXN1TLE1 913 1450 1819 583 • GSK3ATLE1 940 1135 1107 1028 • SLC9A3R1TLE1 945 1508 2080 2165 • DVL1TLE1 963 1316 1113 2026 • FOSL1TLE1 968 845 1403 671 • BTRCTLE1 973 2069 2128 471 • TLE1WNT5A 978 366 1065 1624 • FOSL1TLE2 982 332 362 1109 • TLE2WNT5A 999 836 1152 165 • CTBP2TLE1 1079 2019 2081 1050 • LRP5TLE2 1091 234 880 394 • NKD1TLE2 1174 2109 1720 1111 • CTBP2TLE2 1188 1129 1300 560 • FBXW4TLE2 1199 2299 1778 483 • APCTLE1 1222 1677 1760 1940 • CCND3TLE2 1227 2157 1447 1277 • FRZBTLE2 1228 2226 992 642 • TLE1TLE2 1231 442 1418 340 • DVL2TLE2 1272 1379 2241 170 • EP300TLE1 1351 842 2476 1650 • EP300TLE2 1355 1332 1807 66 • TLE1WNT4 1406 285 919 1214 • LRP5TLE1 1413 621 1572 65 • TCF7TLE2 1428 1516 1010 718 • NKD1TLE1 1436 1062 2220 1672 • TCF7TLE1 1459 2288 1891 609 • CXXC4TLE2 1478 2360 2041 420 • FBXW4TLE1 1487 1575 2428 1552 • FBXW2TLE2 1491 1483 1917 1340 • PPP2CATLE2 1494 885 979 1317 • FRZBTLE1 1497 1141 1263 226 • DAAM1TLE1 1498 2146 2403 1151 • DVL2TLE1 1507 1663 1855 1892 • DKK1TLE1 1540 416 1298 605 • FBXW11TLE2 1543 881 1400 343 • SFRP1TLE2 1544 133 1995 210 • GSK3BTLE2 1548 179 545 1574 • FZD8TLE2 1552 88 873 280 • NLKTLE2 1564 2478 958 376 • TLE2WNT4 1567 601 1484 205 • CCND3TLE1 1589 1937 1844 1235 • PPP2R1ATLE1 1591 1427 1468 1424 • PITX2TLE2 1618 86 480 1004 • CSNK2A1TLE2 1628 2440 335 1090 • FZD8TLE1 1635 277 375 1244 • DKK1TLE2 1637 257 1391 417 • FZD1TLE1 1648 755 1742 822 • FZD6TLE1 1654 2272 220 1091 • DAAM1TLE2 1686 2427 1044 405 • TLE2WNT2 1690 1505 2444 1493 • FZD7TLE2 1712 313 167 227 • CXXC4TLE1 1715 1684 2326 901 • TCF7L1TLE2 1725 388 2376 1500 • FZD6TLE2 1739 837 231 67 • NLKTLE1 1740 1892 1041 1130 • PPP2CATLE1 1748 761 1876 1408 • PPP2R1ATLE2 1760 1681 772 653 • PITX2TLE1 1762 312 792 63 • TLE1WIF1 1771 1069 1811 1906 • FZD2TLE2 1777 564 1064 75 • TLE1WNT2 1799 530 1521 2037 • FZD2TLE1 1810 1739 380 1200 • FBXW11TLE1 1815 1035 1729 912 • LEF1TLE2 1849 249 1175 80 • GSK3BTLE1 1850 535 894 552 • FGF4TLE2 1856 337 2475 529 • TLE1WNT3A 1877 720 1230 2350 • LRP6TLE1 1893 769 791 818 • FZD7TLE1 1914 998 318 2340 • FZD1TLE2 1934 222 868 1024 • TLE2WNT3A 1977 2064 1916 525 • FBXW2TLE1 1987 2337 1408 1273 • CSNK2A1TLE1 2014 1734 554 1041 • TLE2WNT1 2020 1558 1478 311 • TLE2WIF1 2071 2322 1203 13 • TCF7L1TLE1 2081 659 682 938 • SFRP1TLE1 2105 742 1998 1081 • LRP6TLE2 2129 458 2377 209 • FGF4TLE1 2142 452 1415 1878 • SFRP4TLE2 2166 341 1984 182 • CSNK1G1TLE2 2175 152 1378 131 • TLE1WNT2B 2179 604 401 240 • TLE1WNT1 2218 259 1223 1601 • TLE1WNT3 2245 2057 181 1864 • SFRP4TLE1 2247 488 1838 645 • SENP2TLE1 2275 1096 812 2033 • TLE2WNT3 2280 2014 297 7 • LEF1TLE1 2283 710 1589 1689 • SENP2TLE2 2303 607 1471 121 • TLE2WNT2B 2316 2278 398 17 • CSNK1G1TLE1 2457 471 1132 1909

## AES 2*^nd^* order interaction ranking at different time points

• AESGSK3A 501 178 692 1730 595 • AESCCND1 528 1970 2090 2314 119 • AESSFRP1 682 39 1841 1685 670 • AESFGF4 740 1165 1770 1003 514 • AESJUN 772 1512 1373 333 1826 • AESFZD2 819 1996 1256 1865 1331 • AESWIF1 844 523 2048 324 1278 • AESAXIN1 918 309 1710 24 2292 • AESBCL9 935 1124 481 1952 2271 • AESMYC 955 1847 866 789 2021 • AESDVL1 997 438 2201 175 1178 • AESPORCN 1017 835 1613 605 877 • AESCCND2 1021 220 1190 68 1752 • AESCSNK1A1 1025 473 834 816 1909 • AESFOXN1 1031 390 331 1815 1155 • AESLEF1 1072 559 915 471 1743 • AESWNT1 1134 631 1063 1182 299 • AESLRP6 1150 799 560 2420 922 • AESCTBP2 1167 177 985 1059 899 • AESPYGO1 1224 1765 903 2046 1015 • AESNKD1 1279 146 1696 725 1774 • AESDVL2 1282 779 2189 2469 1839 • AEST 1369 532 824 2252 2250 • AESTLE1 1429 1226 983 892 1392 • AESWNT3 1443 170 2054 1694 1166 • AESBTRC 1450 74 647 92 2226 • AESCTNNB1 1537 1581 2389 285 2177 • AESCTNNBIP1 1548 1283 1569 253 1516 • AESFBXW4 1663 2036 1157 97 2185 • AESFZD6 1673 270 841 1209 1510 • AESFSHB 1694 723 374 2191 1809 • AESCSNK1G1 1715 1221 552 2274 1160 • AESWNT4 1722 1434 1595 1017 515 • AESSLC9A3R1 1723 448 1160 73 911 • AESDAAM1 1769 145 928 647 1001 • AESPITX2 1770 938 1100 1594 1174 • AESPPP2CA 1772 1246 501 303 1200 • AESFBXW2 1783 926 230 1372 1624 • AESCTBP1 1837 2190 2478 216 1514 • AESCSNK1D 1851 1193 1278 2036 1569 • AESWNT2 1872 935 1402 127 1599 • AESEP300 1886 1033 1932 369 2086 • AESDIXDC1 1985 335 1831 469 2059 • AESCSNK2A1 1989 554 1143 2264 2167 • AESCCND3 1991 399 567 1622 2209 • AESWNT3A 2010 1259 1940 2345 573 • AESFZD7 2028 627 2135 2269 1895 • AESKREMEN1 2032 1514 690 927 526 • AESWNT2B 2058 86 549 1783 1330 • AESWNT5A 2062 1767 1221 243 1517 • AESSFRP4 2081 1869 2339 842 1527 • AESFBXW11 2084 786 1246 1608 1398 • AESLRP5 2101 816 1066 1634 1106 • AESFOSL1 2138 2079 2245 140 1374 • AESFZD8 2150 1001 1911 1360 2052 • AESNLK 2187 1778 1410 328 1974 • AESCXXC4 2192 416 1733 1197 1929 • AESFZD5 2204 1086 2290 1160 2473 • AESPPP2R1A 2210 125 936 1790 453 • AESTCF7L1 2224 634 1176 407 1824 • AESFZD1 2235 899 1384 276 497 • AESRHOU 2236 2208 2249 1240 1399 • AESFRAT1 2245 923 1761 207 1454 • AESTCF7 2276 162 422 1436 609 • AESTLE2 2294 1927 1745 761 2239 • AESGSK3B 2300 1473 2469 1728 1878 • AESAPC 2355 1062 2482 537 940 • AESDKK1 2445 1411 642 477 2406 • AESFRZB 2451 1899 2016 307 199 • AESSENP2 2483 115 2078 269 1239

## AES 2*^nd^* order interaction ranking at different durations

• AESSENP2 33 954 282 46 • AESSFRP4 58 2200 1366 35 • AESCSNK1G1 123 2086 911 156 • AESPPP2R1A 195 1651 1416 1431 • AESLEF1 203 2245 1697 331 • AESFZD2 215 2198 957 283 • AESGSK3B 216 2187 924 892 • AESFZD1 244 1247 2121 805 • AESFBXW11 253 1993 662 989 • AESFGF4 260 1245 1191 183 • AESTCF7L1 276 2332 1834 1985 • AESLRP6 281 2208 1584 89 • AESPITX2 304 1468 1126 1001 • AESWNT5A 312 1576 2187 837 • AESSFRP1 329 1881 2274 82 • AESCSNK2A1 339 1702 2257 548 • AESFZD7 362 1498 826 219 • AESFZD8 374 1261 1517 984 • AESFBXW2 376 758 647 339 • AESCXXC4 430 461 502 322 • AESFZD6 436 2159 1823 9 • AESTLE2 458 2148 1627 1978 • AESDAAM1 485 2042 1523 10 • AESNLK 501 1269 320 863 • AESTCF7 539 739 257 1757 • AESPPP2CA 576 1581 1365 2479 • AESFBXW4 628 715 966 442 • AESCCND3 665 624 1842 908 • AESDKK1 676 2263 948 1097 • AESEP300 727 695 1871 285 • AESWNT4 741 1309 1577 127 • AESDVL2 765 2385 2404 31 • AESFRZB 804 1002 742 1545 • AESTLE1 822 2141 2222 646 • AESAPC 849 384 166 1164 • AESFOSL1 885 2006 1353 358 • AESLRP5 926 1442 2000 1369 • AESGSK3A 1000 2310 1753 290 • AESBTRC 1001 1073 2315 1158 • AESWNT1 1032 1313 1599 1475 • AESWNT2 1046 2317 1197 144 • AESNKD1 1066 1370 149 676 • AESMYC 1071 819 406 484 • AESKREMEN1 1075 2030 1216 997 • AESCTBP2 1097 1785 779 256 • AESFRAT1 1141 2113 823 2310 • AESFSHB 1165 924 1244 68 • AESFOXN1 1167 1011 819 1449 • AESWIF1 1230 1537 1556 6 • AESCTNNBIP1 1254 536 2316 45 • AEST 1306 2417 1843 1086 • AESDVL1 1307 618 2309 36 • AESCSNK1D 1332 2377 1540 1364 • AESSLC9A3R1 1377 2111 1352 74 • AESCCND2 1379 764 308 39 • AESBCL9 1388 2443 1712 176 • AESJUN 1422 2439 968 1401 • AESCCND1 1460 2180 2202 268 • AESAXIN1 1496 1787 1949 49 • AESWNT2B 1514 2339 730 30 • AESWNT3A 1576 1763 2024 987 • AESWNT3 1659 1807 644 179 • AESCTBP1 1751 689 97 1252 • AESPYGO1 1755 2062 2278 2087 • AESPORCN 1847 1778 2364 208 • AESRHOU 2048 2021 2090 2177 • AESDIXDC1 2097 1790 2344 215 • AESCTNNB1 2106 229 1219 721 • AESCSNK1A1 2289 516 233 1993 • AESFZD5 2410 2183 199 21

## APC 2*^nd^* order interaction ranking at different time points

• APCGSK3A 30 1591 124 1708 913 • APCSFRP1 39 2431 1945 2009 455 • APCFGF4 41 1592 1827 1895 255 • APCCCND2 59 1759 1426 785 603 • APCJUN 61 1317 1857 701 2309 • APCCCND1 81 2340 1295 1390 21 • APCBTRC 103 2024 259 556 163 • APCWNT1 104 2384 80 365 29 • APCMYC 116 2248 619 1855 2105 • APCWNT3 130 1907 1291 1399 86 • APCWIF1 148 2463 1119 1778 550 • APCLEF1 157 470 1387 387 374 • APCPYGO1 169 1188 303 604 717 • APCFZD2 187 843 2115 567 1048 • APCFOXN1 195 2386 434 436 884 • APCPORCN 216 1139 1531 1249 245 • APCCTBP2 217 2410 2039 826 1884 • APCCSNK1A1 235 2035 684 47 500 • APCDVL1 239 2352 1209 203 1920 • APCT 248 1850 282 2337 322 • APCBCL9 260 1293 1643 246 591 • APCLRP6 305 1753 934 905 689 • APCDVL2 307 2041 994 189 1402 • APCCSNK1G1 332 1558 809 2204 260 • APCAXIN1 347 484 1125 748 1211 • APCDAAM1 372 2361 352 448 93 • APCTLE1 373 2454 2475 1710 638 • APCFSHB 399 1318 278 420 962 • APCCTNNBIP1 420 2177 2464 1656 2002 • APCNKD1 454 1839 379 1824 1283 • APCCTNNB1 457 1988 862 1686 412 • APCFZD6 480 2229 189 480 971 • APCPITX2 490 1292 2334 1843 259 • APCCTBP1 519 1607 2015 2093 2342 • APCFBXW2 562 2434 767 710 154 • APCFBXW4 567 1833 257 2238 402 • APCSFRP4 580 647 2303 474 489 • APCPPP2CA 586 2263 452 388 1224 • APCKREMEN1 604 1973 986 1247 357 • APCCCND3 610 1689 125 1193 204 • APCFZD8 624 2116 2160 1058 316 • APCFZD5 637 729 1301 1851 1439 • APCEP300 642 952 1360 1189 1384 • APCTCF7 665 1986 177 2281 37 • APCWNT2 680 243 1022 908 910 • APCGSK3B 700 2255 1553 1445 525 • APCWNT3A 701 1398 1674 2409 614 • APCFRAT1 704 2355 1872 1937 2391 • APCNLK 710 2319 95 643 256 • APCCXXC4 712 275 759 201 425 • APCSLC9A3R1 717 1487 2313 1786 643 • APCWNT2B 752 1568 319 2415 1418 • APCWNT4 754 694 1992 833 2221 • APCPPP2R1A 777 1312 425 667 138 • APCDIXDC1 786 1409 2136 1919 684 • APCCSNK1D 808 2236 514 1814 630 • APCWNT5A 847 2159 391 2305 106 • APCLRP5 861 1079 1027 2277 496 • APCFOSL1 882 695 2432 55 2093 • APCTLE2 947 1088 1794 1499 1066 • APCFZD7 950 2406 1625 1367 903 • APCCSNK2A1 1012 2069 723 1914 947 • APCFZD1 1065 1913 1395 525 1255 • APCFBXW11 1090 2348 447 744 1026 • APCRHOU 1106 832 1098 1831 709 • APCTCF7L1 1174 1819 1610 1801 974 • APCDKK1 1940 2098 323 1332 1954 • APCFRZB 2106 958 2211 17 1291 • AESAPC 2355 1062 2482 537 940 • APCSENP2 2371 2432 655 1128 287

## APC 2*^nd^* order interaction ranking at different durations

• APCSENP2 87 1074 2100 521 • APCSFRP4 166 1128 2091 549 • APCFGF4 363 1156 990 2062 • APCCSNK1G1 407 1533 2295 2081 • APCLEF1 428 1381 1542 1758 • APCWNT5A 477 2063 1686 1488 • APCFZD2 478 1569 673 2040 • APCLRP6 495 1624 1270 1708 • APCFBXW11 542 2199 1851 2444 • APCPPP2R1A 554 1481 1501 2010 • APCGSK3B 555 1188 386 1681 • APCCSNK2A1 563 2142 670 2441 • APCFZD7 577 1380 609 1639 • APCFBXW2 581 2325 2146 1662 • APCFZD1 594 1194 1594 719 • APCCXXC4 636 2103 2254 1255 • APCFZD8 658 1045 586 2449 • APCTCF7L1 684 1317 2093 2313 • APCDKK1 687 778 2468 1392 • APCSFRP1 688 1189 1002 631 • APCPITX2 700 1057 705 2385 • APCNLK 735 1731 1988 2484 • APCFZD6 762 1232 971 886 • APCDAAM1 776 1826 2022 1464 • APCTCF7 815 1780 1652 1231 • APCPPP2CA 824 1772 1975 2014 • AESAPC 849 384 166 1164 • APCTLE2 859 1030 1477 812 • APCFBXW4 904 2149 1575 1957 • APCFRZB 939 2436 2327 1767 • APCWNT4 992 2228 485 2391 • APCDVL2 1081 2402 1596 896 • APCEP300 1094 1040 1047 1994 • APCNKD1 1147 1784 1355 1815 • APCCCND3 1163 2300 2199 2447 • APCLRP5 1169 1547 2286 374 • APCTLE1 1222 1677 1760 1940 • APCBTRC 1304 2091 1557 1842 • APCCTBP2 1382 1196 1913 1885 • APCSLC9A3R1 1440 936 1444 1907 • APCBCL9 1515 1878 797 2154 • APCFOSL1 1521 1235 2345 923 • APCCTNNBIP1 1547 1343 848 909 • APCWNT2 1549 1597 1758 1012 • APCWNT3A 1561 1125 798 2294 • APCWNT1 1590 654 2423 2420 • APCWIF1 1624 1777 1882 1868 • APCKREMEN1 1633 2163 1095 804 • APCGSK3A 1639 1670 885 1587 • APCAXIN1 1640 1769 1929 2141 • APCFOXN1 1666 1886 1427 1261 • APCCSNK1D 1698 899 1960 1653 • APCDVL1 1699 2484 679 843 • APCCCND1 1717 1281 931 2187 • APCJUN 1761 1308 1870 1567 • APCMYC 1766 2369 830 2290 • APCCTNNB1 1772 1467 1313 791 • APCFRAT1 1822 1743 2367 2382 • APCCTBP1 1828 600 1912 2101 • APCT 1906 2077 1746 1137 • APCFSHB 1930 1760 473 771 • APCWNT2B 1982 1256 1309 599 • APCCSNK1A1 2042 1909 825 246 • APCCCND2 2102 1187 2198 1979 • APCDIXDC1 2140 1406 729 1381 • APCPYGO1 2156 1482 2439 1865 • APCWNT3 2332 1469 1276 801 • APCRHOU 2362 1347 564 425 • APCPORCN 2372 2366 960 2024 • APCFZD5 2414 1853 1429 1032

## AXIN1 2*^nd^* order interaction ranking at different time points

• APCAXIN1 347 484 1125 748 1211 • AXIN1GSK3A 458 1443 527 255 342 • AXIN1LEF1 497 1330 1519 853 2262 • AXIN1FOXN1 546 1824 313 117 2424 • AXIN1CSNK1A1 551 1603 1908 2244 2014 • AXIN1JUN 572 1361 1187 1331 1459 • AXIN1MYC 584 2418 696 1253 700 • AXIN1FZD2 655 87 1678 156 2200 • AXIN1PYGO1 656 1745 1135 85 778 • AXIN1BCL9 674 794 2404 1518 1153 • AXIN1CCND1 692 2087 2328 411 22 • AXIN1SFRP1 751 1338 1727 780 437 • AXIN1FGF4 753 2106 2425 1775 436 • AXIN1WNT1 761 1943 285 465 1099 • AXIN1WIF1 854 2360 2326 1242 11 • AXIN1PORCN 864 2194 1102 851 378 • AXIN1CCND2 893 1509 1259 1046 1061 • AESAXIN1 918 309 1710 24 2292 • AXIN1WNT3 969 1665 901 1257 1144 • AXIN1DVL1 1042 2472 1812 751 1105 • AXIN1CSNK1G1 1141 892 1008 354 882 • AXIN1NKD1 1186 2091 875 1757 1997 • AXIN1BTRC 1214 2378 748 309 619 • AXIN1T 1256 1174 1016 771 1243 • AXIN1CTNNBIP1 1263 1198 771 1507 2441 • AXIN1CTBP2 1306 1987 1043 1989 1636 • AXIN1DVL2 1307 1493 1264 242 1425 • AXIN1CTNNB1 1413 1167 2074 131 1769 • AXIN1LRP6 1428 2138 844 543 1567 • AXIN1TLE1 1520 877 1265 500 1669 • AXIN1EP300 1536 937 1628 2063 1171 • AXIN1PITX2 1569 2063 1594 100 652 • AXIN1FBXW4 1573 1495 929 1150 1183 • AXIN1CCND3 1603 1901 857 621 673 • AXIN1CTBP1 1610 934 1728 712 1414 • AXIN1FSHB 1643 2251 475 182 2087 • AXIN1FBXW2 1708 1181 2206 752 269 • AXIN1SFRP4 1729 1199 2471 190 954 • AXIN1LRP5 1740 628 396 56 2062 • AXIN1WNT3A 1773 1277 2237 721 1448 • AXIN1FZD6 1809 866 730 449 227 • AXIN1NLK 1812 2363 493 254 1383 • AXIN1KREMEN1 1842 374 2232 862 719 • AXIN1SLC9A3R1 1857 744 2455 2098 2319 • AXIN1WNT2 1865 696 1359 399 734 • AXIN1PPP2CA 1868 2296 506 827 2213 • AXIN1FZD8 1914 1213 1197 1211 583 • AXIN1PPP2R1A 1920 1290 1491 732 394 • AXIN1WNT4 1934 800 1403 2431 2096 • AXIN1FOSL1 1948 2100 1224 736 892 • AXIN1DAAM1 1952 1095 1829 808 491 • AXIN1WNT5A 2006 1266 1106 330 1195 • AXIN1CSNK1D 2012 428 1128 252 926 • AXIN1TCF7 2051 2104 1150 798 1600 • AXIN1GSK3B 2059 1490 1194 161 864 • AXIN1CSNK2A1 2069 888 1233 1238 1822 • AXIN1DIXDC1 2120 1676 1691 2249 228 • AXIN1FZD7 2125 1417 1624 1705 1390 • AXIN1TLE2 2131 2076 2127 2386 2055 • AXIN1RHOU 2133 1729 1249 211 979 • AXIN1FZD5 2137 1252 1853 1875 2449 • AXIN1CXXC4 2142 1066 1671 2325 796 • AXIN1FRAT1 2145 1466 1503 72 1991 • AXIN1WNT2B 2167 547 115 1108 828 • AXIN1TCF7L1 2194 1538 1356 497 1783 • AXIN1FZD1 2218 1719 1789 1081 1050 • AXIN1FBXW11 2228 2061 613 458 924 • AXIN1DKK1 2432 2362 494 916 1397 • AXIN1FRZB 2440 1308 2256 1478 1244 • AXIN1SENP2 2478 1724 2427 565 815

## AXIN1 2*^nd^* order interaction ranking at different durations

• AXIN1SENP2 50 1039 321 2172 • AXIN1SFRP4 106 2009 596 794 • AXIN1LRP6 141 1869 1295 2249 • AXIN1CSNK2A1 171 657 2004 2117 • AXIN1FBXW11 209 725 557 2148 • AXIN1FZD1 235 1911 372 1003 • AXIN1FBXW2 254 939 416 2093 • AXIN1FZD2 274 806 2054 2209 • AXIN1CXXC4 287 141 606 1717 • AXIN1FZD7 294 2384 1364 2039 • AXIN1FGF4 300 2376 749 1438 • AXIN1FZD8 303 2211 2129 1804 • AXIN1CSNK1G1 305 1001 1049 1873 • AXIN1SFRP1 315 2423 2110 2194 • AXIN1LEF1 317 2205 1021 2253 • AXIN1PPP2R1A 320 965 796 2133 • AXIN1TCF7L1 345 1895 744 1140 • AXIN1PPP2CA 351 923 977 2031 • AXIN1WNT5A 364 365 667 2107 • AXIN1GSK3B 390 2351 2353 34 • AXIN1TLE2 412 1123 1649 230 • AXIN1PITX2 439 2070 1090 1030 • AXIN1NLK 465 301 189 1666 • AXIN1DAAM1 508 870 2469 2399 • AXIN1FBXW4 544 628 760 1344 • AXIN1FZD6 548 1590 2058 1477 • AXIN1DVL2 595 232 1694 2378 • AXIN1TCF7 620 574 357 1827 • AXIN1TLE1 690 992 1388 816 • AXIN1WNT4 780 637 1845 753 • AXIN1SLC9A3R1 794 615 734 2127 • AXIN1FRZB 797 404 450 284 • AXIN1DKK1 801 1988 633 1565 • AXIN1CCND3 827 547 422 2259 • AXIN1LRP5 848 2156 1997 372 • AXIN1CTBP2 876 1471 625 1807 • AXIN1EP300 900 273 1907 1459 • AXIN1WNT1 951 675 1625 1503 • AXIN1BTRC 1025 1201 1150 1304 • AXIN1CSNK1D 1049 1610 762 344 • AXIN1CCND1 1052 1090 842 2122 • AXIN1NKD1 1076 308 99 1423 • AXIN1WNT2 1080 513 330 1454 • AXIN1GSK3A 1085 2434 1529 1935 • AXIN1FOSL1 1092 2305 840 424 • AXIN1BCL9 1118 869 395 2261 • AXIN1JUN 1128 493 884 134 • AXIN1DVL1 1130 520 1006 1326 • AXIN1KREMEN1 1201 507 1248 1196 • AXIN1FSHB 1202 1531 489 1167 • AXIN1CCND2 1236 167 258 1622 • AXIN1WIF1 1257 1815 754 1391 • AXIN1FRAT1 1284 823 522 885 • AXIN1FOXN1 1337 830 1586 940 • AXIN1MYC 1421 880 433 2212 • AESAXIN1 1496 1787 1949 49 • AXIN1T 1517 1148 681 390 • AXIN1DIXDC1 1575 2307 1872 2225 • AXIN1WNT2B 1622 2431 915 2251 • APCAXIN1 1640 1769 1929 2141 • AXIN1CTNNBIP1 1683 323 1167 761 • AXIN1WNT3 1743 2241 533 1753 • AXIN1RHOU 1788 2233 2296 87 • AXIN1CTBP1 1816 79 237 1824 • AXIN1CTNNB1 1830 171 1675 1611 • AXIN1WNT3A 1853 2007 1288 703 • AXIN1CSNK1A1 2031 386 17 633 • AXIN1PORCN 2181 820 872 2096 • AXIN1PYGO1 2211 1478 1653 327 • AXIN1FZD5 2388 1177 327 1418

## BCL9 2*^nd^* order interaction ranking at different time points

• APCBCL9 260 1293 1643 246 591 • AXIN1BCL9 674 794 2404 1518 1153 • BCL9GSK3A 790 1599 412 1164 1409 • BCL9CCND1 922 2093 1048 1093 257 • AESBCL9 935 1124 481 1952 2271 • BCL9PYGO1 1100 974 1065 2104 2475 • BCL9SFRP1 1146 1071 2271 1073 1766 • BCL9FGF4 1156 1123 1483 1610 578 • BCL9WIF1 1177 1985 1835 135 1691 • BCL9FZD2 1178 301 1726 804 2019 • BCL9T 1225 758 836 232 1540 • BCL9MYC 1226 2114 284 1799 893 • BCL9JUN 1230 1682 1170 337 968 • BCL9WNT3 1300 975 2477 1374 1197 • BCL9BTRC 1354 1597 439 1557 727 • BCL9FOXN1 1374 2250 167 985 951 • BCL9PORCN 1382 1800 456 2058 1198 • BCL9LEF1 1398 2308 2327 13 1981 • BCL9WNT1 1469 2265 573 267 599 • BCL9LRP6 1485 1054 408 2095 546 • BCL9DVL1 1551 2000 2283 1032 1860 • BCL9CSNK1A1 1578 1735 942 1732 1184 • BCL9CCND2 1628 1965 479 2352 351 • BCL9DVL2 1689 1667 649 71 2060 • BCL9CTBP2 1737 1797 2123 2120 1159 • BCL9CSNK1G1 1787 924 301 1978 439 • BCL9FZD6 1836 2330 446 222 1969 • BCL9FSHB 1869 114 139 1086 1261 • BCL9CTNNBIP1 1906 25 1390 569 1796 • BCL9KREMEN1 1954 364 828 2037 996 • BCL9CTNNB1 1970 1783 2050 2128 110 • BCL9NKD1 2003 2247 711 1555 2396 • BCL9SLC9A3R1 2021 1294 1961 2154 2348 • BCL9EP300 2036 1788 1416 1221 2045 • BCL9FBXW4 2038 2477 677 2031 1956 • BCL9TCF7 2044 1809 133 1660 521 • BCL9SFRP4 2048 951 2252 950 1112 • BCL9GSK3B 2060 1306 2119 264 2409 • BCL9TLE1 2066 1775 1024 1105 313 • BCL9FZD8 2070 1621 1354 1347 2199 • BCL9PPP2CA 2094 1602 402 2250 1423 • BCL9FBXW2 2119 1989 436 1944 917 • BCL9CCND3 2159 2140 519 1909 471 • BCL9CTBP1 2165 1690 1603 59 1829 • BCL9FZD5 2180 970 1736 2137 2169 • BCL9CSNK2A1 2198 2237 950 2416 2015 • BCL9CXXC4 2208 255 1028 788 729 • BCL9WNT2 2211 873 1683 379 1635 • BCL9TCF7L1 2213 2196 1230 1013 1060 • BCL9WNT4 2220 2084 458 1953 1187 • BCL9DAAM1 2230 1810 1520 2209 1336 • BCL9NLK 2231 1971 657 431 822 • BCL9FRAT1 2239 325 1811 125 2380 • BCL9DIXDC1 2256 578 1828 1307 2035 • BCL9PITX2 2257 1974 2351 1983 1443 • BCL9PPP2R1A 2275 2457 416 2074 2116 • BCL9CSNK1D 2281 1823 1453 1126 2433 • BCL9WNT3A 2313 1496 1687 2061 2379 • BCL9LRP5 2317 1121 1199 2083 2075 • BCL9WNT2B 2336 996 305 2038 2467 • BCL9WNT5A 2340 1646 1252 2072 918 • BCL9FZD7 2346 654 1757 163 522 • BCL9FOSL1 2383 144 2080 1195 312 • BCL9TLE2 2391 131 1220 214 2369 • BCL9RHOU 2395 2416 2354 1826 1546 • BCL9FZD1 2396 1764 1144 1214 2193 • BCL9FBXW11 2430 1009 638 1977 2079 • BCL9DKK1 2452 543 666 1123 1799 • BCL9FRZB 2455 421 2371 138 1481 • BCL9SENP2 2466 1992 1408 907 1387

## BCL9 2*^nd^* order interaction ranking at different durations

• BCL9SENP2 66 1598 1024 1668 • BCL9SFRP4 103 1549 703 850 • BCL9LRP6 186 1742 1832 930 • BCL9FGF4 198 1958 1347 1731 • BCL9LEF1 224 1224 1136 1899 • BCL9CSNK1G1 230 2247 1677 1582 • BCL9FBXW11 272 1351 2463 1550 • BCL9PPP2R1A 275 773 2409 1630 • BCL9FZD7 295 1430 560 103 • BCL9CSNK2A1 308 1231 860 1064 • BCL9PITX2 321 1144 1585 1405 • BCL9FBXW2 324 1009 1993 657 • BCL9FZD1 325 2268 2333 1070 • BCL9SFRP1 334 1270 2340 785 • BCL9WNT5A 354 2155 2483 2036 • BCL9FZD2 382 1168 2194 1053 • BCL9CXXC4 399 412 675 123 • BCL9NLK 427 1049 283 955 • BCL9TCF7L1 448 2393 1787 2073 • BCL9DAAM1 466 1845 1655 2414 • BCL9FZD6 483 1588 1151 819 • BCL9FZD8 518 2303 1437 2112 • BCL9GSK3B 558 2323 1604 795 • BCL9TLE1 599 2397 1085 1527 • BCL9TLE2 633 2061 2349 298 • BCL9FBXW4 634 1065 1857 2009 • BCL9CCND3 655 631 2391 1263 • BCL9FRZB 672 1025 974 675 • BCL9PPP2CA 675 1821 2150 1300 • BCL9DKK1 726 2261 1442 2163 • BCL9NKD1 751 705 500 2254 • BCL9WNT4 774 986 1966 318 • BCL9TCF7 795 1286 352 1779 • BCL9WNT2 860 912 1668 2129 • BCL9EP300 872 495 2201 1088 • BCL9DVL2 910 1047 1987 2358 • BCL9CTBP2 961 1574 1703 740 • BCL9LRP5 1009 1999 1670 857 • BCL9FOSL1 1078 1403 1315 364 • AXIN1BCL9 1118 869 395 2261 • BCL9CCND1 1125 1646 709 1420 • BCL9SLC9A3R1 1157 1357 1825 1871 • BCL9DVL1 1187 809 1412 611 • BCL9WNT1 1195 1992 2441 441 • BCL9FSHB 1196 2413 1642 1694 • BCL9CCND2 1215 528 2079 1384 • BCL9GSK3A 1241 2399 1908 778 • BCL9CSNK1D 1288 2470 2263 1788 • BCL9WIF1 1303 1686 1818 462 • BCL9FOXN1 1325 2078 2177 1802 • BCL9JUN 1335 2207 1555 1025 • BCL9FRAT1 1352 1540 2268 741 • AESBCL9 1388 2443 1712 176 • BCL9MYC 1414 2320 1264 1296 • BCL9BTRC 1431 2238 2276 1484 • BCL9WNT3A 1433 890 1435 1866 • BCL9CTNNBIP1 1471 1068 1332 1504 • APCBCL9 1515 1878 797 2154 • BCL9KREMEN1 1524 1691 1330 1663 • BCL9CTBP1 1536 402 2119 1928 • BCL9WNT2B 1684 1400 693 597 • BCL9CSNK1A1 1694 629 690 1183 • BCL9DIXDC1 1802 1327 1948 2076 • BCL9PORCN 1854 1789 1657 2027 • BCL9RHOU 1892 1855 1852 654 • BCL9WNT3 1954 2115 883 713 • BCL9T 1994 1695 1296 241 • BCL9CTNNB1 2066 176 2375 511 • BCL9FZD5 2331 1094 1013 2077 • BCL9PYGO1 2387 2476 2413 1243

## BTRC 2*^nd^* order interaction ranking at different time points

• APCBTRC 103 2024 259 556 163 • BTRCGSK3A 538 827 1786 1889 2366 • BTRCPYGO1 549 363 1493 10 1777 • BTRCFOXN1 623 2218 1756 1721 1551 • BTRCCCND1 634 397 2366 2273 303 • BTRCFZD2 641 466 2031 1152 1257 • BTRCJUN 694 699 2036 754 1313 • BTRCFGF4 698 593 1944 428 1785 • BTRCCCND2 707 793 2289 106 135 • BTRCPORCN 839 591 997 1917 1251 • BTRCCSNK1A1 846 544 2355 769 2331 • BTRCMYC 901 1125 1250 963 2306 • BTRCWIF1 923 601 2120 417 1444 • BTRCWNT1 930 412 1923 67 2004 • BTRCSFRP1 940 527 2178 627 1234 • BTRCWNT3 951 1282 2153 1924 1934 • BTRCLRP6 1078 910 1773 1573 2148 • BTRCLEF1 1151 371 2416 1070 441 • AXIN1BTRC 1214 2378 748 309 619 • BTRCT 1341 2284 2437 503 2202 • BCL9BTRC 1354 1597 439 1557 727 • BTRCCSNK1G1 1372 1672 1784 1653 1333 • BTRCDVL1 1424 749 2190 99 983 • AESBTRC 1450 74 647 92 2226 • BTRCCTNNBIP1 1474 857 2433 951 1042 • BTRCCTBP2 1584 834 2062 800 1610 • BTRCDVL2 1590 1309 1200 2359 2378 • BTRCFSHB 1621 281 1860 2152 2017 • BTRCCTNNB1 1627 1015 1500 1599 1308 • BTRCSLC9A3R1 1688 166 1081 1358 2413 • BTRCTLE1 1707 587 1836 110 801 • BTRCDAAM1 1726 360 2415 145 960 • BTRCFZD8 1731 2385 2322 844 433 • BTRCFBXW2 1768 48 1686 1888 1559 • BTRCPPP2CA 1800 2191 648 2242 2241 • BTRCCXXC4 1802 1386 1814 566 1858 • BTRCFBXW4 1807 1429 2081 520 2360 • BTRCWNT4 1821 984 2367 195 977 • BTRCPPP2R1A 1831 677 2435 2196 567 • BTRCCTBP1 1833 1142 1498 331 1205 • BTRCNLK 1835 263 1407 1386 1520 • BTRCKREMEN1 1841 183 973 2040 1149 • BTRCFZD6 1880 498 1452 1391 923 • BTRCNKD1 1887 174 1995 795 1323 • BTRCSFRP4 1918 2210 1487 996 928 • BTRCCCND3 1933 760 2147 1071 555 • BTRCEP300 1941 95 2297 248 1938 • BTRCTCF7 1942 1126 1508 553 906 • BTRCFZD1 1978 55 1764 1516 2120 • BTRCPITX2 1981 716 2247 665 1173 • BTRCWNT2 1982 901 2315 394 2112 • BTRCGSK3B 2018 1359 1685 2167 2344 • BTRCFRAT1 2027 1203 2365 1021 2454 • BTRCWNT3A 2034 1180 1755 1997 2401 • BTRCDIXDC1 2054 1153 1304 1572 2154 • BTRCRHOU 2089 1976 1482 664 1657 • BTRCFZD5 2090 1421 1973 1192 768 • BTRCTCF7L1 2134 27 1718 1067 2162 • BTRCCSNK1D 2141 1773 2079 2051 2157 • BTRCWNT2B 2158 940 1401 1076 1565 • BTRCWNT5A 2162 922 2476 1862 2327 • BTRCFBXW11 2176 286 480 1222 769 • BTRCLRP5 2197 613 1577 807 1615 • BTRCTLE2 2214 826 1880 376 1953 • BTRCCSNK2A1 2222 1841 617 2082 1844 • BTRCFOSL1 2270 1163 1248 506 1808 • BTRCFZD7 2286 521 1631 843 1588 • BTRCDKK1 2407 2279 1225 312 2403 • BTRCFRZB 2457 2356 1615 29 2078 • BTRCSENP2 2476 1848 2112 49 1484

## BTRC 2*^nd^* order interaction ranking at different durations

• BTRCSENP2 82 1713 440 236 • BTRCSFRP4 154 2256 1252 129 • BTRCLEF1 313 2025 1991 275 • BTRCLRP6 415 1655 1510 1103 • BTRCCSNK2A1 418 707 1108 2421 • BTRCFGF4 422 2108 828 664 • BTRCSFRP1 446 2140 1327 929 • BTRCFZD2 473 1964 1861 1661 • BTRCCSNK1G1 486 2324 1504 891 • BTRCPITX2 490 1088 1954 1839 • BTRCGSK3B 532 1819 1185 1233 • BTRCFZD7 538 1253 769 115 • BTRCWNT5A 543 1541 1671 1336 • BTRCTCF7L1 566 2170 2239 2208 • BTRCPPP2R1A 580 1968 2265 1926 • BTRCFZD1 591 2102 1342 378 • BTRCFZD6 592 2234 1304 447 • BTRCFBXW2 605 876 2002 1355 • BTRCFBXW11 625 978 82 1632 • BTRCDAAM1 648 1390 1189 42 • BTRCDKK1 697 2215 264 1716 • BTRCTLE2 723 1147 2355 823 • BTRCFZD8 757 1850 1900 2125 • BTRCCXXC4 772 134 413 214 • BTRCPPP2CA 810 1170 2051 1848 • BTRCFRZB 839 441 1977 1908 • BTRCTCF7 841 1396 217 1525 • BTRCNLK 845 1783 378 856 • BTRCDVL2 932 247 1384 1323 • BTRCTLE1 973 2069 2128 471 • BTRCCCND3 987 407 2224 1485 • AESBTRC 1001 1073 2315 1158 • BTRCEP300 1008 50 783 515 • BTRCWNT4 1013 1399 1658 1837 • BTRCNKD1 1016 636 1061 1000 • AXIN1BTRC 1025 1201 1150 1304 • BTRCFBXW4 1029 566 2459 557 • BTRCDVL1 1065 1229 1678 137 • BTRCCTBP2 1124 577 1940 2295 • BTRCLRP5 1178 1552 2179 1437 • BTRCWNT2 1205 2284 1301 2128 • BTRCSLC9A3R1 1299 993 1425 990 • APCBTRC 1304 2091 1557 1842 • BTRCFOSL1 1324 1990 1098 1342 • BTRCGSK3A 1326 2447 1621 487 • BTRCMYC 1361 1101 1025 2183 • BTRCJUN 1370 1022 1165 641 • BTRCWIF1 1417 2121 2431 474 • BTRCKREMEN1 1430 1084 2118 2281 • BCL9BTRC 1431 2238 2276 1484 • BTRCCSNK1D 1446 1622 1821 1763 • BTRCWNT1 1452 592 2337 1797 • BTRCCCND1 1504 1198 1669 777 • BTRCFOXN1 1574 846 1894 834 • BTRCT 1613 1359 2347 532 • BTRCFSHB 1632 849 2204 351 • BTRCFRAT1 1685 1437 2159 1227 • BTRCCTNNBIP1 1708 1020 2323 95 • BTRCWNT2B 1765 2022 1506 229 • BTRCCCND2 1804 643 478 1120 • BTRCCTBP1 1868 826 1046 1696 • BTRCWNT3A 1916 2286 1590 2017 • BTRCDIXDC1 1923 1589 2226 271 • BTRCCTNNB1 1962 239 2055 845 • BTRCFZD5 2223 1551 1508 1745 • BTRCCSNK1A1 2259 973 726 1019 • BTRCWNT3 2335 1698 466 685 • BTRCPYGO1 2437 1081 1690 898 • BTRCPORCN 2445 754 1482 949 • BTRCRHOU 2462 2395 1919 1240

## CXXC4 2*^nd^* order interaction ranking at different time points

• CXXC4GSK3A 74 147 114 415 2259 • CXXC4FOXN1 106 2260 148 349 1045 • CXXC4WNT1 125 2455 113 338 1558 • CXXC4JUN 140 2439 1222 1454 1984 • CXXC4SFRP1 166 187 1411 236 2207 • CXXC4PORCN 194 1364 1185 2303 1035 • CXXC4FGF4 243 2206 2275 339 2415 • CXXC4LEF1 255 1040 1315 568 1379 • CXXC4MYC 261 2077 181 1102 2057 • CXXC4FZD2 275 1573 1211 1103 792 • CXXC4WIF1 335 2147 2285 1933 2048 • CXXC4PYGO1 423 2003 138 505 1851 • CXXC4DVL1 438 2280 1280 1239 1312 • CXXC4WNT3 441 1930 1555 2452 1726 • CXXC4LRP6 477 2408 488 2279 1944 • CXXC4DVL2 487 1004 1342 794 406 • CXXC4FBXW4 559 2056 500 1147 656 • CXXC4T 561 1994 610 708 50 • APCCXXC4 712 275 759 201 425 • CSNK2A1CXXC4 758 60 2296 2457 124 • CXXC4TLE1 769 1601 1273 1848 1735 • CXXC4NKD1 770 2484 590 1198 827 • CXXC4FBXW2 813 2183 535 536 697 • CXXC4KREMEN1 826 838 891 490 1903 • CXXC4DAAM1 836 2105 353 1898 1375 • CXXC4PITX2 892 770 1369 1540 1685 • CXXC4FSHB 914 1402 110 1027 1299 • CSNK1DCXXC4 948 688 1059 430 921 • CXXC4FZD6 987 1430 183 2220 2421 • CXXC4TCF7 1001 1287 336 1751 2443 • FZD5CXXC4 1028 667 1192 124 52 • CXXC4NLK 1035 2141 179 251 1971 • CXXC4SFRP4 1066 825 2382 2362 1745 • CXXC4EP300 1076 637 1255 1006 1329 • CXXC4FZD8 1093 2022 1178 1871 2466 • CXXC4PPP2CA 1098 2404 162 82 2214 • CXXC4SLC9A3R1 1132 2471 1817 1680 2277 • CXXC4PPP2R1A 1152 919 377 1602 646 • CXXC4WNT2 1188 172 740 2353 2314 • CXXC4WNT4 1207 2245 880 2412 1427 • CXXC4WNT5A 1275 1858 295 348 1886 • CXXC4WNT3A 1304 2295 843 2361 1637 • CCND3CXXC4 1327 757 1413 626 2088 • CXXC4DIXDC1 1355 1186 1876 729 1591 • CXXC4WNT2B 1365 1998 150 1706 1503 • CXXC4LRP5 1366 290 1223 1561 1169 • CXXC4GSK3B 1394 2112 1849 1278 2477 • CXXC4FRAT1 1448 869 998 1954 1935 • CXXC4TLE2 1456 987 1124 1836 2204 • CXXC4RHOU 1459 500 1457 454 2435 • CTBP1CXXC4 1477 862 2033 1554 101 • CXXC4FZD1 1528 1444 1568 1325 1219 • CXXC4FBXW11 1572 2137 385 1398 1704 • CXXC4TCF7L1 1611 1104 859 173 90 • CTNNBIP1CXXC4 1652 1070 1865 1841 17 • CXXC4FOSL1 1658 207 2277 1958 2140 • CXXC4FZD7 1753 2465 1467 1493 2147 • CTNNB1CXXC4 1796 119 2186 1636 848 • BTRCCXXC4 1802 1386 1814 566 1858 • CTBP2CXXC4 1815 562 1942 809 625 • CSNK1G1CXXC4 1965 322 2318 1921 569 • CCND2CXXC4 2002 2029 870 486 2085 • CSNK1A1CXXC4 2112 767 1530 1556 103 • CCND1CXXC4 2128 928 2480 784 1367 • AXIN1CXXC4 2142 1066 1671 2325 796 • CXXC4DKK1 2147 1648 46 711 1864 • AESCXXC4 2192 416 1733 1197 1929 • BCL9CXXC4 2208 255 1028 788 729 • CXXC4SENP2 2405 2451 2187 1406 1948 • CXXC4FRZB 2413 1766 2333 1272 1880

## CXXC4 2*^nd^* order interaction ranking at different durations

• FZD5CXXC4 75 46 1869 1502 • CSNK1A1CXXC4 151 449 1661 352 • CTNNB1CXXC4 191 1404 1597 409 • CTBP1CXXC4 227 1252 2359 1404 • AXIN1CXXC4 287 141 606 1717 • CXXC4SENP2 343 2045 1629 1507 • BCL9CXXC4 399 412 675 123 • CCND2CXXC4 429 225 1057 1887 • AESCXXC4 430 461 502 322 • CXXC4SFRP4 468 1377 2195 1270 • CCND1CXXC4 517 57 353 1055 • CSNK1DCXXC4 523 205 816 178 • APCCXXC4 636 2103 2254 1255 • CTNNBIP1CXXC4 653 2248 247 1201 • BTRCCXXC4 772 134 413 214 • CXXC4LRP6 843 1473 1918 1202 • CXXC4LEF1 891 1688 2086 2327 • CXXC4SFRP1 983 1521 1800 1742 • CXXC4FBXW2 990 2017 2165 2222 • CXXC4FZD7 1014 1077 497 1688 • CTBP2CXXC4 1015 91 983 380 • CXXC4PPP2R1A 1023 1884 1841 2050 • CXXC4FZD2 1028 2056 984 1378 • CXXC4FGF4 1088 1180 2082 2351 • CXXC4GSK3B 1096 1542 750 239 • CXXC4FBXW11 1109 2013 2059 2042 • CXXC4FZD1 1135 1620 1747 429 • CXXC4WNT5A 1144 2318 1259 1771 • CXXC4PITX2 1223 1021 908 1211 • CCND3CXXC4 1261 696 363 1059 • CXXC4NLK 1265 2350 581 303 • CXXC4TCF7L1 1283 1436 674 1495 • CXXC4FZD6 1295 2379 1036 1174 • CXXC4FZD8 1298 1572 929 1816 • CXXC4DAAM1 1356 2382 550 2201 • CXXC4FBXW4 1404 1485 999 2199 • CXXC4TLE2 1478 2360 2041 420 • CXXC4PPP2CA 1486 1664 1812 519 • CXXC4DKK1 1493 1573 959 1199 • CXXC4NKD1 1554 1802 1385 1879 • CXXC4LRP5 1562 1012 2193 612 • CXXC4FRZB 1568 1897 2185 576 • CXXC4TCF7 1596 1931 2013 1628 • CXXC4DVL2 1600 1013 2228 1915 • CXXC4EP300 1689 1038 1257 1316 • CXXC4WNT2 1704 1662 1466 2095 • CXXC4TLE1 1715 1684 2326 901 • CXXC4WNT4 1721 1920 1163 1215 • CXXC4FOSL1 1776 2147 2417 1415 • CSNK2A1CXXC4 1787 77 25 555 • CXXC4WNT3A 1862 892 1144 1747 • CXXC4FRAT1 1886 1451 2067 458 • CXXC4WIF1 1938 1207 1651 2058 • CXXC4WNT2B 1953 2398 863 1553 • CXXC4JUN 1990 1940 2039 312 • CXXC4DVL1 2007 2105 1337 1707 • CXXC4WNT1 2030 1697 1755 1614 • CSNK1G1CXXC4 2044 54 928 958 • CXXC4MYC 2055 1617 1568 809 • CXXC4GSK3A 2100 1621 529 979 • CXXC4PYGO1 2136 1776 1723 1077 • CXXC4FSHB 2172 1287 1294 1353 • CXXC4FOXN1 2180 1828 2206 796 • CXXC4SLC9A3R1 2215 1701 1141 2044 • CXXC4DIXDC1 2232 1407 927 1781 • CXXC4T 2307 1603 723 461 • CXXC4RHOU 2368 2239 763 565 • CXXC4KREMEN1 2378 2354 879 1615 • CXXC4WNT3 2400 1300 1015 2060 • CXXC4PORCN 2466 1418 2421 2241

## DAAM1 2*^nd^* order interaction ranking at different time points

• DAAM1GSK3A 132 1331 534 2471 1771 • DAAM1FGF4 171 218 2446 228 958 • DAAM1FOXN1 174 1147 745 993 1241 • DAAM1WNT1 231 1245 965 102 2453 • DAAM1JUN 249 66 1542 2294 1526 • DAAM1WIF1 265 1637 1578 1840 2077 • DAAM1PYGO1 334 733 953 296 1819 • DAAM1SFRP1 354 285 1441 2433 1728 • APCDAAM1 372 2361 352 448 93 • DAAM1PORCN 375 494 2402 679 1671 • DAAM1LEF1 435 1012 832 820 1686 • DAAM1MYC 472 1530 1324 1313 732 • DAAM1FZD2 512 1433 2391 720 952 • DAAM1WNT3 516 1500 1191 1583 1611 • DAAM1T 517 512 409 1858 995 • FZD5DAAM1 534 692 262 1426 94 • DAAM1LRP6 569 451 1078 896 1713 • DAAM1DVL1 606 1479 1456 1949 1318 • DAAM1DVL2 638 1112 1597 275 2304 • CSNK2A1DAAM1 711 155 2444 1863 563 • CXXC4DAAM1 836 2105 353 1898 1375 • CTBP1DAAM1 851 1204 393 972 64 • CSNK1DDAAM1 865 1565 902 1575 167 • DAAM1NKD1 889 1845 916 202 2339 • CCND3DAAM1 921 351 545 757 2311 • DAAM1FSHB 989 1091 286 459 2189 • DAAM1FBXW4 1020 1522 272 692 1792 • DAAM1TLE1 1034 515 2075 728 1690 • DAAM1EP300 1162 69 712 2339 733 • DAAM1FBXW2 1184 476 1344 1054 1680 • DAAM1SLC9A3R1 1198 1937 2273 1442 1081 • DAAM1KREMEN1 1253 299 1433 1525 1748 • DAAM1PITX2 1261 1156 1132 21 1572 • DAAM1SFRP4 1266 2240 1111 1404 1022 • DAAM1WNT2 1273 1740 2231 2100 1122 • DAAM1FZD8 1348 1424 2067 803 1708 • DAAM1FZD6 1378 1011 424 539 2390 • DAAM1WNT3A 1393 503 1430 1929 1471 • DAAM1PPP2CA 1434 1351 829 37 1457 • DAAM1TCF7 1439 608 497 2187 1366 • DAAM1DIXDC1 1446 67 1892 1350 2141 • DAAM1RHOU 1455 971 1632 98 2281 • DAAM1NLK 1457 35 720 2005 1229 • DAAM1WNT2B 1465 375 555 1597 2074 • DAAM1WNT4 1472 751 1839 1201 1391 • DAAM1TLE2 1530 950 2164 1287 2196 • CTNNB1DAAM1 1540 1979 812 1641 512 • DAAM1LRP5 1547 403 2139 1837 1778 • CTBP2DAAM1 1556 456 136 28 530 • DAAM1GSK3B 1575 846 1151 332 2252 • DAAM1PPP2R1A 1592 516 625 1867 701 • CTNNBIP1DAAM1 1619 2444 774 444 3 • CSNK1G1DAAM1 1642 2062 2457 540 757 • DAAM1WNT5A 1653 430 135 1063 1687 • DAAM1FRAT1 1674 2290 1787 1798 1804 • DAAM1FBXW11 1718 1019 1182 1581 1528 • BTRCDAAM1 1726 360 2415 145 960 • DAAM1FOSL1 1736 518 2152 2096 2274 • AESDAAM1 1769 145 928 647 1001 • DAAM1TCF7L1 1822 1460 2393 2097 418 • DAAM1FZD1 1882 1693 1677 2032 1570 • DAAM1FZD7 1898 1748 2199 1494 2064 • CCND2DAAM1 1928 1211 1960 966 1811 • AXIN1DAAM1 1952 1095 1829 808 491 • CSNK1A1DAAM1 2053 1666 1511 2007 73 • BCL9DAAM1 2230 1810 1520 2209 1336 • CCND1DAAM1 2237 77 653 818 1850 • DAAM1DKK1 2330 995 826 1570 1303 • DAAM1SENP2 2353 1249 1314 1723 982 • DAAM1FRZB 2398 1281 1943 1450 1865

## DAAM1 2*^nd^* order interaction ranking at different durations

• CSNK1A1DAAM1 57 1676 1086 94 • FZD5DAAM1 162 623 1933 2030 • CTNNB1DAAM1 271 2407 1728 730 • CCND1DAAM1 373 698 1895 1691 • CTBP1DAAM1 381 2341 1018 396 • CTNNBIP1DAAM1 408 1800 2361 1526 • DAAM1SENP2 411 1227 52 1922 • CCND2DAAM1 443 2428 938 1532 • BCL9DAAM1 466 1845 1655 2414 • AESDAAM1 485 2042 1523 10 • CSNK1DDAAM1 504 608 2116 18 • AXIN1DAAM1 508 870 2469 2399 • BTRCDAAM1 648 1390 1189 42 • CTBP2DAAM1 670 1036 1567 1047 • DAAM1SFRP4 680 2107 427 1760 • APCDAAM1 776 1826 2022 1464 • DAAM1LRP6 829 2252 508 1853 • DAAM1LEF1 895 1860 1142 2041 • DAAM1FZD7 929 2356 1357 2280 • DAAM1FZD2 960 1263 1986 731 • DAAM1SFRP1 1021 2277 923 2371 • DAAM1FZD1 1035 1360 2312 537 • DAAM1PPP2R1A 1039 850 981 1633 • DAAM1TCF7L1 1095 1524 405 1831 • DAAM1FGF4 1132 1522 1493 2016 • DAAM1PITX2 1149 944 1348 220 • DAAM1FBXW11 1173 1903 172 590 • DAAM1GSK3B 1180 1564 1293 158 • DAAM1FZD6 1209 908 1386 1104 • DAAM1WNT5A 1240 1005 1023 1698 • DAAM1FZD8 1302 1342 1491 1239 • DAAM1FBXW2 1313 1369 541 1329 • CXXC4DAAM1 1356 2382 550 2201 • CCND3DAAM1 1357 1459 847 1699 • DAAM1DKK1 1362 2459 215 377 • DAAM1NLK 1376 1093 122 1595 • DAAM1TLE1 1498 2146 2403 1151 • DAAM1TCF7 1500 1918 112 1221 • DAAM1PPP2CA 1503 2133 479 874 • DAAM1FRZB 1532 798 492 544 • DAAM1EP300 1557 154 585 1654 • DAAM1LRP5 1652 1951 582 342 • DAAM1TLE2 1686 2427 1044 405 • DAAM1WNT4 1728 2004 1543 1390 • DAAM1FBXW4 1736 340 660 1952 • DAAM1DVL2 1737 1219 1735 1973 • CSNK2A1DAAM1 1814 1765 592 1345 • DAAM1GSK3A 1834 1477 2225 2425 • DAAM1FOSL1 1898 1902 171 2210 • DAAM1NKD1 1957 406 212 1115 • DAAM1MYC 1988 977 152 1991 • DAAM1T 2029 799 659 71 • DAAM1FRAT1 2043 913 291 970 • DAAM1SLC9A3R1 2057 2441 2163 1461 • DAAM1WNT1 2061 1112 832 1039 • DAAM1DVL1 2063 922 2154 858 • DAAM1FSHB 2141 2097 811 1051 • CSNK1G1DAAM1 2152 254 1033 781 • DAAM1JUN 2198 1167 526 232 • DAAM1WNT2 2243 1601 2266 414 • DAAM1FOXN1 2253 1385 239 855 • DAAM1PORCN 2254 1222 678 1813 • DAAM1DIXDC1 2291 1858 1595 286 • DAAM1WNT3A 2302 1962 2405 1195 • DAAM1KREMEN1 2310 1526 2053 1172 • DAAM1WIF1 2317 2119 607 2458 • DAAM1WNT2B 2339 2283 265 2053 • DAAM1RHOU 2436 1425 2098 143 • DAAM1WNT3 2444 1494 120 2032 • DAAM1PYGO1 2461 883 446 854

## DIXDC1 2*^nd^* order interaction ranking at different time points

• DIXDC1SFRP1 100 1886 468 2322 622 • DIXDC1LEF1 122 251 145 1891 2482 • DIXDC1FOXN1 138 2044 10 687 1977 • DIXDC1JUN 145 878 200 2235 1485 • DIXDC1GSK3A 151 260 4 2044 2448 • DIXDC1FGF4 176 108 757 2292 449 • DIXDC1PYGO1 191 1384 61 367 1511 • DIXDC1FZD2 196 338 347 396 674 • DIXDC1PORCN 201 2201 87 1585 912 • DIXDC1WNT1 202 2172 12 516 2000 • DIXDC1MYC 272 2276 21 2461 1712 • DIXDC1WIF1 290 2332 572 1567 2040 • DIXDC1DVL1 328 2055 176 1035 671 • DIXDC1WNT3 331 847 332 755 1294 • DIXDC1DVL2 421 491 72 1144 1286 • DIXDC1T 459 1520 73 1648 1070 • DIXDC1LRP6 513 2038 15 2214 724 • DIXDC1NKD1 522 1380 20 478 2284 • DIXDC1TLE1 590 1921 256 2478 672 • DIXDC1FBXW4 603 1793 51 2313 1843 • DIXDC1SFRP4 636 2189 1156 2103 2240 • DIXDC1FSHB 646 1959 1 653 2155 • DIXDC1FZD6 755 1829 3 2080 944 • DIXDC1EP300 785 1047 265 1897 1643 • APCDIXDC1 786 1409 2136 1919 684 • DIXDC1KREMEN1 823 82 56 1114 2335 • DIXDC1FBXW2 829 1425 39 1735 683 • DIXDC1PITX2 845 179 196 1235 2129 • DIXDC1PPP2CA 869 2380 6 2302 1899 • DIXDC1WNT4 908 1106 88 2394 2170 • DIXDC1FZD8 934 1325 401 1091 1541 • DIXDC1SLC9A3R1 973 1736 431 373 1912 • DIXDC1WNT5A 996 1710 53 2116 2389 • DIXDC1WNT3A 1030 1286 170 1420 2065 • DIXDC1WNT2B 1036 1494 9 1095 1365 • DIXDC1PPP2R1A 1044 434 70 1228 1344 • DIXDC1NLK 1051 2197 2 838 371 • DIXDC1WNT2 1079 43 199 938 1421 • CSNK1DDIXDC1 1114 986 1774 866 1627 • CSNK2A1DIXDC1 1126 413 1316 1947 58 • DIXDC1GSK3B 1127 1352 252 11 2478 • DIXDC1FRAT1 1154 1708 193 2387 2171 • DIXDC1TCF7 1213 1189 13 1532 1304 • DIXDC1FOSL1 1242 34 299 1927 1573 • CTBP1DIXDC1 1301 1866 1166 2467 191 • DIXDC1RHOU 1326 1919 221 1212 1989 • DIXDC1LRP5 1336 1029 25 610 337 • FZD5DIXDC1 1337 813 2006 835 484 • DIXDC1TLE2 1345 45 236 1289 1254 • CXXC4DIXDC1 1355 1186 1876 729 1591 • DIXDC1FZD7 1384 1801 540 2370 1842 • CCND3DIXDC1 1392 1878 1604 1621 1810 • DIXDC1FBXW11 1400 871 7 1337 1602 • DIXDC1TCF7L1 1402 1671 349 283 1296 • DAAM1DIXDC1 1446 67 1892 1350 2141 • DIXDC1FZD1 1559 993 290 841 2434 • CTNNB1DIXDC1 1705 1345 1684 1744 774 • CTNNBIP1DIXDC1 1853 2428 1752 393 141 • CSNK1G1DIXDC1 1925 811 1672 1424 2225 • AESDIXDC1 1985 335 1831 469 2059 • BTRCDIXDC1 2054 1153 1304 1572 2154 • CTBP2DIXDC1 2061 1052 1927 1085 338 • CCND2DIXDC1 2076 354 2026 802 969 • DIXDC1DKK1 2105 690 11 1488 1699 • AXIN1DIXDC1 2120 1676 1691 2249 228 • CSNK1A1DIXDC1 2250 1734 1767 426 168 • BCL9DIXDC1 2256 578 1828 1307 2035 • DIXDC1SENP2 2366 2012 651 1409 2361 • DIXDC1FRZB 2388 316 742 2440 1396 • CCND1DIXDC1 2408 1658 1605 386 1536

## DIXDC1 2*^nd^* order interaction ranking at different durations

• DIXDC1SENP2 25 475 76 1700 • DIXDC1SFRP4 60 677 196 1682 • DIXDC1LRP6 85 553 531 1924 • DIXDC1FGF4 111 802 1361 2083 • DIXDC1FZD2 124 248 1200 1194 • DIXDC1LEF1 137 627 757 1349 • DIXDC1FBXW2 140 18 205 1422 • DIXDC1PPP2R1A 146 26 374 1684 • DIXDC1FZD1 156 788 1067 875 • DIXDC1FZD7 160 808 1898 2142 • DIXDC1WNT5A 164 30 442 1764 • DIXDC1NLK 170 15 151 1744 • DIXDC1DKK1 202 1889 168 1301 • DIXDC1GSK3B 213 953 1928 113 • FZD5DIXDC1 229 1223 1143 1387 • DIXDC1FBXW11 233 38 107 1604 • DIXDC1TCF7L1 250 906 242 1241 • CSNK1A1DIXDC1 256 1532 756 243 • DIXDC1PITX2 264 1619 1899 662 • DIXDC1PPP2CA 268 65 195 1061 • DIXDC1SFRP1 277 785 947 2427 • DIXDC1FZD8 289 1371 1680 1150 • DIXDC1FZD6 307 306 1736 789 • DIXDC1TLE2 311 74 778 163 • DIXDC1TLE1 326 294 1278 2476 • DIXDC1DVL2 332 22 2072 1010 • DIXDC1FRZB 383 4 129 249 • DIXDC1TCF7 431 47 26 848 • DIXDC1EP300 447 2 801 1346 • DIXDC1LRP5 481 1410 666 48 • DIXDC1WNT2 522 244 1647 350 • DIXDC1WNT4 572 24 695 1067 • DIXDC1FBXW4 579 12 391 1036 • DIXDC1SLC9A3R1 587 221 551 827 • DIXDC1NKD1 681 11 47 1547 • DIXDC1WNT1 715 9 630 762 • DIXDC1FRAT1 788 96 469 338 • DIXDC1WIF1 832 595 629 1959 • DIXDC1DVL1 861 13 1837 1321 • DIXDC1FOXN1 958 17 567 289 • DIXDC1FOSL1 967 548 323 1386 • DIXDC1MYC 994 27 180 709 • DIXDC1KREMEN1 1002 51 1743 523 • DIXDC1GSK3A 1012 440 2018 1279 • DIXDC1T 1018 61 515 451 • DIXDC1JUN 1027 126 388 839 • DIXDC1FSHB 1031 414 578 1411 • CCND2DIXDC1 1348 960 1081 1257 • DIXDC1WNT2B 1477 430 227 1581 • DIXDC1WNT3 1537 529 206 1341 • CCND1DIXDC1 1541 1067 2190 334 • DIXDC1PYGO1 1546 165 1164 388 • DIXDC1WNT3A 1572 350 1688 371 • AXIN1DIXDC1 1575 2307 1872 2225 • CTBP1DIXDC1 1577 1350 1127 1313 • CTNNB1DIXDC1 1615 860 2143 216 • CSNK1DDIXDC1 1742 2290 2300 389 • BCL9DIXDC1 1802 1327 1948 2076 • DIXDC1RHOU 1823 300 2424 439 • CTNNBIP1DIXDC1 1876 1901 1449 1888 • DIXDC1PORCN 1879 118 351 1937 • BTRCDIXDC1 1923 1589 2226 271 • CTBP2DIXDC1 1978 1166 1221 1254 • AESDIXDC1 2097 1790 2344 215 • APCDIXDC1 2140 1406 729 1381 • CSNK1G1DIXDC1 2160 2361 1945 776 • CXXC4DIXDC1 2232 1407 927 1781 • CCND3DIXDC1 2267 1518 1253 1006 • DAAM1DIXDC1 2291 1858 1595 286 • CSNK2A1DIXDC1 2440 1388 1376 554

## DKK1 2*^nd^* order interaction ranking at different time points

• DKK1GSK3A 6 2313 689 1907 1362 • DKK1WNT1 15 2415 1260 2300 1302 • DKK1FGF4 16 2089 1345 1490 1928 • DKK1JUN 17 2148 1629 864 1279 • DKK1WIF1 21 2425 2359 119 2159 • DKK1PYGO1 22 2018 1054 280 876 • DKK1FOXN1 24 2405 1251 1434 404 • DKK1T 27 2275 2174 727 1376 • DKK1WNT3 31 2040 2041 906 1998 • DKK1LEF1 33 2318 2474 876 2307 • DKK1SFRP1 35 2383 2293 1403 427 • DKK1DVL1 36 2426 2175 75 1642 • DKK1MYC 37 2349 1497 196 2337 • DKK1LRP6 50 2347 2208 669 2353 • DKK1PORCN 63 2367 2012 1196 1898 • DKK1NKD1 79 2381 1377 1159 2308 • DKK1FZD6 80 2163 2122 690 2355 • DKK1DVL2 96 2214 2220 1654 1932 • DKK1FZD2 97 2073 2046 30 2377 • DKK1NLK 118 2401 1436 1231 1918 • DKK1FBXW4 137 2358 1667 1492 2268 • DKK1FSHB 146 1423 1909 385 2047 • DKK1EP300 162 2328 2241 723 1711 • DKK1KREMEN1 180 2154 1815 31 2398 • DKK1SLC9A3R1 186 2173 2116 750 2420 • DKK1TLE1 188 2390 1622 198 975 • DKK1PPP2R1A 204 2066 2151 2449 1450 • DKK1PITX2 206 2357 1818 1099 2149 • DKK1SFRP4 254 111 1607 121 737 • DKK1FZD8 262 2433 1479 2290 2224 • DKK1GSK3B 279 2389 1937 2198 2201 • DKK1PPP2CA 296 2422 1285 2212 1236 • DKK1WNT4 312 1968 1719 185 208 • DKK1WNT3A 317 1408 2321 2430 2283 • DKK1LRP5 345 1339 1690 2288 782 • DKK1FBXW2 365 2065 1349 1007 1968 • DKK1WNT2 370 1201 2098 702 1287 • DKK1WNT5A 382 2269 1464 2219 1479 • DKK1FBXW11 391 2423 1179 1124 2382 • DKK1FOSL1 404 2205 1798 699 2280 • DKK1WNT2B 406 1982 1099 1683 2383 • DKK1TCF7 407 2338 1936 1761 1438 • DKK1FZD1 427 2034 1539 1005 1775 • DKK1FRAT1 439 2042 1573 578 1096 • DKK1FZD7 455 2188 2058 1511 1816 • DKK1TLE2 461 2324 2057 361 2368 • DKK1RHOU 491 2399 1489 1385 1950 • DKK1TCF7L1 550 1958 2221 740 1641 • DKK1FRZB 1727 1704 2225 1148 2022 • DKK1SENP2 1758 2446 1996 577 2451 • CSNK1DDKK1 1843 1698 884 770 1346 • APCDKK1 1940 2098 323 1332 1954 • FZD5DKK1 1943 1328 258 2084 1010 • CSNK2A1DKK1 2056 1813 1147 1268 460 • DIXDC1DKK1 2105 690 11 1488 1699 • CXXC4DKK1 2147 1648 46 711 1864 • CCND3DKK1 2253 1058 704 2078 2295 • DAAM1DKK1 2330 995 826 1570 1303 • CTBP1DKK1 2335 1590 96 1298 207 • CTBP2DKK1 2351 1993 310 1261 966 • CTNNBIP1DKK1 2389 1910 84 1601 44 • CTNNB1DKK1 2390 1931 276 1671 839 • BTRCDKK1 2407 2279 1225 312 2403 • CSNK1G1DKK1 2415 2322 847 66 2005 • CCND2DKK1 2417 1387 141 657 1043 • CSNK1A1DKK1 2427 2004 318 2179 349 • AXIN1DKK1 2432 2362 494 916 1397 • AESDKK1 2445 1411 642 477 2406 • BCL9DKK1 2452 543 666 1123 1799 • CCND1DKK1 2459 999 562 2437 1091

## DKK1 2*^nd^* order interaction ranking at different durations

• FZD5DKK1 16 2181 2313 2462 • CSNK1A1DKK1 179 1283 1312 8 • CTNNB1DKK1 184 996 507 2200 • DIXDC1DKK1 202 1889 168 1301 • DKK1SENP2 267 886 1031 959 • CTBP1DKK1 358 1389 1779 2471 • DKK1SFRP4 528 1109 1951 207 • CCND2DKK1 573 1366 341 639 • CSNK1DDKK1 651 2372 211 1806 • AESDKK1 676 2263 948 1097 • APCDKK1 687 778 2468 1392 • BTRCDKK1 697 2215 264 1716 • BCL9DKK1 726 2261 1442 2163 • CTNNBIP1DKK1 728 1774 138 1121 • DKK1FGF4 748 905 1367 1328 • CTBP2DKK1 766 2099 2191 2007 • DKK1FZD7 768 1260 400 1489 • DKK1LRP6 785 859 799 1334 • CCND1DKK1 798 2216 158 2134 • AXIN1DKK1 801 1988 633 1565 • DKK1FZD2 847 422 1093 2231 • DKK1LEF1 866 2264 2151 1776 • DKK1FZD1 878 1602 2200 1413 • DKK1PPP2R1A 954 260 1796 1911 • DKK1FBXW2 966 256 939 1318 • DKK1GSK3B 971 1405 685 411 • DKK1SFRP1 996 1367 1679 617 • DKK1TCF7L1 1005 1640 2437 2338 • DKK1FZD8 1055 487 781 2256 • DKK1FBXW11 1070 155 1941 2052 • DKK1TCF7 1119 246 2106 1166 • DKK1WNT5A 1129 264 2270 1893 • DKK1PITX2 1177 903 1283 2434 • DKK1NLK 1250 172 126 1738 • DKK1FRZB 1273 48 850 1599 • CCND3DKK1 1276 1720 1012 2345 • DKK1FZD6 1280 466 737 494 • DKK1PPP2CA 1309 173 1479 2442 • DAAM1DKK1 1362 2459 215 377 • CXXC4DKK1 1493 1573 959 1199 • DKK1TLE1 1540 416 1298 605 • DKK1FBXW4 1559 147 1776 655 • DKK1GSK3A 1581 888 987 231 • DKK1DVL2 1611 64 1815 392 • DKK1EP300 1626 3 1617 488 • DKK1TLE2 1637 257 1391 417 • DKK1LRP5 1727 1155 2339 1470 • CSNK2A1DKK1 1733 2269 106 2234 • DKK1WNT4 1735 238 1492 2470 • DKK1FSHB 1819 909 1229 538 • DKK1FOSL1 1826 1535 2328 2239 • DKK1NKD1 1857 49 2248 1655 • DKK1KREMEN1 1858 161 1455 1027 • DKK1WIF1 1865 969 2292 1206 • CSNK1G1DKK1 1866 1746 1205 962 • DKK1WNT1 1918 201 2303 1299 • DKK1MYC 1940 263 835 2224 • DKK1WNT2 1947 395 510 2366 • DKK1SLC9A3R1 1960 405 780 309 • DKK1WNT2B 2008 649 2135 264 • DKK1DVL1 2013 150 1436 150 • DKK1JUN 2032 270 1119 1858 • DKK1FOXN1 2089 302 2342 2464 • DKK1FRAT1 2116 279 2346 2386 • DKK1PORCN 2125 220 2445 486 • DKK1RHOU 2239 848 1160 1332 • DKK1WNT3 2312 2229 2060 1265 • DKK1WNT3A 2343 943 903 2477 • DKK1T 2344 289 2287 810 • DKK1PYGO1 2374 346 2144 2235

## EP300 2*^nd^* order interaction ranking at different time points

• DKK1EP300 162 2328 2241 723 1711 • EP300GSK3A 218 1350 521 1506 794 • EP300SFRP1 266 1639 2453 2169 254 • EP300MYC 294 592 668 734 2126 • EP300WNT1 297 717 204 488 536 • EP300FOXN1 300 898 882 997 1915 • EP300FGF4 315 1097 2170 1791 1191 • EP300PYGO1 327 1064 1005 676 1555 • EP300LEF1 333 1662 1525 796 723 • EP300FZD2 374 1467 1879 574 2363 • EP300LRP6 389 1329 1023 1963 1080 • EP300WNT3 419 3 2055 1083 348 • EP300PORCN 424 1625 1292 1387 424 • EP300JUN 428 65 1419 1281 2150 • EP300WIF1 482 1192 1357 1229 1437 • EP300T 484 440 1034 1699 1603 • CSNK1DEP300 617 1008 1917 760 108 • APCEP300 642 952 1360 1189 1384 • CSNK2A1EP300 675 2133 1563 1980 2151 • DIXDC1EP300 785 1047 265 1897 1643 • FZD5EP300 824 870 1855 1878 830 • EP300TLE1 877 1379 1350 2295 1008 • EP300SFRP4 898 2274 2449 2439 1821 • EP300NKD1 912 1856 529 903 865 • EP300FSHB 984 361 451 176 2248 • EP300FBXW2 994 1076 589 1127 580 • EP300KREMEN1 1006 1853 1521 281 639 • EP300FBXW4 1008 1461 781 2265 1673 • CXXC4EP300 1076 637 1255 1006 1329 • EP300GSK3B 1107 276 1935 622 1949 • EP300PPP2CA 1120 230 287 1381 1923 • EP300SLC9A3R1 1129 244 1749 1322 1683 • EP300NLK 1148 1684 432 982 507 • DAAM1EP300 1162 69 712 2339 733 • EP300PITX2 1172 339 1091 315 1736 • EP300WNT4 1185 1341 1068 2427 1537 • EP300FZD8 1211 72 2336 1463 200 • EP300PPP2R1A 1217 1206 660 1626 386 • EP300FZD7 1232 897 2213 1098 1420 • EP300WNT3A 1236 669 1550 948 2080 • EP300FZD6 1245 734 362 561 1230 • EP300FRAT1 1292 1560 2240 1649 2440 • CTBP1EP300 1323 1830 1669 2089 2228 • EP300WNT2B 1352 10 688 475 2340 • EP300TCF7 1357 1256 437 832 252 • EP300TLE2 1361 1703 2379 571 1893 • EP300WNT2 1362 2143 1950 1715 1723 • CCND3EP300 1381 673 1600 256 2235 • EP300FOSL1 1476 795 2103 2403 2459 • EP300LRP5 1497 2122 898 813 1815 • EP300FBXW11 1505 37 741 1666 1426 • CSNK1G1EP300 1521 1360 1339 1785 2386 • EP300RHOU 1524 598 1852 65 831 • AXIN1EP300 1536 937 1628 2063 1171 • CTNNB1EP300 1541 785 1982 1620 1926 • EP300WNT5A 1562 1523 577 1584 657 • EP300TCF7L1 1576 1098 2314 1772 1801 • EP300FZD1 1597 88 1953 512 2076 • CTNNBIP1EP300 1733 1669 1485 1433 385 • CTBP2EP300 1754 916 1019 973 1203 • DVL2EP300 1876 804 1361 225 1262 • AESEP300 1886 1033 1932 369 2086 • BTRCEP300 1941 95 2297 248 1938 • CSNK1A1EP300 2020 2231 1906 2224 1231 • DVL1EP300 2031 68 786 210 1568 • BCL9EP300 2036 1788 1416 1221 2045 • CCND2EP300 2046 763 1895 2155 2243 • CCND1EP300 2148 801 1267 464 1127 • EP300FRZB 2308 137 1891 967 1472 • EP300SENP2 2394 680 1959 1181 1134

## EP300 2*^nd^* order interaction ranking at different durations

• EP300SENP2 163 1732 548 1360 • CSNK1A1EP300 249 511 2314 51 • FZD5EP300 285 178 2138 2349 • EP300SFRP4 340 1667 396 1466 • CTBP1EP300 442 776 1756 1324 • DIXDC1EP300 447 2 801 1346 • CTNNB1EP300 524 970 1979 520 • EP300LRP6 600 1582 461 2318 • EP300SFRP1 669 1217 817 2045 • EP300LEF1 686 1692 2436 1912 • EP300FZD7 712 1722 1222 1112 • AESEP300 727 695 1871 285 • CTNNBIP1EP300 734 2482 1341 1772 • EP300WNT5A 740 2090 964 2422 • EP300FBXW11 750 872 425 651 • CCND2EP300 767 362 1942 1063 • EP300FZD2 771 1914 1968 1715 • EP300GSK3B 781 1586 1952 393 • EP300FGF4 787 1249 2122 1442 • EP300FBXW2 831 2218 1637 1576 • CCND1EP300 834 32 1389 2387 • EP300TCF7L1 864 1193 1005 1984 • EP300FZD1 870 1788 2373 1380 • BCL9EP300 872 495 2201 1088 • EP300PITX2 894 1100 1659 217 • AXIN1EP300 900 273 1907 1459 • EP300FZD8 928 1330 2211 1463 • EP300PPP2R1A 953 2185 1215 2359 • EP300TCF7 985 1105 28 1729 • BTRCEP300 1008 50 783 515 • EP300NLK 1010 2435 35 2246 • CSNK1DEP300 1042 112 1944 27 • EP300FZD6 1086 2130 2137 502 • APCEP300 1094 1040 1047 1994 • EP300PPP2CA 1211 2114 1082 1383 • EP300FBXW4 1217 1781 1419 1291 • EP300FRZB 1218 1628 300 270 • EP300LRP5 1277 1296 2432 242 • DVL1EP300 1293 1186 1745 294 • EP300TLE1 1351 842 2476 1650 • EP300TLE2 1355 1332 1807 66 • CTBP2EP300 1400 53 2462 1268 • EP300WNT4 1410 2335 1713 1722 • DVL2EP300 1449 320 1194 2021 • EP300FOSL1 1462 1236 1601 1446 • EP300NKD1 1505 2067 1111 1652 • CCND3EP300 1534 399 2425 1246 • DAAM1EP300 1557 154 585 1654 • EP300WIF1 1609 1133 2107 1247 • DKK1EP300 1626 3 1617 488 • EP300FOXN1 1653 542 1247 1110 • EP300SLC9A3R1 1664 1472 2365 2398 • EP300WNT2 1679 1721 1859 1456 • CXXC4EP300 1689 1038 1257 1316 • EP300FSHB 1793 2258 1976 1022 • EP300WNT1 1837 1126 2394 1560 • EP300GSK3A 1845 990 1373 495 • EP300FRAT1 1910 1750 2166 689 • EP300T 1931 1395 1274 705 • EP300MYC 1944 1908 311 434 • EP300KREMEN1 1946 816 1527 788 • EP300JUN 2077 1959 1715 186 • EP300WNT2B 2078 1506 782 1185 • EP300WNT3A 2086 1431 2317 1869 • EP300RHOU 2113 1136 1989 190 • CSNK2A1EP300 2153 129 339 814 • CSNK1G1EP300 2159 6 2447 1971 • EP300PORCN 2171 1935 2183 2344 • EP300PYGO1 2235 1919 456 1430 • EP300WNT3 2401 1642 514 1372

## FGF4 2*^nd^* order interaction ranking at different time points

• DKK1FGF4 16 2089 1345 1490 1928 • APCFGF4 41 1592 1827 1895 255 • FBXW11FGF4 46 824 1340 1194 1737 • CSNK2A1FGF4 102 805 2073 2111 89 • DAAM1FGF4 171 218 2446 228 958 • DIXDC1FGF4 176 108 757 2292 449 • CXXC4FGF4 243 2206 2275 339 2415 • CSNK1DFGF4 281 949 1626 1169 2310 • EP300FGF4 315 1097 2170 1791 1191 • CCND3FGF4 342 966 2040 1448 1101 • CTNNB1FGF4 352 1427 1780 1693 56 • FZD5FGF4 362 93 2072 1655 359 • CTBP1FGF4 392 931 1118 1816 248 • CSNK1G1FGF4 468 707 1862 596 620 • CTNNBIP1FGF4 499 176 1680 409 41 • DVL2FGF4 536 797 2255 1082 642 • FBXW2FGF4 548 2118 1771 2071 1465 • FGF4GSK3A 576 1670 28 1651 1038 • DVL1FGF4 633 1184 2150 1149 1117 • CTBP2FGF4 661 1497 2473 1109 502 • BTRCFGF4 698 593 1944 428 1785 • FGF4FZD2 708 302 156 1 286 • AESFGF4 740 1165 1770 1003 514 • AXIN1FGF4 753 2106 2425 1775 436 • FGF4PYGO1 779 1407 78 1643 137 • FGF4MYC 863 1271 245 241 1034 • FGF4WNT1 874 1889 75 136 1793 • FGF4SFRP1 1061 1272 721 629 909 • FGF4WIF1 1080 1957 1480 1877 2082 • FGF4FOXN1 1111 2013 18 1768 1129 • CSNK1A1FGF4 1123 386 1581 2142 32 • BCL9FGF4 1156 1123 1483 1610 578 • CCND2FGF4 1192 1506 2350 2156 1092 • FGF4JUN 1203 1288 776 1591 1090 • FGF4WNT3 1270 1484 1404 2463 415 • FGF4T 1385 1782 93 2411 1202 • FGF4LEF1 1397 906 411 1250 2264 • CCND1FGF4 1425 432 1590 2042 2131 • FGF4PORCN 1486 1209 392 1866 399 • FGF4LRP6 1606 2200 172 1750 1412 • FGF4TLE1 1719 2134 1271 2109 1668 • FGF4FSHB 1892 26 76 239 1703 • FGF4FZD6 1931 1826 37 1876 347 • FGF4WNT2 1947 83 1647 1474 1069 • FGF4TCF7 1979 777 106 1923 821 • FGF4FBXW4 1994 1499 132 1094 2001 • FGF4NKD1 1996 2320 104 703 2321 • FGF4SLC9A3R1 2030 1557 874 1774 311 • FGF4PPP2CA 2077 2121 8 1529 1855 • FGF4WNT5A 2108 738 143 1139 2033 • FGF4WNT2B 2173 342 14 526 2404 • FGF4WNT3A 2186 442 818 1769 798 • FGF4NLK 2188 823 19 2350 175 • FGF4SFRP4 2191 665 1105 1582 1074 • FGF4LRP5 2201 617 216 1981 1182 • FGF4FZD8 2217 1519 729 837 2144 • FGF4GSK3B 2233 1622 1058 1255 2247 • FGF4KREMEN1 2241 1240 563 1828 1538 • FGF4WNT4 2242 2146 406 2399 1145 • FGF4PITX2 2246 788 364 599 1761 • FGF4RHOU 2268 1679 1296 353 1532 • FGF4FOSL1 2279 284 1635 2232 1702 • FGF4PPP2R1A 2287 1618 247 2241 262 • FGF4TCF7L1 2293 1483 733 1849 1055 • FGF4FRAT1 2321 2058 317 1776 2408 • FGF4FZD1 2332 1692 474 1994 1820 • FGF4TLE2 2356 1543 395 104 1872 • FGF4FZD7 2404 1413 1596 678 1137 • FGF4FRZB 2460 876 1584 485 33 • FGF4SENP2 2470 1975 1237 1718 1939

## FGF4 2*^nd^* order interaction ranking at different durations

• FZD5FGF4 15 2225 1399 2305 • CSNK1A1FGF4 59 1625 1965 234 • CCND2FGF4 109 1594 1473 2139 • DIXDC1FGF4 111 802 1361 2083 • CTNNB1FGF4 190 1341 1600 357 • BCL9FGF4 198 1958 1347 1731 • CTBP1FGF4 220 1510 773 985 • CCND1FGF4 257 1852 1982 1920 • AESFGF4 260 1245 1191 183 • CSNK1DFGF4 283 1220 1489 211 • AXIN1FGF4 300 2376 749 1438 • APCFGF4 363 1156 990 2062 • CTNNBIP1FGF4 378 1733 1547 994 • BTRCFGF4 422 2108 828 664 • CTBP2FGF4 547 2330 2188 1756 • DVL1FGF4 557 688 1289 635 • DVL2FGF4 647 1438 2197 1975 • FGF4SENP2 745 419 441 1517 • DKK1FGF4 748 905 1367 1328 • EP300FGF4 787 1249 2122 1442 • CCND3FGF4 946 1382 2231 1289 • FGF4SFRP4 950 935 755 1309 • FGF4LEF1 1050 1034 2180 1533 • CXXC4FGF4 1088 1180 2082 2351 • DAAM1FGF4 1132 1522 1493 2016 • FBXW2FGF4 1151 2075 1051 1266 • FBXW11FGF4 1198 2217 1578 751 • FGF4GSK3B 1219 1066 2096 383 • FGF4LRP6 1256 708 523 2215 • FGF4PPP2R1A 1323 211 1708 879 • FGF4FZD2 1342 390 1808 2223 • FGF4TCF7L1 1389 1679 418 1163 • CSNK2A1FGF4 1399 2294 746 881 • FGF4FZD1 1403 491 1193 2191 • CSNK1G1FGF4 1438 771 2389 995 • FGF4FZD7 1448 1364 1513 737 • FGF4SFRP1 1586 1507 637 2314 • FGF4NLK 1601 298 307 2394 • FGF4FZD8 1646 1716 1242 1626 • FGF4WNT5A 1700 333 1481 1660 • FGF4FZD6 1705 476 1717 2000 • FGF4FBXW4 1714 122 1583 769 • FGF4PITX2 1732 1848 2062 716 • FGF4LRP5 1795 1054 1958 967 • FGF4TCF7 1843 90 87 1967 • FGF4TLE2 1856 337 2475 529 • FGF4MYC 1867 109 256 1365 • FGF4PPP2CA 1870 85 1102 1331 • FGF4WNT4 1907 138 2057 802 • FGF4FSHB 2023 537 1133 1754 • FGF4WNT1 2047 114 1212 569 • FGF4PORCN 2104 266 1462 1894 • FGF4FRZB 2119 183 579 1038 • FGF4SLC9A3R1 2132 474 472 2430 • FGF4TLE1 2142 452 1415 1878 • FGF4GSK3A 2145 1055 677 982 • FGF4JUN 2173 325 1306 1234 • FGF4WIF1 2178 866 1224 1609 • FGF4FOSL1 2234 762 1000 1646 • FGF4NKD1 2252 59 1155 1658 • FGF4FRAT1 2266 394 834 921 • FGF4WNT2 2269 378 1406 2130 • FGF4FOXN1 2386 228 1476 1116 • FGF4PYGO1 2389 282 914 1159 • FGF4WNT3A 2395 679 1962 1597 • FGF4WNT2B 2419 512 350 1577 • FGF4KREMEN1 2459 342 833 2120 • FGF4T 2460 278 623 400 • FGF4WNT3 2475 1912 214 449 • FGF4RHOU 2480 587 2109 726

## FOSL1 2*^nd^* order interaction ranking at different time points

• FOSL1GSK3A 73 1202 62 1115 461 • FOSL1PORCN 93 1875 294 311 120 • FOSL1FOXN1 123 2435 17 484 872 • FOSL1WNT1 143 2476 55 325 61 • FOSL1FZD2 159 854 367 208 989 • FOSL1LEF1 163 439 342 1661 1204 • FOSL1PYGO1 165 1492 182 453 654 • FOSL1MYC 227 2171 40 776 1746 • FOSL1SFRP1 230 261 1567 1911 223 • FOSL1WNT3 233 796 526 227 218 • FOSL1JUN 240 1448 988 2026 1768 • FOSL1T 292 1253 266 1226 1235 • FOSL1WIF1 295 2329 1693 2425 251 • FOSL1LRP6 326 2095 111 834 60 • DKK1FOSL1 404 2205 1798 699 2280 • FOSL1TLE1 449 1415 951 694 213 • FOSL1SFRP4 481 1096 1226 1645 2115 • FOSL1NKD1 495 2311 330 1072 1097 • FOSL1FBXW4 541 2233 215 1847 276 • FOSL1SLC9A3R1 553 1225 356 1846 1574 • FOSL1PPP2R1A 575 1617 298 845 244 • FOSL1KREMEN1 587 1541 710 429 30 • FOSL1FZD8 593 1516 876 199 660 • FOSL1FZD6 615 812 83 2065 1424 • FOSL1WNT2 626 549 933 2130 738 • FOSL1PITX2 645 701 672 93 408 • FOSL1FSHB 720 1588 33 262 1780 • FOSL1WNT5A 728 306 188 943 183 • FOSL1PPP2CA 746 2176 24 377 1408 • FOSL1RHOU 780 1043 593 974 561 • FOSL1WNT3A 784 691 1053 1439 1054 • FOSL1TCF7 789 1761 60 79 1133 • FOSL1NLK 833 1486 16 935 280 • FOSL1WNT4 840 765 478 2323 1564 • FOSL1LRP5 866 168 79 1292 1856 • APCFOSL1 882 695 2432 55 2093 • FOSL1FZD7 897 2059 1663 1956 704 • FOSL1WNT2B 917 558 30 749 1087 • FOSL1FRAT1 972 2151 751 1971 904 • FOSL1GSK3B 976 2305 872 313 123 • FOSL1TLE2 999 181 1002 2397 842 • FOSL1FZD1 1060 1458 779 1014 2230 • FBXW11FOSL1 1175 661 1044 867 1802 • CSNK1DFOSL1 1233 1920 2451 1558 1787 • CSNK2A1FOSL1 1240 648 1392 541 1441 • DIXDC1FOSL1 1242 34 299 1927 1573 • FOSL1TCF7L1 1328 2216 736 1968 2439 • EP300FOSL1 1476 795 2103 2403 2459 • FZD5FOSL1 1478 1298 2157 1051 1919 • FBXW2FOSL1 1570 336 2085 1111 838 • CXXC4FOSL1 1658 207 2277 1958 2140 • CCND3FOSL1 1660 1862 1704 731 915 • CTBP1FOSL1 1728 657 2244 2331 2269 • DAAM1FOSL1 1736 518 2152 2096 2274 • AXIN1FOSL1 1948 2100 1224 736 892 • CSNK1G1FOSL1 1962 1860 1651 513 984 • DVL2FOSL1 1974 130 2142 143 2176 • CTNNB1FOSL1 1997 1036 2065 975 1827 • CTBP2FOSL1 2033 2156 1904 2438 1233 • DVL1FOSL1 2109 425 1196 395 547 • AESFOSL1 2138 2079 2245 140 1374 • CTNNBIP1FOSL1 2174 664 1571 14 1078 • CSNK1A1FOSL1 2226 6 1804 406 253 • BTRCFOSL1 2270 1163 1248 506 1808 • FGF4FOSL1 2279 284 1635 2232 1702 • FOSL1FRZB 2314 1396 1026 912 1695 • CCND2FOSL1 2323 465 2094 1304 1614 • FOSL1SENP2 2338 2209 1138 1400 122 • CCND1FOSL1 2369 1647 2468 768 857 • BCL9FOSL1 2383 144 2080 1195 312

## FOSL1 2*^nd^* order interaction ranking at different durations

• FOSL1SENP2 114 1139 1632 1073 • FZD5FOSL1 161 1882 1878 1981 • FOSL1SFRP4 207 1214 617 145 • FOSL1LRP6 433 1422 803 1800 • FOSL1LEF1 516 946 2189 1674 • FOSL1PITX2 520 1216 1761 1285 • FOSL1FZD2 556 387 1963 1305 • FOSL1PPP2R1A 613 212 1052 2311 • FOSL1FZD8 615 2028 1334 2416 • FOSL1GSK3B 616 1480 995 947 • FOSL1FZD1 623 2192 2234 2429 • CSNK1A1FOSL1 643 417 2401 1600 • FOSL1SFRP1 660 1092 1934 1005 • FOSL1WNT5A 691 231 1864 1751 • FOSL1FZD7 701 1497 571 956 • FOSL1NLK 717 262 390 1476 • FOSL1TCF7L1 729 1225 793 752 • FOSL1PPP2CA 789 316 2481 2423 • CTNNB1FOSL1 814 1160 1020 516 • FOSL1FZD6 821 496 1541 155 • AESFOSL1 885 2006 1353 358 • CCND2FOSL1 888 1383 1994 2006 • CTNNBIP1FOSL1 893 1816 1014 1118 • DIXDC1FOSL1 967 548 323 1386 • FOSL1TLE1 968 845 1403 671 • FOSL1TCF7 969 353 727 2110 • FOSL1TLE2 982 332 362 1109 • CTBP1FOSL1 1006 1673 1333 2452 • FOSL1WNT4 1060 307 751 1447 • CSNK1DFOSL1 1064 1413 2460 1440 • BCL9FOSL1 1078 1403 1315 364 • FOSL1FBXW4 1087 309 2408 1259 • AXIN1FOSL1 1092 2305 840 424 • CCND1FOSL1 1138 2116 299 1844 • FOSL1LRP5 1142 746 1182 838 • FOSL1FRZB 1234 318 203 2331 • BTRCFOSL1 1324 1990 1098 1342 • DVL1FOSL1 1367 2253 561 613 • DVL2FOSL1 1381 1391 1514 1613 • FOSL1FSHB 1423 1230 2181 274 • EP300FOSL1 1462 1236 1601 1446 • FOSL1NKD1 1512 108 477 2160 • APCFOSL1 1521 1235 2345 923 • FOSL1GSK3A 1533 1137 1522 1087 • FOSL1MYC 1569 369 1719 1759 • FOSL1WNT3A 1578 543 1535 2158 • FOSL1WIF1 1592 467 2318 169 • CCND3FOSL1 1604 2232 967 1607 • FOSL1SLC9A3R1 1605 534 2149 1605 • FOSL1WNT1 1649 219 2484 1766 • FOSL1WNT2 1669 352 1773 1856 • FOSL1JUN 1674 206 2221 910 • CTBP2FOSL1 1676 2237 1734 2205 • FOSL1KREMEN1 1697 209 1626 763 • FOSL1FOXN1 1718 188 1531 650 • CXXC4FOSL1 1776 2147 2417 1415 • FOSL1FRAT1 1817 303 786 2315 • DKK1FOSL1 1826 1535 2328 2239 • FOSL1T 1888 453 2471 1015 • DAAM1FOSL1 1898 1902 171 2210 • FOSL1WNT2B 1949 768 962 185 • FOSL1PYGO1 2038 276 2170 1620 • FBXW11FOSL1 2072 2194 892 1566 • FOSL1WNT3 2090 810 525 1056 • FBXW2FOSL1 2150 1859 1765 1457 • FGF4FOSL1 2234 762 1000 1646 • CSNK1G1FOSL1 2293 1939 1078 2174 • FOSL1PORCN 2299 315 1672 1874 • CSNK2A1FOSL1 2367 763 273 2202 • FOSL1RHOU 2447 646 2289 1585

## FOXN1 2*^nd^* order interaction ranking at different time points

• DKK1FOXN1 24 2405 1251 1434 404 • CSNK1DFOXN1 51 2149 868 1245 2365 • FZD5FOXN1 99 2379 89 45 1767 • CXXC4FOXN1 106 2260 148 349 1045 • FBXW11FOXN1 120 1501 346 224 2249 • FOSL1FOXN1 123 2435 17 484 872 • DIXDC1FOXN1 138 2044 10 687 1977 • DAAM1FOXN1 174 1147 745 993 1241 • APCFOXN1 195 2386 434 436 884 • CSNK2A1FOXN1 205 1624 645 1088 826 • CCND3FOXN1 232 1038 1711 2442 2397 • FBXW2FOXN1 269 531 558 1827 2134 • EP300FOXN1 300 898 882 997 1915 • CTBP1FOXN1 350 2271 149 322 1550 • CTNNB1FOXN1 351 2270 300 1168 938 • CSNK1G1FOXN1 385 1700 913 1749 2227 • CTBP2FOXN1 486 469 329 1586 868 • DVL2FOXN1 532 1451 372 2378 1828 • AXIN1FOXN1 546 1824 313 117 2424 • DVL1FOXN1 566 196 130 91 2098 • BTRCFOXN1 623 2218 1756 1721 1551 • CTNNBIP1FOXN1 658 2083 186 1729 309 • CSNK1A1FOXN1 871 624 324 1458 462 • FOXN1GSK3A 880 1274 2450 272 994 • FOXN1LEF1 885 872 2228 118 624 • CCND2FOXN1 899 557 69 970 1389 • AESFOXN1 1031 390 331 1815 1155 • FGF4FOXN1 1111 2013 18 1768 1129 • FOXN1WIF1 1171 891 2010 160 534 • FOXN1FZD2 1220 267 1806 86 1027 • FOXN1PYGO1 1244 852 2028 2371 1309 • FOXN1PORCN 1247 402 1697 925 473 • BCL9FOXN1 1374 2250 167 985 951 • FOXN1JUN 1406 2099 1418 1632 1905 • FOXN1WNT1 1436 1099 1465 1327 358 • FOXN1WNT3 1507 819 1546 2280 758 • FOXN1SFRP1 1513 424 1702 791 65 • FOXN1LRP6 1571 1157 1599 714 1217 • CCND1FOXN1 1577 1635 472 2481 1738 • FOXN1MYC 1595 437 1254 1303 1100 • FOXN1NKD1 1755 293 1018 502 1662 • FOXN1FBXW4 1861 414 2020 412 1359 • FOXN1T 1902 2359 1217 1438 2142 • FOXN1KREMEN1 1957 1505 2414 2056 1086 • FOXN1PITX2 1961 1236 1715 2160 871 • FOXN1FSHB 1972 1349 1848 148 1596 • FOXN1TCF7 1976 1780 888 707 808 • FOXN1TLE1 1995 605 1776 1514 129 • FOXN1WNT4 2039 2192 1366 154 1140 • FOXN1WNT5A 2079 1108 2378 230 2011 • FOXN1FZD8 2099 349 2111 2475 715 • FOXN1SLC9A3R1 2175 129 1675 1040 894 • FOXN1FZD6 2200 345 1094 18 1018 • FOXN1SFRP4 2202 2211 1833 495 694 • FOXN1WNT3A 2234 831 2124 1984 1242 • FOXN1WNT2B 2247 138 900 2138 2180 • FOXN1TLE2 2277 1368 1261 663 1625 • FOXN1WNT2 2296 1026 2362 178 1335 • FOXN1TCF7L1 2305 444 608 743 1132 • FOXN1LRP5 2325 1075 2045 883 1498 • FOXN1FRAT1 2329 959 865 438 1051 • FOXN1FZD7 2333 708 2113 172 590 • FOXN1GSK3B 2334 731 1382 2059 2219 • FOXN1NLK 2343 291 1372 741 184 • FOXN1PPP2CA 2359 1881 905 1788 1983 • FOXN1PPP2R1A 2363 214 1779 1437 112 • FOXN1RHOU 2392 640 1837 2444 1791 • FOXN1FZD1 2421 1477 961 410 2347 • FOXN1FRZB 2464 1678 1302 433 1594 • FOXN1SENP2 2473 427 1964 197 1578

## FOXN1 2*^nd^* order interaction ranking at different durations

• FOXN1SENP2 48 2041 316 73 • FOXN1SFRP4 91 1544 520 102 • FOXN1PPP2R1A 222 1346 1375 491 • FOXN1SFRP1 247 2189 2311 413 • CSNK1A1FOXN1 255 1457 1786 1352 • FOXN1FZD2 297 1966 1646 867 • FOXN1FZD7 330 1323 822 159 • FOXN1LRP6 357 1414 574 975 • FZD5FOXN1 367 692 1610 1098 • FOXN1TCF7L1 371 2468 788 931 • FOXN1LEF1 386 1433 1362 379 • FOXN1FZD1 387 2250 1660 2453 • FOXN1FZD6 403 2438 1335 154 • FOXN1WNT5A 463 1016 1983 951 • FOXN1NLK 467 665 262 1040 • FOXN1PPP2CA 529 736 364 1732 • FOXN1PITX2 568 1747 1226 668 • FOXN1FZD8 603 2174 1325 680 • FOXN1GSK3B 631 1917 1346 2226 • FOXN1TLE2 758 2416 602 1810 • FOXN1WNT4 796 443 2171 2136 • FOXN1FRZB 808 423 824 2121 • FOXN1TCF7 833 1179 338 1573 • FOXN1WNT2 903 896 1559 1435 • FOXN1TLE1 913 1450 1819 583 • FOXN1LRP5 914 1857 1759 2221 • DIXDC1FOXN1 958 17 567 289 • FOXN1FBXW4 1024 1417 1822 526 • CTBP1FOXN1 1037 1936 2015 2383 • FOXN1WNT1 1057 781 2320 1665 • FOXN1SLC9A3R1 1090 1107 2322 581 • CTNNBIP1FOXN1 1110 1511 1432 684 • AESFOXN1 1167 1011 819 1449 • FOXN1NKD1 1185 741 407 345 • FOXN1MYC 1207 787 1297 882 • FOXN1WIF1 1208 855 1307 130 • CTNNB1FOXN1 1244 1392 1316 1149 • FOXN1T 1278 1082 1368 2372 • FOXN1GSK3A 1290 1079 1909 149 • CCND1FOXN1 1292 655 1299 1192 • FOXN1FRAT1 1317 1314 1704 723 • BCL9FOXN1 1325 2078 2177 1802 • CCND2FOXN1 1331 749 621 817 • AXIN1FOXN1 1337 830 1586 940 • FOXN1FSHB 1345 1609 1972 22 • FOXN1WNT3A 1468 1877 2114 1002 • FOXN1JUN 1476 1653 2219 2176 • BTRCFOXN1 1574 846 1894 834 • CTBP2FOXN1 1651 774 1241 1515 • EP300FOXN1 1653 542 1247 1110 • APCFOXN1 1666 1886 1427 1261 • CSNK1DFOXN1 1703 503 1239 2067 • FOSL1FOXN1 1718 188 1531 650 • CCND3FOXN1 1883 2408 1180 1082 • FOXN1WNT2B 1889 1798 717 111 • FOXN1KREMEN1 1905 1324 2031 1901 • DVL1FOXN1 1952 1335 137 180 • FOXN1WNT3 1973 1083 108 480 • DVL2FOXN1 2027 1424 767 2282 • DKK1FOXN1 2089 302 2342 2464 • FBXW11FOXN1 2107 389 813 2188 • FBXW2FOXN1 2130 589 2162 1841 • CSNK2A1FOXN1 2162 194 303 742 • FOXN1PORCN 2165 2015 1475 844 • CXXC4FOXN1 2180 1828 2206 796 • FOXN1PYGO1 2203 1871 1693 663 • DAAM1FOXN1 2253 1385 239 855 • CSNK1G1FOXN1 2369 372 972 2116 • FGF4FOXN1 2386 228 1476 1116 • FOXN1RHOU 2415 1159 1718 847

## FRAT1 2*^nd^* order interaction ranking at different time points

• FRAT1PYGO1 56 2336 339 783 1213 • FRAT1FZD2 153 2202 2185 87 2073 • FRAT1GSK3A 161 1129 194 2043 1807 • FRAT1PORCN 183 2187 924 746 250 • FRAT1SFRP1 197 2157 394 1335 505 • FRAT1WNT1 200 2131 592 709 440 • FRAT1WIF1 221 2152 2344 184 1269 • FRAT1WNT3 252 2372 1439 1524 1058 • FRAT1JUN 313 1270 686 2318 2020 • FRAT1LRP6 316 2224 616 1259 181 • FRAT1T 340 2150 283 1220 2282 • FRAT1MYC 364 2081 714 1707 2334 • FRAT1LEF1 369 2286 1415 470 2113 • FRAT1NKD1 433 2180 232 2351 2107 • DKK1FRAT1 439 2042 1573 578 1096 • FRAT1TLE1 539 1849 2202 347 1273 • FRAT1FSHB 595 2304 238 362 2108 • FRAT1KREMEN1 681 2395 1928 170 554 • FRAT1PITX2 689 1610 1673 221 613 • APCFRAT1 704 2355 1872 1937 2391 • FRAT1FZD8 745 2344 1974 212 787 • FRAT1FZD6 757 1720 268 1629 574 • FRAT1SLC9A3R1 783 2272 1887 1682 242 • FRAT1PPP2CA 792 2225 98 1041 1142 • FRAT1SFRP4 806 1275 2203 217 2007 • FRAT1FBXW4 831 2047 491 468 653 • FRAT1WNT5A 849 1874 355 805 540 • FRAT1NLK 900 2020 366 1512 517 • FRAT1TCF7 915 2075 74 2153 1996 • FBXW11FRAT1 932 1177 1783 245 1651 • FRAT1PPP2R1A 956 1900 1269 1611 197 • FOSL1FRAT1 972 2151 751 1971 904 • FRAT1TLE2 988 1078 926 1991 1592 • FRAT1WNT4 991 122 623 1882 1507 • FRAT1WNT2 1019 1653 2179 1746 1786 • FRAT1WNT2B 1046 2107 443 563 1361 • FRAT1WNT3A 1054 1951 1739 1029 1486 • FRAT1GSK3B 1058 2294 1754 880 859 • FRAT1LRP5 1144 1050 814 2267 2102 • DIXDC1FRAT1 1154 1708 193 2387 2171 • FZD5FRAT1 1169 1414 1574 2145 2069 • FRAT1FZD7 1180 2178 2330 509 458 • CSNK1DFRAT1 1181 1175 1017 616 1959 • CSNK2A1FRAT1 1190 802 2155 1453 837 • EP300FRAT1 1292 1560 2240 1649 2440 • FRAT1RHOU 1334 1609 2104 1022 361 • FRAT1FZD1 1363 1707 496 911 847 • FRAT1TCF7L1 1427 2111 879 875 666 • CXXC4FRAT1 1448 869 998 1954 1935 • FBXW2FRAT1 1641 1422 2017 1002 2184 • CCND3FRAT1 1655 1381 2452 1382 1947 • DAAM1FRAT1 1674 2290 1787 1798 1804 • CTBP1FRAT1 1742 1825 943 952 1003 • CTNNB1FRAT1 1747 1321 1664 960 1370 • CSNK1G1FRAT1 1900 2023 2484 1024 400 • DVL2FRAT1 1909 1003 1131 1305 1887 • CTNNBIP1FRAT1 1964 2195 701 1401 706 • BTRCFRAT1 2027 1203 2365 1021 2454 • CTBP2FRAT1 2052 2377 2129 1145 2267 • CCND2FRAT1 2116 963 1202 618 1763 • AXIN1FRAT1 2145 1466 1503 72 1991 • DVL1FRAT1 2164 1593 1161 1413 1531 • BCL9FRAT1 2239 325 1811 125 2380 • AESFRAT1 2245 923 1761 207 1454 • FRAT1FRZB 2289 1537 2439 1804 1782 • FGF4FRAT1 2321 2058 317 1776 2408 • FOXN1FRAT1 2329 959 865 438 1051 • CSNK1A1FRAT1 2367 1006 2286 2453 972 • CCND1FRAT1 2385 1310 1832 1509 1995 • FRAT1SENP2 2435 2342 1501 1489 354

## FRAT1 2*^nd^* order interaction ranking at different durations

• FRAT1SENP2 31 2132 1509 118 • FRAT1SFRP4 73 1120 1996 197 • FRAT1LEF1 155 2334 1507 291 • FRAT1LRP6 212 2458 1138 1616 • FRAT1SFRP1 217 1957 2451 402 • FRAT1PPP2R1A 238 721 1472 1069 • CSNK1A1FRAT1 251 1050 1935 1921 • FRAT1PITX2 298 2131 1245 2263 • FRAT1FZD2 341 991 827 1603 • FRAT1FZD7 365 1202 859 128 • FRAT1FZD6 384 1528 1125 117 • FRAT1TCF7L1 400 1306 583 2015 • FRAT1FZD1 437 1348 1383 328 • FRAT1GSK3B 453 1714 1154 2336 • FRAT1FZD8 472 1150 1269 2102 • FRAT1WNT5A 505 552 862 1990 • FRAT1NLK 535 385 130 835 • FRAT1FBXW4 561 376 1783 643 • FRAT1TLE1 567 2230 955 832 • FRAT1PPP2CA 570 501 891 1529 • FRAT1WNT4 612 480 2242 1290 • FZD5FRAT1 617 775 1210 1579 • FRAT1TCF7 662 678 725 300 • FRAT1LRP5 674 1865 875 1876 • FRAT1FRZB 739 704 259 1786 • FRAT1TLE2 746 1570 1544 934 • DIXDC1FRAT1 788 96 469 338 • CTBP1FRAT1 835 2306 1816 1986 • FRAT1WNT2 912 844 2252 1162 • FRAT1NKD1 930 311 584 135 • FRAT1SLC9A3R1 943 1447 1664 456 • FRAT1WIF1 1017 2481 1955 213 • FRAT1WNT1 1020 560 2253 1320 • CCND1FRAT1 1102 2073 611 936 • FRAT1GSK3A 1108 1566 2178 440 • AESFRAT1 1141 2113 823 2310 • CTNNBIP1FRAT1 1160 2210 814 506 • FRAT1MYC 1268 1072 790 1656 • AXIN1FRAT1 1284 823 522 885 • FRAT1FSHB 1310 1985 1356 116 • FOXN1FRAT1 1317 1314 1704 723 • CCND2FRAT1 1320 1238 1317 1375 • BCL9FRAT1 1352 1540 2268 741 • FRAT1KREMEN1 1365 597 1574 2326 • CSNK1DFRAT1 1384 672 2240 1789 • FRAT1WNT3A 1396 904 1581 1276 • CTNNB1FRAT1 1483 1704 1519 2347 • FRAT1T 1614 1042 1569 749 • FRAT1JUN 1677 915 1354 1048 • BTRCFRAT1 1685 1437 2159 1227 • CTBP2FRAT1 1716 951 1762 913 • FRAT1WNT3 1746 2136 815 960 • FOSL1FRAT1 1817 303 786 2315 • APCFRAT1 1822 1743 2367 2382 • FRAT1WNT2B 1833 2138 743 247 • CXXC4FRAT1 1886 1451 2067 458 • FBXW2FRAT1 1896 2066 1518 637 • EP300FRAT1 1910 1750 2166 689 • FRAT1PORCN 1941 1078 2145 920 • DVL1FRAT1 1950 1994 289 430 • CCND3FRAT1 1980 1462 511 1034 • DVL2FRAT1 1985 2196 1255 1963 • DAAM1FRAT1 2043 913 291 970 • FRAT1RHOU 2091 1208 2157 1782 • DKK1FRAT1 2116 279 2346 2386 • CSNK2A1FRAT1 2128 2186 325 615 • FGF4FRAT1 2266 394 834 921 • FBXW11FRAT1 2281 2204 2130 2146 • CSNK1G1FRAT1 2373 355 710 2061 • FRAT1PYGO1 2404 1325 2334 783

## FRZB 2*^nd^* order interaction ranking at different time points

• FRZBSFRP1 4 1416 1440 1205 217 • FRZBGSK3A 5 1482 90 1544 403 • FRZBFZD2 9 930 923 955 10 • FRZBPYGO1 10 2291 243 346 113 • FRZBJUN 11 2412 644 507 409 • FRZBLEF1 12 629 987 1090 1958 • FRZBPORCN 13 1981 800 1938 85 • FRZBWIF1 18 2483 218 1864 608 • FRZBMYC 23 2341 320 206 501 • FRZBLRP6 26 1067 123 2202 294 • FRZBWNT1 28 2478 63 528 144 • FRZBWNT3 29 2155 630 1213 153 • FRZBNKD1 48 2430 159 1677 1491 • FRZBT 49 2398 263 900 1130 • FRZBTLE1 52 1914 1270 2055 1385 • FRZBFSHB 62 355 210 588 2037 • FRZBFZD6 64 1953 44 2054 610 • FRZBPITX2 66 1794 947 1256 985 • FRZBPPP2R1A 67 2232 413 806 178 • FRZBFBXW4 68 2391 495 1781 171 • FRZBWNT4 76 1273 155 2047 1647 • FRZBSLC9A3R1 77 2230 1013 2165 1334 • FRZBPPP2CA 78 2468 5 1286 226 • FRZBFZD8 88 2113 1353 1131 373 • FRZBTCF7 101 2467 241 2087 866 • FRZBGSK3B 107 2165 470 1773 1518 • FRZBWNT3A 114 1233 1049 1780 145 • FRZBSFRP4 121 775 1514 1484 524 • FRZBWNT2 131 520 796 579 1608 • FRZBWNT5A 135 2354 235 1726 518 • FRZBKREMEN1 139 1894 471 1101 623 • FRZBRHOU 141 1844 896 772 1079 • FRZBTLE2 155 1945 691 980 887 • FRZBWNT2B 158 1756 27 1589 157 • FRZBFZD1 173 1890 727 46 879 • FRZBLRP5 175 1511 309 1019 446 • FRZBNLK 203 2317 140 1844 187 • FRZBFZD7 311 2241 1529 811 635 • FRZBTCF7L1 349 2309 348 1223 1940 • FRZBSENP2 1525 2475 1121 858 452 • DKK1FRZB 1727 1704 2225 1148 2022 • APCFRZB 2106 958 2211 17 1291 • FBXW11FRZB 2118 1065 2214 101 2457 • CSNK2A1FRZB 2127 859 1805 891 1483 • FRAT1FRZB 2289 1537 2439 1804 1782 • FZD5FRZB 2303 506 1000 1185 2343 • EP300FRZB 2308 137 1891 967 1472 • FOSL1FRZB 2314 1396 1026 912 1695 • CCND3FRZB 2337 1895 1212 633 988 • CSNK1DFRZB 2386 1933 2394 1299 2183 • DIXDC1FRZB 2388 316 742 2440 1396 • CSNK1G1FRZB 2397 231 2222 1939 707 • DAAM1FRZB 2398 1281 1943 1450 1865 • CTBP1FRZB 2412 575 1772 1692 961 • CXXC4FRZB 2413 1766 2333 1272 1880 • DVL1FRZB 2414 1080 1528 1465 956 • DVL2FRZB 2416 1342 1545 1206 1164 • FBXW2FRZB 2425 2345 1642 1273 1867 • CTNNBIP1FRZB 2428 264 2043 2340 121 • CTNNB1FRZB 2436 809 2458 1883 1817 • AXIN1FRZB 2440 1308 2256 1478 1244 • AESFRZB 2451 1899 2016 307 199 • CTBP2FRZB 2453 2226 1112 261 1838 • BCL9FRZB 2455 421 2371 138 1481 • BTRCFRZB 2457 2356 1615 29 2078 • FGF4FRZB 2460 876 1584 485 33 • CCND2FRZB 2461 764 2310 1854 1963 • CSNK1A1FRZB 2462 560 1884 167 325 • FOXN1FRZB 2464 1678 1302 433 1594 • CCND1FRZB 2475 1944 2196 2146 1400

## FRZB 2*^nd^* order interaction ranking at different durations

• FZD5FRZB 19 525 343 963 • CSNK1A1FRZB 80 2231 2485 1638 • FRZBSENP2 200 2104 925 50 • CTBP1FRZB 356 1950 1380 1736 • DIXDC1FRZB 383 4 129 249 • FRZBSFRP4 454 2249 2397 12 • CTNNB1FRZB 469 2053 411 916 • CCND2FRZB 666 2340 715 381 • BCL9FRZB 672 1025 974 675 • FRZBFZD2 703 1026 1634 864 • FRZBLEF1 707 2051 1588 292 • FRZBLRP6 708 2084 2208 437 • CSNK1DFRZB 730 329 1534 2192 • FRAT1FRZB 739 704 259 1786 • FRZBSFRP1 791 1894 1083 53 • AXIN1FRZB 797 404 450 284 • AESFRZB 804 1002 742 1545 • FOXN1FRZB 808 423 824 2121 • CCND1FRZB 818 568 213 1770 • BTRCFRZB 839 441 1977 1908 • FRZBFZD7 840 1841 1262 54 • CTNNBIP1FRZB 889 2221 230 248 • FRZBNLK 919 887 1881 141 • FRZBPITX2 922 1138 1612 942 • FRZBPPP2R1A 924 1119 2374 1727 • APCFRZB 939 2436 2327 1767 • FRZBGSK3B 949 1834 663 601 • FRZBTCF7L1 957 1843 2335 1190 • FRZBWNT5A 988 1687 1593 944 • FRZBFZD1 993 1273 2458 1570 • FRZBFZD6 1004 2479 1448 33 • CTBP2FRZB 1030 766 1187 728 • FRZBPPP2CA 1033 541 2381 2106 • DVL1FRZB 1040 1149 128 258 • FRZBFZD8 1072 1944 1069 1774 • FRZBTCF7 1121 2074 708 469 • EP300FRZB 1218 1628 300 270 • FRZBTLE2 1228 2226 992 642 • FOSL1FRZB 1234 318 203 2331 • DKK1FRZB 1273 48 850 1599 • FRZBWNT4 1319 782 1017 2153 • FRZBLRP5 1341 1693 1285 2463 • DVL2FRZB 1373 838 495 237 • CCND3FRZB 1408 1639 146 865 • FRZBNKD1 1464 612 2077 251 • FRZBFBXW4 1473 1301 1964 55 • FRZBTLE1 1497 1141 1263 226 • DAAM1FRZB 1532 798 492 544 • FRZBWNT3A 1551 1847 1490 1659 • CXXC4FRZB 1568 1897 2185 576 • CSNK2A1FRZB 1627 606 296 1080 • FRZBGSK3A 1629 2038 1392 397 • FRZBWNT2 1650 1048 2016 1306 • FRZBSLC9A3R1 1681 1152 2223 422 • FRZBWNT1 1711 723 2443 1099 • FRZBMYC 1744 1520 1502 265 • FRZBFSHB 1752 1142 2273 43 • FBXW11FRZB 1764 446 1407 1146 • FRZBWIF1 1770 1839 1974 147 • CSNK1G1FRZB 1809 121 1645 1060 • FRZBJUN 1838 2162 1768 1482 • FBXW2FRZB 1887 709 989 1314 • FRZBKREMEN1 1979 2274 1619 1790 • FGF4FRZB 2119 183 579 1038 • FRZBT 2143 1291 1563 1583 • FRZBWNT2B 2220 1455 1809 16 • FRZBWNT3 2224 1874 2455 435 • FRZBPORCN 2333 2401 1607 691 • FRZBRHOU 2390 1218 1650 600 • FRZBPYGO1 2435 813 2123 1220

## FSHB 2*^nd^* order interaction ranking at different time points

• FRZBFSHB 62 355 210 588 2037 • DKK1FSHB 146 1423 1909 385 2047 • FSHBGSK3A 303 307 1173 487 641 • FSHBPYGO1 330 200 1847 2263 682 • FSHBFZD2 337 1393 2426 986 720 • APCFSHB 399 1318 278 420 962 • FSHBMYC 413 2306 996 1170 2023 • FSHBSFRP1 426 1241 1620 304 161 • FSHBWIF1 460 2261 2071 762 1098 • FSHBJUN 466 1786 1289 233 182 • FSHBWNT1 471 2394 761 928 234 • FSHBWNT3 506 1255 2009 378 100 • FSHBLEF1 508 616 1870 36 1041 • FSHBPORCN 515 164 1990 155 265 • CSNK1DFSHB 583 659 345 48 2258 • FRAT1FSHB 595 2304 238 362 2108 • DIXDC1FSHB 646 1959 1 653 2155 • FBXW11FSHB 651 1546 1867 1011 2263 • CSNK2A1FSHB 699 384 2053 1936 785 • FOSL1FSHB 720 1588 33 262 1780 • FSHBLRP6 762 1626 2215 1479 780 • FSHBNKD1 771 2096 2259 793 1924 • FZD5FSHB 802 915 144 22 1672 • FSHBT 803 1941 1461 719 1372 • CCND3FSHB 842 1814 887 753 49 • CTBP1FSHB 857 1027 231 840 2418 • CXXC4FSHB 914 1402 110 1027 1299 • EP300FSHB 984 361 451 176 2248 • DAAM1FSHB 989 1091 286 459 2189 • FSHBPITX2 1068 344 1527 1427 1562 • FSHBTLE1 1075 1418 2242 1568 685 • FSHBKREMEN1 1122 279 1004 2296 365 • FBXW2FSHB 1161 118 571 611 1029 • FSHBFBXW4 1165 1304 990 976 2101 • FSHBWNT2 1202 57 2398 319 1727 • FSHBSFRP4 1239 2168 2096 164 1260 • FSHBSLC9A3R1 1276 759 2121 1343 1306 • FSHBPPP2CA 1302 1918 1417 659 2290 • CTNNB1FSHB 1312 198 387 1295 1654 • FSHBWNT5A 1344 1383 1747 1738 148 • FSHBNLK 1430 2219 2361 1868 375 • FSHBFZD8 1435 841 2239 2245 482 • CTBP2FSHB 1479 780 509 244 1248 • FSHBFZD6 1484 519 1108 473 784 • FSHBWNT3A 1490 689 1505 2355 1698 • FSHBGSK3B 1516 2167 2254 2206 498 • FSHBWNT4 1517 1564 1969 888 1917 • FSHBTCF7 1543 1260 1113 2024 856 • FSHBTLE2 1586 980 1659 562 1619 • FSHBLRP5 1600 410 2266 2455 557 • BTRCFSHB 1621 281 1860 2152 2017 • FSHBPPP2R1A 1630 475 1889 1497 383 • FSHBWNT2B 1640 2393 1443 879 2041 • AXIN1FSHB 1643 2251 475 182 2087 • CTNNBIP1FSHB 1648 1346 91 946 747 • CSNK1G1FSHB 1676 676 1859 1896 925 • FSHBFZD7 1680 1668 1681 278 1222 • DVL1FSHB 1684 1018 217 321 2436 • AESFSHB 1694 723 374 2191 1809 • FSHBRHOU 1724 1623 2177 2170 391 • DVL2FSHB 1732 110 602 1161 2330 • FSHBFZD1 1739 1832 1918 613 2095 • CSNK1A1FSHB 1846 697 227 1674 1512 • BCL9FSHB 1869 114 139 1086 1261 • FGF4FSHB 1892 26 76 239 1703 • FSHBTCF7L1 1895 1449 2343 1015 902 • CCND2FSHB 1915 464 583 522 1729 • FOXN1FSHB 1972 1349 1848 148 1596 • CCND1FSHB 2262 1906 64 1835 488 • FSHBSENP2 2441 1138 1662 132 832

## FSHB 2*^nd^* order interaction ranking at different durations

• FSHBSENP2 79 625 458 1122 • FSHBSFRP4 107 829 1792 1363 • FSHBLRP6 214 1700 2217 1675 • FSHBLEF1 219 1444 1318 1486 • FSHBFZD7 344 1579 700 1863 • FSHBPPP2R1A 346 663 2087 1791 • FSHBFZD2 409 593 2049 416 • FSHBFZD6 426 751 986 1441 • FSHBPITX2 432 1304 1925 362 • FSHBTCF7L1 451 2374 527 1397 • FSHBFZD1 460 1685 2175 904 • FSHBGSK3B 494 2432 1630 272 • FSHBSFRP1 507 1553 1795 860 • FSHBFZD8 588 2024 1235 974 • FSHBWNT5A 626 349 704 1725 • FSHBTLE2 645 358 1740 126 • CSNK1A1FSHB 650 2289 2124 100 • FSHBNLK 652 207 1060 1427 • FSHBPPP2CA 659 383 953 406 • FZD5FSHB 725 907 1565 1267 • FSHBTCF7 736 473 370 1014 • FSHBFBXW4 763 214 2395 2118 • FSHBWNT4 764 334 1644 932 • FSHBTLE1 856 731 2354 1743 • FSHBLRP5 952 1365 1371 90 • CTNNBIP1FSHB 989 2329 1157 1282 • CCND1FSHB 995 2060 292 756 • DIXDC1FSHB 1031 414 578 1411 • CTNNB1FSHB 1045 1868 1275 204 • FSHBGSK3A 1084 1493 1880 1832 • FSHBNKD1 1104 170 359 1510 • CCND2FSHB 1122 1565 2174 507 • AESFSHB 1165 924 1244 68 • BCL9FSHB 1196 2413 1642 1694 • AXIN1FSHB 1202 1531 489 1167 • FSHBWNT1 1212 267 2260 438 • CSNK1DFSHB 1221 2012 994 96 • FSHBWIF1 1251 930 1156 1155 • CTBP1FSHB 1252 2124 2369 296 • FSHBWNT2 1294 683 543 890 • FRAT1FSHB 1310 1985 1356 116 • FOXN1FSHB 1345 1609 1972 22 • FSHBSLC9A3R1 1395 521 2277 1939 • FOSL1FSHB 1423 1230 2181 274 • FSHBWNT3A 1424 938 1732 535 • FSHBMYC 1435 280 319 1302 • FSHBWNT2B 1501 807 698 1487 • FSHBT 1597 558 1186 398 • DVL1FSHB 1610 1942 589 1429 • BTRCFSHB 1632 849 2204 351 • FSHBKREMEN1 1656 218 1188 1101 • CTBP2FSHB 1719 1656 2014 880 • FSHBPORCN 1722 460 1310 1187 • FSHBJUN 1734 348 2290 407 • CCND3FSHB 1741 1492 809 370 • FRZBFSHB 1752 1142 2273 43 • FSHBWNT3 1763 2171 1426 1275 • EP300FSHB 1793 2258 1976 1022 • DVL2FSHB 1805 1279 1810 928 • DKK1FSHB 1819 909 1229 538 • APCFSHB 1930 1760 473 771 • CSNK2A1FSHB 1993 2036 345 683 • FSHBPYGO1 2021 660 2416 468 • FGF4FSHB 2023 537 1133 1754 • FSHBRHOU 2053 667 2271 454 • DAAM1FSHB 2141 2097 811 1051 • CXXC4FSHB 2172 1287 1294 1353 • FBXW11FSHB 2318 1373 2232 780 • FBXW2FSHB 2363 1812 1515 245 • CSNK1G1FSHB 2364 952 2477 1114

## JUN 2*^nd^* order interaction ranking at different time points

• FRZBJUN 11 2412 644 507 409 • DKK1JUN 17 2148 1629 864 1279 • FZD1JUN 43 1296 1287 2106 7 • APCJUN 61 1317 1857 701 2309 • FZD5JUN 94 741 1033 1110 2013 • FBXW2JUN 110 1284 2462 545 1631 • GSK3BJUN 133 1510 851 1724 1168 • CXXC4JUN 140 2439 1222 1454 1984 • DIXDC1JUN 145 878 200 2235 1485 • FBXW11JUN 156 332 1707 1135 2405 • FZD6JUN 179 1265 2243 1910 1013 • FOSL1JUN 240 1448 988 2026 1768 • FZD7JUN 241 1223 1279 2162 1665 • CSNK1DJUN 246 2221 2422 897 1316 • CCND3JUN 247 746 2338 290 318 • DAAM1JUN 249 66 1542 2294 1526 • FZD8JUN 259 1289 852 284 1451 • CSNK2A1JUN 270 1151 2192 2372 1908 • CTBP2JUN 285 567 1396 425 1806 • FRAT1JUN 313 1270 686 2318 2020 • CTNNB1JUN 390 798 2100 1934 2260 • CTBP1JUN 405 2160 878 1365 1747 • EP300JUN 428 65 1419 1281 2150 • DVL2JUN 440 486 1575 179 1845 • FSHBJUN 466 1786 1289 233 182 • AXIN1JUN 572 1361 1187 1331 1459 • CSNK1G1JUN 608 1606 2217 1296 1456 • CTNNBIP1JUN 632 1390 1614 1543 558 • DVL1JUN 644 46 2029 1394 844 • BTRCJUN 694 699 2036 754 1313 • AESJUN 772 1512 1373 333 1826 • CSNK1A1JUN 832 321 1169 1777 1201 • JUNPYGO1 853 1183 845 1285 592 • CCND1JUN 942 1578 797 2143 617 • JUNWNT1 1000 985 328 270 35 • JUNMYC 1014 206 489 655 1946 • CCND2JUN 1043 1643 1473 2329 559 • FGF4JUN 1203 1288 776 1591 1090 • BCL9JUN 1230 1682 1170 337 968 • FZD2JUN 1377 638 680 956 970 • FOXN1JUN 1406 2099 1418 1632 1905 • JUNSFRP1 1492 548 1921 404 24 • GSK3AJUN 1554 233 2047 609 869 • JUNLEF1 1563 508 2110 639 542 • JUNNKD1 1615 1250 428 695 410 • JUNWIF1 1637 1650 2022 2049 1268 • JUNPORCN 1699 1005 1148 1563 23 • JUNLRP6 1702 195 1477 2262 80 • JUNWNT3 1738 32 1721 2458 54 • JUNTLE1 1814 1948 1902 2090 83 • JUNT 1873 939 596 371 1249 • JUNPITX2 2013 126 1967 738 31 • JUNFBXW4 2029 511 107 2393 379 • JUNPPP2CA 2042 341 99 860 1430 • JUNSFRP4 2135 1701 2070 26 4 • JUNWNT2 2161 507 1458 1010 435 • JUNNLK 2182 2046 414 2085 509 • JUNLRP5 2183 711 511 1505 271 • JUNKREMEN1 2199 1452 1038 1579 104 • JUNSLC9A3R1 2264 1532 2236 1999 649 • JUNPPP2R1A 2265 569 614 1908 42 • JUNWNT3A 2288 467 2002 1880 800 • JUNWNT5A 2319 1401 659 1922 98 • JUNTLE2 2345 376 1517 1338 277 • JUNWNT4 2348 2102 1434 2382 1141 • JUNTCF7 2362 96 311 1993 516 • JUNWNT2B 2380 158 211 829 765 • JUNTCF7L1 2384 1691 1186 2021 1413 • JUNRHOU 2403 670 2171 1447 224 • JUNSENP2 2484 1385 2300 1590 364

## JUN 2*^nd^* order interaction ranking at different durations

• JUNSENP2 24 2466 2351 157 • JUNSFRP4 54 2392 1043 5 • JUNSFRP1 113 1873 1613 83 • JUNLRP6 127 1970 2214 1322 • JUNLEF1 131 1875 1950 104 • JUNPITX2 173 1891 1135 1094 • JUNWNT5A 194 1515 1716 38 • JUNTCF7L1 240 1863 2085 2230 • JUNPPP2R1A 252 2251 1749 88 • JUNTLE2 322 948 2299 198 • JUNWNT4 360 1840 1467 499 • JUNNLK 392 2370 309 445 • JUNFBXW4 398 812 2338 140 • JUNPPP2CA 506 1282 2056 421 • JUNTCF7 578 916 434 708 • JUNLRP5 598 1053 1817 625 • JUNTLE1 775 1669 1931 622 • JUNNKD1 786 1361 451 138 • JUNWNT2 806 1513 1721 2168 • JUNSLC9A3R1 875 681 837 301 • JUNT 909 1717 2410 1229 • JUNWNT1 933 2473 471 1197 • JUNWIF1 937 2168 2368 37 • CSNK1A1JUN 959 1599 2466 1950 • FZD5JUN 1007 582 2426 770 • DIXDC1JUN 1027 126 388 839 • AXIN1JUN 1128 493 884 134 • JUNKREMEN1 1134 2092 1684 1083 • JUNMYC 1172 843 532 1017 • JUNWNT2B 1239 2419 1453 173 • CTNNBIP1JUN 1271 1837 459 640 • CTBP1JUN 1327 2302 2385 973 • BCL9JUN 1335 2207 1555 1025 • GSK3AJUN 1340 305 431 467 • CCND2JUN 1369 2112 1854 998 • BTRCJUN 1370 1022 1165 641 • JUNWNT3A 1383 1151 1532 436 • AESJUN 1422 2439 968 1401 • FOXN1JUN 1476 1653 2219 2176 • CTNNB1JUN 1511 1387 913 1531 • CSNK1DJUN 1522 271 2419 2328 • CCND1JUN 1529 622 275 1641 • FOSL1JUN 1674 206 2221 910 • FRAT1JUN 1677 915 1354 1048 • FSHBJUN 1734 348 2290 407 • APCJUN 1761 1308 1870 1567 • JUNWNT3 1784 898 2061 278 • JUNPORCN 1794 2381 2372 108 • FRZBJUN 1838 2162 1768 1482 • JUNRHOU 1882 1949 2233 2478 • CXXC4JUN 1990 1940 2039 312 • DKK1JUN 2032 270 1119 1858 • EP300JUN 2077 1959 1715 186 • DVL2JUN 2114 738 496 859 • DVL1JUN 2122 2485 290 196 • GSK3BJUN 2155 135 80 2298 • FZD7JUN 2164 163 44 373 • FZD1JUN 2169 195 806 679 • FGF4JUN 2173 325 1306 1234 • CTBP2JUN 2194 1401 1889 876 • DAAM1JUN 2198 1167 526 232 • JUNPYGO1 2206 2222 2134 466 • FBXW11JUN 2217 527 1775 1281 • FBXW2JUN 2306 1134 2046 314 • FZD2JUN 2308 697 285 1347 • FZD6JUN 2321 648 161 307 • CCND3JUN 2355 825 553 423 • FZD8JUN 2365 72 334 1773 • CSNK2A1JUN 2371 651 332 911 • CSNK1G1JUN 2408 304 1459 2433

## KREMEN1 2*^nd^* order interaction ranking at

• FRZBKREMEN1 139 1894 471 1101 623 • DKK1KREMEN1 180 2154 1815 31 2398 • KREMEN1WNT1 228 2123 242 2436 824 • KREMEN1PYGO1 318 1547 1057 259 1123 • KREMEN1SFRP1 339 1791 2291 1948 878 • KREMEN1WIF1 358 1938 1317 532 939 • KREMEN1PORCN 397 774 354 2395 840 • KREMEN1LEF1 402 686 544 1970 779 • KREMEN1MYC 431 1517 631 2188 1111 • KREMEN1WNT3 545 2051 2108 158 219 • FOSL1KREMEN1 587 1541 710 429 30 • APCKREMEN1 604 1973 986 1247 357 • FZD7KREMEN1 612 792 578 2166 1587 • KREMEN1T 628 1932 1056 1736 2056 • FBXW11KREMEN1 647 343 1611 2008 2172 • CSNK2A1KREMEN1 666 683 2023 1467 1639 • KREMEN1LRP6 676 490 557 1100 528 • FRAT1KREMEN1 681 2395 1928 170 554 • KREMEN1NKD1 715 1969 656 817 852 • FZD1KREMEN1 738 586 1740 1630 474 • FZD5KREMEN1 773 530 513 2347 79 • DIXDC1KREMEN1 823 82 56 1114 2335 • CXXC4KREMEN1 826 838 891 490 1903 • FZD8KREMEN1 860 14 420 2259 1965 • KREMEN1FBXW4 907 1905 333 340 881 • CSNK1DKREMEN1 952 1723 1481 2099 296 • KREMEN1TLE1 957 1480 1158 855 387 • GSK3BKREMEN1 998 161 2138 489 330 • EP300KREMEN1 1006 1853 1521 281 639 • CCND3KREMEN1 1073 382 1142 2396 2003 • KREMEN1PITX2 1074 441 542 1668 320 • FBXW2KREMEN1 1083 542 1986 1452 2414 • FSHBKREMEN1 1122 279 1004 2296 365 • KREMEN1SFRP4 1179 956 1913 180 2039 • KREMEN1PPP2CA 1201 1634 423 1797 2471 • KREMEN1PPP2R1A 1209 318 658 2293 1386 • KREMEN1WNT2B 1216 468 82 1553 1925 • CTBP1KREMEN1 1221 1835 1127 1687 760 • KREMEN1TCF7 1238 1608 146 1187 2194 • DAAM1KREMEN1 1253 299 1433 1525 1748 • KREMEN1SLC9A3R1 1255 271 1668 258 1179 • KREMEN1NLK 1278 1774 461 1047 2018 • KREMEN1WNT2 1284 78 871 1146 1347 • KREMEN1WNT4 1299 556 1637 636 841 • FZD6KREMEN1 1305 15 1729 1520 1281 • KREMEN1WNT3A 1404 9 1367 2407 1478 • KREMEN1TLE2 1414 787 1165 335 1052 • KREMEN1LRP5 1444 1575 1243 2077 1282 • KREMEN1RHOU 1452 606 1888 38 2286 • KREMEN1WNT5A 1555 1961 726 2426 963 • DVL2KREMEN1 1601 2002 1968 1697 1694 • KREMEN1TCF7L1 1609 377 1006 1658 2251 • CTNNB1KREMEN1 1634 1397 914 460 1163 • CTNNBIP1KREMEN1 1741 851 971 991 125 • DVL1KREMEN1 1788 171 989 496 109 • CTBP2KREMEN1 1794 1627 1046 2114 1582 • CSNK1G1KREMEN1 1824 1978 1045 1457 1181 • BTRCKREMEN1 1841 183 973 2040 1149 • AXIN1KREMEN1 1842 374 2232 862 719 • CCND2KREMEN1 1884 943 717 1631 2370 • BCL9KREMEN1 1954 364 828 2037 996 • FOXN1KREMEN1 1957 1505 2414 2056 1086 • AESKREMEN1 2032 1514 690 927 526 • FZD2KREMEN1 2082 736 1122 1373 604 • JUNKREMEN1 2199 1452 1038 1579 104 • CSNK1A1KREMEN1 2227 17 622 320 238 • FGF4KREMEN1 2241 1240 563 1828 1538 • **different durations** • CCND1KREMEN1 2360 893 1206 1945 1521 • GSK3AKREMEN1 2399 1248 1868 1771 1025 • KREMEN1SENP2 2448 1039 1502 355 1364

## KREMEN1 2*^nd^* order interaction ranking at different durations

• KREMEN1SENP2 51 2011 249 189 • KREMEN1SFRP4 84 1106 376 238 • CSNK1A1KREMEN1 133 1876 867 1730 • KREMEN1LEF1 168 2279 831 1144 • KREMEN1LRP6 270 2242 731 1327 • KREMEN1WNT5A 333 974 861 2480 • KREMEN1TCF7L1 397 1321 306 1898 • KREMEN1SFRP1 423 2308 854 1492 • KREMEN1PPP2R1A 445 726 475 1625 • KREMEN1PITX2 452 1606 1171 2131 • KREMEN1PPP2CA 497 714 549 2445 • KREMEN1NLK 526 856 55 1618 • KREMEN1TLE1 635 794 245 558 • KREMEN1NKD1 638 613 119 465 • KREMEN1TLE2 737 1823 897 1269 • KREMEN1TCF7 742 1064 65 2424 • KREMEN1FBXW4 800 1130 1417 353 • CTNNB1KREMEN1 805 1213 2343 1534 • FZD5KREMEN1 816 400 1793 1629 • KREMEN1LRP5 857 1608 886 1416 • KREMEN1WNT4 865 925 771 2238 • DIXDC1KREMEN1 1002 51 1743 523 • KREMEN1SLC9A3R1 1003 1803 2196 365 • AESKREMEN1 1075 2030 1216 997 • CSNK1DKREMEN1 1082 472 1561 2075 • KREMEN1WNT2 1131 2236 2139 2001 • JUNKREMEN1 1134 2092 1684 1083 • CTBP1KREMEN1 1150 2058 1177 2448 • KREMEN1WIF1 1179 2437 1042 97 • AXIN1KREMEN1 1201 507 1248 1196 • KREMEN1WNT1 1220 1014 436 2169 • KREMEN1MYC 1279 1007 270 1358 • CCND2KREMEN1 1349 2188 895 1855 • KREMEN1T 1358 2301 322 2332 • FRAT1KREMEN1 1365 597 1574 2326 • CCND1KREMEN1 1378 2027 2366 2247 • BTRCKREMEN1 1430 1084 2118 2281 • CTNNBIP1KREMEN1 1439 2346 1526 826 • BCL9KREMEN1 1524 1691 1330 1663 • DVL1KREMEN1 1625 2313 1370 694 • APCKREMEN1 1633 2163 1095 804 • FSHBKREMEN1 1656 218 1188 1101 • GSK3AKREMEN1 1665 478 1159 2034 • FOSL1KREMEN1 1697 209 1626 763 • KREMEN1WNT3A 1726 1378 1777 2072 • KREMEN1WNT3 1774 2380 159 359 • CTBP2KREMEN1 1836 722 921 2080 • KREMEN1RHOU 1852 1538 1336 1228 • DKK1KREMEN1 1858 161 1455 1027 • FOXN1KREMEN1 1905 1324 2031 1901 • EP300KREMEN1 1946 816 1527 788 • KREMEN1WNT2B 1948 1583 9 181 • FRZBKREMEN1 1979 2274 1619 1790 • FZD6KREMEN1 2005 821 821 665 • DVL2KREMEN1 2011 987 2158 634 • CCND3KREMEN1 2035 2281 1915 2276 • FZD7KREMEN1 2087 113 226 743 • GSK3BKREMEN1 2135 235 1116 2381 • FZD1KREMEN1 2176 139 1959 2097 • FBXW2KREMEN1 2248 1975 615 1983 • KREMEN1PORCN 2300 1846 501 1861 • CSNK1G1KREMEN1 2304 297 1548 2059 • CSNK2A1KREMEN1 2305 988 683 1787 • DAAM1KREMEN1 2310 1526 2053 1172 • FZD8KREMEN1 2323 40 1914 2126 • KREMEN1PYGO1 2334 1813 1635 2043 • CXXC4KREMEN1 2378 2354 879 1615 • FZD2KREMEN1 2416 1248 997 1944 • FBXW11KREMEN1 2425 920 1433 2436 • FGF4KREMEN1 2459 342 833 2120

## LEF1 2*^nd^* order interaction ranking at different time points

• FRZBLEF1 12 629 987 1090 1958 • DKK1LEF1 33 2318 2474 876 2307 • DIXDC1LEF1 122 251 145 1891 2482 • FZD1LEF1 142 1598 1219 544 2103 • APCLEF1 157 470 1387 387 374 • FBXW11LEF1 160 568 1907 1517 2072 • FOSL1LEF1 163 439 342 1661 1204 • FZD8LEF1 167 120 718 1679 1343 • FZD7LEF1 212 1024 1136 3 1355 • CSNK2A1LEF1 214 50 2456 2435 1601 • FZD5LEF1 229 90 1331 1966 310 • CXXC4LEF1 255 1040 1315 568 1379 • GSK3BLEF1 293 52 1003 2222 1566 • EP300LEF1 333 1662 1525 796 723 • CTBP1LEF1 346 1389 725 273 464 • CTNNB1LEF1 366 912 1713 2014 1252 • FRAT1LEF1 369 2286 1415 470 2113 • FBXW2LEF1 398 681 2288 1974 2456 • KREMEN1LEF1 402 686 544 1970 779 • DAAM1LEF1 435 1012 832 820 1686 • FZD6LEF1 452 348 2470 482 1240 • CSNK1DLEF1 474 700 1358 1068 2153 • AXIN1LEF1 497 1330 1519 853 2262 • FSHBLEF1 508 616 1870 36 1041 • CCND3LEF1 514 713 2295 722 2402 • CTNNBIP1LEF1 585 481 850 1154 142 • DVL2LEF1 616 283 2132 895 1110 • CTBP2LEF1 739 447 388 1183 2049 • CSNK1G1LEF1 744 368 2441 2027 2395 • LEF1PYGO1 873 383 565 1527 2132 • FOXN1LEF1 885 872 2228 118 624 • LEF1MYC 894 1410 819 1174 1943 • DVL1LEF1 906 398 752 941 261 • LEF1WNT3 1003 1605 1381 2349 560 • AESLEF1 1072 559 915 471 1743 • LEF1WIF1 1103 2057 1524 1042 2237 • LEF1SFRP1 1142 2459 2269 2006 813 • BTRCLEF1 1151 371 2416 1070 441 • CCND2LEF1 1265 896 1282 2018 2476 • LEF1PORCN 1339 584 1548 344 1545 • LEF1LRP6 1386 1053 1040 1564 107 • FGF4LEF1 1397 906 411 1250 2264 • BCL9LEF1 1398 2308 2327 13 1981 • LEF1WNT1 1453 2312 553 1004 1293 • CSNK1A1LEF1 1498 1110 1850 1487 775 • LEF1T 1533 875 978 898 1505 • JUNLEF1 1563 508 2110 639 542 • GSK3ALEF1 1567 784 1281 637 2242 • CCND1LEF1 1695 528 1116 2333 761 • FZD2LEF1 1697 495 570 1078 823 • LEF1FBXW4 1700 1559 773 2157 1543 • LEF1NKD1 1881 2007 1283 1869 1501 • LEF1SLC9A3R1 1893 589 2485 2332 699 • LEF1WNT2 1904 242 1428 515 661 • LEF1PPP2CA 1953 1857 400 797 2090 • LEF1SFRP4 1973 488 2093 62 2081 • LEF1TLE1 2009 1169 1991 499 1513 • LEF1WNT4 2049 2033 1587 351 1194 • LEF1WNT2B 2074 982 47 917 1556 • LEF1PPP2R1A 2124 644 1355 2182 466 • LEF1TCF7 2155 1150 503 2459 1258 • LEF1PITX2 2169 702 1582 1470 1023 • LEF1WNT5A 2171 1122 1029 1552 1490 • LEF1LRP5 2190 457 1311 2428 1598 • LEF1WNT3A 2216 326 2238 1714 1103 • LEF1NLK 2244 1476 273 1018 2447 • LEF1TLE2 2272 1113 2219 606 932 • LEF1TCF7L1 2306 1102 1559 1237 336 • LEF1RHOU 2378 1632 1077 1117 1970 • LEF1SENP2 2479 2006 2172 188 1682

## LEF1 2*^nd^* order interaction ranking at different durations

• FZD5LEF1 13 1838 758 937 • CSNK1A1LEF1 18 1554 922 78 • CTNNB1LEF1 122 1051 2360 1755 • JUNLEF1 131 1875 1950 104 • DIXDC1LEF1 137 627 757 1349 • FRAT1LEF1 155 2334 1507 291 • KREMEN1LEF1 168 2279 831 1144 • CTBP1LEF1 169 1659 1592 518 • CCND1LEF1 174 1206 627 1769 • CCND2LEF1 181 1741 1886 2376 • CSNK1DLEF1 183 1241 1782 1242 • AESLEF1 203 2245 1697 331 • FSHBLEF1 219 1444 1318 1486 • BCL9LEF1 224 1224 1136 1899 • BTRCLEF1 313 2025 1991 275 • AXIN1LEF1 317 2205 1021 2253 • CTNNBIP1LEF1 361 2223 1789 1398 • FOXN1LEF1 386 1433 1362 379 • APCLEF1 428 1381 1542 1758 • CTBP2LEF1 440 1906 2104 2082 • GSK3ALEF1 496 2065 598 1728 • FOSL1LEF1 516 946 2189 1674 • DVL1LEF1 525 1727 1766 666 • EP300LEF1 686 1692 2436 1912 • FRZBLEF1 707 2051 1588 292 • DVL2LEF1 709 1195 1764 1997 • LEF1SENP2 738 531 223 849 • DKK1LEF1 866 2264 2151 1776 • CCND3LEF1 890 2173 1733 1157 • CXXC4LEF1 891 1688 2086 2327 • DAAM1LEF1 895 1860 1142 2041 • FZD6LEF1 1022 2293 474 828 • FGF4LEF1 1050 1034 2180 1533 • FZD8LEF1 1058 979 170 1720 • GSK3BLEF1 1077 1203 676 604 • FBXW2LEF1 1112 1499 2297 1519 • CSNK2A1LEF1 1158 2275 800 991 • FZD7LEF1 1190 2179 218 1175 • LEF1SFRP4 1194 392 672 536 • FZD1LEF1 1229 717 2113 594 • FBXW11LEF1 1242 2193 1195 479 • FZD2LEF1 1374 980 1198 1799 • LEF1LRP6 1447 686 874 2008 • CSNK1G1LEF1 1480 1973 1465 2428 • LEF1PPP2R1A 1535 216 1308 1821 • LEF1NLK 1558 180 182 2418 • LEF1SFRP1 1560 1015 1939 964 • LEF1TCF7L1 1617 1010 1423 2114 • LEF1WNT5A 1641 191 1058 2236 • LEF1PPP2CA 1782 240 1117 750 • LEF1PITX2 1806 2085 1943 1433 • LEF1TLE2 1849 249 1175 80 • LEF1WIF1 1937 335 1663 1783 • LEF1LRP5 1969 1503 2227 744 • LEF1SLC9A3R1 1995 518 2482 1368 • LEF1WNT4 1998 162 2293 945 • LEF1NKD1 2039 101 104 1406 • LEF1TCF7 2052 192 27 1037 • LEF1FBXW4 2112 128 2235 1180 • LEF1WNT2 2138 374 2350 1210 • LEF1RHOU 2146 590 2321 983 • LEF1WNT1 2192 107 2238 490 • LEF1WNT3 2250 873 140 2240 • LEF1WNT3A 2273 712 2168 505 • LEF1TLE1 2283 710 1589 1689 • LEF1MYC 2286 187 209 1733 • LEF1WNT2B 2345 363 85 582 • LEF1PYGO1 2409 217 1279 1343 • LEF1T 2429 198 642 277 • LEF1PORCN 2479 158 878 1778

## MYC 2*^nd^* order interaction ranking at different time points

• FRZBMYC 23 2341 320 206 501 • DKK1MYC 37 2349 1497 196 2337 • FZD7MYC 84 1160 43 2336 1274 • FZD1MYC 112 1291 756 266 2412 • APCMYC 116 2248 619 1855 2105 • GSK3BMYC 190 752 429 1812 2427 • FBXW11MYC 192 979 687 684 1419 • LRP5MYC 193 2244 516 1739 1889 • CSNK2A1MYC 222 1837 1012 2176 2384 • FOSL1MYC 227 2171 40 776 1746 • CSNK1DMYC 250 2298 1695 1460 2125 • CXXC4MYC 261 2077 181 1102 2057 • DIXDC1MYC 272 2276 21 2461 1712 • FZD5MYC 274 2249 398 1323 2160 • EP300MYC 294 592 668 734 2126 • FZD6MYC 319 1119 2250 1669 1280 • CCND3MYC 323 1673 1154 1534 1580 • FZD8MYC 344 381 587 1297 2253 • FRAT1MYC 364 2081 714 1707 2334 • FBXW2MYC 381 1630 1319 641 1292 • CTBP2MYC 401 352 433 1765 2024 • CTBP1MYC 411 2011 580 683 1227 • FSHBMYC 413 2306 996 1170 2023 • CTNNB1MYC 425 2028 576 884 2422 • KREMEN1MYC 431 1517 631 2188 1111 • DAAM1MYC 472 1530 1324 1313 732 • AXIN1MYC 584 2418 696 1253 700 • CSNK1G1MYC 588 2186 782 1172 2293 • DVL2MYC 622 625 739 2421 2302 • MYCWNT1 687 2365 1041 1603 96 • CTNNBIP1MYC 729 1744 597 1905 1432 • LRP6MYC 821 750 1055 2400 2432 • FGF4MYC 863 1271 245 241 1034 • LEF1MYC 894 1410 819 1174 1943 • BTRCMYC 901 1125 1250 963 2306 • MYCPYGO1 919 1954 931 799 15 • AESMYC 955 1847 866 789 2021 • MYCSFRP1 983 337 2374 2270 8 • JUNMYC 1014 206 489 655 1946 • CCND2MYC 1037 1196 536 2283 1238 • DVL1MYC 1094 199 380 933 1986 • MYCPORCN 1095 1084 1409 2358 258 • MYCWNT3 1149 660 1692 1973 26 • CSNK1A1MYC 1163 1566 634 1569 177 • BCL9MYC 1226 2114 284 1799 893 • FZD2MYC 1258 1629 344 1180 1530 • GSK3AMYC 1464 2169 1538 745 2288 • MYCWIF1 1504 2321 1883 977 1349 • CCND1MYC 1593 1459 823 2312 1218 • FOXN1MYC 1595 437 1254 1303 1100 • MYCT 1608 1185 1617 552 919 • MYCTLE1 1827 1550 2370 915 916 • MYCTCF7 1917 1611 713 672 70 • MYCNKD1 1936 839 956 1418 900 • MYCSFRP4 1966 1960 1676 122 1206 • MYCPPP2R1A 2057 1855 1714 1539 97 • MYCPITX2 2067 1977 2060 2356 476 • MYCFBXW4 2071 1863 799 1227 860 • MYCNLK 2139 1805 1732 1965 59 • MYCTLE2 2143 1572 1809 1264 1488 • MYCWNT5A 2156 1327 1547 1639 606 • MYCSLC9A3R1 2163 94 1962 625 725 • MYCWNT2 2170 1508 1861 530 1595 • MYCWNT2B 2179 246 1240 1232 541 • MYCPPP2CA 2225 563 499 688 1706 • MYCWNT4 2232 1307 1459 2139 825 • MYCRHOU 2263 499 1730 1050 667 • MYCTCF7L1 2269 2204 2429 2158 2092 • MYCWNT3A 2271 942 858 1820 196 • MYCSENP2 2480 1021 2246 366 302

## MYC 2*^nd^* order interaction ranking at different durations

• MYCSENP2 86 1352 1892 1280 • MYCSFRP4 117 1132 2120 568 • MYCPITX2 410 1058 916 1693 • MYCSFRP1 424 1986 1798 2190 • MYCTCF7L1 480 1678 2094 2166 • MYCPPP2R1A 493 1587 1536 1966 • MYCWNT5A 512 1161 2412 1803 • MYCPPP2CA 571 862 2040 1218 • MYCNLK 646 932 486 2397 • CSNK1A1MYC 649 942 775 355 • FZD5MYC 663 1173 1799 2341 • MYCTLE2 719 539 2267 326 • MYCWNT4 724 747 1019 1836 • CTNNB1MYC 825 1712 423 1237 • MYCFBXW4 828 633 1896 976 • MYCTCF7 877 1694 277 1834 • MYCTLE1 898 1792 1428 803 • CTBP1MYC 980 2321 2387 1558 • DIXDC1MYC 994 27 180 709 • CCND2MYC 1051 1983 2026 2384 • MYCNKD1 1054 360 1022 1883 • MYCWNT2 1069 1185 1231 707 • AESMYC 1071 819 406 484 • CCND1MYC 1074 661 295 1870 • MYCWIF1 1136 2296 2245 547 • JUNMYC 1172 843 532 1017 • CTNNBIP1MYC 1181 2409 379 1927 • FOXN1MYC 1207 787 1297 882 • FRAT1MYC 1268 1072 790 1656 • KREMEN1MYC 1279 1007 270 1358 • MYCWNT1 1289 858 1769 1193 • DVL1MYC 1321 789 241 943 • CSNK1DMYC 1333 509 1450 768 • BTRCMYC 1361 1101 1025 2183 • BCL9MYC 1414 2320 1264 1296 • AXIN1MYC 1421 880 433 2212 • FSHBMYC 1435 280 319 1302 • MYCSLC9A3R1 1525 839 1794 2446 • FOSL1MYC 1569 369 1719 1759 • MYCWNT3A 1588 994 658 2022 • MYCT 1660 510 1992 218 • GSK3AMYC 1692 330 22 831 • LRP5MYC 1738 174 185 60 • FRZBMYC 1744 1520 1502 265 • APCMYC 1766 2369 830 2290 • CTBP2MYC 1848 941 2037 1692 • MYCWNT2B 1863 1145 1092 348 • FGF4MYC 1867 109 256 1365 • FBXW11MYC 1933 1925 1831 1311 • MYCPORCN 1935 1434 2449 1373 • DKK1MYC 1940 263 835 2224 • EP300MYC 1944 1908 311 434 • DVL2MYC 1965 1121 248 2406 • DAAM1MYC 1988 977 152 1991 • MYCWNT3 2004 1356 328 201 • CCND3MYC 2025 1643 696 1095 • CXXC4MYC 2055 1617 1568 809 • MYCRHOU 2062 693 900 450 • FZD2MYC 2095 504 90 1849 • FBXW2MYC 2099 1764 476 2322 • FZD7MYC 2134 193 10 1455 • FZD8MYC 2187 52 42 917 • MYCPYGO1 2209 735 2319 2377 • LRP6MYC 2228 409 1277 1706 • FZD6MYC 2246 1060 56 580 • CSNK2A1MYC 2282 1995 115 2438 • LEF1MYC 2286 187 209 1733 • FZD1MYC 2366 67 906 1366 • GSK3BMYC 2382 70 53 403 • CSNK1G1MYC 2473 370 484 638

## NKD1 2*^nd^* order interaction ranking at different time points

• FRZBNKD1 48 2430 159 1677 1491 • DKK1NKD1 79 2381 1377 1159 2308 • FZD1NKD1 306 392 291 658 2038 • NKD1WNT1 338 7 260 1267 1212 • FZD7NKD1 388 948 169 589 2320 • NKD1PYGO1 393 705 1346 39 618 • FZD5NKD1 432 1438 618 1324 745 • FRAT1NKD1 433 2180 232 2351 2107 • LRP5NKD1 434 2307 981 878 1433 • APCNKD1 454 1839 379 1824 1283 • CSNK2A1NKD1 478 1232 1437 2271 1464 • FOSL1NKD1 495 2311 330 1072 1097 • DIXDC1NKD1 522 1380 20 478 2284 • NKD1SFRP1 535 962 2184 1355 314 • FBXW11NKD1 568 721 1583 999 2423 • GSK3BNKD1 578 753 463 226 1494 • NKD1WNT3 591 53 2430 1271 288 • CSNK1DNKD1 596 2438 640 2215 1868 • FZD8NKD1 649 1827 326 1574 2416 • NKD1WIF1 650 611 1688 2151 1075 • CCND3NKD1 695 836 1294 327 1557 • FBXW2NKD1 714 856 1397 2319 335 • KREMEN1NKD1 715 1969 656 817 852 • NKD1PORCN 749 652 1858 455 388 • CXXC4NKD1 770 2484 590 1198 827 • FSHBNKD1 771 2096 2259 793 1924 • FZD6NKD1 883 1727 1234 586 1554 • DAAM1NKD1 889 1845 916 202 2339 • EP300NKD1 912 1856 529 903 865 • CTBP1NKD1 938 2166 487 1839 128 • NKD1TLE1 963 42 2306 80 1215 • NKD1FBXW4 1016 541 502 1955 1381 • CTNNBIP1NKD1 1053 2442 279 451 229 • CTNNB1NKD1 1081 1158 609 134 2117 • AXIN1NKD1 1186 2091 875 1757 1997 • DVL2NKD1 1251 1717 174 529 2232 • NKD1T 1269 385 1198 964 1656 • AESNKD1 1279 146 1696 725 1774 • CSNK1G1NKD1 1353 2337 607 2136 2358 • CTBP2NKD1 1359 2213 246 1547 1342 • LRP6NKD1 1409 2473 486 1270 2118 • NKD1WNT2 1502 646 1639 297 2133 • NKD1NLK 1538 565 1042 1408 2186 • NKD1SFRP4 1574 2420 1652 638 2238 • NKD1PITX2 1579 80 1082 1066 1360 • JUNNKD1 1615 1250 428 695 410 • CSNK1A1NKD1 1617 1037 707 20 905 • DVL1NKD1 1624 693 382 582 690 • NKD1PPP2CA 1651 160 249 1178 2450 • NKD1SLC9A3R1 1664 1912 2126 661 1907 • NKD1WNT2B 1672 62 1363 2348 1833 • CCND2NKD1 1706 2469 754 1175 2114 • NKD1RHOU 1749 525 2089 462 2212 • FOXN1NKD1 1755 293 1018 502 1662 • NKD1WNT3A 1778 2297 1972 402 886 • NKD1WNT4 1790 1674 1886 1065 2104 • NKD1TLE2 1798 1615 1189 1112 2474 • NKD1PPP2R1A 1805 609 661 1681 87 • NKD1TCF7 1813 165 654 1407 2043 • NKD1WNT5A 1817 91 917 2134 1836 • CCND1NKD1 1864 1316 722 649 888 • LEF1NKD1 1881 2007 1283 1869 1501 • BTRCNKD1 1887 174 1995 795 1323 • MYCNKD1 1936 839 956 1418 900 • FZD2NKD1 1950 1828 466 1613 735 • FGF4NKD1 1996 2320 104 703 2321 • BCL9NKD1 2003 2247 711 1555 2396 • NKD1TCF7L1 2096 1068 1257 2346 2127 • GSK3ANKD1 2196 2045 766 1834 1874 • NKD1SENP2 2456 132 1666 1870 1354

## NKD1 2*^nd^* order interaction ranking at different durations

• NKD1SENP2 159 1517 1737 1798 • FZD5NKD1 189 184 687 2270 • CSNK1A1NKD1 232 1630 713 614 • NKD1SFRP4 310 1209 1781 623 • CTBP1NKD1 614 868 1804 2419 • KREMEN1NKD1 638 613 119 465 • DIXDC1NKD1 681 11 47 1547 • CTNNB1NKD1 699 1143 547 1588 • CCND2NKD1 704 1023 1456 1830 • NKD1PPP2R1A 714 1210 2247 894 • NKD1WNT5A 743 1956 1516 2413 • BCL9NKD1 751 705 500 2254 • NKD1SFRP1 752 1200 1546 1007 • JUNNKD1 786 1361 451 138 • NKD1TCF7L1 792 1890 1826 540 • NKD1PITX2 846 1333 594 2374 • CSNK1DNKD1 854 339 1656 1419 • NKD1PPP2CA 868 1795 1696 2267 • FRAT1NKD1 930 311 584 135 • NKD1NLK 970 2457 114 1054 • NKD1WNT4 997 2444 1011 1173 • BTRCNKD1 1016 636 1061 1000 • MYCNKD1 1054 360 1022 1883 • CCND1NKD1 1059 368 184 649 • AESNKD1 1066 1370 149 676 • AXIN1NKD1 1076 308 99 1423 • DVL1NKD1 1103 861 86 755 • FSHBNKD1 1104 170 359 1510 • CTNNBIP1NKD1 1116 1275 173 700 • APCNKD1 1147 1784 1355 1815 • NKD1TCF7 1152 1171 347 2320 • NKD1TLE2 1174 2109 1720 1111 • FOXN1NKD1 1185 741 407 345 • GSK3ANKD1 1259 76 68 1377 • CTBP2NKD1 1301 971 566 1801 • NKD1WIF1 1397 1264 1665 223 • NKD1FBXW4 1412 1419 2207 1445 • NKD1TLE1 1436 1062 2220 1672 • NKD1SLC9A3R1 1442 1277 415 1494 • DVL2NKD1 1457 901 232 2025 • FRZBNKD1 1464 612 2077 251 • NKD1WNT2 1466 1883 2047 813 • EP300NKD1 1505 2067 1111 1652 • FOSL1NKD1 1512 108 477 2160 • CCND3NKD1 1516 961 268 2339 • CXXC4NKD1 1554 1802 1385 1879 • FBXW11NKD1 1608 706 1351 2143 • NKD1T 1687 2291 1591 489 • LRP5NKD1 1730 34 244 212 • NKD1WNT1 1789 1745 1218 1188 • GSK3BNKD1 1818 10 131 765 • NKD1WNT2B 1842 2410 344 513 • DKK1NKD1 1857 49 2248 1655 • FBXW2NKD1 1874 429 192 2274 • FZD1NKD1 1902 66 331 99 • CSNK2A1NKD1 1936 450 96 2316 • FZD7NKD1 1956 16 6 1152 • DAAM1NKD1 1957 406 212 1115 • FZD8NKD1 1975 5 62 2028 • CSNK1G1NKD1 2022 110 540 922 • LEF1NKD1 2039 101 104 1406 • NKD1PORCN 2070 2477 2406 2400 • NKD1RHOU 2083 1181 1404 395 • FZD2NKD1 2096 444 144 1548 • NKD1WNT3A 2103 1612 1039 1315 • FZD6NKD1 2163 411 121 1319 • FGF4NKD1 2252 59 1155 1658 • NKD1PYGO1 2298 2406 1967 593 • LRP6NKD1 2357 29 2140 2369 • NKD1WNT3 2474 1329 2101 656

## NLK 2*^nd^* order interaction ranking at different time points

• NLKPYGO1 69 1980 1391 1064 2220 • NLKWNT1 115 485 1014 560 431 • DKK1NLK 118 2401 1436 1231 1918 • FRZBNLK 203 2317 140 1844 187 • NLKSFRP1 238 1695 2056 680 1011 • NLKWIF1 277 968 2007 343 2446 • NLKPORCN 384 400 2483 821 712 • NLKWNT3 387 107 1795 380 158 • NLKT 479 2119 2267 846 711 • FZD1NLK 594 1311 269 691 290 • FBXW11NLK 659 182 1790 881 174 • NLKTLE1 677 1109 1929 651 479 • APCNLK 710 2319 95 643 256 • NLKPITX2 828 1116 2156 1410 1847 • FOSL1NLK 833 1486 16 935 280 • CSNK2A1NLK 848 668 1601 1918 834 • FZD7NLK 867 2314 50 1333 805 • FRAT1NLK 900 2020 366 1512 517 • LRP5NLK 904 2464 530 1258 565 • NLKSFRP4 916 2109 1948 1498 1764 • NLKFBXW4 959 139 2167 1709 1870 • NLKPPP2R1A 964 1051 2329 2067 570 • GSK3BNLK 974 297 637 1190 2042 • CSNK1DNLK 1018 2142 921 1903 730 • CXXC4NLK 1035 2141 179 251 1971 • NLKSLC9A3R1 1039 633 1952 384 1863 • DIXDC1NLK 1051 2197 2 838 371 • NLKTCF7 1085 732 1653 1379 937 • FZD5NLK 1091 1600 59 670 486 • FZD8NLK 1109 672 351 1612 1890 • NLKWNT2 1117 134 2272 921 2026 • NLKPPP2CA 1124 210 1518 1976 1653 • FZD6NLK 1133 768 2307 1455 1161 • EP300NLK 1148 1684 432 982 507 • NLKWNT3A 1189 1579 2341 926 1298 • FBXW2NLK 1210 533 541 2016 1407 • CTBP1NLK 1227 2323 100 1055 499 • CCND3NLK 1250 1613 1808 2301 1401 • KREMEN1NLK 1278 1774 461 1047 2018 • NLKWNT4 1310 1356 2223 2367 152 • CTNNB1NLK 1417 1715 168 2102 935 • FSHBNLK 1430 2219 2361 1868 375 • DAAM1NLK 1457 35 720 2005 1229 • NLKTLE2 1489 1752 2309 2199 2083 • NLKWNT2B 1491 954 175 1376 1707 • NLKWNT5A 1511 1685 1318 1652 1976 • NKD1NLK 1538 565 1042 1408 2186 • CSNK1G1NLK 1589 2174 2209 116 1196 • CTNNBIP1NLK 1620 2287 476 2210 81 • NLKRHOU 1661 350 1174 129 2152 • DVL2NLK 1703 1094 1871 1334 650 • NLKTCF7L1 1746 59 1592 494 1267 • CTBP2NLK 1751 387 518 608 345 • DVL1NLK 1761 728 126 570 980 • CSNK1A1NLK 1779 2449 532 1171 117 • AXIN1NLK 1812 2363 493 254 1383 • BTRCNLK 1835 263 1407 1386 1520 • LRP6NLK 1894 1539 1768 823 1980 • CCND2NLK 1949 2136 389 1550 102 • FZD2NLK 2093 1589 198 918 72 • MYCNLK 2139 1805 1732 1965 59 • JUNNLK 2182 2046 414 2085 509 • AESNLK 2187 1778 1410 328 1974 • FGF4NLK 2188 823 19 2350 175 • BCL9NLK 2231 1971 657 431 822 • LEF1NLK 2244 1476 273 1018 2447 • FOXN1NLK 2343 291 1372 741 184 • GSK3ANLK 2372 1405 1371 765 1071 • CCND1NLK 2379 890 698 467 2218 • NLKSENP2 2442 38 1565 942 442

## NLK 2*^nd^* order interaction ranking at different durations

• FZD5NLK 68 580 1551 1916 • CSNK1A1NLK 83 1190 899 733 • DIXDC1NLK 170 15 151 1744 • CTBP1NLK 350 945 632 2454 • CTNNB1NLK 368 2367 382 2271 • JUNNLK 392 2370 309 445 • NLKSENP2 420 1550 656 1008 • BCL9NLK 427 1049 283 955 • AXIN1NLK 465 301 189 1666 • FOXN1NLK 467 665 262 1040 • CCND2NLK 488 620 1623 787 • AESNLK 501 1269 320 863 • KREMEN1NLK 526 856 55 1618 • FRAT1NLK 535 385 130 835 • CCND1NLK 560 1435 147 2179 • CSNK1DNLK 575 258 452 1458 • MYCNLK 646 932 486 2397 • FSHBNLK 652 207 1060 1427 • CTNNBIP1NLK 683 2034 84 1954 • FOSL1NLK 717 262 390 1476 • APCNLK 735 1731 1988 2484 • NLKSFRP4 770 1452 1123 629 • GSK3ANLK 783 237 19 1929 • BTRCNLK 845 1783 378 856 • DVL1NLK 873 565 46 1129 • LRP5NLK 882 45 701 263 • FRZBNLK 919 887 1881 141 • CTBP2NLK 941 926 200 841 • NKD1NLK 970 2457 114 1054 • EP300NLK 1010 2435 35 2246 • DVL2NLK 1099 1982 77 1564 • NLKSFRP1 1140 2087 1009 935 • NLKPPP2R1A 1154 653 2402 1819 • NLKTCF7L1 1175 1980 1497 1746 • CCND3NLK 1197 2101 512 1808 • DKK1NLK 1250 172 126 1738 • CXXC4NLK 1265 2350 581 303 • NLKWNT5A 1322 752 1421 2135 • FZD6NLK 1334 502 5 1465 • DAAM1NLK 1376 1093 122 1595 • NLKPITX2 1419 1212 1100 1642 • FZD7NLK 1429 81 16 2203 • FZD1NLK 1465 168 39 767 • CSNK2A1NLK 1472 1449 29 2287 • FBXW2NLK 1489 1243 312 1941 • GSK3BNLK 1506 105 30 52 • LEF1NLK 1558 180 182 2418 • NLKTLE2 1564 2478 958 376 • FZD8NLK 1582 25 7 2409 • NLKPPP2CA 1593 1246 2045 2100 • FGF4NLK 1601 298 307 2394 • NLKFBXW4 1620 1752 898 1933 • FBXW11NLK 1645 1512 1040 1133 • CSNK1G1NLK 1696 63 141 2085 • NLKTCF7 1709 1725 534 2218 • NLKTLE1 1740 1892 1041 1130 • NLKWNT4 1796 1464 858 1724 • FZD2NLK 1821 457 75 2155 • LRP6NLK 1824 243 164 2023 • NLKSLC9A3R1 2067 1595 963 1362 • NLKWNT2 2069 381 893 1677 • NLKWNT1 2073 800 2033 1511 • NLKWNT2B 2120 2098 587 399 • NLKRHOU 2131 2400 865 432 • NLKWIF1 2202 2033 1648 1177 • NLKT 2205 718 1324 739 • NLKPORCN 2226 2206 2075 2396 • NLKWNT3A 2236 2001 920 1919 • NLKWNT3 2287 2373 712 1982 • NLKPYGO1 2319 2175 1236 1792

## PITX2 2*^nd^* order interaction ranking at different time points

• FRZBPITX2 66 1794 947 1256 985 • DKK1PITX2 206 2357 1818 1099 2149 • PITX2SFRP1 276 2019 2086 61 1942 • PITX2PYGO1 280 1085 418 1731 2315 • PITX2WNT1 380 1706 86 2064 1007 • APCPITX2 490 1292 2334 1843 259 • PITX2PORCN 523 1335 574 1348 931 • PITX2WIF1 543 1595 2146 194 1906 • FZD7PITX2 581 1197 769 2320 494 • FZD1PITX2 589 1455 1998 2127 456 • FBXW11PITX2 601 630 1775 747 1803 • FOSL1PITX2 645 701 672 93 408 • PITX2WNT3 685 1224 1466 1378 1473 • FRAT1PITX2 689 1610 1673 221 613 • LRP5PITX2 725 1896 966 1719 999 • FZD5PITX2 726 1000 2099 1166 640 • PITX2T 737 1820 205 2189 2468 • CSNK1DPITX2 782 1911 992 1850 1613 • CSNK2A1PITX2 796 1154 2421 1959 115 • GSK3BPITX2 809 84 1650 1140 845 • NLKPITX2 828 1116 2156 1410 1847 • DIXDC1PITX2 845 179 196 1235 2129 • CXXC4PITX2 892 770 1369 1540 1685 • FBXW2PITX2 924 394 1742 1060 1621 • CCND3PITX2 958 422 1062 521 855 • PITX2TLE1 993 2325 1561 54 341 • FSHBPITX2 1068 344 1527 1427 1562 • KREMEN1PITX2 1074 441 542 1668 320 • CTBP1PITX2 1113 1194 1741 220 417 • PITX2SFRP4 1140 1072 2373 78 1500 • EP300PITX2 1172 339 1091 315 1736 • FZD6PITX2 1173 889 1905 1477 267 • FZD8PITX2 1206 618 1598 573 506 • DAAM1PITX2 1261 1156 1132 21 1572 • PITX2FBXW4 1280 1818 415 1204 1714 • PITX2WNT3A 1286 517 615 1340 1630 • PITX2SLC9A3R1 1368 253 1512 1158 1062 • PITX2PPP2CA 1375 1363 190 1770 2187 • PITX2WNT2 1390 1561 1819 483 742 • PITX2TCF7 1426 1956 102 868 1214 • PITX2WNT2B 1451 21 160 364 1741 • PITX2PPP2R1A 1458 1821 202 271 762 • CTNNBIP1PITX2 1460 346 1541 457 78 • PITX2WNT4 1494 407 784 89 1020 • CTNNB1PITX2 1514 964 1778 1887 332 • PITX2TLE2 1553 1322 1576 1084 773 • AXIN1PITX2 1569 2063 1594 100 652 • NKD1PITX2 1579 80 1082 1066 1360 • DVL2PITX2 1618 1268 1268 517 1716 • CSNK1G1PITX2 1631 599 1796 2088 1461 • LRP6PITX2 1633 28 1379 527 1193 • PITX2RHOU 1650 1305 1939 2002 2246 • PITX2WNT5A 1717 1347 440 293 754 • AESPITX2 1770 938 1100 1594 1174 • DVL1PITX2 1775 1680 703 169 1314 • CTBP2PITX2 1799 884 612 215 890 • CSNK1A1PITX2 1922 1721 2316 1856 189 • PITX2TCF7L1 1935 1251 873 962 1259 • FZD2PITX2 1956 1285 1427 1810 264 • CCND2PITX2 1960 1278 1204 23 2123 • FOXN1PITX2 1961 1236 1715 2160 871 • BTRCPITX2 1981 716 2247 665 1173 • JUNPITX2 2013 126 1967 738 31 • MYCPITX2 2067 1977 2060 2356 476 • CCND1PITX2 2078 149 2077 735 1369 • LEF1PITX2 2169 702 1582 1470 1023 • GSK3APITX2 2212 1865 1735 742 1466 • FGF4PITX2 2246 788 364 599 1761 • BCL9PITX2 2257 1974 2351 1983 1443 • PITX2SENP2 2447 1815 2130 162 1773

## PITX2 2*^nd^* order interaction ranking at different durations

• CSNK1A1PITX2 41 1284 1271 1602 • FZD5PITX2 95 2002 1131 1123 • CTNNB1PITX2 138 783 2301 1357 • JUNPITX2 173 1891 1135 1094 • CTBP1PITX2 234 1334 591 1287 • DIXDC1PITX2 264 1619 1899 662 • FRAT1PITX2 298 2131 1245 2263 • AESPITX2 304 1468 1126 1001 • CCND2PITX2 309 1527 1076 1178 • BCL9PITX2 321 1144 1585 1405 • CSNK1DPITX2 379 1885 1169 1840 • MYCPITX2 410 1058 916 1693 • FSHBPITX2 432 1304 1925 362 • AXIN1PITX2 439 2070 1090 1030 • KREMEN1PITX2 452 1606 1171 2131 • BTRCPITX2 490 1088 1954 1839 • CCND1PITX2 510 1529 2470 825 • FOSL1PITX2 520 1216 1761 1285 • PITX2SENP2 531 322 41 262 • FOXN1PITX2 568 1747 1226 668 • CTNNBIP1PITX2 593 1880 1016 595 • PITX2SFRP4 692 327 238 172 • DVL1PITX2 693 1674 998 758 • APCPITX2 700 1057 705 2385 • GSK3APITX2 720 2295 2155 868 • CTBP2PITX2 731 1910 1331 1292 • NKD1PITX2 846 1333 594 2374 • DVL2PITX2 884 1703 2281 1608 • EP300PITX2 894 1100 1659 217 • LRP5PITX2 911 1312 1643 628 • FRZBPITX2 922 1138 1612 942 • DAAM1PITX2 1149 944 1348 220 • PITX2PPP2R1A 1159 80 135 1367 • DKK1PITX2 1177 903 1283 2434 • CCND3PITX2 1191 1453 1505 626 • CXXC4PITX2 1223 1021 908 1211 • PITX2SFRP1 1243 438 887 779 • FZD8PITX2 1269 1080 1147 257 • PITX2WNT5A 1330 68 638 1701 • FZD6PITX2 1336 2260 759 574 • NLKPITX2 1419 1212 1100 1642 • PITX2TCF7L1 1451 605 188 729 • FZD1PITX2 1453 1340 1893 2113 • FBXW11PITX2 1469 1756 877 1859 • PITX2PPP2CA 1470 21 178 1170 • FBXW2PITX2 1482 1375 1290 1823 • GSK3BPITX2 1495 1525 559 669 • CSNK2A1PITX2 1523 1649 1674 1862 • FZD7PITX2 1538 2059 624 1421 • PITX2TLE2 1618 86 480 1004 • FZD2PITX2 1672 1397 1250 1205 • CSNK1G1PITX2 1702 1448 1560 1974 • FGF4PITX2 1732 1848 2062 716 • PITX2TLE1 1762 312 792 63 • PITX2TCF7 1779 78 20 1471 • PITX2WNT4 1792 55 1862 1854 • PITX2FBXW4 1798 8 313 692 • LEF1PITX2 1806 2085 1943 1433 • PITX2WIF1 1928 561 493 79 • LRP6PITX2 1959 2000 844 1719 • PITX2WNT2 1976 189 1208 1702 • PITX2WNT3 2015 524 58 589 • PITX2SLC9A3R1 2079 199 985 606 • PITX2T 2144 44 201 1189 • PITX2WNT3A 2225 215 1750 1554 • PITX2WNT1 2249 33 467 1382 • PITX2PORCN 2342 19 371 2137 • PITX2WNT2B 2358 343 92 44 • PITX2RHOU 2448 293 849 2195 • PITX2PYGO1 2485 144 369 2304

## PORCN 2*^nd^* order interaction ranking at different time points

• FRZBPORCN 13 1981 800 1938 85 • DKK1PORCN 63 2367 2012 1196 1898 • FZD7PORCN 83 188 673 445 170 • FOSL1PORCN 93 1875 294 311 120 • FZD1PORCN 113 415 734 2401 273 • FBXW11PORCN 178 63 1087 2181 637 • FRAT1PORCN 183 2187 924 746 250 • CSNK1DPORCN 184 2144 2399 1417 1154 • CXXC4PORCN 194 1364 1185 2303 1035 • DIXDC1PORCN 201 2201 87 1585 912 • APCPORCN 216 1139 1531 1249 245 • CSNK2A1PORCN 219 2117 1782 2180 47 • FZD5PORCN 301 1354 2004 1414 53 • FBXW2PORCN 356 1763 2387 2110 1285 • LRP5PORCN 367 1016 1851 2485 149 • DAAM1PORCN 375 494 2402 679 1671 • FZD8PORCN 379 450 464 2190 504 • NLKPORCN 384 400 2483 821 712 • KREMEN1PORCN 397 774 354 2395 840 • GSK3BPORCN 412 1010 314 1646 327 • CTNNB1PORCN 418 1811 357 792 111 • EP300PORCN 424 1625 1292 1387 424 • FZD6PORCN 448 551 1335 1252 55 • CCND3PORCN 453 1324 2418 1961 611 • CTBP1PORCN 492 1654 421 1369 62 • FSHBPORCN 515 164 1990 155 265 • PITX2PORCN 523 1335 574 1348 931 • DVL2PORCN 544 167 791 592 99 • LRP6PORCN 724 1056 1149 186 319 • PORCNWIF1 741 1950 2261 624 1700 • NKD1PORCN 749 652 1858 455 388 • CTNNBIP1PORCN 822 973 490 52 28 • CCND2PORCN 837 2407 755 302 139 • BTRCPORCN 839 591 997 1917 1251 • CTBP2PORCN 862 553 827 1008 147 • AXIN1PORCN 864 2194 1102 851 378 • CSNK1A1PORCN 962 1055 1954 1314 39 • PORCNSFRP1 977 1946 2200 724 2255 • PORCNPYGO1 990 272 700 673 566 • PORCNWNT1 1004 1747 510 924 2388 • AESPORCN 1017 835 1613 605 877 • CSNK1G1PORCN 1047 1269 1229 2233 698 • PORCNWNT3 1088 1604 2052 1315 2298 • MYCPORCN 1095 1084 1409 2358 258 • DVL1PORCN 1105 726 525 294 68 • FOXN1PORCN 1247 402 1697 925 473 • LEF1PORCN 1339 584 1548 344 1545 • BCL9PORCN 1382 1800 456 2058 1198 • FGF4PORCN 1486 1209 392 1866 399 • FZD2PORCN 1496 64 719 1449 84 • PORCNT 1509 44 945 2247 1353 • JUNPORCN 1699 1005 1148 1563 23 • PORCNFBXW4 1774 1885 1114 932 1800 • GSK3APORCN 1854 278 2481 1352 1575 • PORCNTLE1 1862 1220 1177 662 1841 • PORCNSLC9A3R1 1866 1842 2405 1353 1315 • PORCNSFRP4 1916 1883 2165 19 1661 • PORCNWNT5A 1984 2267 869 437 2316 • CCND1PORCN 1993 185 671 2124 2326 • PORCNWNT3A 2008 240 2323 914 1650 • PORCNPPP2R1A 2019 1170 1047 213 2393 • PORCNWNT2 2088 524 1813 419 1951 • PORCNPPP2CA 2103 186 275 274 897 • PORCNRHOU 2221 1872 2117 1541 1719 • PORCNTCF7 2260 98 906 2211 2312 • PORCNWNT4 2267 730 1399 57 1576 • PORCNTLE2 2282 2016 1840 336 1859 • PORCNWNT2B 2283 22 467 120 1931 • PORCNTCF7L1 2292 1382 731 1405 1475 • PORCNSENP2 2477 1934 1963 400 1275

## PORCN 2*^nd^* order interaction ranking at different durations

• PORCNPPP2CA 228 1809 2291 1052 • PORCNWNT2 231 1928 1576 2289 • PORCNSFRP4 236 1924 470 329 • PORCNSENP2 241 2197 1027 1160 • PORCNWNT4 246 937 1873 1945 • PORCNTCF7 261 1647 279 2440 • PORCNFBXW4 265 373 1258 1078 • PORCNTCF7L1 284 2273 1305 1648 • PORCNSFRP1 296 1810 1702 1045 • PORCNTLE1 299 2311 1860 1335 • PORCNWNT5A 318 1854 601 2286 • PORCNTLE2 372 793 1849 577 • PORCNPPP2R1A 402 1111 445 2329 • PORCNSLC9A3R1 499 1559 2411 1134 • PORCNWNT1 514 1393 2078 1575 • PORCNT 744 964 1422 714 • PORCNRHOU 880 1879 1631 1016 • PORCNWNT2B 974 2480 384 333 • PORCNWIF1 976 2152 1924 1272 • FZD5PORCN 1011 863 1856 2460 • PORCNPYGO1 1062 1176 1848 647 • PORCNWNT3A 1170 949 2304 2150 • PORCNWNT3 1467 1946 518 2227 • CCND1PORCN 1701 851 728 2297 • FSHBPORCN 1722 460 1310 1187 • CTNNBIP1PORCN 1778 2358 622 2244 • JUNPORCN 1794 2381 2372 108 • AESPORCN 1847 1778 2364 208 • BCL9PORCN 1854 1789 1657 2027 • CTBP1PORCN 1859 1801 2027 1718 • DIXDC1PORCN 1879 118 351 1937 • MYCPORCN 1935 1434 2449 1373 • FRAT1PORCN 1941 1078 2145 920 • LRP5PORCN 1984 290 2357 323 • CSNK1A1PORCN 2002 1439 1884 509 • CCND3PORCN 2040 1979 2259 2392 • CCND2PORCN 2064 1318 2132 690 • NKD1PORCN 2070 2477 2406 2400 • FGF4PORCN 2104 266 1462 1894 • CTBP2PORCN 2115 882 1946 1796 • DKK1PORCN 2125 220 2445 486 • FOXN1PORCN 2165 2015 1475 844 • EP300PORCN 2171 1935 2183 2344 • AXIN1PORCN 2181 820 872 2096 • DVL1PORCN 2195 1153 650 321 • NLKPORCN 2226 2206 2075 2396 • FZD6PORCN 2251 795 228 1960 • DAAM1PORCN 2254 1222 678 1813 • DVL2PORCN 2261 1163 882 2437 • LRP6PORCN 2265 296 1469 1667 • CSNK1DPORCN 2296 447 2386 1606 • FOSL1PORCN 2299 315 1672 1874 • KREMEN1PORCN 2300 1846 501 1861 • FRZBPORCN 2333 2401 1607 691 • PITX2PORCN 2342 19 371 2137 • FBXW11PORCN 2352 1091 2275 2364 • APCPORCN 2372 2366 960 2024 • CTNNB1PORCN 2377 1611 409 833 • CSNK1G1PORCN 2398 251 1978 1687 • FBXW2PORCN 2399 727 2452 2301 • CSNK2A1PORCN 2403 1104 302 2403 • FZD8PORCN 2405 143 394 2439 • GSK3BPORCN 2407 97 439 727 • GSK3APORCN 2433 354 628 470 • FZD1PORCN 2439 153 1710 497 • BTRCPORCN 2445 754 1482 949 • FZD2PORCN 2464 1204 276 1676 • CXXC4PORCN 2466 1418 2421 2241 • FZD7PORCN 2477 145 133 2132 • LEF1PORCN 2479 158 878 1778

## PYGO1 2*^nd^* order interaction ranking at different time points

• FRZBPYGO1 10 2291 243 346 113 • DKK1PYGO1 22 2018 1054 280 876 • FZD7PYGO1 54 1940 166 939 413 • FRAT1PYGO1 56 2336 339 783 1213 • NLKPYGO1 69 1980 1391 1064 2220 • CSNK2A1PYGO1 82 388 2066 1551 1693 • GSK3BPYGO1 91 972 292 2229 539 • FZD1PYGO1 95 56 308 2171 1220 • CSNK1DPYGO1 128 2403 908 2260 1612 • FBXW11PYGO1 149 224 840 631 2346 • FOSL1PYGO1 165 1492 182 453 654 • APCPYGO1 169 1188 303 604 717 • FZD5PYGO1 170 1711 454 1243 1406 • PPP2R1APYGO1 189 193 1877 193 1170 • DIXDC1PYGO1 191 1384 61 367 1511 • FZD6PYGO1 208 331 1966 1269 896 • LRP5PYGO1 213 1489 641 978 450 • FZD8PYGO1 224 1063 280 398 1663 • CCND3PYGO1 234 580 1064 391 1756 • PPP2CAPYGO1 271 1161 2001 1293 1221 • PITX2PYGO1 280 1085 418 1731 2315 • KREMEN1PYGO1 318 1547 1057 259 1123 • EP300PYGO1 327 1064 1005 676 1555 • FSHBPYGO1 330 200 1847 2263 682 • DAAM1PYGO1 334 733 953 296 1819 • FBXW2PYGO1 357 241 1492 1080 1715 • NKD1PYGO1 393 705 1346 39 618 • CXXC4PYGO1 423 2003 138 505 1851 • CTNNB1PYGO1 429 1059 524 1962 194 • CTBP1PYGO1 507 860 255 403 114 • LRP6PYGO1 521 455 1293 901 1927 • BTRCPYGO1 549 363 1493 10 1777 • CTBP2PYGO1 600 1633 626 1657 1210 • CSNK1G1PYGO1 631 2315 1941 828 2285 • AXIN1PYGO1 656 1745 1135 85 778 • DVL2PYGO1 673 23 1297 2193 647 • CTNNBIP1PYGO1 716 1114 358 920 34 • DVL1PYGO1 730 201 335 77 1131 • FZD2PYGO1 759 1301 384 1276 508 • CCND2PYGO1 776 1873 1534 1184 1529 • FGF4PYGO1 779 1407 78 1643 137 • JUNPYGO1 853 1183 845 1285 592 • LEF1PYGO1 873 383 565 1527 2132 • MYCPYGO1 919 1954 931 799 15 • PYGO1WNT1 975 1549 441 1906 1617 • PORCNPYGO1 990 272 700 673 566 • PYGO1SFRP1 1029 1279 2092 1230 572 • PYGO1WIF1 1064 911 1414 103 936 • BCL9PYGO1 1100 974 1065 2104 2475 • GSK3APYGO1 1121 411 1993 1052 1717 • CSNK1A1PYGO1 1145 366 1753 63 179 • AESPYGO1 1224 1765 903 2046 1015 • FOXN1PYGO1 1244 852 2028 2371 1309 • PYGO1WNT3 1526 262 1638 318 285 • CCND1PYGO1 1587 150 442 1044 1118 • PYGO1T 1850 571 1290 1560 355 • PYGO1TLE1 1852 840 2063 944 2119 • PYGO1SFRP4 1903 1513 1133 314 1975 • PYGO1WNT2 2085 771 1593 152 2205 • PYGO1SLC9A3R1 2095 643 2396 12 1534 • PYGO1FBXW4 2115 2139 842 947 1341 • PYGO1TCF7 2243 1746 1075 981 1059 • PYGO1TCF7L1 2249 1439 2258 1038 2323 • PYGO1TLE2 2266 597 2346 645 421 • PYGO1RHOU 2299 990 2051 1028 1784 • PYGO1WNT3A 2307 1074 1660 1025 883 • PYGO1WNT4 2315 418 1523 5 1307 • PYGO1WNT2B 2318 828 559 697 1076 • PYGO1WNT5A 2424 1394 663 1089 1428 • PYGO1SENP2 2485 2126 1227 94 1945

## PYGO1 2*^nd^* order interaction ranking at different durations

• PYGO1SENP2 5 2467 752 453 • PYGO1WNT5A 6 479 506 1847 • PYGO1FBXW4 14 245 1608 571 • PYGO1SFRP4 30 1398 569 24 • PYGO1WNT4 61 772 1971 2417 • PYGO1SFRP1 63 2345 2393 464 • PYGO1TCF7 90 656 157 1555 • PYGO1TCF7L1 93 1930 1059 228 • PYGO1TLE2 110 1446 2384 2164 • PYGO1WNT2 149 1948 1153 1710 • PYGO1SLC9A3R1 237 1075 1714 404 • PYGO1T 242 448 1724 603 • PYGO1TLE1 245 1445 1639 698 • PYGO1WIF1 316 1660 1319 428 • PYGO1WNT3A 474 1475 2256 1142 • PYGO1WNT2B 492 1709 1209 151 • PYGO1WNT1 661 357 2102 2333 • PYGO1RHOU 667 1337 2407 1932 • FZD5PYGO1 925 1205 2156 2193 • PYGO1WNT3 962 1103 890 899 • PORCNPYGO1 1062 1176 1848 647 • CSNK1A1PYGO1 1363 2212 1835 2401 • DIXDC1PYGO1 1546 165 1164 388 • AESPYGO1 1755 2062 2278 2087 • CTNNBIP1PYGO1 1855 2368 1443 1085 • CTNNB1PYGO1 1891 1824 2280 2292 • CTBP1PYGO1 1927 1984 1268 1537 • FSHBPYGO1 2021 660 2416 468 • FOSL1PYGO1 2038 276 2170 1620 • CCND1PYGO1 2046 1240 1667 1141 • CXXC4PYGO1 2136 1776 1723 1077 • APCPYGO1 2156 1482 2439 1865 • FOXN1PYGO1 2203 1871 1693 663 • JUNPYGO1 2206 2222 2134 466 • CSNK1DPYGO1 2208 397 2398 2140 • MYCPYGO1 2209 735 2319 2377 • AXIN1PYGO1 2211 1478 1653 327 • EP300PYGO1 2235 1919 456 1430 • DVL2PYGO1 2255 2464 1397 2051 • PPP2R1APYGO1 2260 1339 1970 1154 • CCND2PYGO1 2271 2048 2103 766 • FZD8PYGO1 2278 115 293 1546 • NKD1PYGO1 2298 2406 1967 593 • NLKPYGO1 2319 2175 1236 1792 • FBXW2PYGO1 2327 2363 2236 1679 • KREMEN1PYGO1 2334 1813 1635 2043 • GSK3BPYGO1 2347 148 455 2250 • CTBP2PYGO1 2350 634 2479 2214 • DKK1PYGO1 2374 346 2144 2235 • LRP6PYGO1 2384 481 1190 1226 • BCL9PYGO1 2387 2476 2413 1243 • FGF4PYGO1 2389 282 914 1159 • FRAT1PYGO1 2404 1325 2334 783 • LEF1PYGO1 2409 217 1279 1343 • FZD7PYGO1 2412 213 278 1057 • FBXW11PYGO1 2417 1539 1074 2283 • CCND3PYGO1 2418 1961 857 1762 • CSNK1G1PYGO1 2420 103 1683 2461 • FRZBPYGO1 2435 813 2123 1220 • BTRCPYGO1 2437 1081 1690 898 • LRP5PYGO1 2441 361 774 902 • FZD1PYGO1 2450 181 2133 1563 • FZD6PYGO1 2452 1028 269 591 • DVL1PYGO1 2455 2145 1035 747 • DAAM1PYGO1 2461 883 446 854 • GSK3APYGO1 2463 486 640 686 • PPP2CAPYGO1 2467 832 2064 1949 • CSNK2A1PYGO1 2470 2316 463 1258 • FZD2PYGO1 2476 1027 608 1467 • PITX2PYGO1 2485 144 369 2304

## RHOU 2*^nd^* order interaction ranking at different time points

• RHOUWNT1 89 2193 151 2203 693 • RHOUSFRP1 108 1587 1788 698 527 • FRZBRHOU 141 1844 896 772 1079 • RHOUT 287 1334 899 2048 2164 • RHOUWIF1 304 2039 1766 234 1941 • RHOUWNT3 343 1376 1052 341 1189 • DKK1RHOU 491 2399 1489 1385 1950 • RHOUFBXW4 502 2333 498 815 2068 • RHOUSFRP4 579 855 1825 1380 1539 • RHOUTLE1 640 2299 2397 773 2325 • FOSL1RHOU 780 1043 593 974 561 • RHOUSLC9A3R1 825 1362 1429 83 908 • RHOUWNT2 843 1535 1661 153 687 • RHOUWNT5A 886 2070 531 2311 1579 • RHOUWNT4 895 1168 2265 287 1482 • FZD7RHOU 965 287 1405 326 663 • RHOUTCF7 1002 1326 359 603 1896 • RHOUWNT3A 1010 288 2335 2221 1452 • FZD1RHOU 1071 513 941 2285 564 • RHOUWNT2B 1082 453 293 150 1678 • APCRHOU 1106 832 1098 1831 709 • FBXW11RHOU 1110 378 2144 1995 2294 • CSNK2A1RHOU 1187 1542 1484 2443 819 • RHOUTLE2 1199 1280 1606 368 927 • FZD5RHOU 1315 1871 1474 706 1373 • DIXDC1RHOU 1326 1919 221 1212 1989 • RHOUTCF7L1 1329 2234 788 43 2428 • FRAT1RHOU 1334 1609 2104 1022 361 • LRP5RHOU 1370 1239 1337 2381 2137 • CSNK1DRHOU 1379 2235 2319 1766 1814 • GSK3BRHOU 1441 1854 551 1689 1121 • KREMEN1RHOU 1452 606 1888 38 2286 • DAAM1RHOU 1455 971 1632 98 2281 • CXXC4RHOU 1459 500 1457 454 2435 • PPP2CARHOU 1488 2115 1869 1130 1284 • FBXW2RHOU 1501 921 1894 310 1049 • EP300RHOU 1524 598 1852 65 831 • FZD8RHOU 1542 698 1544 2185 2381 • FZD6RHOU 1550 1300 1958 656 160 • CCND3RHOU 1625 1030 1821 2011 426 • PITX2RHOU 1650 1305 1939 2002 2246 • NLKRHOU 1661 350 1174 129 2152 • PPP2R1ARHOU 1701 623 1843 1576 1644 • FSHBRHOU 1724 1623 2177 2170 391 • NKD1RHOU 1749 525 2089 462 2212 • CTNNB1RHOU 1791 2316 1141 1121 1345 • LRP6RHOU 1797 2253 2218 1375 1549 • CTBP2RHOU 1810 1465 1107 984 793 • CTNNBIP1RHOU 1860 2027 1478 654 201 • CTBP1RHOU 1889 2017 895 423 1357 • DVL2RHOU 1955 615 2364 2454 767 • CSNK1G1RHOU 2000 1041 2212 531 835 • DVL1RHOU 2047 1888 1035 237 929 • BTRCRHOU 2089 1976 1482 664 1657 • FZD2RHOU 2092 1155 1763 1043 795 • AXIN1RHOU 2133 1729 1249 211 979 • CCND2RHOU 2146 372 2299 595 2410 • PORCNRHOU 2221 1872 2117 1541 1719 • AESRHOU 2236 2208 2249 1240 1399 • RHOUSENP2 2251 1884 1515 107 1310 • CSNK1A1RHOU 2254 2227 778 501 430 • MYCRHOU 2263 499 1730 1050 667 • FGF4RHOU 2268 1679 1296 353 1532 • PYGO1RHOU 2299 990 2051 1028 1784 • LEF1RHOU 2378 1632 1077 1117 1970 • FOXN1RHOU 2392 640 1837 2444 1791 • BCL9RHOU 2395 2416 2354 1826 1546 • JUNRHOU 2403 670 2171 1447 224 • CCND1RHOU 2431 853 993 777 1300 • GSK3ARHOU 2449 2162 2332 1308 2417

## RHOU 2*^nd^* order interaction ranking at different durations

• RHOUSENP2 20 1349 145 25 • RHOUSFRP4 27 1514 113 56 • RHOUTLE2 37 538 1122 2277 • RHOUTLE1 56 914 982 28 • RHOUWIF1 102 2046 487 15 • RHOUSFRP1 134 2054 770 57 • RHOUTCF7L1 135 1328 163 836 • RHOUWNT5A 152 573 590 607 • RHOUTCF7 175 410 31 784 • RHOUWNT4 197 291 556 561 • RHOUWNT2 225 523 975 192 • RHOUFBXW4 355 331 1480 559 • PYGO1RHOU 667 1337 2407 1932 • RHOUWNT2B 702 956 366 70 • RHOUSLC9A3R1 713 803 1985 105 • RHOUWNT1 793 292 383 1153 • PORCNRHOU 880 1879 1631 1016 • RHOUT 892 445 284 1713 • CSNK1A1RHOU 979 1008 1539 2275 • RHOUWNT3A 986 1501 2264 632 • RHOUWNT3 998 2240 91 177 • FZD5RHOU 1067 840 1063 992 • CTNNB1RHOU 1769 1584 2446 1961 • AXIN1RHOU 1788 2233 2296 87 • CTBP1RHOU 1797 2093 1390 706 • CCND2RHOU 1813 2023 1827 790 • DIXDC1RHOU 1823 300 2424 439 • CCND1RHOU 1844 2161 2467 332 • KREMEN1RHOU 1852 1538 1336 1228 • JUNRHOU 1882 1949 2233 2478 • BCL9RHOU 1892 1855 1852 654 • CTNNBIP1RHOU 1939 1997 1947 221 • LRP5RHOU 2003 494 1829 1930 • AESRHOU 2048 2021 2090 2177 • DVL2RHOU 2049 2079 1322 386 • FSHBRHOU 2053 667 2271 454 • MYCRHOU 2062 693 900 450 • NKD1RHOU 2083 1181 1404 395 • GSK3ARHOU 2085 867 1814 710 • FRAT1RHOU 2091 1208 2157 1782 • CSNK1DRHOU 2108 1237 1711 1809 • EP300RHOU 2113 1136 1989 190 • GSK3BRHOU 2126 356 570 673 • NLKRHOU 2131 2400 865 432 • LEF1RHOU 2146 590 2321 983 • FBXW2RHOU 2151 801 1957 760 • CCND3RHOU 2177 1164 2003 682 • DVL1RHOU 2214 997 1411 188 • CSNK1G1RHOU 2229 484 2001 463 • DKK1RHOU 2239 848 1160 1332 • FZD2RHOU 2279 1266 2294 993 • CSNK2A1RHOU 2292 2471 669 806 • FZD6RHOU 2315 1635 1162 273 • FZD8RHOU 2322 401 2213 2115 • PPP2R1ARHOU 2324 1615 1498 320 • APCRHOU 2362 1347 564 425 • CXXC4RHOU 2368 2239 763 565 • FRZBRHOU 2390 1218 1650 600 • LRP6RHOU 2397 737 2044 592 • FOXN1RHOU 2415 1159 1718 847 • CTBP2RHOU 2427 1791 2246 1453 • DAAM1RHOU 2436 1425 2098 143 • FZD1RHOU 2438 464 1890 1891 • FBXW11RHOU 2442 1489 1234 674 • FOSL1RHOU 2447 646 2289 1585 • PITX2RHOU 2448 293 849 2195 • FZD7RHOU 2449 569 657 122 • BTRCRHOU 2462 2395 1919 1240 • PPP2CARHOU 2468 1536 1240 2094 • FGF4RHOU 2480 587 2109 726

## SENP2 2*^nd^* order interaction ranking at different time points

• SENP2WNT1 1 887 42 16 407 • SENP2SFRP1 2 844 564 1691 202 • SENP2WNT3 3 461 861 992 751 • SENP2T 7 1140 116 126 1083 • SENP2WIF1 8 1471 547 2184 2070 • SENP2TLE1 14 105 1586 1853 339 • SENP2SFRP4 19 2375 620 1713 2313 • SENP2SLC9A3R1 25 679 1109 786 679 • SENP2WNT2B 32 848 45 572 529 • SENP2FBXW4 34 308 225 1062 1175 • SENP2WNT3A 38 546 1146 583 964 • SENP2WNT5A 40 203 97 630 1124 • SENP2WNT4 45 830 492 2308 2211 • SENP2TCF7 57 1107 117 945 472 • SENP2TLE2 65 1060 407 2039 1987 • SENP2WNT2 86 1136 863 824 2485 • SENP2TCF7L1 134 885 1070 292 1861 • FRZBSENP2 1525 2475 1121 858 452 • DKK1SENP2 1758 2446 1996 577 2451 • RHOUSENP2 2251 1884 1515 107 1310 • FOSL1SENP2 2338 2209 1138 1400 122 • DAAM1SENP2 2353 1249 1314 1723 982 • FZD7SENP2 2354 883 2216 1132 1032 • FZD1SENP2 2357 266 1235 165 451 • DIXDC1SENP2 2366 2012 651 1409 2361 • APCSENP2 2371 2432 655 1128 287 • LRP5SENP2 2377 2301 2463 959 1894 • FBXW2SENP2 2382 1297 1854 205 155 • FBXW11SENP2 2387 1355 1933 60 308 • EP300SENP2 2394 680 1959 1181 1134 • CXXC4SENP2 2405 2451 2187 1406 1948 • FZD5SENP2 2406 1420 1009 1642 25 • CSNK2A1SENP2 2409 420 1987 704 367 • PPP2CASENP2 2411 1750 2087 1097 369 • CSNK1DSENP2 2420 2222 1751 600 1138 • GSK3BSENP2 2422 1441 1885 42 991 • FRAT1SENP2 2435 2342 1501 1489 354 • FZD6SENP2 2439 2010 1213 2258 1102 • FSHBSENP2 2441 1138 1662 132 832 • NLKSENP2 2442 38 1565 942 442 • CTNNBIP1SENP2 2443 2388 1842 1635 57 • CCND3SENP2 2444 216 2181 644 858 • PPP2R1ASENP2 2446 2262 1926 696 2266 • PITX2SENP2 2447 1815 2130 162 1773 • KREMEN1SENP2 2448 1039 1502 355 1364 • CSNK1G1SENP2 2450 2164 1406 147 1311 • CTNNB1SENP2 2454 561 2308 413 164 • NKD1SENP2 2456 132 1666 1870 1354 • FZD8SENP2 2458 2417 1971 137 880 • DVL1SENP2 2463 1087 1092 756 1560 • CTBP1SENP2 2465 1852 1205 1526 180 • BCL9SENP2 2466 1992 1408 907 1387 • DVL2SENP2 2467 502 1920 51 998 • FZD2SENP2 2468 1803 2109 27 483 • CTBP2SENP2 2469 256 2301 447 419 • FGF4SENP2 2470 1975 1237 1718 1939 • CCND1SENP2 2471 555 1824 53 1640 • LRP6SENP2 2472 2268 877 4 659 • FOXN1SENP2 2473 427 1964 197 1578 • CCND2SENP2 2474 1527 1866 597 1670 • BTRCSENP2 2476 1848 2112 49 1484 • PORCNSENP2 2477 1934 1963 400 1275 • AXIN1SENP2 2478 1724 2427 565 815 • LEF1SENP2 2479 2006 2172 188 1682 • MYCSENP2 2480 1021 2246 366 302 • GSK3ASENP2 2481 1798 1323 250 2261 • CSNK1A1SENP2 2482 989 2158 994 247 • AESSENP2 2483 115 2078 269 1239 • JUNSENP2 2484 1385 2300 1590 364 • PYGO1SENP2 2485 2126 1227 94 1945

## SENP2 2*^nd^* order interaction ranking at different durations

• CSNK1A1SENP2 1 1971 2108 98 • PYGO1SENP2 5 2467 752 453 • FZD5SENP2 7 1110 2167 1402 • RHOUSENP2 20 1349 145 25 • CTBP1SENP2 21 2029 2136 101 • JUNSENP2 24 2466 2351 157 • DIXDC1SENP2 25 475 76 1700 • FRAT1SENP2 31 2132 1509 118 • CTNNB1SENP2 32 1344 234 546 • AESSENP2 33 954 282 46 • CSNK1DSENP2 36 1952 1363 133 • FOXN1SENP2 48 2041 316 73 • AXIN1SENP2 50 1039 321 2172 • KREMEN1SENP2 51 2011 249 189 • BCL9SENP2 66 1598 1024 1668 • CCND1SENP2 72 2154 95 1956 • CCND2SENP2 76 1613 649 1811 • CTNNBIP1SENP2 78 2396 74 624 • FSHBSENP2 79 625 458 1122 • BTRCSENP2 82 1713 440 236 • MYCSENP2 86 1352 1892 1280 • APCSENP2 87 1074 2100 521 • DVL1SENP2 98 1004 94 2405 • GSK3ASENP2 99 968 15 1795 • CTBP2SENP2 108 1288 2071 1540 • FOSL1SENP2 114 1139 1632 1073 • NKD1SENP2 159 1517 1737 1798 • EP300SENP2 163 1732 548 1360 • DVL2SENP2 167 2190 236 570 • FRZBSENP2 200 2104 925 50 • LRP5SENP2 206 917 1530 167 • PORCNSENP2 241 2197 1027 1160 • CCND3SENP2 258 1754 684 793 • DKK1SENP2 267 886 1031 959 • CXXC4SENP2 343 2045 1629 1507 • DAAM1SENP2 411 1227 52 1922 • PPP2CASENP2 413 2453 988 152 • NLKSENP2 420 1550 656 1008 • GSK3BSENP2 457 591 105 58 • FZD8SENP2 471 379 60 287 • FZD6SENP2 489 2405 11 807 • PITX2SENP2 531 322 41 262 • FBXW2SENP2 550 2336 993 337 • FZD1SENP2 584 432 438 184 • FBXW11SENP2 606 1861 1887 996 • CSNK2A1SENP2 627 2257 51 508 • FZD2SENP2 695 517 197 542 • FZD7SENP2 711 1423 8 978 • LEF1SENP2 738 531 223 849 • FGF4SENP2 745 419 441 1517 • LRP6SENP2 761 328 970 961 • PPP2R1ASENP2 779 2450 402 1310 • CSNK1G1SENP2 837 911 393 1058 • SENP2SFRP4 1707 1833 1070 1264 • SENP2WNT5A 2000 483 1550 1943 • SENP2SFRP1 2056 2472 1883 808 • SENP2TCF7L1 2082 2483 1906 1734 • SENP2TLE1 2275 1096 812 2033 • SENP2FBXW4 2297 269 1705 1913 • SENP2TLE2 2303 607 1471 121 • SENP2T 2326 586 1865 225 • SENP2WIF1 2329 664 1897 2175 • SENP2TCF7 2370 567 904 1238 • SENP2WNT1 2393 505 2173 361 • SENP2WNT2 2411 562 1345 1102 • SENP2WNT4 2426 347 1616 585 • SENP2WNT3 2454 2123 2420 1444 • SENP2WNT3A 2456 1372 1273 693 • SENP2WNT2B 2469 857 2068 2393 • SENP2SLC9A3R1 2472 576 1199 981

## T 2*^nd^* order interaction ranking at different time points

• SENP2T 7 1140 116 126 1083 • DKK1T 27 2275 2174 727 1376 • FRZBT 49 2398 263 900 1130 • APCT 248 1850 282 2337 322 • FZD1T 264 458 883 863 789 • RHOUT 287 1334 899 2048 2164 • FOSL1T 292 1253 266 1226 1235 • PPP2R1AT 314 58 789 885 1795 • FZD5T 320 159 683 1644 1082 • CSNK1DT 329 1374 2278 1696 1108 • FBXW11T 336 550 2163 1752 1999 • FRAT1T 340 2150 283 1220 2282 • LRP5T 371 2049 1900 1049 556 • FZD7T 394 1162 457 1879 1750 • SLC9A3R1T 414 169 381 548 2215 • DIXDC1T 459 1520 73 1648 1070 • GSK3BT 473 1531 762 602 2452 • NLKT 479 2119 2267 846 711 • CSNK2A1T 483 2129 2005 1113 493 • EP300T 484 440 1034 1699 1603 • FBXW2T 485 228 1903 1163 901 • DAAM1T 517 512 409 1858 995 • CCND3T 531 564 890 2073 726 • FZD8T 555 719 427 1119 1692 • CXXC4T 561 1994 610 708 50 • SFRP4T 565 2458 390 848 548 • FBXW4T 592 622 681 1260 1597 • FZD6T 598 92 1558 1495 2392 • KREMEN1T 628 1932 1056 1736 2056 • TWNT3 635 2026 2044 2135 40 • CTBP1T 667 1524 944 983 2233 • PPP2CAT 721 1128 2008 2287 422 • PITX2T 737 1820 205 2189 2468 • TWIF1 766 2327 633 1811 1458 • TWNT1 797 2080 708 1595 5 • FSHBT 803 1941 1461 719 1372 • CTBP2T 1139 2293 192 2125 634 • CTNNBIP1T 1155 2009 520 442 301 • DVL2T 1159 735 853 953 692 • CTNNB1T 1215 1772 1535 1053 1487 • BCL9T 1225 758 836 232 1540 • AXIN1T 1256 1174 1016 771 1243 • NKD1T 1269 385 1198 964 1656 • LRP6T 1287 1571 1875 1821 1115 • CSNK1G1T 1289 1771 1365 1872 593 • BTRCT 1341 2284 2437 503 2202 • AEST 1369 532 824 2252 2250 • FGF4T 1385 1782 93 2411 1202 • DVL1T 1454 1548 643 1283 1666 • TTLE1 1481 2277 2392 218 544 • PORCNT 1509 44 945 2247 1353 • CSNK1A1T 1518 320 1834 389 633 • LEF1T 1533 875 978 898 1505 • TWNT2 1566 109 2442 70 2333 • CCND2T 1581 709 1549 1760 1445 • MYCT 1608 1185 1617 552 919 • TWNT4 1829 1967 1533 1012 1377 • PYGO1T 1850 571 1290 1560 355 • JUNT 1873 939 596 371 1249 • SFRP1T 1879 1964 248 9 2027 • FZD2T 1890 909 515 2050 206 • FOXN1T 1902 2359 1217 1438 2142 • TWNT3A 1938 581 2369 1829 1470 • CCND1T 1967 1789 803 2272 1547 • TWNT5A 2015 1222 1712 854 444 • TWNT2B 2025 790 735 1578 2192 • TTCF7 2086 1882 864 1982 790 • TTCF7L1 2153 2108 1509 2317 651 • TTLE2 2205 1942 1665 1354 1853 • GSK3AT 2223 254 2114 2377 2411

## T 2*^nd^* order interaction ranking at different durations

• PYGO1T 242 448 1724 603 • TWNT5A 292 957 2456 501 • TTCF7L1 328 1729 1286 821 • TTLE2 491 1146 2097 527 • TTLE1 533 1098 1413 199 • TTCF7 590 827 78 203 • TWNT1 722 581 2112 562 • TWNT4 809 641 2074 1472 • CSNK1A1T 812 2469 2182 2091 • CTNNB1T 850 2080 1911 1179 • RHOUT 892 445 284 1713 • JUNT 909 1717 2410 1229 • TWIF1 921 2371 1265 3 • CCND2T 956 1974 2308 136 • DIXDC1T 1018 61 515 451 • TWNT2 1089 1037 1225 120 • FZD5T 1182 616 1738 1523 • CTNNBIP1T 1233 1463 688 282 • CTBP1T 1264 1456 2210 1278 • FOXN1T 1278 1082 1368 2372 • AEST 1306 2417 1843 1086 • KREMEN1T 1358 2301 322 2332 • AXIN1T 1517 1148 681 390 • FSHBT 1597 558 1186 398 • GSK3AT 1603 578 468 293 • BTRCT 1613 1359 2347 532 • FRAT1T 1614 1042 1569 749 • CCND1T 1634 563 356 206 • MYCT 1660 510 1992 218 • NKD1T 1687 2291 1591 489 • DVL1T 1710 1070 229 259 • CTBP2T 1713 1458 1833 512 • DVL2T 1820 1415 881 786 • SLC9A3R1T 1841 544 576 253 • TWNT3 1851 1363 435 2 • FOSL1T 1888 453 2471 1015 • LRP5T 1890 224 1788 1439 • APCT 1906 2077 1746 1137 • EP300T 1931 1395 1274 705 • CSNK1DT 1963 497 1424 297 • FBXW2T 1968 2460 1685 522 • CCND3T 1983 1811 738 764 • BCL9T 1994 1695 1296 241 • FBXW4T 2006 1805 488 354 • GSK3BT 2080 111 191 573 • FRZBT 2143 1291 1563 1583 • PITX2T 2144 44 201 1189 • FBXW11T 2148 1905 2218 1737 • FZD6T 2149 744 156 62 • NLKT 2205 718 1324 739 • PPP2R1AT 2237 1568 838 1631 • PPP2CAT 2241 1124 721 244 • CSNK1G1T 2288 319 2302 887 • FZD2T 2294 477 123 1046 • CXXC4T 2307 1603 723 461 • FZD1T 2320 253 1374 1594 • SENP2T 2326 586 1865 225 • FZD8T 2328 82 73 754 • LRP6T 2336 156 2030 168 • SFRP4T 2340 203 2380 139 • DKK1T 2344 289 2287 810 • SFRP1T 2376 100 794 325 • CSNK2A1T 2392 1226 142 1521 • LEF1T 2429 198 642 277 • FZD7T 2446 288 32 717 • FGF4T 2460 278 623 400

## WIF1 2*^nd^* order interaction ranking at different time points

• SENP2WIF1 8 1471 547 2184 2070 • FRZBWIF1 18 2483 218 1864 608 • DKK1WIF1 21 2425 2359 119 2159 • TCF7L1WIF1 60 566 2194 668 1176 • FZD7WIF1 98 2037 1274 231 2303 • FBXW11WIF1 117 510 1930 350 1724 • APCWIF1 148 2463 1119 1778 550 • FZD5WIF1 154 1984 1110 2468 853 • CSNK2A1WIF1 177 1227 2281 1676 714 • TLE2WIF1 199 2339 1706 2150 1057 • GSK3BWIF1 211 1840 1988 260 1125 • FZD1WIF1 215 1929 2149 1822 1338 • FRAT1WIF1 221 2152 2344 184 1269 • CSNK1DWIF1 257 2461 2352 910 2094 • DAAM1WIF1 265 1637 1578 1840 2077 • SLC9A3R1WIF1 268 1257 1494 538 1638 • NLKWIF1 277 968 2007 343 2446 • FZD8WIF1 278 1843 1623 370 1266 • FZD6WIF1 284 2285 2173 1157 1072 • DIXDC1WIF1 290 2332 572 1567 2040 • PPP2R1AWIF1 291 1990 1823 1155 2341 • FOSL1WIF1 295 2329 1693 2425 251 • RHOUWIF1 304 2039 1766 234 1941 • PPP2CAWIF1 310 573 1845 2408 2336 • TCF7WIF1 324 1432 2083 2255 1812 • CXXC4WIF1 335 2147 2285 1933 2048 • KREMEN1WIF1 358 1938 1317 532 939 • LRP5WIF1 363 2440 1312 682 2463 • SFRP4WIF1 386 2480 968 1186 1410 • CCND3WIF1 403 570 2101 620 1854 • CTBP1WIF1 436 2350 1924 342 810 • CTNNB1WIF1 446 1651 1633 1664 1209 • FSHBWIF1 460 2261 2071 762 1098 • TLE1WIF1 476 2001 1609 1519 1740 • EP300WIF1 482 1192 1357 1229 1437 • FBXW4WIF1 493 215 1748 1341 553 • PITX2WIF1 543 1595 2146 194 1906 • CTNNBIP1WIF1 557 2419 2287 1606 392 • DVL2WIF1 619 408 892 308 372 • FBXW2WIF1 625 1013 1649 2248 2208 • NKD1WIF1 650 611 1688 2151 1075 • CSNK1G1WIF1 702 2409 2030 877 1348 • PORCNWIF1 741 1950 2261 624 1700 • TWIF1 766 2327 633 1811 1458 • LRP6WIF1 816 2266 2372 790 1515 • CTBP2WIF1 830 1880 1101 705 2053 • AESWIF1 844 523 2048 324 1278 • AXIN1WIF1 854 2360 2326 1242 11 • DVL1WIF1 876 1034 1444 1753 990 • BTRCWIF1 923 601 2120 417 1444 • WIF1WNT3 944 370 959 34 1056 • WIF1WNT1 985 1244 36 149 6 • CCND2WIF1 1027 1657 2400 2066 1084 • PYGO1WIF1 1064 911 1414 103 936 • FGF4WIF1 1080 1957 1480 1877 2082 • LEF1WIF1 1103 2057 1524 1042 2237 • FOXN1WIF1 1171 891 2010 160 534 • BCL9WIF1 1177 1985 1835 135 1691 • SFRP1WIF1 1358 1877 1241 2388 2158 • CCND1WIF1 1380 626 932 919 215 • CSNK1A1WIF1 1420 1702 2263 1700 263 • FZD2WIF1 1466 1586 1526 825 1658 • MYCWIF1 1504 2321 1883 977 1349 • JUNWIF1 1637 1650 2022 2049 1268 • GSK3AWIF1 1693 1515 2204 2342 1317 • WIF1WNT2B 1776 433 31 619 1024 • WIF1WNT2 1838 1688 885 2482 1930 • WIF1WNT3A 2154 1574 1083 263 1350 • WIF1WNT4 2157 1267 507 2276 2272 • WIF1WNT5A 2285 2175 271 1233 1649

## WIF1 2*^nd^* order interaction ranking at different durations

• RHOUWIF1 102 2046 487 15 • CSNK1A1WIF1 153 1118 1172 20 • PYGO1WIF1 316 1660 1319 428 • WIF1WNT5A 562 241 1836 2013 • FZD5WIF1 601 2106 2092 889 • CTBP1WIF1 694 1806 2029 871 • DIXDC1WIF1 832 595 629 1959 • TWIF1 921 2371 1265 3 • CCND2WIF1 931 1671 2434 2161 • JUNWIF1 937 2168 2368 37 • WIF1WNT4 975 418 1233 1469 • PORCNWIF1 976 2152 1924 1272 • FRAT1WIF1 1017 2481 1955 213 • CTNNB1WIF1 1061 1113 1701 363 • MYCWIF1 1136 2296 2245 547 • KREMEN1WIF1 1179 2437 1042 97 • FOXN1WIF1 1208 855 1307 130 • AESWIF1 1230 1537 1556 6 • FSHBWIF1 1251 930 1156 1155 • AXIN1WIF1 1257 1815 754 1391 • BCL9WIF1 1303 1686 1818 462 • GSK3AWIF1 1328 831 457 2307 • CSNK1DWIF1 1339 817 1007 109 • DVL1WIF1 1343 2150 1183 587 • CCND1WIF1 1359 1632 454 905 • CTNNBIP1WIF1 1387 1689 505 732 • NKD1WIF1 1397 1264 1665 223 • BTRCWIF1 1417 2121 2431 474 • CTBP2WIF1 1510 2164 2440 1125 • WIF1WNT1 1573 424 1806 349 • FOSL1WIF1 1592 467 2318 169 • SLC9A3R1WIF1 1606 1331 910 2184 • EP300WIF1 1609 1133 2107 1247 • WIF1WNT2 1619 462 1463 1709 • APCWIF1 1624 1777 1882 1868 • LRP5WIF1 1671 790 2032 125 • FRZBWIF1 1770 1839 1974 147 • TLE1WIF1 1771 1069 1811 1906 • FBXW4WIF1 1781 1421 605 2069 • DKK1WIF1 1865 969 2292 1206 • DVL2WIF1 1881 1989 1640 2207 • TCF7WIF1 1900 1887 2186 1009 • CCND3WIF1 1913 1736 1204 1182 • PITX2WIF1 1928 561 493 79 • LEF1WIF1 1937 335 1663 1783 • CXXC4WIF1 1938 1207 1651 2058 • WIF1WNT3A 1955 927 2262 426 • PPP2CAWIF1 1958 1618 954 711 • FBXW2WIF1 1970 1707 2115 1845 • GSK3BWIF1 1992 526 426 77 • FZD2WIF1 2012 1862 869 310 • FBXW11WIF1 2019 1290 2160 954 • FZD1WIF1 2033 685 1579 319 • FZD6WIF1 2051 1250 216 1136 • TLE2WIF1 2071 2322 1203 13 • FZD8WIF1 2076 733 271 1410 • PPP2R1AWIF1 2117 1735 1784 1514 • WIF1WNT2B 2123 489 720 1817 • CSNK2A1WIF1 2147 2010 315 1225 • FGF4WIF1 2178 866 1224 1609 • CSNK1G1WIF1 2189 658 2418 1298 • NLKWIF1 2202 2033 1648 1177 • WIF1WNT3 2204 1548 246 792 • TCF7L1WIF1 2262 984 1673 539 • LRP6WIF1 2290 1197 2396 1412 • DAAM1WIF1 2317 2119 607 2458 • SENP2WIF1 2329 664 1897 2175 • SFRP1WIF1 2346 701 2184 2375 • SFRP4WIF1 2406 635 2164 1775 • FZD7WIF1 2421 533 93 578

